# Assessment of the residential Finnish wolf population combines DNA captures, citizen observations and mortality data using a Bayesian state-space model

**DOI:** 10.1101/2021.12.21.473527

**Authors:** Samu Mäntyniemi, Inari Helle, Ilpo Kojola

## Abstract

Assessment of the Finnish wolf population relies on multiple sources of information. This paper describes how Bayesian inference is used to pool the information contained in different data sets (point observations, non-invasive genetics, known mortalities) for the estimation of the number of territories occupied by family packs and pairs. The output of the assessment model is a joint probability distribution, which describes current knowledge about the number of wolves within each territory. The joint distribution can be used to derive probability distributions for the total number of wolves in all territories and for the pack status within each territory. Most of the data set comprises of both voluntary-provided point observations and DNA samples provided by volunteers and research personnel. The new method reduces the role of expert judgement in the assessment process, providing increased transparency and repeatability.

## 1 Introduction

An important prerequisite for effective population management is reliable population monitoring, because population counts are imperative for several activities like assessing the conservation status of a species and setting hunting quotas. However, large carnivores pose a challenge for population monitoring, because they typically inhabit large remote areas at low densities (Herfindal et al, 2005; Kindberg et al, 2011; Mattisson et al, 2013), which is why species observations often accumulate unevenly and with varying precision. Voluntary- provided data are often used in monitoring of large carnivore populations (Kindberg et al, 2011; Bragina et al, 2015; Cretois et al, 2020). Thus, any population monitoring scheme focusing on large carnivores needs to cope with varying levels of uncertainty.

The wolf (*Canis lupus*) has experienced major population collapses in its native range in Europe and North America due to human activities but is now recolonising many areas (Chapron et al, 2014; Ripple et al, 2014). This is the case also in Finland, where the wolf population was hunted down from approximately 1 000 individuals to only some dozens in the second half of the 1800s (Mykrä et al, 2017). In the 1990s, the wolf re-established a permanent population. The population has been increasing since 2017 being 32-38 packs (90% probability interval,PI) and 18-25 pairs (90% PI) in March 2021 (Heikkinen et al, 2021).

The recolonization has resulted in conflicts, as the wolf causes damages to domestic animals, attacks hunting dogs and evokes fear in people (Marucco and Boitani, 2012; Flykt et al, 2013; Johansson et al, 2016; Olson et al, 2019; Tikkunen and Kojola, 2020; Bassi et al, 2021). Damage from wolves often generates displeasure and frustration and may fuel the illegal killing of wolves (Liberg et al, 2012; Pohja-Mykrä and Kurki, 2014; Suutarinen and Kojola, 2017; Liberg et al, 2020; Nowak et al, 2021). In Finland, the wolf is classified as endangered species (Liukko et al, 2019). Furthermore, outside the reindeer husbandry area, it is included in Annex IV of the EU’s Habitats Directive, which requires strict protection of the species. However, the wolf is also a game species in Finland, and the latest management hunting season was implemented in 2015/2016 (Ministry of Agriculture and Forestry, 2019).

Given the complex situation and conflicting aims and views present in the society, it is evident that there is a strong need for reliable monitoring of the wolf population in Finland. Today the monitoring is based on volunteer-provided point observations, non-invasive DNA samples collected by volunteers and wildlife professionals, information acquired from GPS-collared animals, and knowledge on known annual mortality (Kojola et al, 2018). Until recently, these data were used by experts to delineate the wolf territories, to estimate the number of individuals per territory, and by using the proportion of non-residents reported in the literature (Fuller et al, 2003)), to produce the final population estimate (Kojola et al, 2018). However, as this process is heavily expert-driven, the need for a method that integrates available knowledge in a more transparent and repeatable manner while explicitly expressing uncertainty has been recognized.

Here we describe a new method for the estimation of the annual size of the Finnish wolf population. The method is based on Bayesian inference, which is well suited to the question at hand, as it enables efficient integration of multiple data sources and handles uncertainty explicitly. In recent years Bayesian modelling has been applied to estimate wolf populations based on, for instance, tracking surveys, sign surveys, howling sessions and multistate hierarchical site occupancy models (Jiménez et al, 2016; Stauffer et al, 2021), spatial DNA capture-recapture (Bischof et al, 2020; López-Bao et al, 2018), and on an individual-based model which uses the number of packs, reproductions and pairs as data (Chapron et al, 2016). In our approach the starting point for the analysis is a set of wolf territories, which are delineated by experts every year as described by Kojola et al (2018). Our aim is to answer the question: Based on data accumulated within each of these wolf territories, what is the pack status and number of wolves per each territory, and what is the total number of wolves occupying these territories in Finland, given the associated uncertainties?

## 2 Material and methods

The Finnish wolf population assessment has two phases. In the first phase potential territory areas are estimated by a panel of experts based on all the available information (clusters of point observations of packs and twosomes, DNA samples, and GPS-locations of collared wolves; Kojola et al (2018)). In the second phase territory specific data are used to infer the number of wolves occupying each territory using the Bayesian state-space model, which we describe in this section (Fig 1). We illustrate the model by assessing the number of pairs, packs and total number of wolves in Finnish wolf territories in March 2020.

**Fig. 1.**
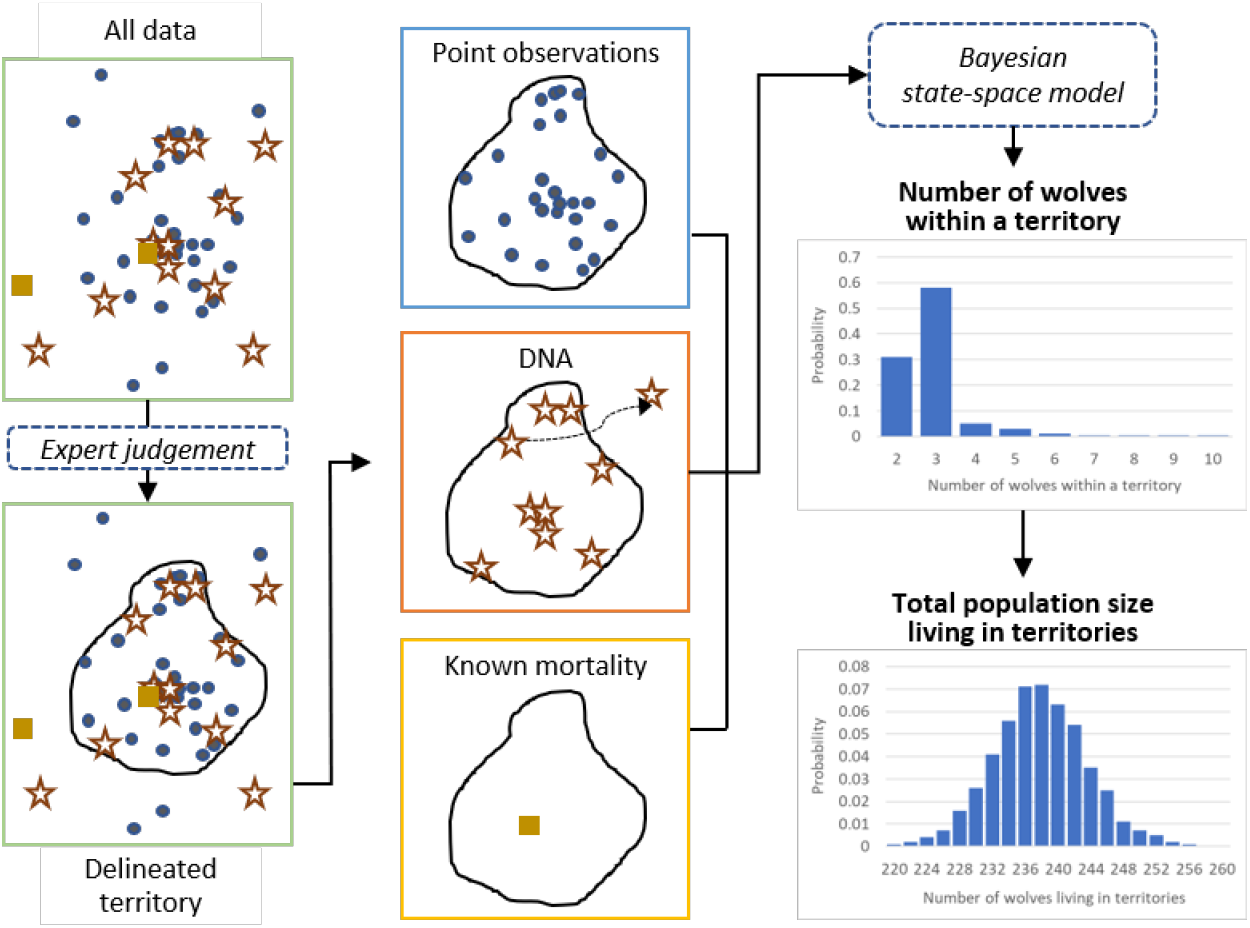
The role of the Bayesian state-space model in the process of annual Finnish wolf population assessment. In the first phase, the territories are delineated by experts. In the second phase, different data are used to infer the number of wolves within each territory using the developed Bayesian state-space model. The results of the analysis are provided as probability distributions.

### 2.1 Data

The purpose of the state-space model is to pool information about the number of wolves within each territory from three different data sources: point observations, non-invasive DNA samples and reported mortality. The data collection protocol is described in detail in Kojola et al (2018). Here we describe different data sets at the level sufficient to understand the modelling work in focus. Table 1 shows how data are organised for the two example territories that we refer to throughout the paper. The full data set is available as online resource.

**Table 1.** Data for two territories (Kallioluoma and Halivaara). DNA data associated to autumn season covers both autumn and spring season samples. Full data set for all territories is provided as online resource.

| Territory name | Season | Number of wolves observed together (count of observations) |  |  |  |  |  |  |  |  |  |  |  |  |  |  | DNA samples | Unique individuals | Known mortality |
| --- | --- | --- | --- | --- | --- | --- | --- | --- | --- | --- | --- | --- | --- | --- | --- | --- | --- | --- | --- |
|  |  | 1 | 2 | 3 | 4 | 5 | 6 | 7 | 8 | 9 | 10 | 11 | 12 | 13 | 14 | 15 |  |  |  |
| Halivaara | Autumn | 59 | 55 | 9 | 5 | 8 | 7 | 0 | 0 | 0 | 0 | 0 | 0 | 0 | 0 | 0 | 25 | 7 | 0 |
| Halivaara | Spring | 9 | 8 | 5 | 8 | 4 | 0 | 0 | 0 | 0 | 0 | 0 | 0 | 0 | 0 | 0 | 15 | 6 | 1 |
| Kallioliuoma | Autumn | 13 | 2 | 6 | 0 | 5 | 0 | 0 | 0 | 0 | 0 | 0 | 0 | 0 | 0 | 0 | NA | NA | 0 |
| Kallioliuoma | Spring | 0 | 0 | 0 | 0 | 0 | 0 | 0 | 0 | 0 | 0 | 0 | 0 | 0 | 0 | 0 | NA | NA | 2 |

#### Point observations

Since 2009 point observations of wolves in Finland have been reported in the digital large carnivore monitoring system called Tassu (“the Paw”). The observations are recorded by approximately 2 000 volunteer large carnivore contact persons, who have training in wolf ecology, behaviour, and paw print identification. For each observation the type (sighting, track, prey kill site, or livestock depredation), date, location, and the number of animals are reported, and if feasible, also the age status of animals and front paw print dimensions. The number of individuals associated with each observation is estimated by the contact person submitting the data. Contact persons have their own networks of local people, who are mostly hunters with capability in species identification. Such a network is particularly relevant for sightings, because sight observations cannot usually be validated afterwards in snow-free conditions. The observations are divided into observations made before and after the turn of the year, i.e., in autumn season (from August to the end of December, hereafter simply “autumn”) and in spring season (January-March, hereafter simply “spring”). All types of observations are treated similarly. Highest reported pack size, number of pair observations and number of observations concerning more than two wolves (packs) are used as model input.

The data set for March 2020 population assessment includes 2301 pair or pack observations gathered from 55 territories. The number of point observations reported per territory ranges from 1 to 149. The median number of observations is 25. The highest reported pack size observation over all territories is 11 wolves.

#### DNA samples

Non-invasive DNA samples of wolves have been extensively collected for population monitoring purposes from 2016 onwards. Samples (mainly scats) are collected by volunteers (from November to February) and wildlife professionals (from November to March). The samples are used for identification of individuals and for kinship analyses. For this analysis, the data are divided into samples collected in autumn and in spring seasons. Total number of successful DNA samples per territory and the number of different individuals found within these samples are used as model input.

The data set for March 2020 population assessment includes a total of 610 successful DNA samples, from which 190 different individuals were identified. The number of successful DNA samples per territory ranges from 0 to 43. The maximum number of individuals found from a territory is 10. Fifteen out of 55 territories have no DNA samples taken, whereas 50% of all territories have 10 or more successful samples. Samples from which enough DNA could be extracted for microsatellite genotyping were considered successful.

#### Known mortality

Wolves that are hunted with a derogation or damage control licence, found dead, or removed by police order, are sent to laboratory for autopsy. Regarding these individuals, several attributes are reported, including the location, date and cause of death, sex, and age. Tissue samples are taken for DNA analysis. The number of individuals found dead within each territory is used as model input.

#### Previous expert estimates of pack sizes

Before the development of the model presented here the number of individuals within each territory was assessed by the expert panel based on all the data available from each territory. The expert panel gave particular emphasis on the number of unique individuals found from DNA samples and on the highest observed pack size reported by volunteers. The panel also considered both the total number of DNA samples collected and the number of point observations provided. Data gathered in spring had more weight than the autumn data on the final estimate provided by the panel. Data on known mortality was used by the panel. In the face of uncertainty, the panel expressed the size of the pack as a range with minimum and maximum numbers of wolves. For each territory, minimum and maximum estimated numbers of wolves were reported: these figures are used as model input in this analysis. These types of data are available from wolf population assessments conducted in 2018 and 2019. The data set contains estimated pack sizes for 88 cases. This data set is used only as training data.

#### Training data and assessment data

The data sets are divided into two parts, which are modelled separately. *Training data* consists of point observations and previous expert estimates of pack sizes from years prior to the target year (2018-2019). The purpose of these data is to provide information about the link between the point observations and expert assessments. *Assessment data* consists of point observations, DNA samples, and known mortality for each territory in the target year (2020).

### 2.2 Model structure for wolf population assessment

Here we provide an overview of the model structure and the modelling process. Detailed technical description of the model is given in Appendix A. The computer code and data necessary for replicating our example analysis are provided at the bioR*χ*iv repository at https://www.biorxiv.org/content/10.1101/2021.12.21.473527v1.supplementary-material.

A prerequisite for the model is wolf territories inferred by the experts as described in Kojola et al (2018). The experts delineate the territories based on the spatial distribution of point observations of pairs and potential family packs as well as DNA samples, also using information on potential natural boundaries such as lakes and urban areas. We built our model within the Bayesian state-space modelling framework, which incorporates both process variation and the uncertainty related to observations in a single model (Mäntyniemi et al, 2015). The process model is used to describe the survival process within each territory and the observation models describe data generating processes related to point observations, DNA samples and known mortality. We assign prior distributions to the model parameters, and this prior knowledge is then updated with the information contained in data. The result is a joint posterior probability distribution of model parameters which contains the current state of knowledge on the number of individuals inhabiting each territory.

#### 2.2.1 Process model for survival within a territory

The survival model describes how the number of wolves within a territory can change over time. No reproduction takes place in wolf populations during the winter season. Thus, we assume that the number of resident wolves in a territory can only decrease. Although this assumption may not always hold because of the adoption behaviour (e.g., Mech and Boitani (2003)), we assume such behaviour to be relatively rare in Finland. Survival rate is assumed to vary between individual wolves around a common mean, which is not known exactly. This assumption creates a hierarchical structure where information can flow between territories. The mean survival rate of wolves is a model parameter which becomes estimated when the model is fitted to data from autumn and spring. Territories with more precise knowledge about the number of wolves in autumn and spring will provide more precise knowledge about the mean survival of wolves. This information is passed on to territories with less data when fitting the model to all territories.

Prior distribution for the number of wolves in each territory has a similar hierarchical structure. Territories are assumed to be exchangeable *a priori* in terms of the number of wolves that they contain: we do not know which territory might have larger or smaller number of wolves than the others. This assumption means that the number of wolves in each territory can be thought of as a random draw from a hypothetical superpopulation of wolf territories. In other words, the probability of having, say, 6 wolves in a territory is equal to relative frequency of territories containing 6 individuals in a very large population of wolf territories. These relative frequencies are not known precisely, but they are assigned a prior distribution based on the distribution of wolf pack sizes in the past. This hierarchical structure provides another path by which territories can exchange information about the number of individuals. Hence, the prior distribution for each territory can be thought to be based on the distribution of pack sizes in other territories in the past and in the current year.

#### 2.2.2 Observation model for point observations from Tassu

The observation model for point observations describes the link between the true number of wolves and the point observations made by citizens within a territory. Prior knowledge about this link is available from earlier wolf population assessments, where the number of wolves within each territory was assessed by experts based on point observations, DNA data, and known mortality. Finding a statistical model between the expert estimates and point observations formalises the logic of expert interpretation of point observation data. After this, the data can be automatically applied and included as a source of information in the state-space model.

Based on data from years 2018-2019, it was found that the expert estimated pack size is positively associated with the highest pack size reported by the volunteers. This relationship was modelled as a linear regression through origin (Fig 2). The expert estimated pack size was negatively associated with the proportion of pair observations of all observations concerning packs or pairs. This relationship was modelled with logistic regression (Fig 2).

**Fig. 2.**
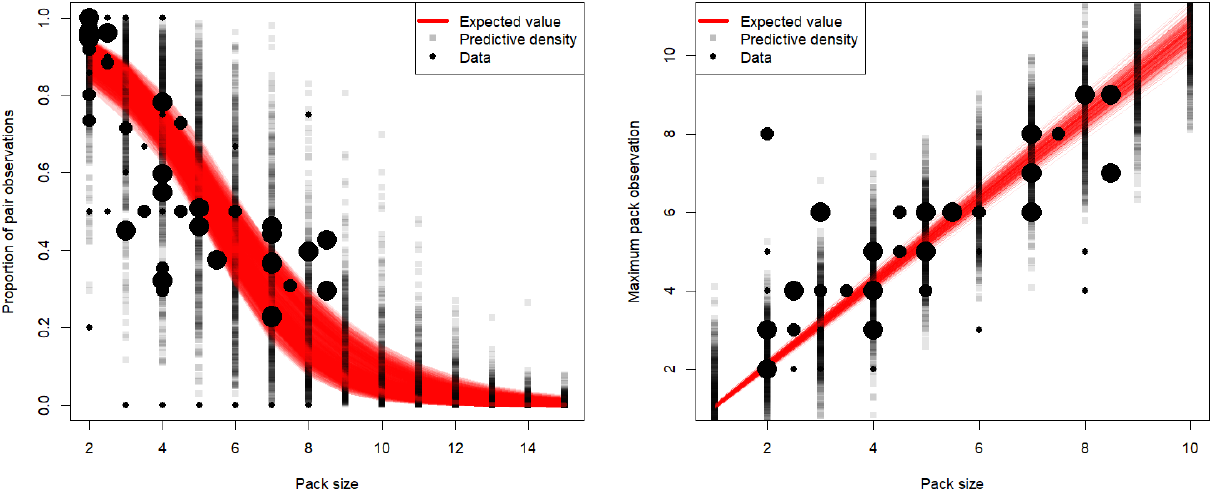
Model fit for training data on the relationship between proportion of pair observations (left), observed maximum pack size (right) and estimated pack size (x-axis). Red lines are based on random draws of regression parameters from their joint posterior distribution. Grey squares are random draws from posterior predictive distribution, that will be used in population estimation in subsequent analysis. Black circles represent training data. Larger circle indicates larger amount of observations. The estimated pack size is plotted by using the mean estimate, when the estimated pack size was uncertain.

Posterior distributions of linear and logistic regression parameters based on the fit to data from 2018 and 2019 contain the information about the link between the true number of wolves and point observations. When fitting the state-space model to the target year data, these posteriors are used as prior distributions. These priors will be further updated when fitting the model to the target year data, provided that there are at least some territories from which lot of DNA data are available. This structure provides the third mechanism by which territories can exchange information through common parameters.

#### 2.2.3 Observation model for DNA samples

Miller et al (2005) presented a model by which the size of a small population can be estimated based on DNA recaptures. The simplest form of the model assumes that all individuals have equal probability of being captured by DNA sampling. While this assumption is hardly true in the case of a wolf population spanning over a very large area where the sampling effort is highly variable, it can be realistic within a single wolf territory, where all the pack members are well mixed and move around the same area (but see Discussion). In this case, individual sampling histories are not needed. The number of successfully analysed samples and the number of different individuals found serve as sufficient statistics, on which the inference can be based.

We use the likelihood function derived by Miller et al (2005) as a link between the true number of wolves within a territory and DNA data. DNA samples collected during autumn and spring are assumed to be informative about the number of wolves in autumn. This accounts for the fact that individuals present in spring must have been present also in autumn. However, only samples collected in spring are assumed to be informative about the number of wolves present in spring. This accounts for the fact that individuals found in autumn but not in spring may have died during the winter.

#### 2.2.4 Observation model for known mortality

Wolves found dead within a territory provide a lower bound for the total number of wolves that may have died during the winter. At the same time, they also provide a lower bound for the total number of wolves that were initially alive in autumn. Known mortality is treated as a censored observation about the total number of dead wolves in a territory. This can provide information about all the model parameters.

### 2.3 Expert assessment

In order to compare the results of the new model to results obtained using the previous method based on expert assessment, the same data set was shown to I. Kojola, who has to most experience on the logic used in previous wolf population assessments in Finland. He assessed the number of wolves in each territory by giving the minimum and maximum estimates for the pack sizes in autumn and in spring. We used the mean of the range to compare the expert estimate to the posterior mean from model output for each territory. The total number of wolves inhabiting the territories was described as a range obtained by summing up the territory specific minimum and maximum estimates.

## 3 Results

### 3.1 Training data

We fitted the model first to training data from 2018 and 2019 and used the resulting posterior distribution of linear and logistic regression parameters as a prior when fitting the model to assessment data from 2020. Model fit for training data is shown in Fig 2, which also illustrates the predictive density of data values conditional on pack size. When estimating the pack size given an observed value, the likelihood function implied for the pack size can be visually seen from the graph by fixing a horizontal line at the observation. For example, proportion of pair observations equal to 0.4 would support pack sizes from 2 to 10 with most weight on 4 to 7.

### 3.2 Territories

Fitting the model to assessment data produces a posterior distribution for the number of wolves in each territory. The posterior distributions were approximated using Markov chain Monte Carlo simulation, which means that summaries of the population can be derived simply by calculating the summary statistic for each iteration of the simulation and examining the distribution of the summary statistic.

Posterior distributions of the total number of wolves inhabiting the territories in autumn and spring are shown in Fig 3 together with the posterior distribution of the mean survival rate of all wolves. In addition to density and probability plots, posterior distributions can also be summarised with probability intervals (PI) that contain a specific amount of probability mass. For example, 90% PI for the number of individuals in autumn is [256,278]. The interval has the interpretation that the true value is thought to lie within this range with 90% probability. This is different from the concept of a confidence interval, which does not have interpretation as a formal statement of uncertainty about the variable of interest. The most probable number of individuals inhabiting the territories in spring is 218 with 90% PI of [209,228]. The mean survival rate of wolves over the winter is estimated to be [0.77,0.86] with 90% probability.

**Fig. 3.**
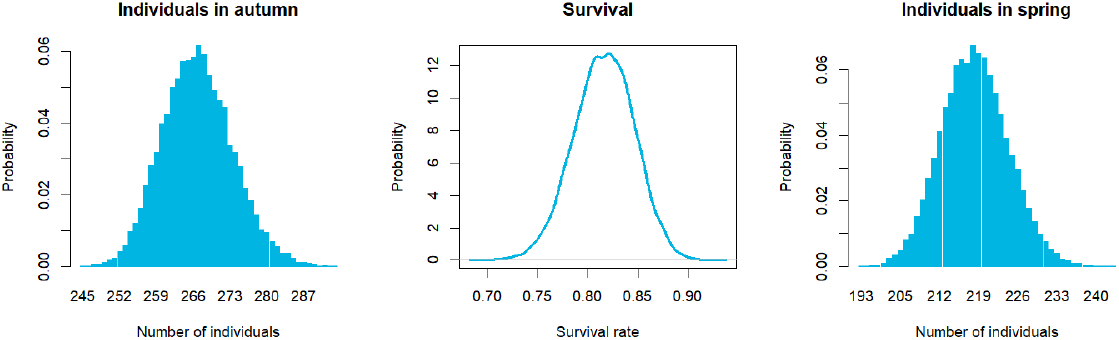
Marginal posterior distributions of the number of wolves in territories in autumn, survival rate from autumn to spring, and the number of wolves in territories in spring.

Expert assessment based on the same data resulted in lower estimates for the number of individuals compared to model based posterior distributions. Based on the expert assessment the range for autumn was [216,247] which does not intersect the 90% PI of the posterior distribution. The expert estimate for spring was [176,204] which is lower than the 90% posterior PI.

Posterior distributions of total numbers of packs, pairs, and empty (less than two wolves) territories are shown in Fig 4. These variables have negative posterior correlation, because there are territories for which the classification is uncertain but which can only belong to one class. Non-zero posterior correlation means that there is information about probable combinations of packs, pairs and empty territories that could not be seen directly from the marginal distributions of these variables. For example, combinations where both packs and pairs are either high or low are less probable than combinations where either one is high and the other one is low.

**Fig. 4.**
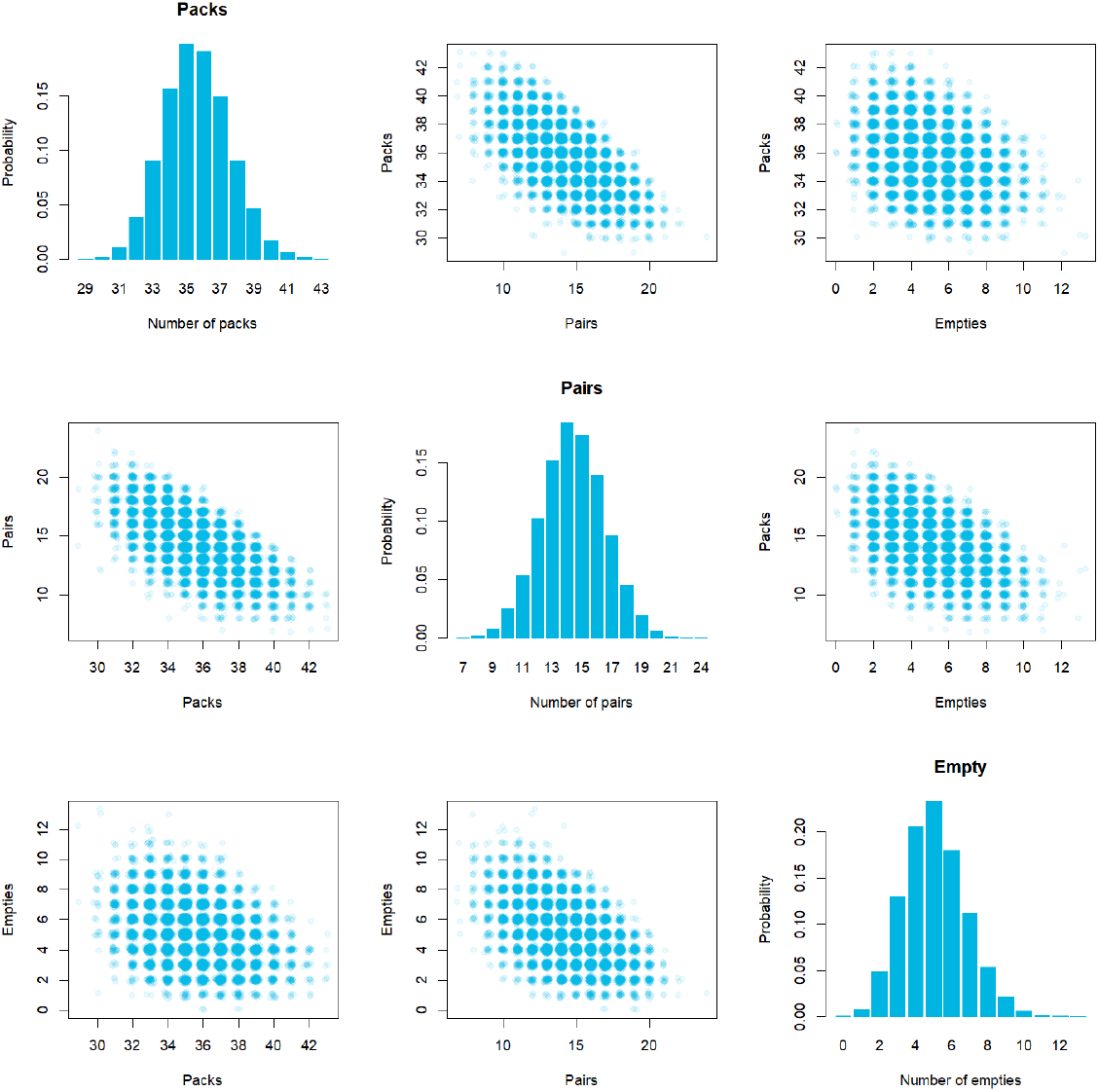
Marginal posterior distributions and posterior correlations of the number of territories occupied by packs, the number of territories occupied by pairs, and the number of empty territories.

The functioning of the model is demonstrated with two territories, Halivaara (Fig 5) and Kallioluoma (Fig 6). Halivaara is an example of a territory with high amount of data. It has total of 27 DNA samples collected throughout the season, from which 7 different individuals were identified. This provides strong information that places almost 100% probability for 7 wolves in autumn (Fig 5). 15 out of 27 DNA samples were collected in spring, from which 6 different individuals were identified. In addition, one individual was found dead, which further corroborates the inference that the number of wolves decreased during the winter (Fig 5). Because there is high certainty that there were more than two wolves in the territory, the territory is classified as a pack with 100% probability (Fig 5).

**Fig. 5.**
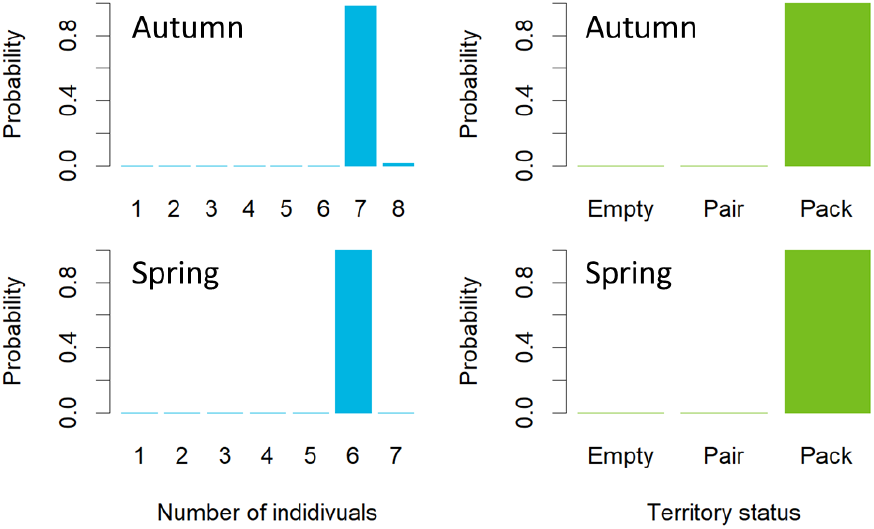
Posterior distributions for the number of wolves (on the left) and territory status (on the right) in Halivaara territory in autumn (upper row) and in spring (lower row).

**Fig. 6.**
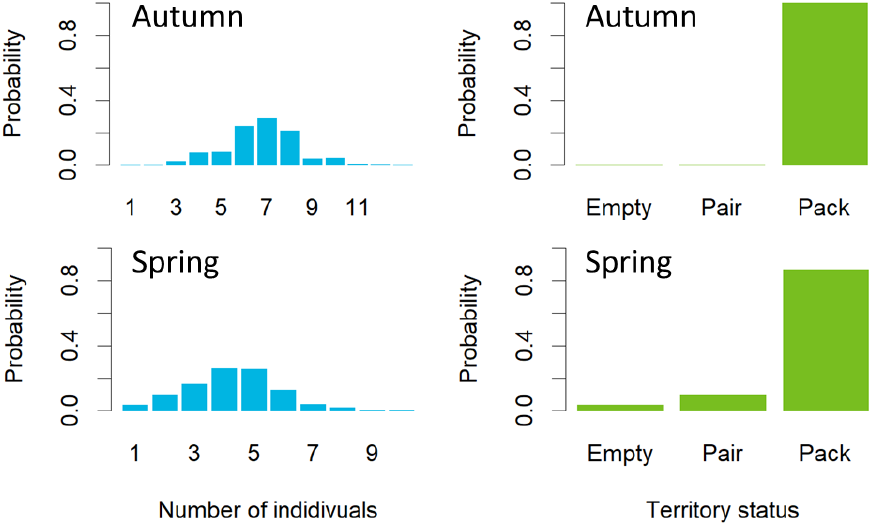
Posterior distributions for the number of wolves (on the left) and territory status (on the right) in Kallioluoma territory in autumn (upper row) and in spring (lower row).

Kallioluoma territory provides a contrasting example with no DNA samples (Fig 6). There were 13 point observations reported on 2 or more wolves, and all point observations were made in autumn. One wolf was found dead during the winter. The posterior distribution of the number of wolves in autumn is wide, with the highest probability on 7 individuals (Fig 6). Highest reported point observation is 5 wolves, but the low proportion of pair observations (15%) is typical for pack sizes larger than 5 (Fig 2). Hence, the probability of such packs increases. The posterior distribution for the number of wolves in spring (Fig 6) is mostly based on estimated survival rate of wolves across all territories and on the number of wolves in autumn. This is backed up also by the one observed death. Kallioluoma territory serves also as an example of a territory for which the status is not certain. Occurrence of territories with uncertain status creates the negative correlation (Fig 4) between the total numbers of packs, pairs, and empty territories.

Data and results from all territories can be downloaded from the bioR*χ*iv repository at https://www.biorxiv.org/content/10.1101/2021.12.21.473527v1.supplementary-material.

Comparison of posterior means from the model against the means of expert assessments for each territory shows a clear, almost one to one, positive relationship both in autumn and spring seasons (Fig 7). The model tends to give slightly higher pack sizes on average than the expert estimation. However, for a few territories the difference seems quite large. See Discussion for potential reasons for these differences.

**Fig. 7.**
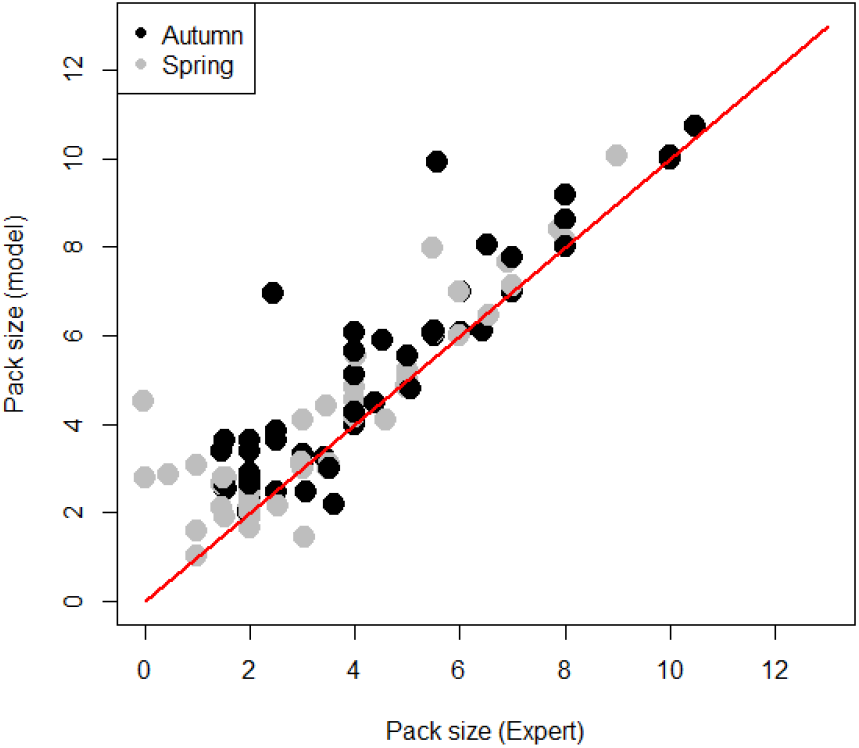
Comparison of expert assessment (x-axis) and model output (y-axis) for the estimated pack size in each territory. Mean of the posterior distribution of the pack size is shown for the model output. Expert estimates are the mean of minimum and maximum estimates for each territory. One to one relationship is shown as a red line.

## 4 Discussion

It is evident that the management of species like wolf, which evokes conflicts while being vulnerable to various human activities, requires reliable population assessment methods. It is also important to acknowledge the strong demand coming from various stakeholder groups in the society concerning the objectivity and transparency of population assessments and existing uncertainties. Further, focusing on a large carnivore which prefers forested areas and exhibits elusive behaviour, the assessment methods substantially benefit from as comprehensive use of all existing data as possible.

By adopting the Bayesian approach, we have been able to meet these demands. The method provides the result as a probability distribution, enabling intuitive communication of uncertainty. When the result is depicted as a probability distribution, the most probable number of wolves per territory is easily interpreted, but also the uncertainty related to the estimate is straightforward to perceive. By communicating results this way, also the less probable but still possible numbers of wolves are made visible. The hierarchical structure of the model allows for efficient use of all the available information, where data rich territories can share their information with data poor territories (Punt et al, 2011).

The model integrates different types of citizen-provided data. DNA samples provide information about the pack size based on the number of different individuals found from the same territory. The higher the sample size, the higher the probability that all individuals become sampled. It is worth noting that also other kinds of observations, from which different individuals can be identified, could be used in the model in addition to or in place of DNA samples. For example, Mattioli et al (2018) used individual wolves identified from camera traps as individual recaptures. However, for these to be informative in the sense of individual identification, the pack must include enough phenotypic variation so that at least some of the individuals can be distinguished from the rest.

Citizen-provided point observations provide information about the pack size based on two factors: the maximum number of wolves observed at once, and the proportion of pair observations compared to all observations concerning two or more wolves. The total number of point observations made from a territory increases the precision of the pack size estimation. It also increases the weight of this information source when combined with the information provided by DNA samples.

We believe that the seamless integration of different information sources using a Bayesian state-space model can be beneficial also in other cases, where different types of data are collected to infer the wolf population size. For example, Ausband et al (2022) compared the use of camera traps and DNA samples, and Stenglein et al (2010) compared the use of DNA samples and GPS surveys. In both cases, it would be possible to embed both data sources into a single model in the same way as we did for citizen-provided DNA samples and point observations. Different data sources become calibrated against each other, and they can then be flexibly used in the analysis to different degree in different territories.

Direct comparison of the developed model with other quantitative methods is not possible. To our knowledge, the sampling design involving a network of volunteer citizens is unique to Finland, and comparable data is not collected anywhere else. Other estimation methods typically rely on known observer effort (e.g. Bischof et al, 2020; Jiménez et al, 2016; López-Bao et al, 2018), which is not available in Finland. This means that models for analysing this type of data have not been presented before. However, we outline potential paths for development and similarities to other modelling approaches that have been proposed for the population estimation using different types of data.

The only alternative approach for this particular estimation problem is the expert panel approach used previously in Finland. We have identified at least two cases, where the model could estimate slightly higher pack sizes than the expert panel. First, when the average number of DNA samples per wolf is low in a territory, there is a considerable chance that not all wolves have been sampled during the season. In this case, the model correctly gives posterior probability also for the pack sizes larger than the number of unique individuals found in the DNA samples. However, this can be difficult to grasp for human mind, and the expert estimate is typically equal to the number of individuals identified by DNA, especially if the point observations do not strongly suggest higher pack sizes.

In such a situation, the model typically produces a probability distribution with a posterior mode corresponding with the number of unique individuals found in the DNA samples and the pack sizes larger than the mode having smaller probabilities. The posterior mean is then higher than the mode. This phenomenon is visible in Fig 7 where the posterior means are slightly higher than the expert estimates. This matters when we add up skew probability distributions from multiple territories. When the number of territories increases, the distribution of the sum starts to resemble a Gaussian distribution, and the mode of the sum will be larger than the sum of the modes from individual territories. In this case, the model works logically and the expert judgement could be considered to underestimate the true number of wolves.

Another situation where the expert judgment and the model might disagree to some extent, can occur when there are only very few (or zero) observations in spring, but the territory is included in the assessment based on observations made in autumn. Our example from the Kallioluoma territory (Fig 6) falls into this category. In such a case, the expert panel would be prone to conclude that wolves have disappeared from the territory during the winter. For the model, however, the reason for the absence of data is that volunteers are not reporting observations, and hence, there is no new information about the status of the pack. Thus, the status of the territory in spring is predicted based on two factors: the status in autumn and the estimated survival rate of wolves in other territories. The correct interpretation varies case by case and cannot be deduced beforehand. Few cases like this are visible in Fig 7 where the expert has estimated pack size of zero, but the model provides considerably higher estimate.

The expert panel may take into consideration the amount of data gathered in autumn as well as some external knowledge on environmental conditions. For example, if the reporting intensity was already low in autumn and environmental conditions for wolf observing were challenging in spring, the panel might conclude that wolves can still be present in the territory. Conversely, a sudden collapse in previously high reporting activity under favourable observing conditions would mean that the pack has vanished. Currently the model does not consider the temporal variation in reporting activity nor does it utilise data on environmental conditions. These are targets for future development of the model. For territories for which there are no data available in spring, the model can overestimate the number of wolves. We suggest that such cases (no data in spring) should be considered by an expert panel for an ad hoc decision on whether the territory can be included in the model or not.

The model has other limitations and potential targets for development. First, the assumption that all wolves within a territory have an equal probability to end up in the DNA sample may not always be exactly correct. For example, adult wolves may use their feces as territory marks more often than younger individuals by leaving them on forest roads and paths, which may serve as territory boundaries (Vilà et al, 1994). Humans may tend to use the same routes and may thus have higher chance of spotting these feces for DNA sampling.

Generally, violating the assumption of equal detectability leads to under-estimation of population size (Cubaynes et al, 2010; Mäntyniemi et al, 2005; Marescot et al, 2011). In the context of our model, such violation could have some relevance in territories occupied by large family packs, where offspring may potentially have smaller detectability in DNA sampling than adults. Territories inhabited by pairs without offspring would not suffer from this problem. High reporting intensity related to point observations is expected to mitigate the impact of unequal detectability in DNA sampling. Further, DNA sampling with high intensity is expected to have a similar effect, as it increases the probability that individuals with low detectability become sampled at least once.

In summary, large family packs with low point observation and DNA sampling intensity can be prone to the underestimation of the population size. The model could be further developed to account for unequal sampling probability, potentially by utilising the model structure presented by Miller et al (2005) for unequal sampling. However, it is worth noting that Stenglein et al (2010) used the method of Miller et al (2005) for the estimation of wolf abundance based on DNA samples (from wolf feces) but did not find any evidence of unequal sampling probability.

Second, the model does not consider the history of territories. Coupling consecutive years together would enable a structure where the past state of the territory could be used to inform the prior distribution for the number of wolves in the territory next year. For example, an empty territory could have higher probability of being empty also in the next year. If such an autocorrelation in the territory history was found, it would also increase the precision of the population assessment, or the same assessment precision could be achieved with smaller number of observations.

A natural extension of a model with a territorial autocorrelation would be an individual based model for each territory. This would provide a platform where individual DNA captures and survival probabilities could be explicitly modelled. Adults, sub-adults and pups could be identified by following the DNA samples over years.

Third, further analysis of the relationship between point observations and the pack size might focus on exploring other functional shapes than linear and logistic regression. For example, anecdotal evidence suggests that the relationship between the observed maximum pack size and the pack size could be, at least to some extent, non-linear: for territories with large packs (confirmed by DNA), the observed maximum pack size tends to be smaller than the actual pack size. This might reflect a potential tendency of large packs to split into subgroups more often than smaller packs. Furthermore, for the largest packs the estimate for pack size based on tracks in snow tends to be smaller than the real pack size.

Another line of future research would be to explore the possibility to develop open-population spatial capture-recapture (OPSCR) models (Bischof et al, 2020) for Finnish wolf population assessment. The main difference between the model presented here and the OPSCR model developed by Bischof et al (2020) is the focus of the assessment. Our main target of inference is the number of packs and pairs occupying the expert-provided territories, whereas the OPSCR model provides estimates of wolf density and total abundance on a spatial grid. Another key difference is the data available for the analysis. The OPSCR model relies on extensive genetic monitoring and known mortality with mostly known observer effort, whereas the model presented here integrates also point observations reported by volunteers and does not require known observer effort, which is not available in Finland.

We have assumed that the size of a wolf pack does not increase during the winter. This assumption would be violated, if a family pack accepted a dispersing wolf as an adoptee. While such behaviour is possible (e.g., Mech and Boitani (2003)), it is believed to be too rare in Finland to significantly affect the population estimation. If an additional wolf joins the pack during winter, it can be expected that the estimate of the number of wolves already present in autumn goes up, or the estimate of wolf survival goes up, or both. What happens in a particular territory mainly depends on the amount of information available about the number of wolves in autumn. However, as the DNA data accumulates in the future, the extent of adoptions in Finland can be better studied with kinship analyses.

Currently, the monitoring of wolf population in Finland is heavily dependent on point observations reported and DNA samples collected by volunteers. Both types of data are prone to changes in external circumstances. In most parts of Finland, the snow-covered season has shortened (Luomaranta et al, 2019), which may affect the number of observations in the future. Further, the number of collected DNA samples can be highly dependent on, e.g., the general opinion on the usefulness of DNA collection for volunteers. Such potential changes in available data does not affect the developed model as such, but they may increase the uncertainty associated with the results. This emphasizes the need to maintain and reinforce further the collaboration between the research and volunteers.

## Supporting information

Code, data and all results

## Supplementary information

Full data sets, the computer code for running the analysis and detailed territory-specific results are available for download from the bioR*χ*iv repository at https://www.biorxiv.org/content/10.1101/2021.12.21.473527v1.supplementary-material.

## Acknowledgments

We wish to thank all volunteers and research personnel, who have contributed to the collection of data used in this study. We also wish to thank Mia Valtonen, Helena Johansson and the anonymous referees whose comments greatly improved the quality of this manuscript.

## Declarations

Authors declare no conflict of interest.

## Appendix A Technical description of the Bayesian state-space model

This appendix provides assumptions and mathematical details of the Bayesian state-space model used to estimate the number of wolves in Finnish wolf territories. List of symbols used for modelling the training data is presented in table A1. Symbols used for the state-space model are listed in tables A2 and A3.

## A.1 Training data

The relationship between volunteer made observations and the estimated number of wolves in a territory was analysed from Finnish wolf population assessments conducted in 2018 and 2019. Data comprises of 88 territories from which the minimum and maximum estimates of the number of wolves in each territory in spring was available. The estimated pack size was found to be positively associated with the highest observed wolf pack size reported by the volunteers (Fig 2). This relationship was modelled as a linear regression through the origin as follows

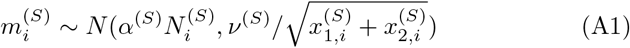

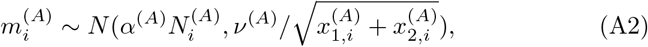

where 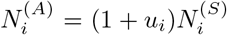. This structure reflects the fact that the number of wolves in the territory in autumn must have been larger or equal than the number of wolves in the following spring. The residual standard deviation is scaled by the square root of the total number of pack and pair observations made from the territory. This structure implies that the maximum observed pack size is more informative about the pack size when more data is reported from the territory.

The estimated pack size was found to be negatively associated with the proportion of pair observations from all pack or pair observations (Fig 2). This relationship was modelled using a random effects logistic regression in spring

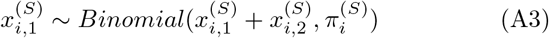

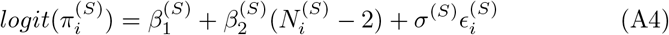

and in autumn:

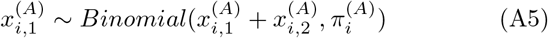

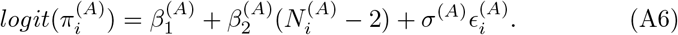

## A.2 State-space model structure for wolf population assessment

Territories are first inferred using expert judgement as in Kojola et al (2018). This model infers the number of individuals within each territory.

This is a Bayesian state-space model which describes the latent survival process in each territory. Different data sets are linked to the relevant states of the population using observation models.

## A.2.1 Survival process

Winter is divided into two phases: autumn and spring. Survival events of wolves are assumed to be independent of each other. The survival probability of all wolves across all territories is assumed to vary between individuals with mean survival probability *θ*. As shown by Mäntyniemi et al (2015), these assumptions lead to binomial model for the number of wolves 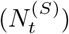 that survive from autumn 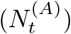 to spring. We approximate the Binomial model with a scaled Beta distribution for computational convenience (Mäntyniemi et al, 2015):

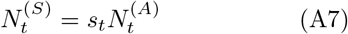

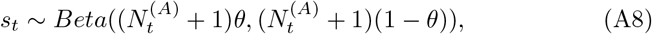

where *s_t_* is the proportion of wolves that survive to spring in territory *t*. Number of wolves in spring 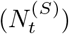 is rounded to nearest integer. If the number of wolves is zero after the rounding, then the number of wolves is set to one. This is for computational convenience, that does not affect the results, because only pack sizes of two or more are of interest.

Prior distribution for the number of wolves in a territory in autumn is defined using a hierarchical structure. Vector of proportions *ω*_1_,…, *ω*_15_ describes the relative frequencies of different pack sizes in a hypothetical super-population of wolf packs, from which each pack is considered to be a random draw with replacement. The maximum possible wolf pack size is assumed to be 15. A Dirichlet prior is assigned to the vector of proportions

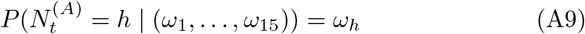

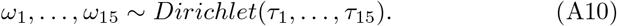

where parameters *τ*_1_,…, *τ*_15_ are fixed based on the distribution of pack sizes in earlier years. The hierarchical structure enables information to flow from territories with lots of data to territories with smaller number of observations.

The number of wolves that die during the winter is given by

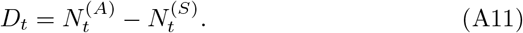

## Model for volunteer observations

The observation model for the maximum observed pack size is identical to the observation model used for the training data:

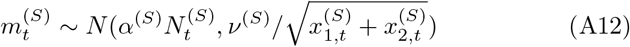

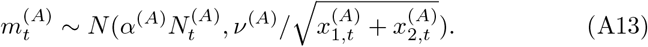

The observation model for the number of pair observations given the number of wolves in the territory and the number of pack observations is identical to the observation model used for the training data:

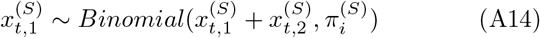

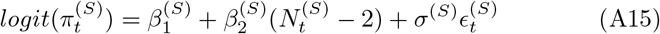

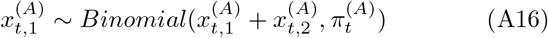

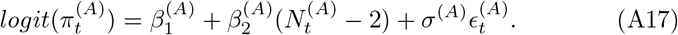

Prior distribution for the parameter vector

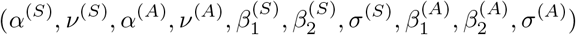

is a multivariate normal distribution with mean vector and covariance matrix equal to posterior mean vector and covariance matrix obtained by analysing the training data from 2018 and 2019 wolf population assessments.

## DNA observations

All wolves within a territory are assumed to have equal chance to end up in the DNA sample. Sampling is conducted with replacement and getting sampled is assumed to not affect the future chance of getting sampled. Under these assumptions the likelihood function for the number of wolves in a territory given the number of successful samples collected and the number of different individuals found is proportional to a multinomial distribution (Miller et al, 2005). Dropping constant terms yields the following likelihood for the number of wolves in territory *t* in autumn

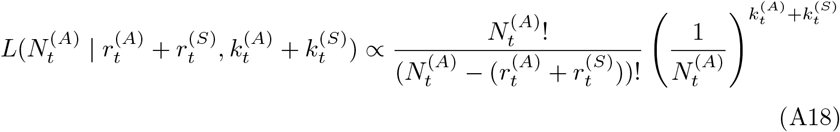

where 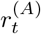 and 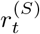 are the number of different individuals found in territory *t* in autumn and in spring, respectively. Number of successful DNA samples collected in autumn and spring are denoted 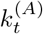 and 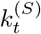. The spring samples are added to autumn samples, because any pack member alive in spring must have been a pack member also in previous autumn. The likelihood for the number of wolves in a territory in spring depends only on the samples collected in spring:

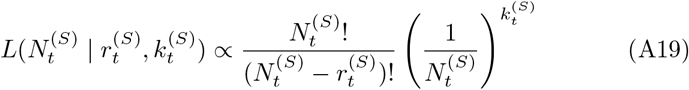

## Known mortality and dispersal

Observed number of dead wolves (*d_t_*) in a territory is treated as an interval censored observation about the true number of dead wolves, so that the observed number of dead wolves gives the lower bound for the true number of wolves that died in the territory.

If the last DNA sample of a wolf is found far away from a territory where the other samples where found, then the DNA sample is excluded from the data. This is because the target is to estimate the number of wolves that occupy the territory in spring.

## A.2.2 Computation

The joint posterior distribution of model parameters was approximated using Markov chain Monte Carlo (MCMC) simulation. The simulation was implemented using JAGS version 4.3.0 (Plummer, 2003). Pre-processing of data and post-processing of simulation results was conducted using R 3.6.0.

The burn-in period of the MCMC simulation for the population assessment was run using two chains for 10000 iterations, after which 1000000 iterations were produced with thinning of 1000. Convergence of the simulation was assessed using visual inspection of the chains.

---

**Table A1.**
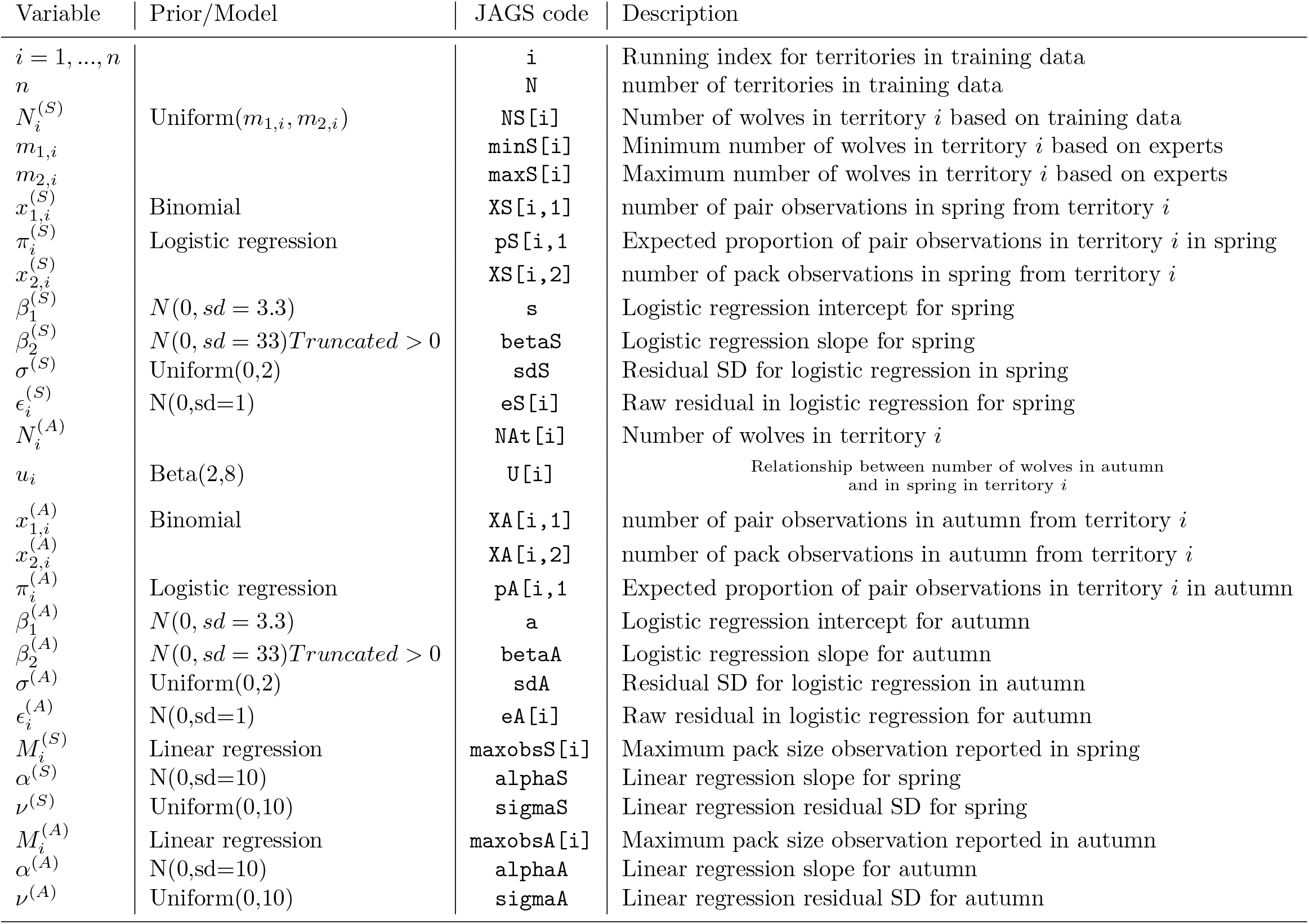
Description of variables and notation for the model used to analyse the training data. Prior distributions are reported when relevant. Corresponding JAGS variables are also given for reference.

**Table A2.** Description of variables and notation for the model used to analyse the population assessment data. Prior distributions are reported when relevant. Corresponding JAGS variables are also given for reference.

| Variable | Prior/Model | JAGS code | Description |
| --- | --- | --- | --- |
| $t = 1, \dots, n_A$ | | **t** | Running index for territories in population assessment data |
| $n_A$ | | **Territories** | number of territories in training data |
| $N_t^{(S)}$ | | **n[t,2]** | Number of wolves in spring in territory $t$ |
| $x_{1,t}^{(S)}$ | Binomial | **pairs[i]** | number of pair observations in spring from territory $t$ |
| $\pi_t^{(S)}$ | Logistic regression | **p[i]** | Expected proportion of pair observations in territory $t$ in spring |
| $x_{2,t}^{(S)}$ | | included in **Sample[i]** | number of pack observations in spring from territory $t$ |
| $\beta_1^{(S)}$ | posterior from training data | **a[2]** | Logistic regression intercept for spring |
| $\beta_2^{(S)}$ | posterior from training data | **beta[2]** | Logistic regression slope for spring |
| $\sigma^{(S)}$ | posterior from training data | **sd[2]** | Residual SD for logistic regression in spring |
| $\epsilon_t^{(S)}$ | $N(0, \text{sd}=1)$ | **eps[i]** | Raw residual in logistic regression for spring |
| $N_t^{(A)}$ | | **n[t,1]** | Number of wolves in territory $t$ in autumn |
| $s_t$ | | **s[t,2]** | Proportion of wolves that survive from autumn to spring in territory $t$ |
| $\theta$ | $\text{Beta}(2,2)$ | **surv** | Survival probability of wolves from autumn to spring in all territories |
| $x_{1,t}^{(A)}$ | Binomial | **pairs[i]** | Number of pair observations in autumn from territory $t$ |
| $x_{2,t}^{(A)}$ | | included in **Sample[i]** | number of pack observations in autumn from territory $t$ |
| $\pi_t^{(A)}$ | Logistic regression | **p[i]** | Expected proportion of pair observations in territory $t$ in autumn |
| $\beta_1^{(A)}$ | posterior from training data | **a[1]** | Logistic regression intercept for autumn |
| $\beta_2^{(A)}$ | posterior from training data | **beta[1]** | Logistic regression slope for autumn |
| $\sigma^{(A)}$ | posterior from training data | **sd[1]** | Residual SD for logistic regression in autumn |
| $\epsilon_i^{(A)}$ | $N(0, \text{sd}=1)$ | **eps[i]** | Raw residual in logistic regression for autumn |

**Table A3.** Description of variables and notation for the model used to analyse the population assessment data. Prior distributions are reported when relevant. Corresponding JAGS variables are also given for reference.

| Variable | Prior/Model | JAGS code | Description |
| --- | --- | --- | --- |
| $m_t^{(S)}$ | Linear regression | `max[i]` | Maximum pack size observation reported in spring |
| $\alpha^{(S)}$ | posterior from training data | `alpha[2]` | Linear regression slope for spring |
| $\nu^{(S)}$ | posterior from training data | `sigma[2]` | Linear regression residual SD for spring |
| $m_t^{(A)}$ | Linear regression | `max[i]` | Maximum pack size observation reported in autumn |
| $\alpha^{(A)}$ | posterior from training data | `alpha[1]` | Linear regression slope for autumn |
| $\nu^{(A)}$ | posterior from training data | `sigma[1]` | Linear regression residual SD for autumn |
| $\omega_1, \dots, \omega_{15}$ | Dirichlet | `Np[1:15]` | Proportions of wolf pack sizes 1 to 15 in a superpopulation of wolf packs |
| $\tau_1, \dots, \tau_{15}$ | | `alphaN[1:15]` | Prior parameters for $\omega_1, \dots, \omega_{15}$ |
| $D_t$ | | `dead[i]` | Number of wolves that died between autumn and spring in territory $t$ |
| $d_t$ | | `D[i]` | Number of wolves found dead between autumn and spring in territory $t$ |
| $r_t^{(A)}$ | | included in `x[i]` | number of different DNA individuals found from territory $t$ in autumn |
| $r_t^{(S)}$ | | `x[i]` | number of different DNA individuals found from territory $t$ in spring |
| $k_t^{(A)}$ | | included in `k[i]` | number of successful DNA samples from territory $t$ in autumn |
| $k_t^{(S)}$ | | `k[i]` | number of successful DNA samples from territory $t$ in spring |

