## Supplementary material for "Assessment of the residential Finnish wolf population combines DNA captures, citizen observations and mortality data using a Bayesian state-space model": Code, data and all results: data_prep.r

```
## This script prepares the data set for use with the Bayesian state-space model


## Read posteriors mean and covariance matrix estimated using the training data

mu=dget("results/prior_means")
cov=dget("results/prior_covariance")

## Read territory specific observations from excel file

obs=read_excel("data/tassu.xlsx")
names=as.matrix(obs)[,1]
obs=data.matrix(obs)
Sample=rowSums(obs[,4:17])       # Sample: sum of all wolf observations containing at least 2 wolves in a territory
pairs=obs[,4]                    # pairs: number observations containing 2 wolves in a territory
index=Sample==0                  # Find territories with no observations
pairs[index]=NA                  # and set their pair observations to NA

## Find the maximum observed wolf pack size

max=c()
for(i in 1:length(pairs)){
  z=obs[i,4:17]
  m=c()
  for(j in 1:14){
    if(z[j]>0) m[j]=j
    else m[j]=0
  }
  max[i]=max(m)+1
}

### Create data list

data=list(n_obs=length(obs[,1])
          ,Territories=length(obs[,1])/2,Months=2,alphaN=c(0,60,20,20,7,11,11,6,2,1,1,1,1,1,1),
          x=obs[,"x"],k=obs[,"k"],z=obs[,"z"],pairs=obs[,4],
          max=max,missing=obs[,"missing"],
          D=obs[,"removed"]-1,I=obs[,"I"]
          ,Sample=Sample,terr=obs[,"terr"],month=obs[,"Month"],
          mu=mu,tau=ginv(cov))

### Create initial values

s=array(NA,dim=c(data$Territories,2))
n=array(NA,dim=c(data$Territories,2))
for(i in 1:data$Territories){
  s[i,1]=NA
  s[i,2]=0.8
  n[i,1]=15
  n[i,2]=NA
}

## Initial values for two chains

inits=list(list(Np=c(0,0,0,0,0,0,0,0,0,0,0,0,0,0,1),n=n,s=s),list(Np=c(0,0,0,0,0,0,0,0,0,0,0,0,0,1,0),n=n-1,s=0.95*s))
```
