## Supplementary material for "Assessment of the residential Finnish wolf population combines DNA captures, citizen observations and mortality data using a Bayesian state-space model": Code, data and all results: JAGS_model.r

```
model{
  
  
  
    
    
    ## Sum up packs & pairs & individuals
    
    for(m in 1:Months){
    Total[m]=sum(n[,m])
    Singles[m]=sum(single[,m])
    Pairs[m]=sum(pair[,m])
    Packs[m]=sum(pack[,m])
    }
  
  
  for(t in 1:Territories){
    
      for(m in 1:Months){
        
      # Count packs, pairs and single wolf territories based on number of ind in each territory
      
      single[t,m]=equals(n[t,m],1)
      pair[t,m]=equals(n[t,m],2)
      pack[t,m]=step(n[t,m]-3)
      
      }
      
      # Pior for initial population size in November. Possible range 1-15 wolves
    
      n[t,1]~dcat(Np[1:15])
      dead[t,1]~dnorm(0,1)  # just a dummmy variable to fill up the matrix. Not used for anything
      
      # simple population dynamics model for survival of wolves from november to march
      
      for(m in 2:Months){
        n_star[t,m]=round(s[t,m]*n[t,m-1])
        n[t,m]=ifelse(n_star[t,m]==0,1,n_star[t,m])
        s[t,m]~dbeta(surv*n[t,m-1]+0.1,(1-surv)*n[t,m-1]+0.1)
        dead[t,m]=n[t,m-1]-n[t,m]
      }
    
  }
  
  # prior for the proportion of territories with different number of wolves. Hierarchical model
  
  Np[1:15]~ddirich(alphaN[1:15])
  
  # prior for the survival probability over the winter
  
  surv~dbeta(2,2)
  
  # Interval censoring for the number of dead in each territory. Observation gives the lower bound.
  
  for(i in 1:n_obs){
    I[i]~dinterval(dead[terr[i],month[i]],D[i])
  }
  
  
  ## observation model for the DNA samples: custom likelihood using the "ones-trick".
  ## x[i] = number of different wolves observed
  ## k[i] = number of (successful) DNA samples collected
  
  for(i in 1:n_obs){
    N[i]=n[terr[i],month[i]]
    Nstar[i]=ifelse(N[i]<x[i],x[i],N[i])
    L[i]=ifelse(N[i]>=x[i],exp(logfact(Nstar[i])-logfact(Nstar[i]-x[i])+k[i]*log(1/Nstar[i])),0)
    Lstar[i]=ifelse(missing[i]==0,L[i],1)
    z[i]~dbern(Lstar[i])
  }
  
  
  ### Observation model for the highest wolf count observed (max[i])
  # assuming linear dependency from the number of wolves in the territory
  # priors for model parameters come from a separate analysis of old tassu data & estimated pack sizes
  
  for(i in 1:n_obs){
    max[i]~dnorm(alpha[month[i]]*N[i],pow(sigma[month[i]]/sqrt(Sample[i]+0.1),-2))
  }
  
  alpha[1]=par[2]
  alpha[2]=par[3]
  sigma[1]=par[9]
  sigma[2]=par[10]
  
  ### Observation model the proportion of pair observations out of all observations. Priors for model parameters come from a separate
  ## analysis of old data & estimates
  
  for(i in 1:n_obs){
    zi[i]=exp(a[month[i]]-beta[month[i]]*(N[i]-2)+sd[month[i]]*eps[i])
    p[i]=zi[i]/(1+zi[i])
    pairs[i]~dbin(p[i],Sample[i])
    eps[i]~dnorm(0,1)
  }
  
  a[1]=par[1]
  a[2]=par[6]
  beta[1]=par[4]
  beta[2]=par[5]
  sd[1]=par[7]
  sd[2]=par[8]
  
  ## Posterior distribution of previous analysis used as a prior in this analysis.
  
  par[1:10]~dmnorm(mu,tau)
  
}
```
