## Supplementary material for "Assessment of the residential Finnish wolf population combines DNA captures, citizen observations and mortality data using a Bayesian state-space model": Code, data and all results: lukecolors.r

```
LukeOrange=function(a){
  return(rgb(255,130,0,alpha=a,maxColorValue=255))
}

# musta  R84 G88 B90

LukeBlack=function(a){
  return(rgb(84,88,90,maxColorValue=255,alpha=a))
}

# sininen R0 G181 B226

LukeBlue=function(a){
  return(rgb(0,181,226,alpha=a,maxColorValue=255))
}

# tumma sininen  R0 G51 B160

LukeDarkBlue=function(a){
  return(rgb(0,51,160,maxColorValue=255,alpha=a))
}

# Vihre�  R120 G190 B32

LukeGreen=function(a){
  return(rgb(120,190,32,maxColorValue=255,alpha=a))
}
```
