## Supplementary material for "Assessment of the residential Finnish wolf population combines DNA captures, citizen observations and mortality data using a Bayesian state-space model": Code, data and all results: plots.r

```
## This script produces output plots based on MCMC simulation

## Load colors to be used

source("code/lukecolors.r")

d=as.matrix(chains) # Format MCMC chains as a matrix

## set target folder for the plots

if(month==2){
  plot=1
  folder="March"
}      # plot March
if(month==1){
  plot=0 
  folder="Nov"
  
}     # plot Nov

# Plot posterior distribution for each territory
exp=c()

plot(x=c(0,13),y=c(0,13),col="white",xlab="Pack size (Expert)",ylab="Pack size (model)")

for(i in 1:data$n_obs){
  z=d[,paste0("N[",i,"]")]
  e=runif(length(d[,1]),obs[i,"Expert_min"]-0.1,obs[i,"Expert_max"]+0.1)
  expert[i]=mean(e)
  if(i%%2 == 0) pcol="grey"
  if(i%%2 == 1) pcol="black"
  points(mean(e[1:100]),mean(z[1:100]),pch=19,cex=2,col=pcol)
  p=c()
  n=length(z)
  CV=sd(z) #/mean(z)
  if(CV<0.6) col=LukeBlue(255)
  if(CV>0.6) col=LukeBlue(255)
  if(CV>1) col=LukeBlue(255)
  if(plot==0){
    
    png(file=paste0("results/",folder,"/",lang,"/",folder,"_",lang,"_",names[i],".png"),width=1400,height=1400/3)
    par(mfrow=c(1,2),cex=3,mar=c(4,4,1,1))
    
    if(lang=="FI"){
      xlab="Yksilömäärä"
      ylab="Todennäköisyys"
    }
    if(lang=="SWE"){
      xlab="Antal"
      ylab="Sannolikheten"
    }
    if(lang=="ENG"){
      xlab="Number of indidivuals"
      ylab="Probability"
    }
    
    
    
    barplot(table(c(1:min(z),z))/length(z),main="",col=col,border=col,xlab=xlab,ylab=ylab,ylim=c(0,1))
    
    p[1]=sum(z==1)/n
    p[2]=sum(z==2)/n
    p[3]=sum(z>2)/n
    v=var(p)
    if(v>0.2) col=LukeGreen(255)
    if(v<0.2) col=LukeOrange(255)
    if(v<0.1) col="red"
    
    if(lang=="FI"){
      xlab="Reviirin tila"
      ylab="Todennäköisyys"
      leg=c("Yksittäinen","Pari","Lauma")
    }
    if(lang=="SWE"){
      xlab="Revirstatus"
      ylab="Sannolikheten"
      leg=c("Ensam","Par","Flock")
    }
    if(lang=="ENG"){
      xlab="Territory status"
      ylab="Probability"
      leg=c("Empty","Pair","Pack")
    }
    
    barplot(p,names.arg = leg,main="",col=col,border=col,xlab=xlab,ylab=ylab,ylim=c(0,1))
    
    dev.off()
    if(i<3) probs=c(names[i],max(p)*100)             # one of the first two: 1 or 2 occur 
    else probs=rbind(probs,c(names[i],max(p)*100))
    plot=1
  }
  else plot=0
  
  lines(x=c(0,13),y=c(0,13),col="red",lwd=2)
  legend("topleft",c("Autumn","Spring"),col=c("black","grey"),cex=c(1,1),pch=c(19,19))
}


### Plot about number of individuals in territories and survival from autumn to spring
col=LukeBlue(255)

png(file=paste0("results/",folder,"/",lang,"/",folder,"_",lang,"_survival.png"),width=1400,height=1400/3)
par(mfrow=c(1,3),cex=1.5)

x_a=seq(from=1, to=data$n_obs,by=2)
x_s=x_a+1

z4=d[,paste0("Total[",1,"]")]
barplot(table(z4)/length(z4),main="Individuals in autumn",col=col,border=col,xlab="Number of individuals",ylab="Probability")
plot(density(d[,"surv"]),col=LukeBlue(255),main="Survival",xlab="Survival rate",ylab="Probability",lwd=3)
sum(expert[x_a])
sum(expert[x_s])


z5=d[,paste0("Total[",2,"]")]
barplot(table(z5)/length(z5),main="Individuals in spring",col=col,border=col,xlab="Number of individuals",ylab="Probability")
dev.off()

### Plot about total number of packs, pairs and empty territories and their posterior correlation

png(file=paste0("results/",folder,"/",lang,"/",folder,"_",lang,"_territorytotals.png"),width=1400,height=1400)

z1=d[,paste0("Packs[",month,"]")]

z2=d[,paste0("Pairs[",month,"]")]

z3=d[,paste0("Singles[",month,"]")]

x=data.frame(Packs=rnorm(length(z1),z1,0.1),Pairs=rnorm(length(z2),z2,0.1),Empties=rnorm(length(z3),z3,0.1))

par(mfrow=c(3,3),cex=1.5)
barplot(table(z1)/length(z1),main="Packs",col=col,border=col,xlab="Number of packs",ylab="Probability")
plot(x$Pairs,x$Packs,col=LukeBlue(20),pch=19,ylab="Packs",xlab="Pairs")
plot(x$Empties,x$Packs,col=LukeBlue(20),pch=19,ylab="Packs",xlab="Empties")

plot(x$Packs,x$Pairs,col=LukeBlue(20),pch=19,ylab="Pairs",xlab="Packs")
barplot(table(z2)/length(z2),main="Pairs",col=col,border=col,xlab="Number of pairs",ylab="Probability")
plot(x$Empties,x$Pairs,col=LukeBlue(20),pch=19,ylab="Packs",xlab="Empties")

plot(x$Packs,x$Empties,col=LukeBlue(20),pch=19,ylab="Empties",xlab="Packs")
plot(x$Pairs,x$Empties,col=LukeBlue(20),pch=19,ylab="Empties",xlab="Pairs")
barplot(table(z3)/length(z3),main="Empty",col=col,border=col,xlab="Number of empties",ylab="Probability")

dev.off()
```
