## Supplementary material for "Assessment of the residential Finnish wolf population combines DNA captures, citizen observations and mortality data using a Bayesian state-space model": Code, data and all results: main_script_assessment.R

```
### This script runs the entire analysis of the Bayesian state-space model. It loads the data, runs MCMC simulation and produces plots of results


setwd("c:/models/Online_Resource_1_code_results")  # set the working directory. Change this to match  your system

source("Old_Tassu_model.r") # run the model for traing data first. This creates prior distributions for linear and logistic regression
                            # model parameters

## Load required packages
require(rjags)
require(readxl)
require(MASS)

# Point to the JAGS model file

modelfile=("code/jags_model.r")

# Run the data preparation code

source("code/data_prep.r")

# Compile the JAGS model for the data set

jm=jags.model(modelfile,data,inits,n.chains=2)

# Run initial burn-in for the MCMC

chains=coda.samples(jm,c("N","surv","Total","Singles","Pairs","Packs","n","dead","par"),n.iter=10000)

# Run the model for final inference

chains=coda.samples(jm,c("surv","Total","Singles","Pairs","Packs","dead","par","n","N"),n.iter=1000000,thin=1000)

# Create plots from MCMC output
## Plot chains of all variables to assess convergence

pdf("results/mcmc.pdf")

plot(chains)

dev.off()

## Create plots of autumn estimates:
month=1
lang="ENG"
source("code/plots.r")

## Create plots of spring estimates:
month=2
lang="ENG"
source("code/plots.r")
```
