## Supplementary material for "Assessment of the residential Finnish wolf population combines DNA captures, citizen observations and mortality data using a Bayesian state-space model": Code, data and all results: Old_Tassu_model.r

```
require(rjags)
require(readxl)

model3<-"
model{

for(i in 1:N){

XA[i,1:2]~dmulti(pA[i,1:2],TA[i])                  # Pair and pack observations in Autumn  
XS[i,1:2]~dmulti(pS[i,1:2],TS[i])                  # Pair and pack observations in Spring  

# Logistic regression for the proportion of pair observations as a function of the number of wolves in the territory in Autumn

pA[i,1]=exp(a-(NAt[i]-2)*betaA+sdA*eA[i])/(1+exp(a-(NAt[i]-2)*betaA+sdA*eA[i]))
pA[i,2]=1-pA[i,1]

# Logistic regression for the proportion of pair observations as a function of the number of wolves in the territory in Spring

pS[i,1]=exp(s-(NS[i]-2)*betaS+sdS*eS[i])/(1+exp(s-(NS[i]-2)*betaS+sdS*eS[i]))
pS[i,2]=1-pS[i,1]

# Prior for the number of wolves in spring based on the expert estimates

NS[i]~dunif(minS[i],maxS[i]+0.2)

# Prior for the number of wolwes in previous autumn based on the number of wolves in spring

NAt[i]=(1+u[i])*NS[i]  # more wolves in autumn than in spring
u[i]~dbeta(2,8)

# Territory-specific random effects for the logistic regression

eA[i]~dnorm(0,1)
eS[i]~dnorm(0,1)

# Linear model for the highest pack size observation as a function of the number of wolves in the territory

maxobsA[i]~dnorm(alphaA*NAt[i],tauA[i])                 # Autumn
tauA[i]=pow(sigmaA/sqrt(sum(XA[i,1:2])+1),-2)           # Precision depends on the total number of observations made

maxobsS[i]~dnorm(alphaS*NS[i],tauS[i])                  # Spring
tauS[i]=pow(sigmaS/sqrt(sum(XS[i,1:2])+1),-2)           # Precision depends on the total number of observations made
}

## Minimally informative priors for model parameters

betaA~dnorm(0,0.001)T(0,)       # Relationship between number of wolves in territory and proportion of pair observations is assumed
betaS~dnorm(0,0.001)T(0,)       # to be negative a priori (minus sign is in the logistic regression already)

sdA~dunif(0,2)                 # SD of territory specific random effects in Autumn
sdS~dunif(0,2)                 # SD of territory specific random effects in Spring
 
a~dnorm(0,0.1)                 # Intercept for the logistic regression: logit-proportion of pair observations when pack size =2 in autumn
s~dnorm(0,0.1)                 # Intercept for the logistic regression: logit-proportion of pair observations when pack size =2 in spring

alphaA~dnorm(0,0.01)T(0,)      # Relationship between number of wolves in territory and highest observed pack size is assumed
alphaS~dnorm(0,0.01)T(0,)      # to be positive a priori

sigmaS~dunif(0,10)             # Residual SD of maximum pack size observations in spring
sigmaA~dunif(0,10)             # Residual SD of maximum pack size observations in autumn
}

"

#setwd("c:/models/wolf_publ")                    # set working directory to match your system

dat=read_xlsx("data/old_tassu.xlsx")            # Read the data set from Excel file
dat=as.matrix(dat)

data=list(
  N=length(dat[,1]),                           # Number of observations
  XA=dat[,3:4],                                # Pair and pack observations in autumn
  XS=dat[,5:6],                                # Pair and pack observations in spring
  minS=dat[,1],                                # minimum pack size according to expert estimate
  maxS=dat[,2],                                # maximum pack size according to expert estimate
  TA=dat[,3]+dat[,4],                          # Total number of pack & pair observations in autumn
  TS=dat[,5]+dat[,6],                          # Total number of pack & pair observations in spring
  maxobsA=dat[,7],                             # maximum pack observation in autumn
  maxobsS=dat[,8]                              # maximum pack observation in spring
)


## Run the JAGS model

jm=jags.model(textConnection(model3),data,n.chain=2)
chains=coda.samples(jm,c("alphaA","alphaS","sigmaS","sigmaA","betaA","betaS","sdS","sdA","a","s"),n.iter=10000)

## Plot the chains for visual inspection

plot(chains)

## Calculate posteriror means and covariance matrix, save to disk to be used as priors in population assessment

d<-as.matrix(chains)

dput(colMeans(d),file="results/prior_means")
dput(cov(d),file="results/prior_covariance")

## Create a plot that shows the estimated logistic regression between pack size and the expected proportion of pair observations

png("results/packsize_vs_pair_obs.png",width=1500,height=700)
par(mfrow=c(1,2),cex=1.5)
x=2:15
for(k in 1:500){
  P=c()
  muP=c()
  b=c()
  betaS=d[k,"betaS"] 
  sdS=d[k,"sdS"] 
  s=d[k,"s"] 
  
  for(i in 2:15){
    b[i]=rnorm(1,0,sdS)
    P[i]=exp(s-betaS*(i-2)+b[i])/(1+exp(s-betaS*(i-2)+b[i]))
    muP[i]=exp(s-betaS*(i-2))/(1+exp(s-betaS*(i-2)))
    
  }
  if(k==1) plot(x,muP[2:15],type="l",ylim=c(0,1),xlim=c(2,15),col=rgb(1,0,0,alpha=0.1),lwd=5,xlab="Pack size",ylab="Proportion of pair observations")
  else points(x,muP[2:15],type="l",col=rgb(1,0,0,alpha=0.1),lwd=5)
  points(x,P[2:15],pch=15,col=rgb(0,0,0,alpha=0.1))
}


## Overlay the data points

N=(data$XS[,1]+data$XS[,2])>10
N2=(data$XS[,1]+data$XS[,2])>20
points((data$minS+data$maxS)/2,(data$XS[,1])/(data$XS[,1]+data$XS[,2]),pch=19,cex=1+N+N2)

legend("topright",legend=c("Expected value","Predictive density","Data"),lwd=c(5,NA,NA),col=c("red","grey","black"),pch=c(NA,15,19))


### Create a plot that shows the estimated linear regression between pack size and the expected highest pack observation

x=1:10
for(k in 1:500){
alphaS=d[k,"alphaS"]
sigmaS=d[k,"sigmaS"]
mu=c()
obs=c()
for(j in 1:10){
  mu[j]=alphaS*x[j]
  obs[j]=rnorm(1,mu[j],sigmaS/sqrt(20))
}

if(k==1) plot(x,mu,type="l",col=rgb(1,0,0,alpha=0.1),xlab="Pack size",ylab="Maximum pack observation")
else points(x,mu,type="l",col=rgb(1,0,0,alpha=0.1))
points(x,obs,pch=15,col=rgb(0,0,0,alpha=0.1))
}

points((data$maxS+data$minS)/2,data$maxobsS,cex=1+N+N2,xlim=c(0,10),ylim=c(0,10),pch=19)
legend("topleft",legend=c("Expected value","Predictive density","Data"),lwd=c(5,NA,NA),col=c("red","grey","black"),pch=c(NA,15,19))

dev.off()
```
