## Supplementary material for "Assessment of the residential Finnish wolf population combines DNA captures, citizen observations and mortality data using a Bayesian state-space model": Code, data and all results: Read Me.txt

This folder includes all the material necessary to run the example analysis presented in 

Mäntyniemi, S. Helle, I. and Kojola, I. (2022) Assessment of the Finnish wolf population combines DNA captures, 
citizen observations and mortality data using a Bayesian state-space model. European Journal of Wildlife Management.

Software needed:

R 3.6.0 or later
JAGS 4.3.0 or later 

- Save this folder structure to your computer
- The entire analysis can be conducted by running "main_script_assessment.R" in R. Edit the script first by changing the working directory to match your system.
- You may need to install some required packages for R: follow the potential error messages.
- Results can be found from the "results" folder.
