## Supplementary material for "Assessment of the residential Finnish wolf population combines DNA captures, citizen observations and mortality data using a Bayesian state-space model": Code, data and all results: mcmc.pdf

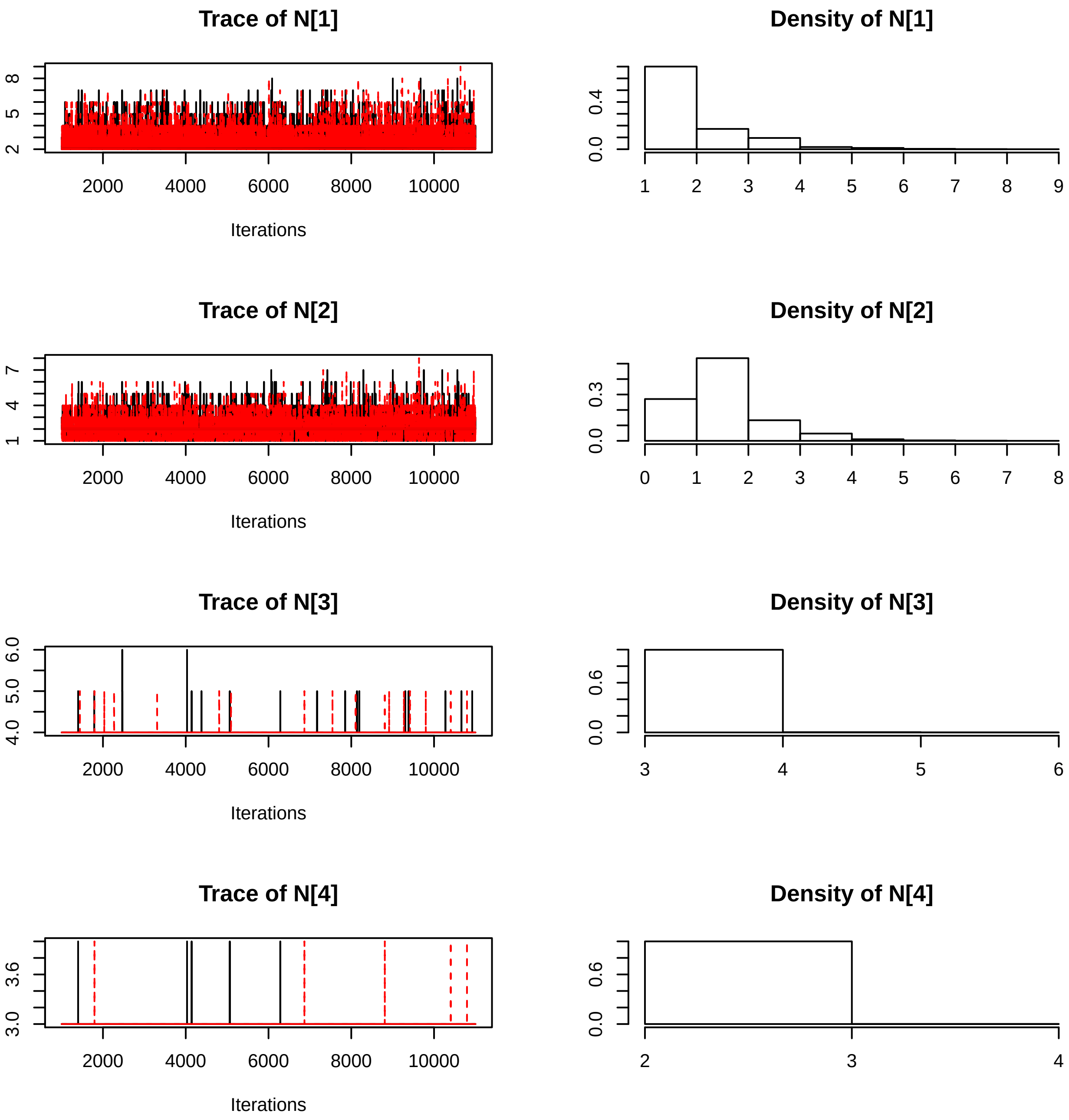
Trace of N[1]
Density of N[1]
Trace of N[2]
Density of N[2]
Trace of N[3]
Density of N[3]
Trace of N[4]
Density of N[4]
Iterations

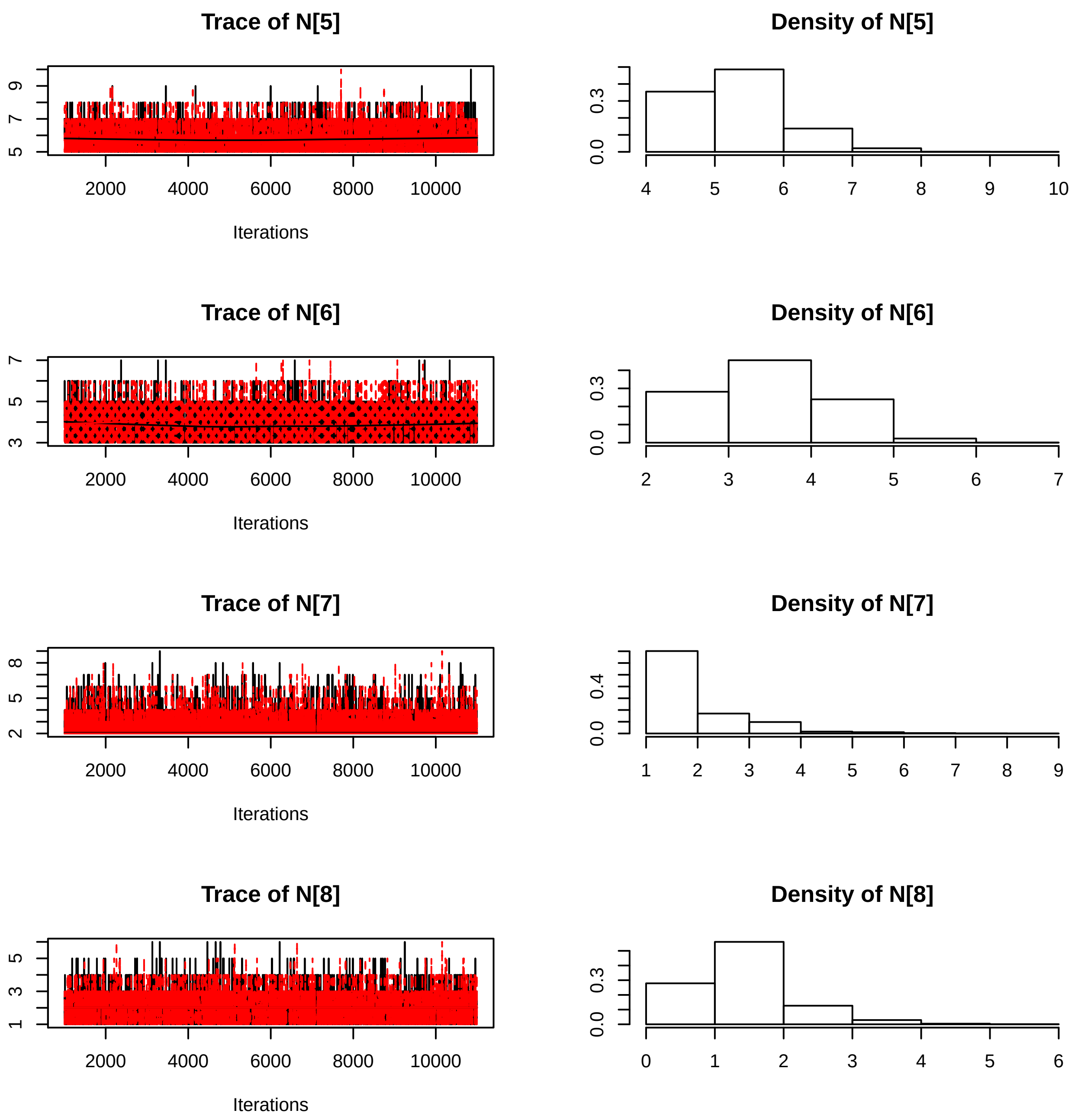
Trace of N[5]
9
7
5
2000
4000
6000
8000
10000
Iterations
Density of N[5]
0.3
0.0
4
5
6
7
8
9
10
Trace of N[6]
7
5
3
2000
4000
6000
8000
10000
Iterations
Density of N[6]
0.3
0.0
2
3
4
5
6
7
Trace of N[7]
8
5
2
2000
4000
6000
8000
10000
Iterations
Density of N[7]
0.4
0.0
1
2
3
4
5
6
7
8
9
Trace of N[8]
5
3
1
2000
4000
6000
8000
10000
Iterations
Density of N[8]
0.3
0.0
0
1
2
3
4
5
6

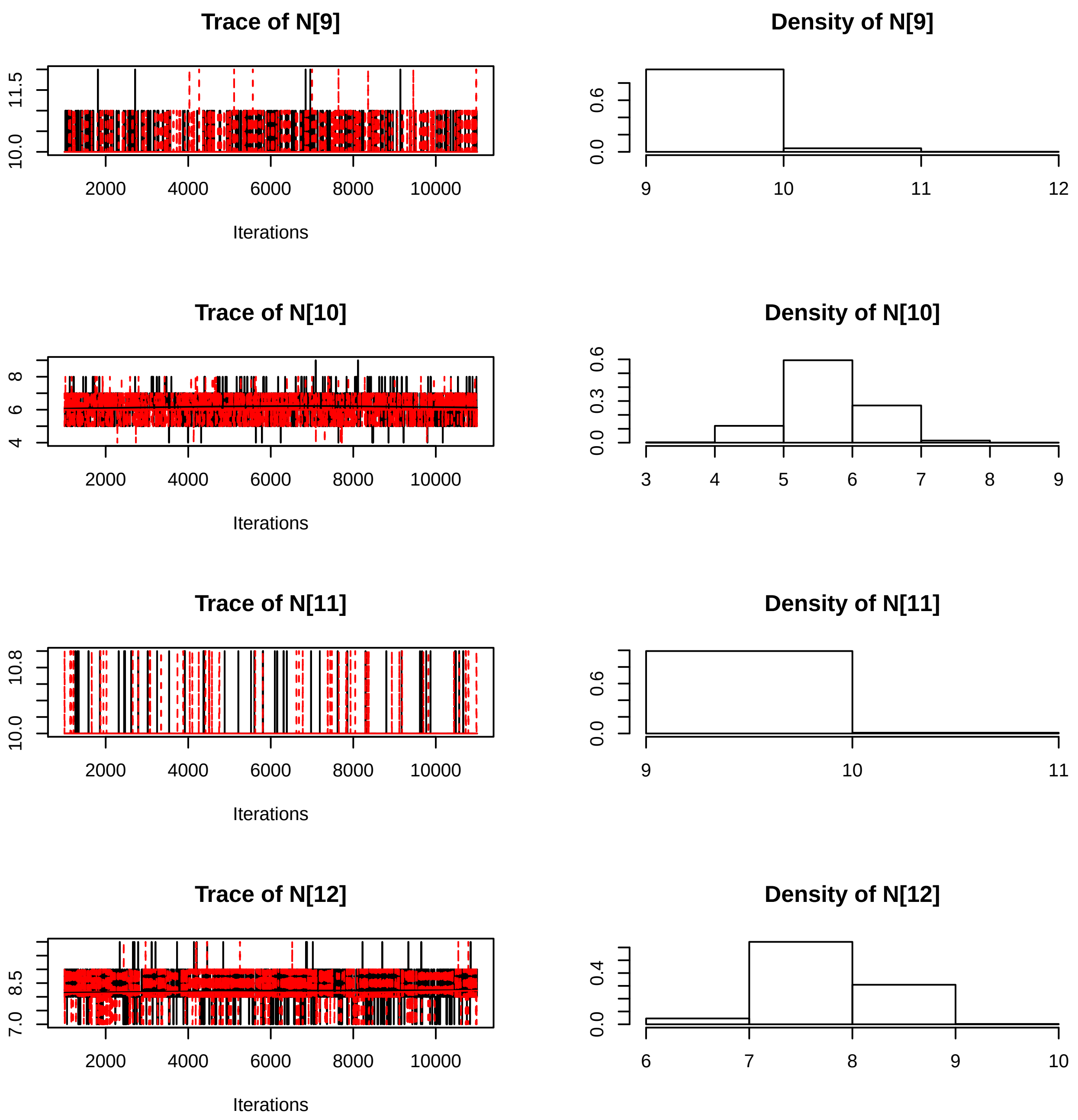
Trace of N[9]
11.5
10.0
2000 4000 6000 8000 10000
Iterations
Density of N[9]
0.6
0.0
9 10 11 12
Trace of N[10]
8
6
4
2000 4000 6000 8000 10000
Iterations
Density of N[10]
0.6
0.3
0.0
3 4 5 6 7 8 9
Trace of N[11]
10.8
10.0
2000 4000 6000 8000 10000
Iterations
Density of N[11]
0.6
0.0
9 10 11
Trace of N[12]
8.5
7.0
2000 4000 6000 8000 10000
Iterations
Density of N[12]
0.4
0.0
6 7 8 9 10

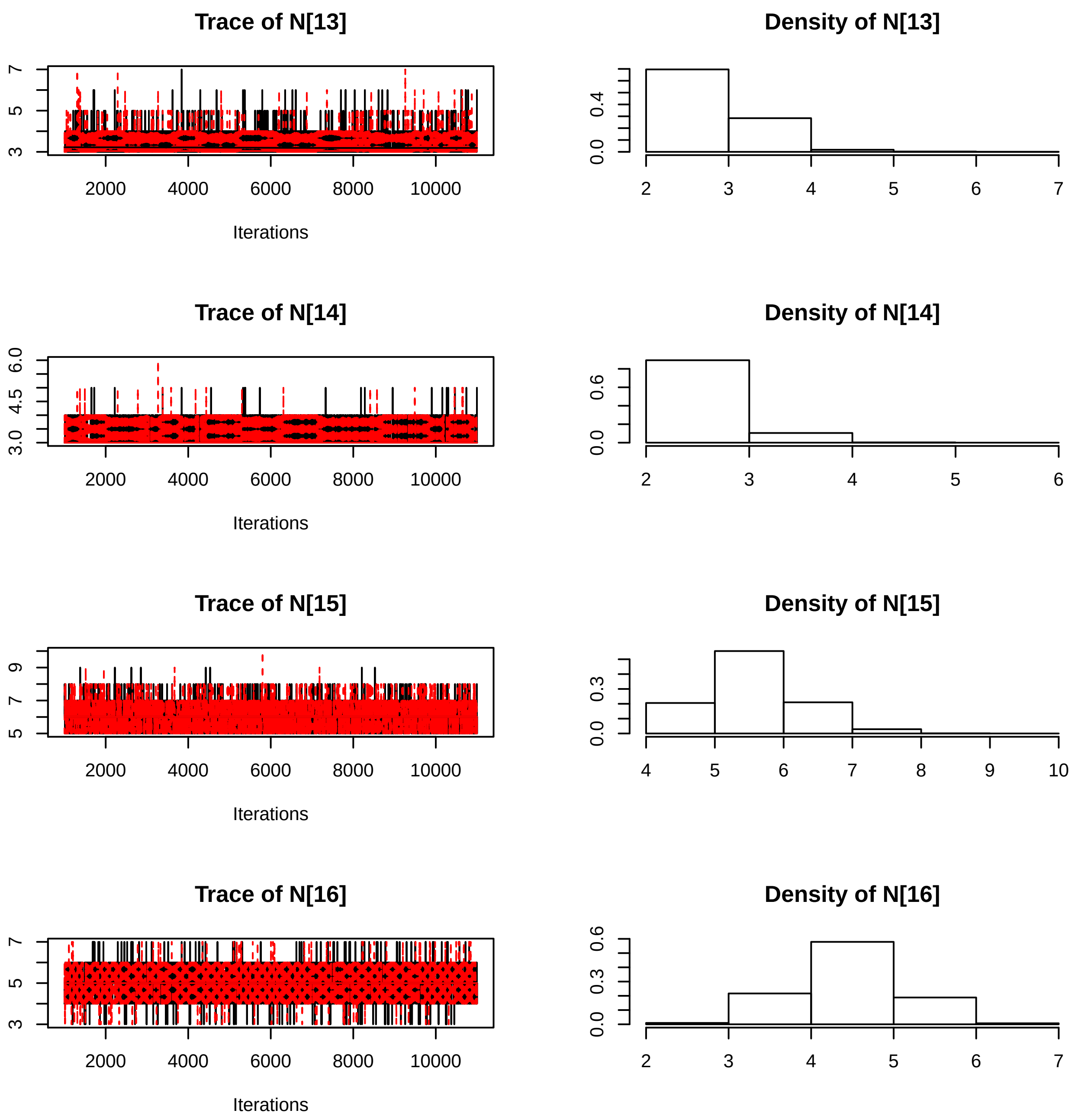
Trace of N[13]
Density of N[13]
Trace of N[14]
Density of N[14]
Trace of N[15]
Density of N[15]
Trace of N[16]
Density of N[16]
Iterations
2000
4000
6000
8000
10000

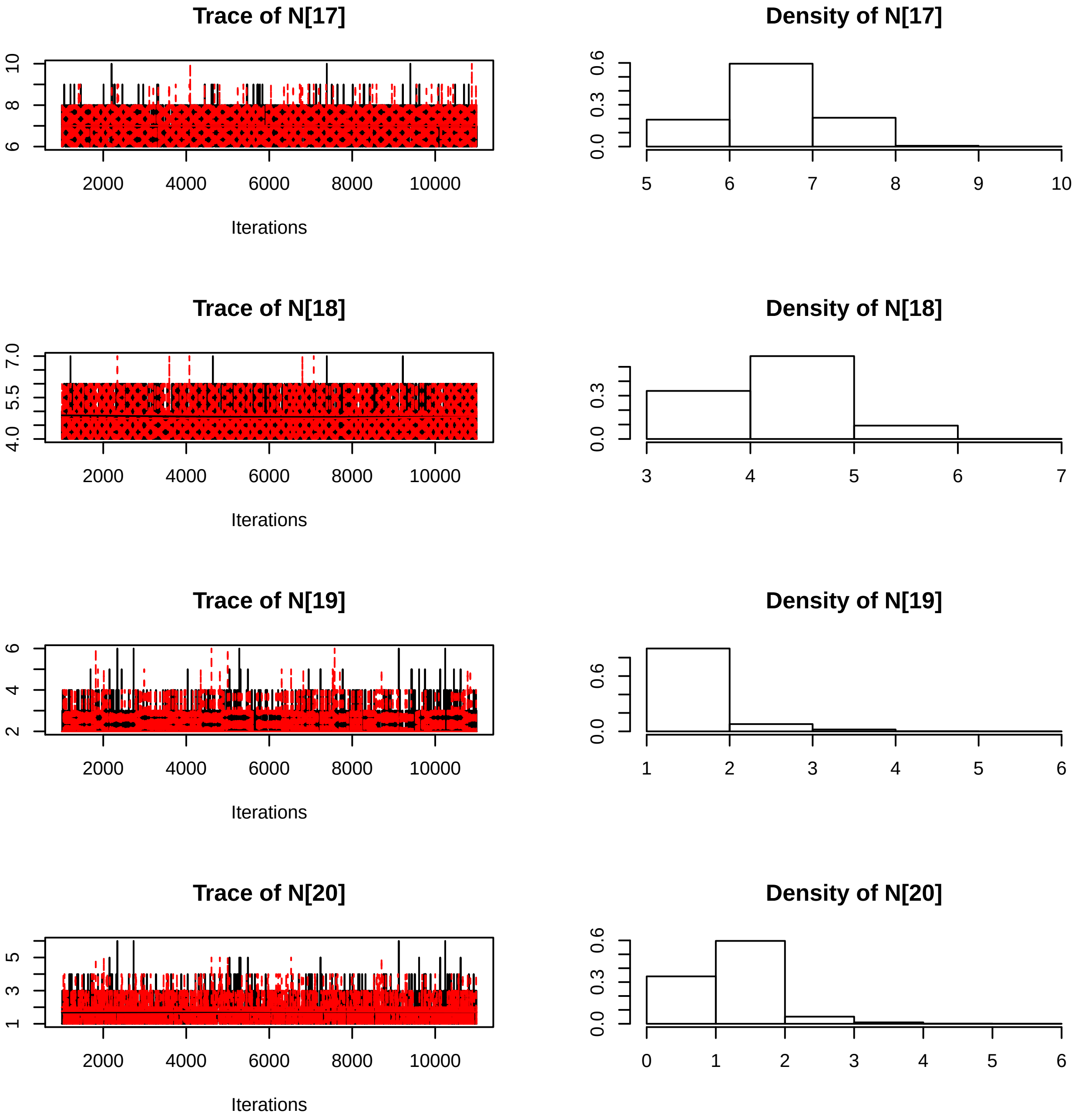

Trace of N[17]
Iterations
Density of N[17]
Trace of N[18]
Iterations
Density of N[18]
Trace of N[19]
Iterations
Density of N[19]
Trace of N[20]
Iterations
Density of N[20]

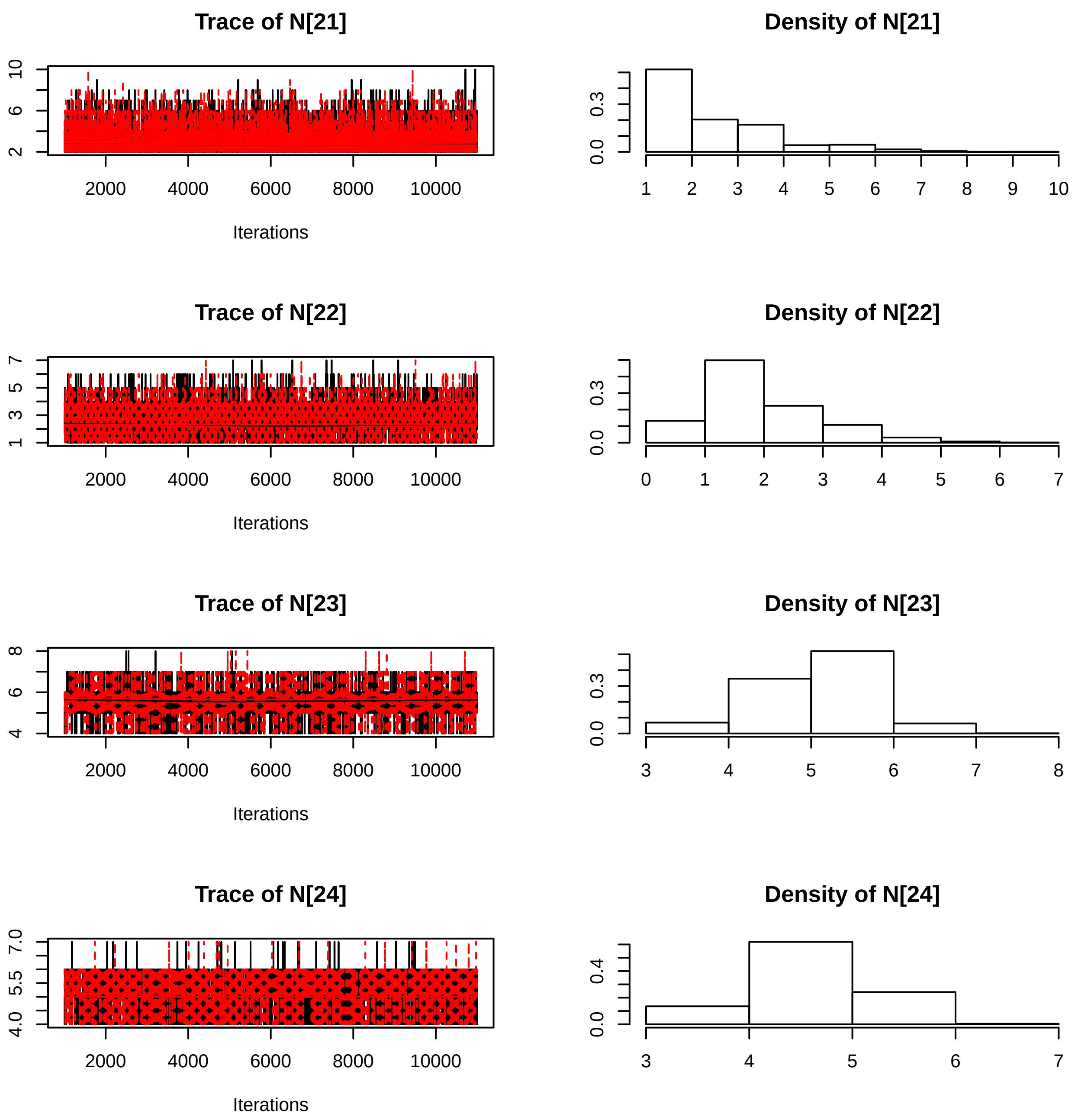
Trace of N[21]
10
6
2
2000
4000
6000
8000
10000
Iterations
Density of N[21]
0.3
0.0
1
2
3
4
5
6
7
8
9
10
Trace of N[22]
7
5
3
1
2000
4000
6000
8000
10000
Iterations
Density of N[22]
0.3
0.0
0
1
2
3
4
5
6
7
Trace of N[23]
8
6
4
2000
4000
6000
8000
10000
Iterations
Density of N[23]
0.3
0.0
3
4
5
6
7
8
Trace of N[24]
7.0
5.5
4.0
2000
4000
6000
8000
10000
Iterations
Density of N[24]
0.4
0.0
3
4
5
6
7

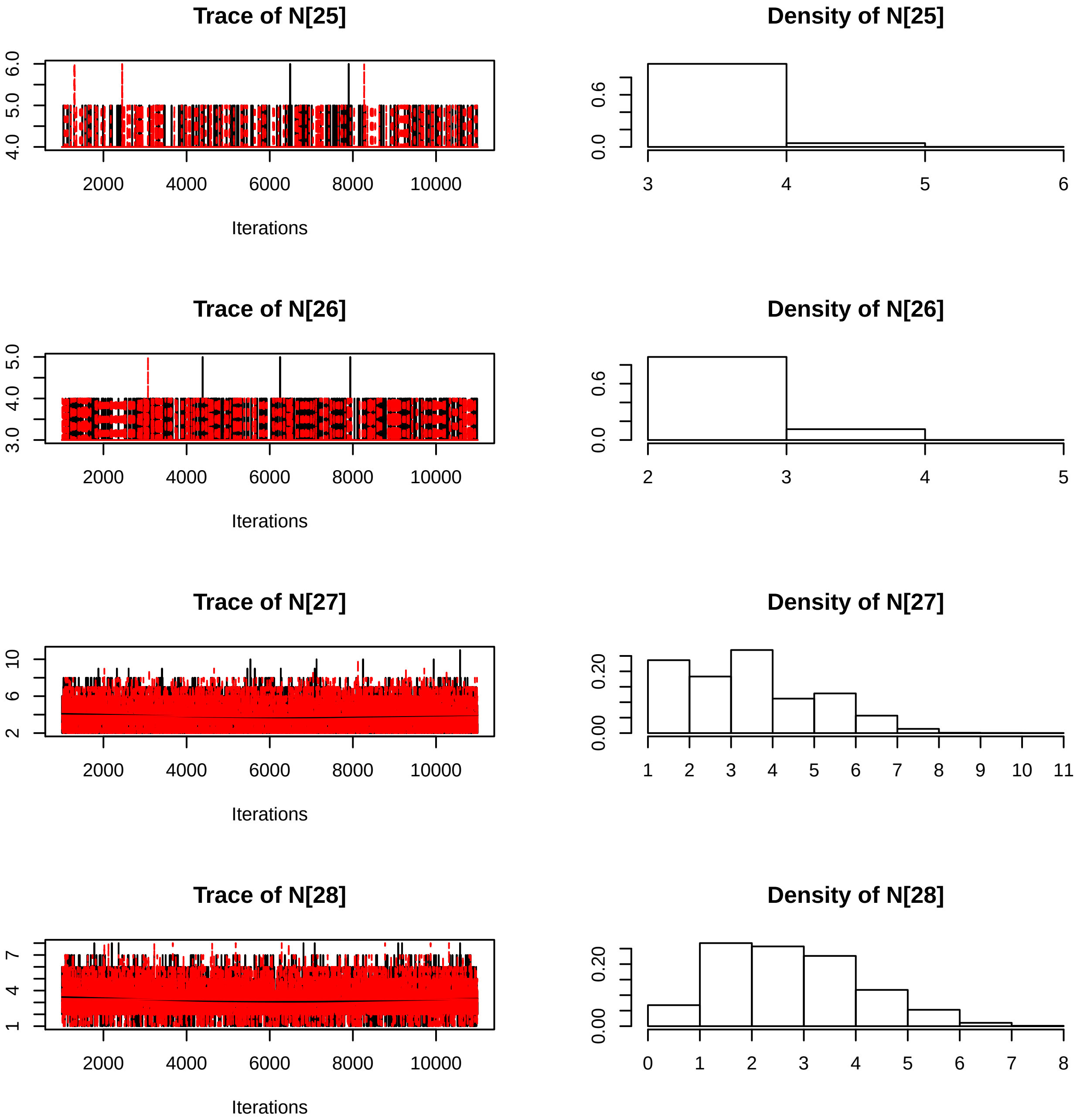

Trace of N[25]
Density of N[25]
Trace of N[26]
Density of N[26]
Trace of N[27]
Density of N[27]
Trace of N[28]
Density of N[28]
Iterations
2000
4000
6000
8000
10000

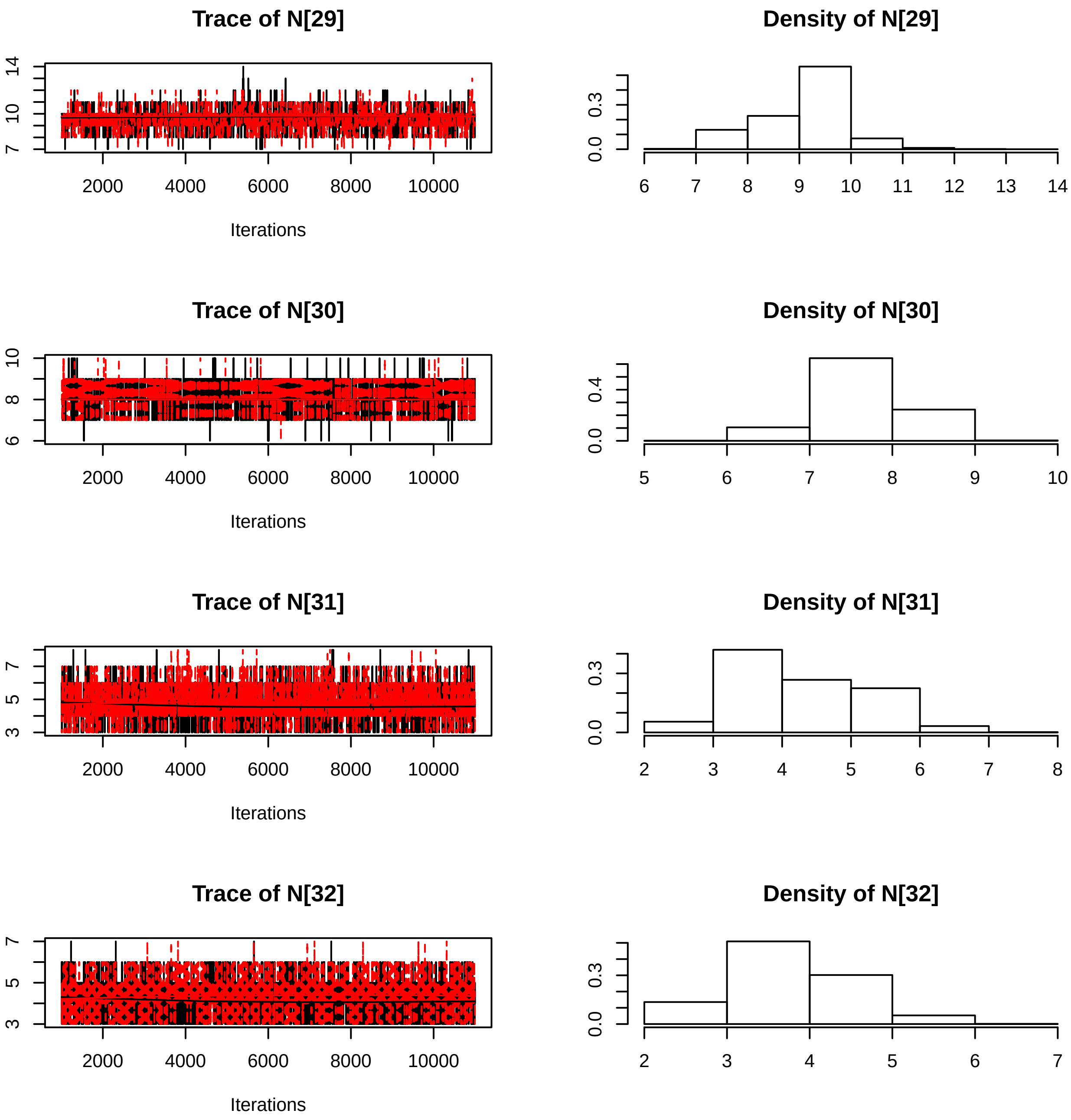
Trace of N[29]
Density of N[29]
Iterations
Trace of N[30]
Density of N[30]
Iterations
Trace of N[31]
Density of N[31]
Iterations
Trace of N[32]
Density of N[32]
Iterations
2000
4000
6000
8000
10000
0.0
0.3
0.4

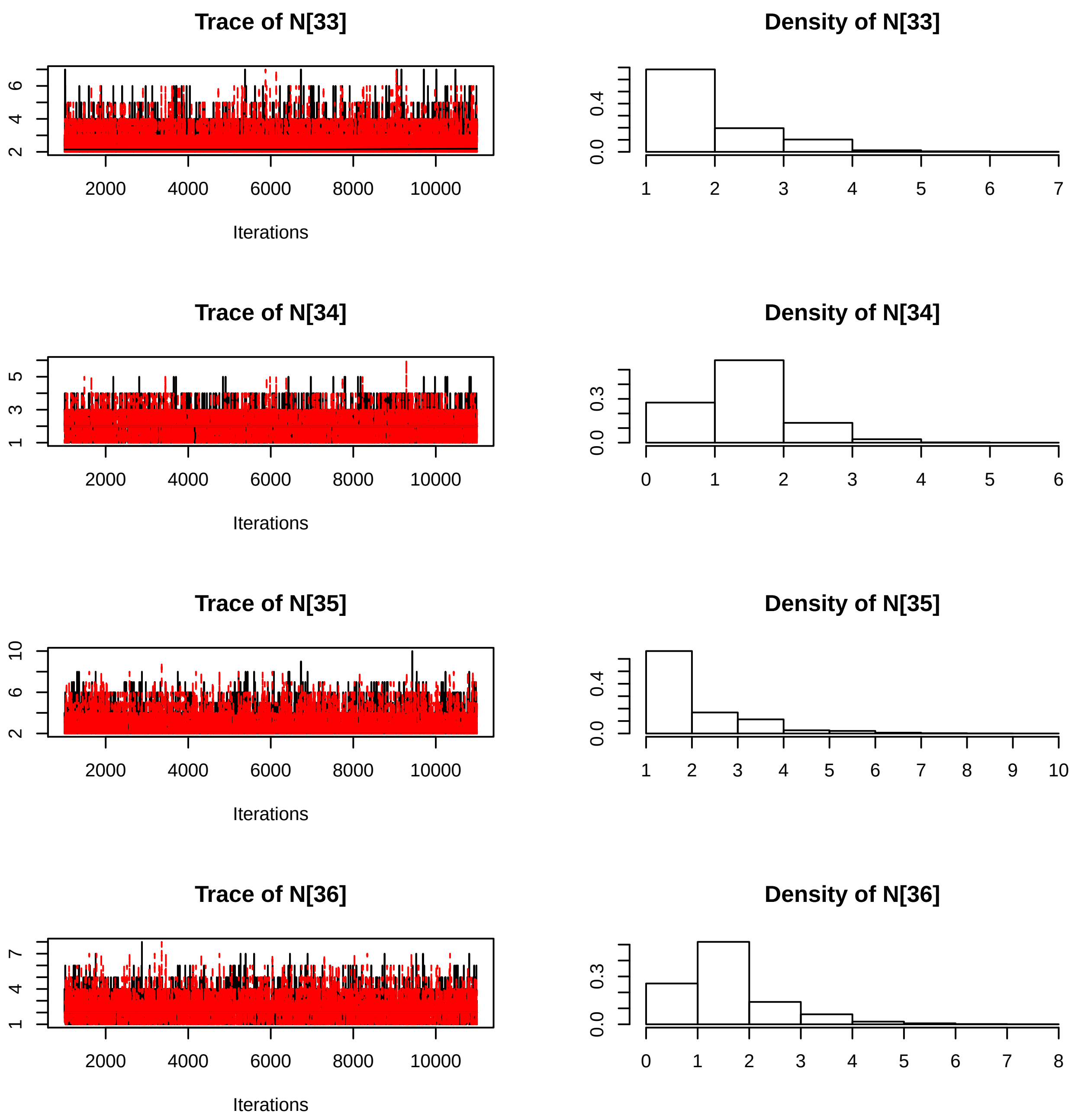

Trace of N[33]
2
4
6
2000
4000
6000
8000
10000
Iterations
Density of N[33]
0.0
0.4
1
2
3
4
5
6
7
Trace of N[34]
1
3
5
2000
4000
6000
8000
10000
Iterations
Density of N[34]
0.0
0.3
0
1
2
3
4
5
6
Trace of N[35]
2
6
10
2000
4000
6000
8000
10000
Iterations
Density of N[35]
0.0
0.4
1
2
3
4
5
6
7
8
9
10
Trace of N[36]
1
4
7
2000
4000
6000
8000
10000
Iterations
Density of N[36]
0.0
0.3
0
1
2
3
4
5
6
7
8

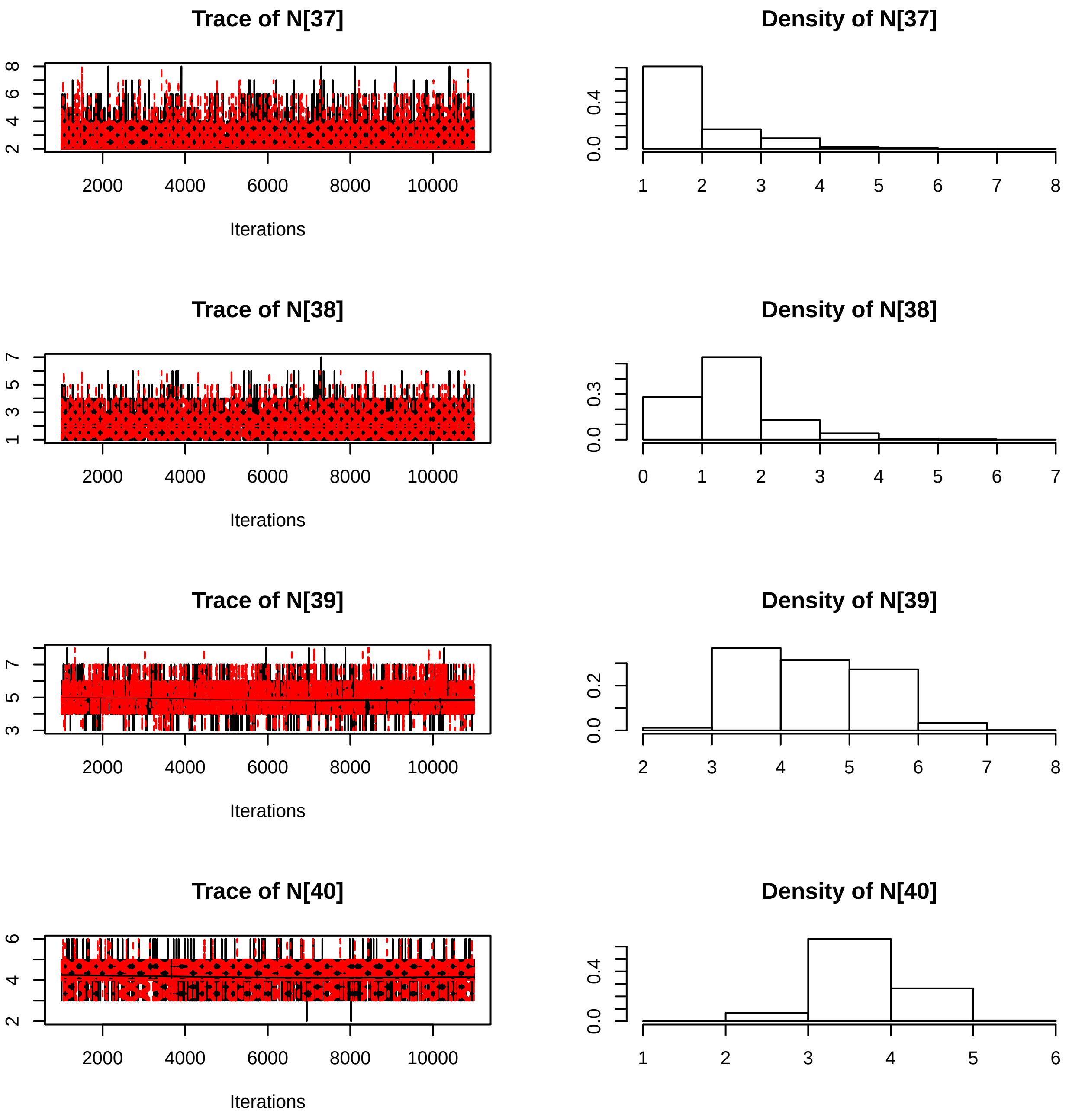
Trace of N[37]
Iterations
Density of N[37]
Trace of N[38]
Iterations
Density of N[38]
Trace of N[39]
Iterations
Density of N[39]
Trace of N[40]
Iterations
Density of N[40]

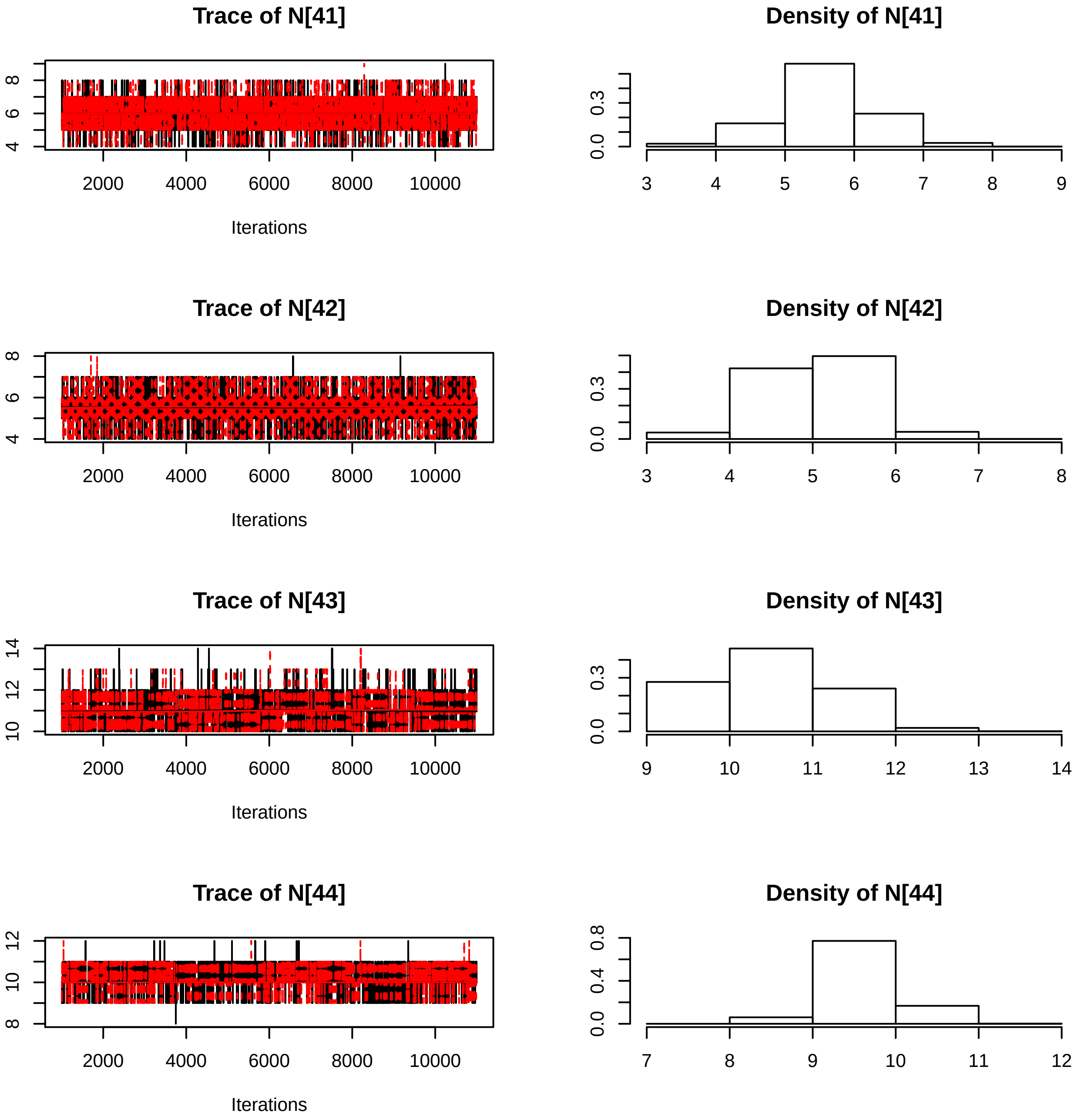

Trace of N[41]
Iterations
Density of N[41]
Trace of N[42]
Iterations
Density of N[42]
Trace of N[43]
Iterations
Density of N[43]
Trace of N[44]
Iterations
Density of N[44]

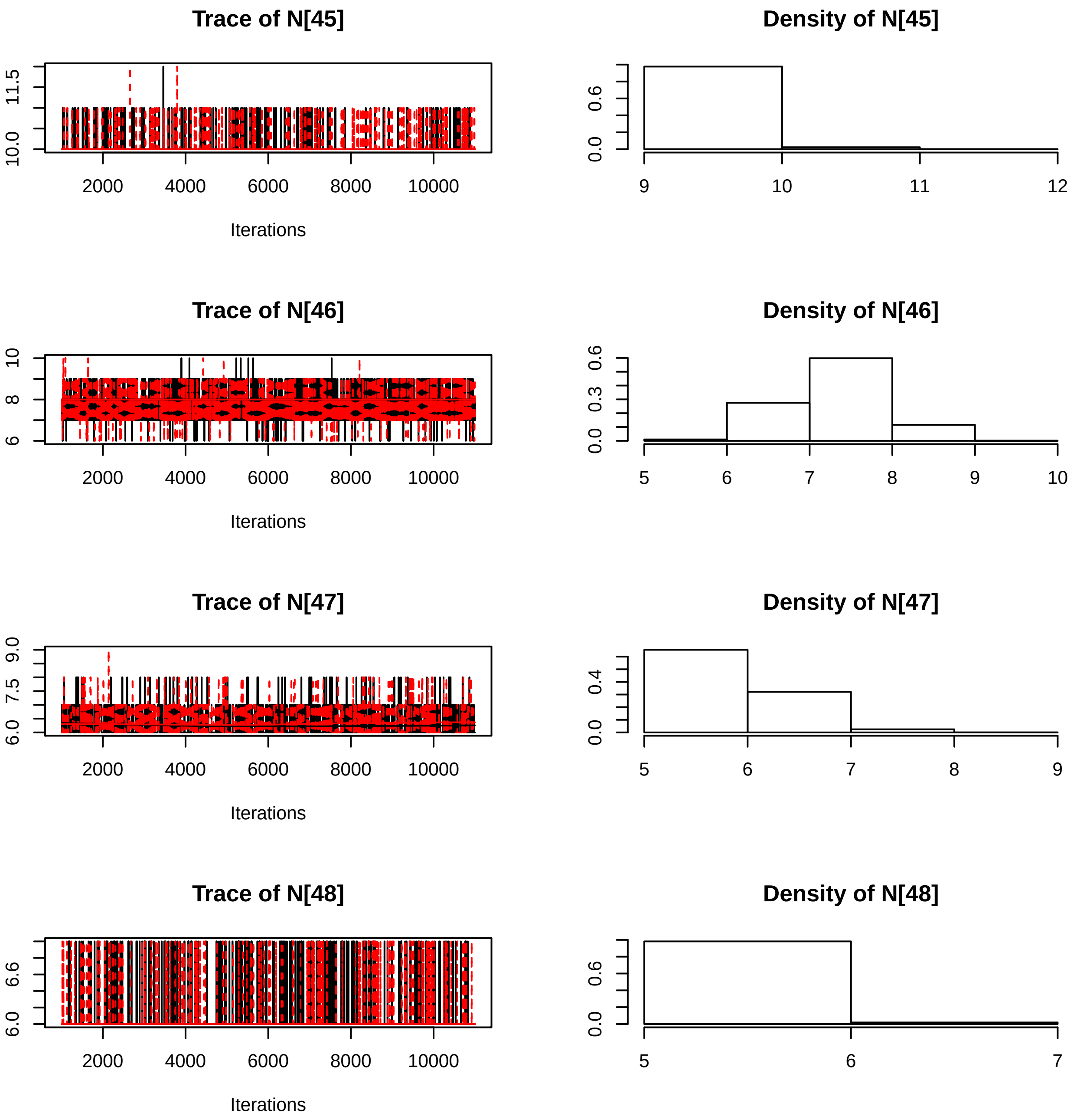
Trace of N[45]
Density of N[45]
Trace of N[46]
Density of N[46]
Trace of N[47]
Density of N[47]
Trace of N[48]
Density of N[48]
Iterations
2000
4000
6000
8000
10000
10.0
11.5
0.0
0.6
9
10
11
12
6
8
10
0.0
0.3
0.6
5
6
7
8
9
10
6.0
7.5
9.0
0.0
0.4
5
6
7
8
9
6.0
6.6
0.0
0.6
5
6
7

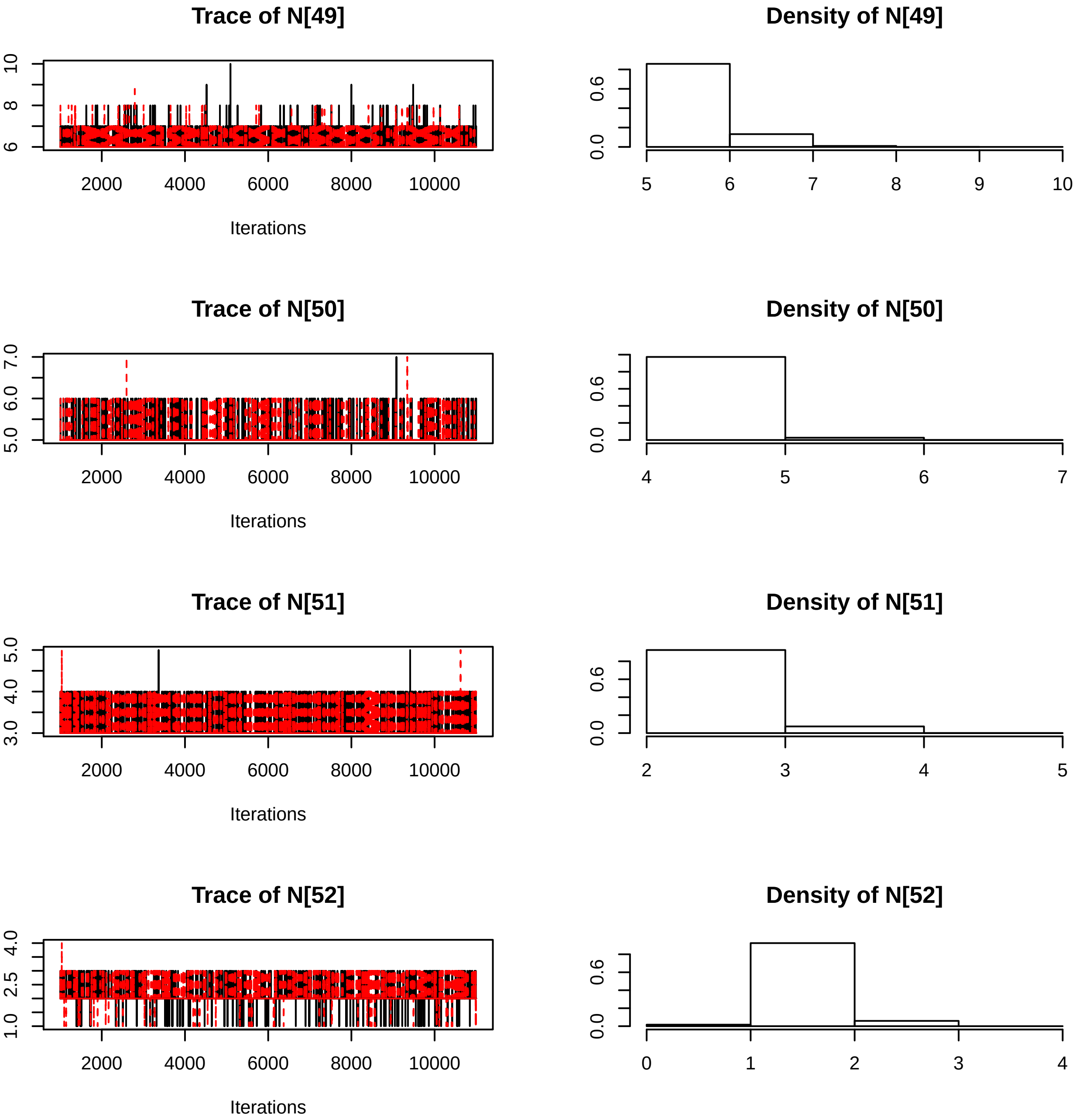
Trace of N[49]
Density of N[49]
Trace of N[50]
Density of N[50]
Trace of N[51]
Density of N[51]
Trace of N[52]
Density of N[52]
Iterations
Iterations
Iterations
Iterations
2000
4000
6000
8000
10000

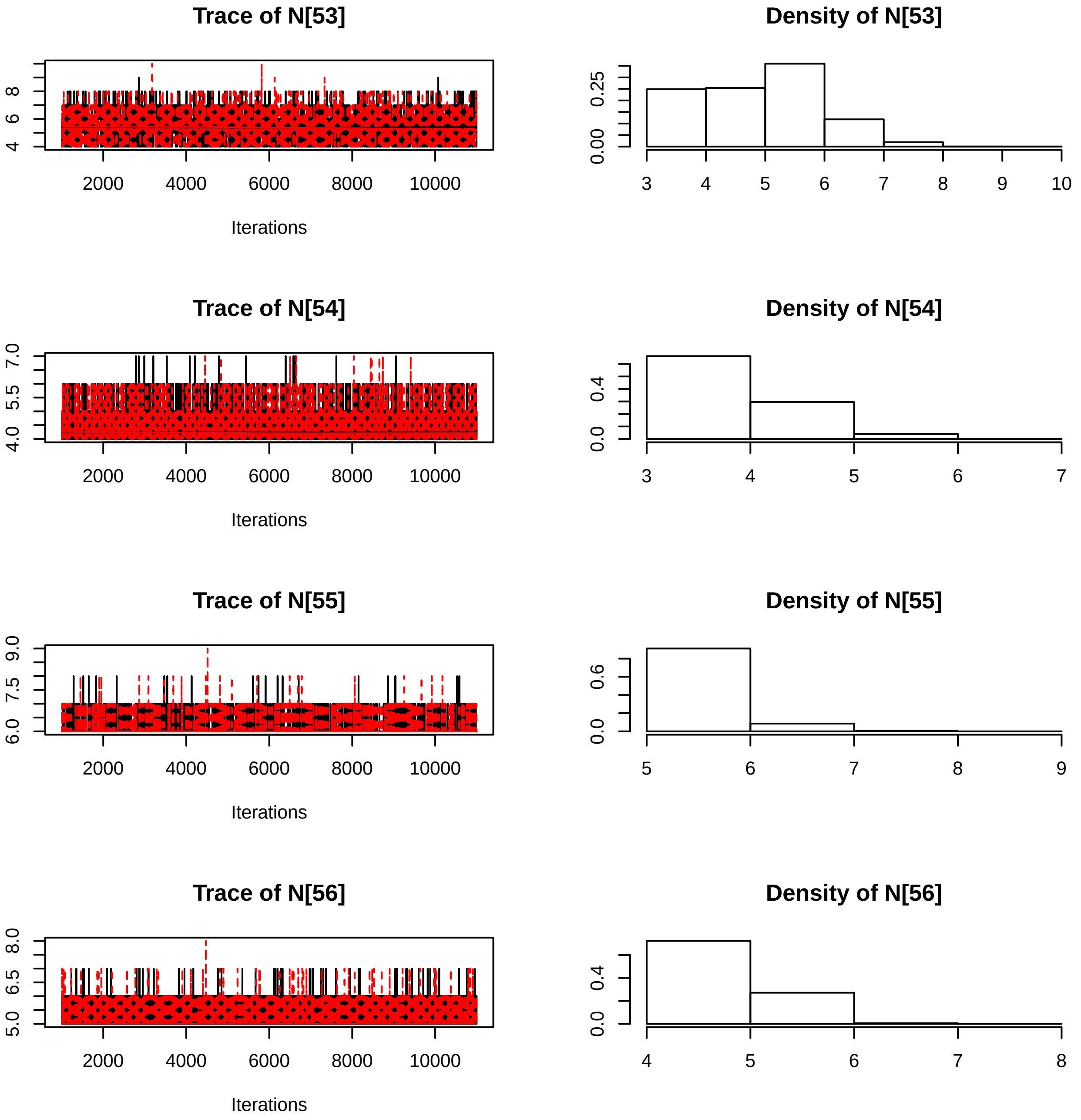

Trace of N[53]
Iterations
Density of N[53]
Trace of N[54]
Iterations
Density of N[54]
Trace of N[55]
Iterations
Density of N[55]
Trace of N[56]
Iterations
Density of N[56]

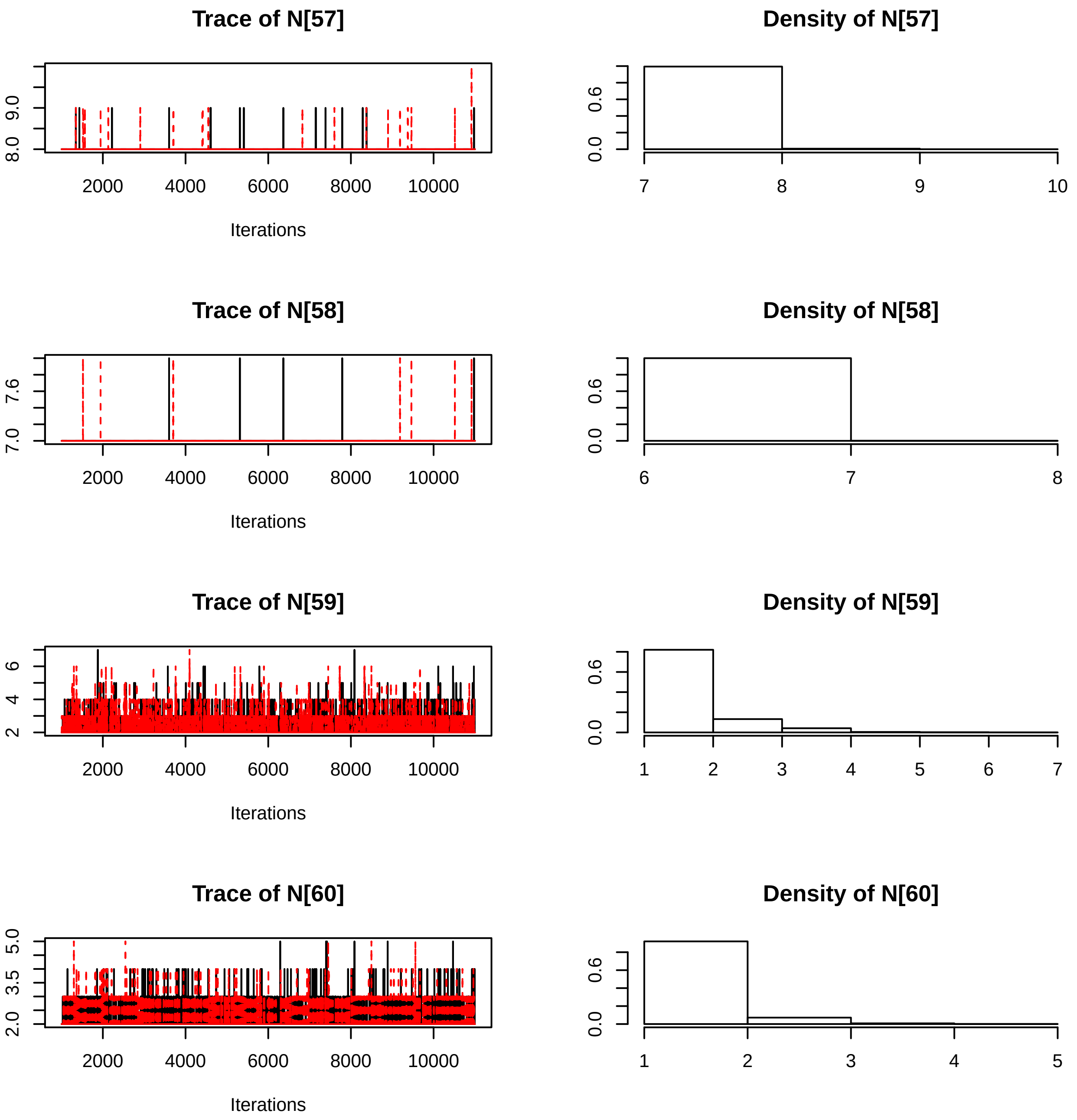

Trace of N[57]
Density of N[57]
Trace of N[58]
Density of N[58]
Trace of N[59]
Density of N[59]
Trace of N[60]
Density of N[60]
Iterations
2000
4000
6000
8000
10000

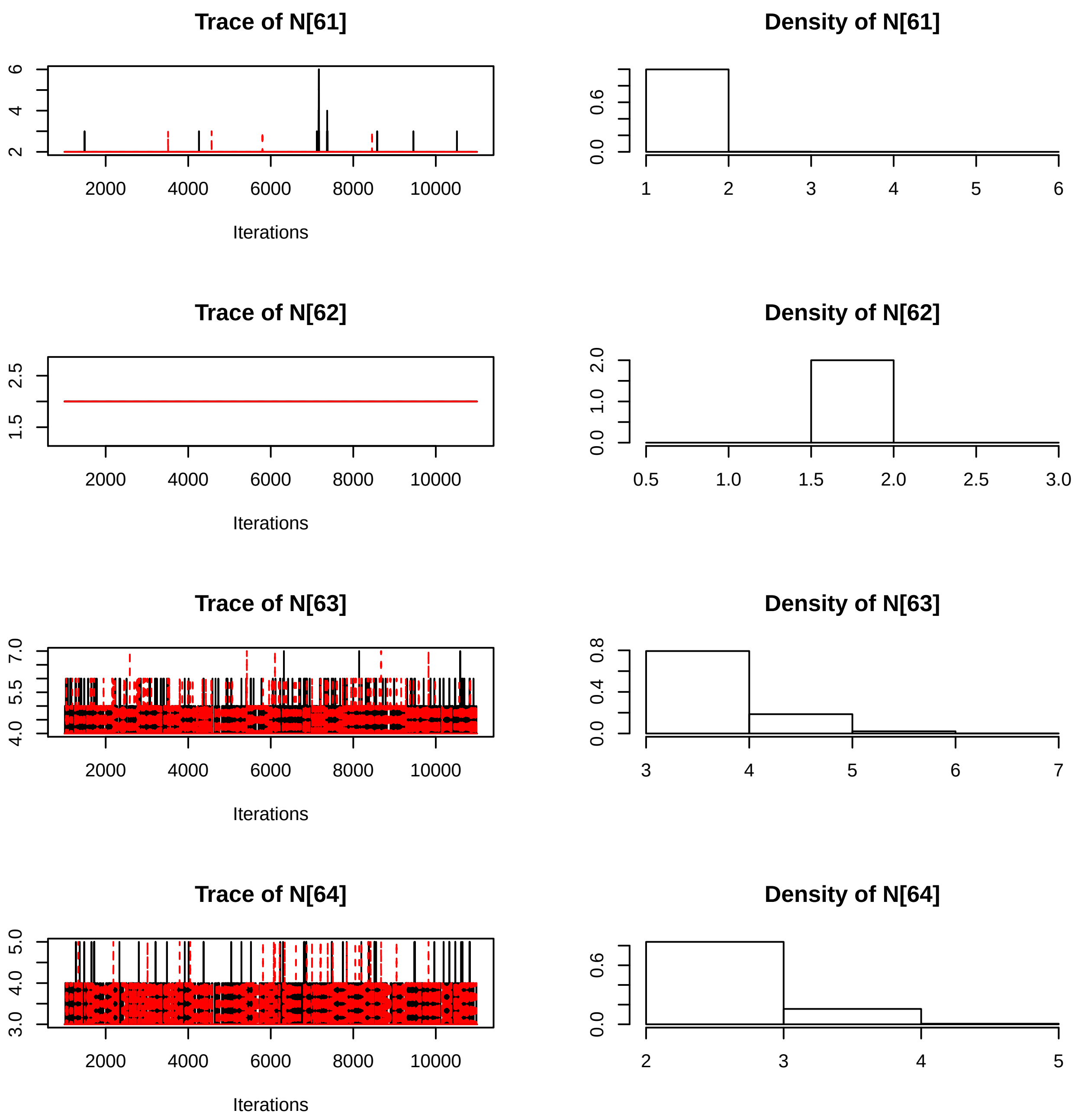
Trace of N[61]
Density of N[61]
Trace of N[62]
Density of N[62]
Trace of N[63]
Density of N[63]
Trace of N[64]
Density of N[64]
Iterations
2000
4000
6000
8000
10000

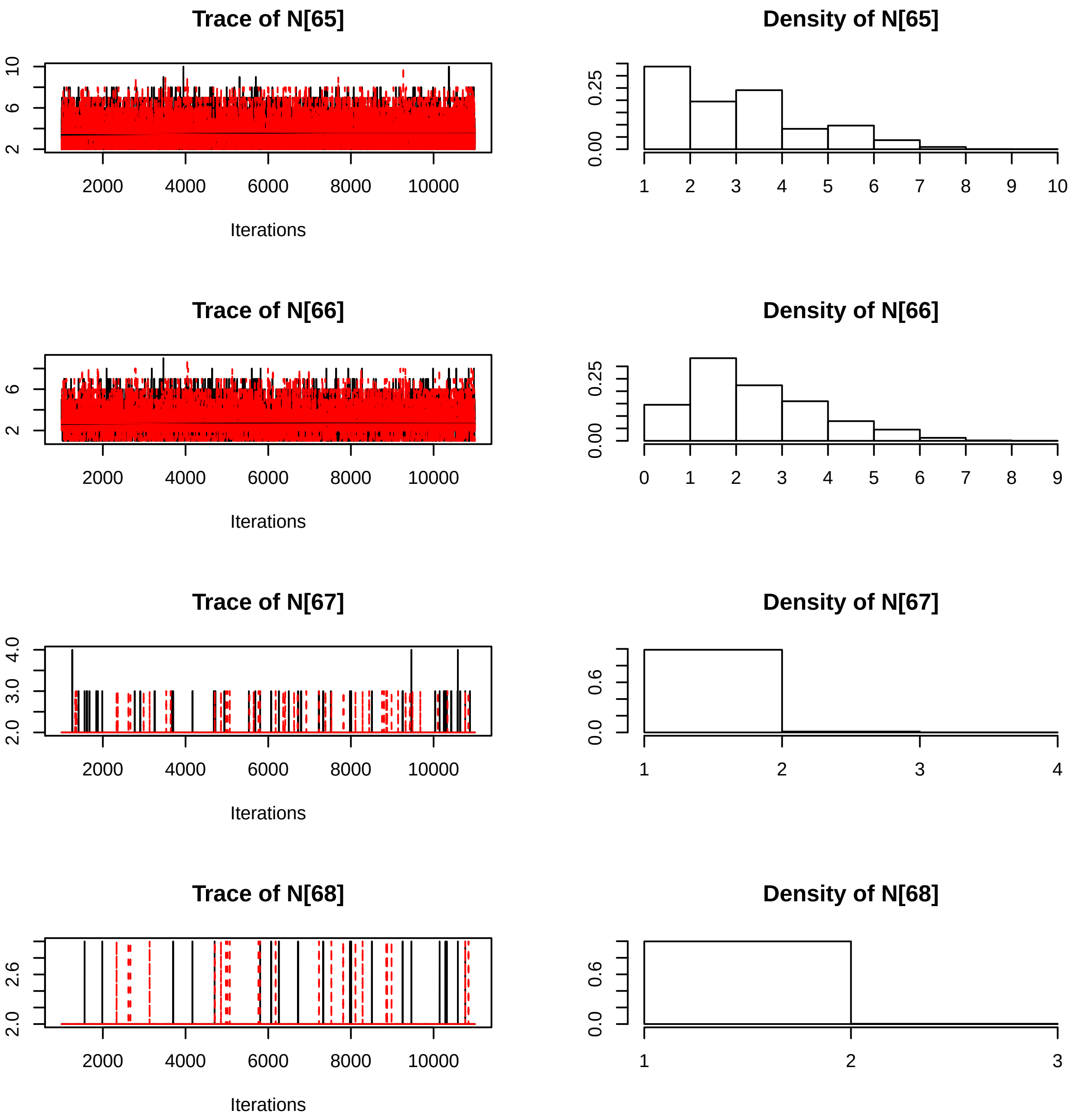
Trace of N[65]
Iterations
Density of N[65]
Trace of N[66]
Iterations
Density of N[66]
Trace of N[67]
Iterations
Density of N[67]
Trace of N[68]
Iterations
Density of N[68]

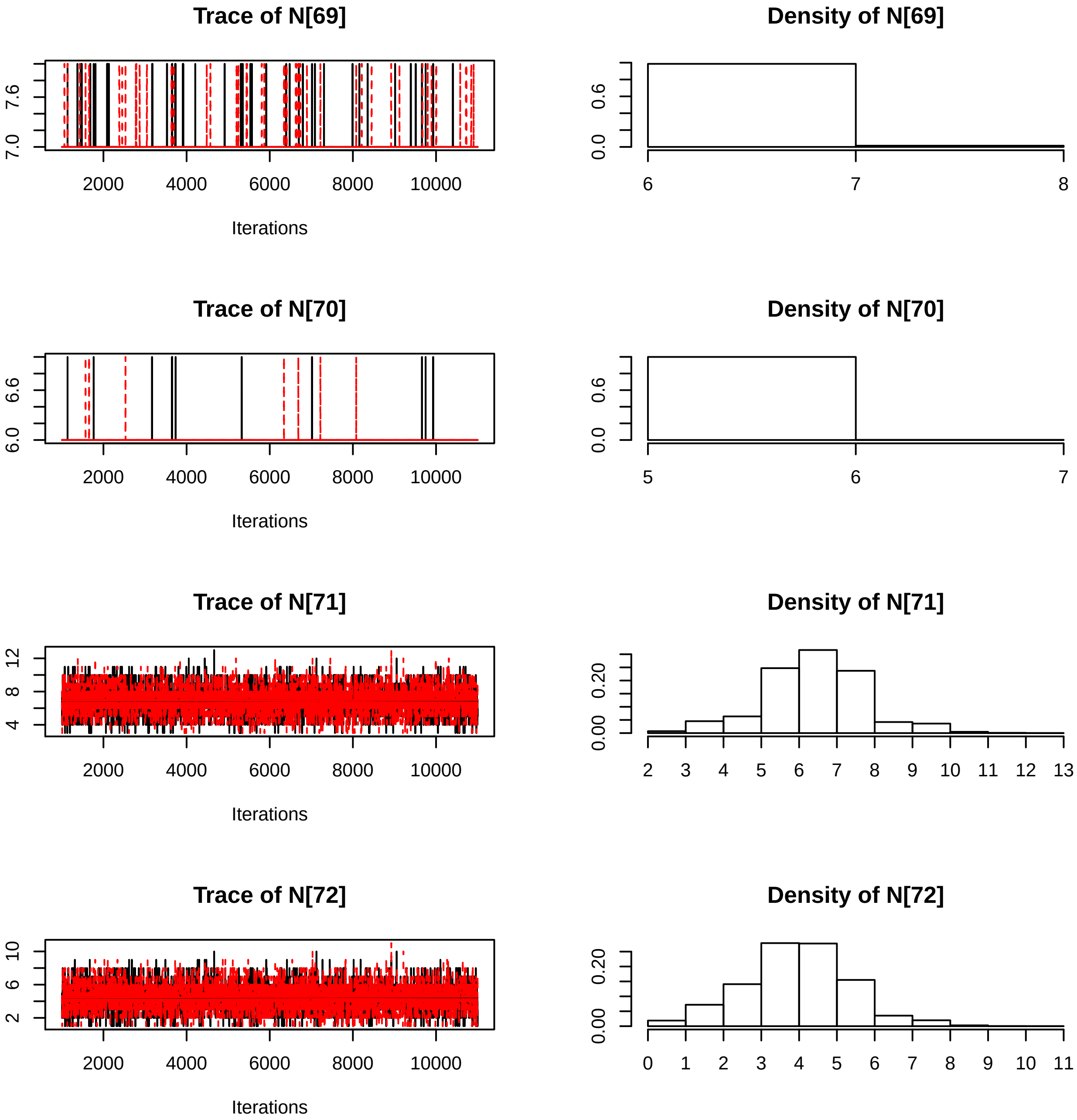
Trace of N[69]
7.0
7.6
2000
4000
6000
8000
10000
Iterations
Density of N[69]
0.0
0.6
6
7
8
Trace of N[70]
6.0
6.6
2000
4000
6000
8000
10000
Iterations
Density of N[70]
0.0
0.6
5
6
7
Trace of N[71]
4
8
12
2000
4000
6000
8000
10000
Iterations
Density of N[71]
0.00
0.20
2 3 4 5 6 7 8 9 10 11 12 13
Trace of N[72]
2
6
10
2000
4000
6000
8000
10000
Iterations
Density of N[72]
0.00
0.20
0 1 2 3 4 5 6 7 8 9 10 11

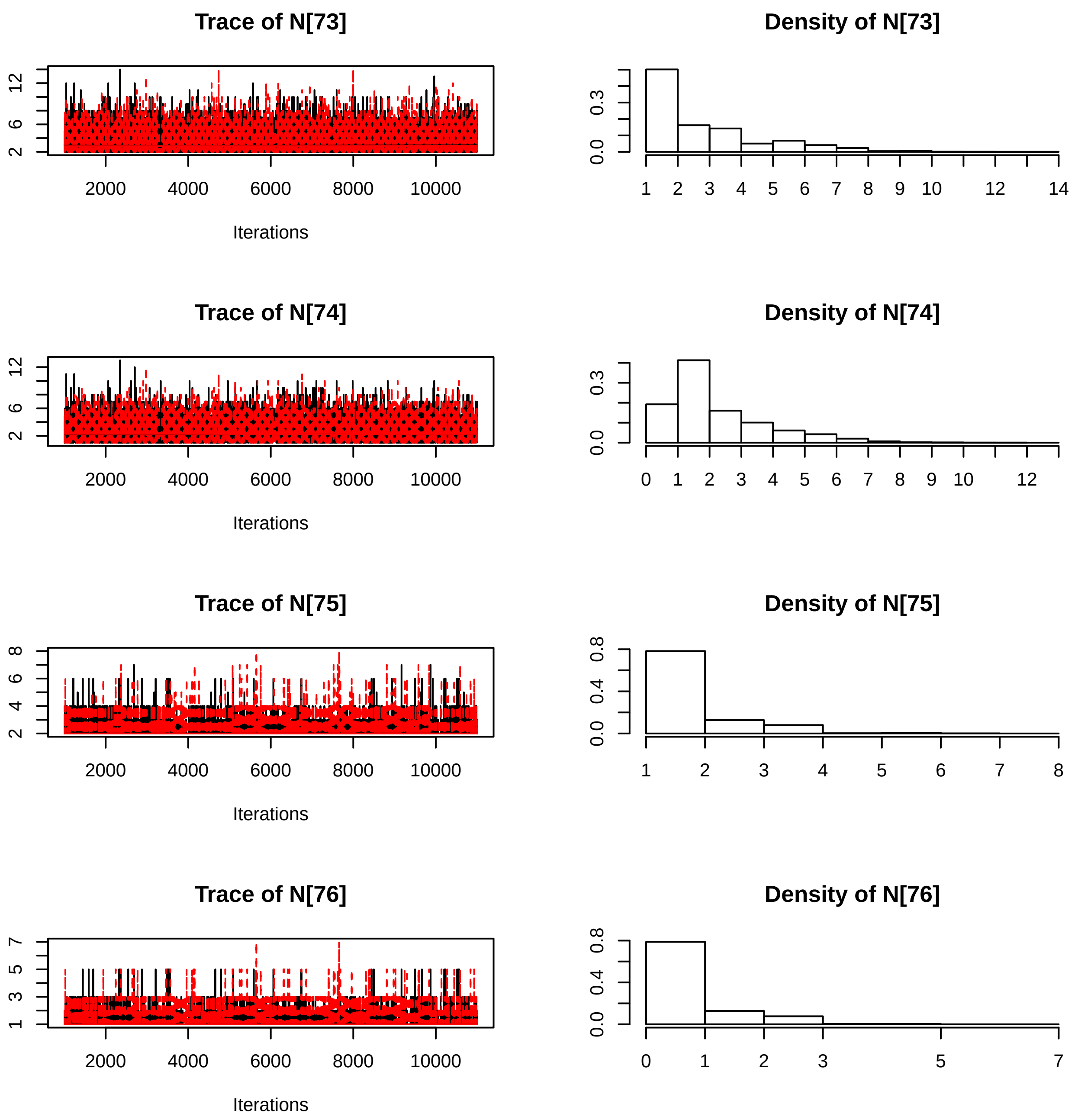
Trace of N[73]
Iterations
Density of N[73]
Trace of N[74]
Iterations
Density of N[74]
Trace of N[75]
Iterations
Density of N[75]
Trace of N[76]
Iterations
Density of N[76]

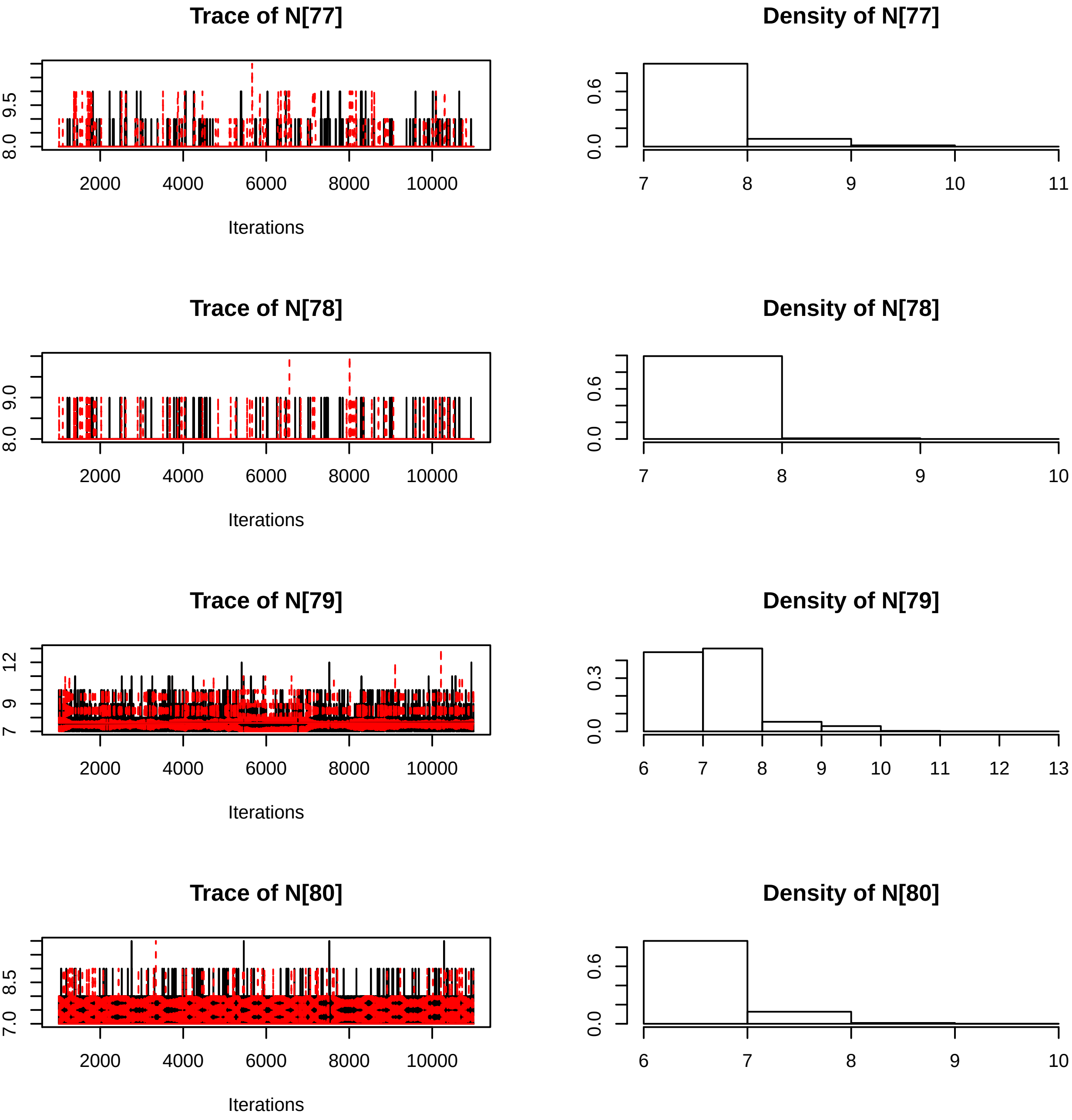
Trace of N[77]
9.5
8.0
2000
4000
6000
8000
10000
Iterations
Density of N[77]
0.6
0.0
7
8
9
10
11
Trace of N[78]
9.0
8.0
2000
4000
6000
8000
10000
Iterations
Density of N[78]
0.6
0.0
7
8
9
10
Trace of N[79]
12
9
7
2000
4000
6000
8000
10000
Iterations
Density of N[79]
0.3
0.0
6
7
8
9
10
11
12
13
Trace of N[80]
8.5
7.0
2000
4000
6000
8000
10000
Iterations
Density of N[80]
0.6
0.0
6
7
8
9
10

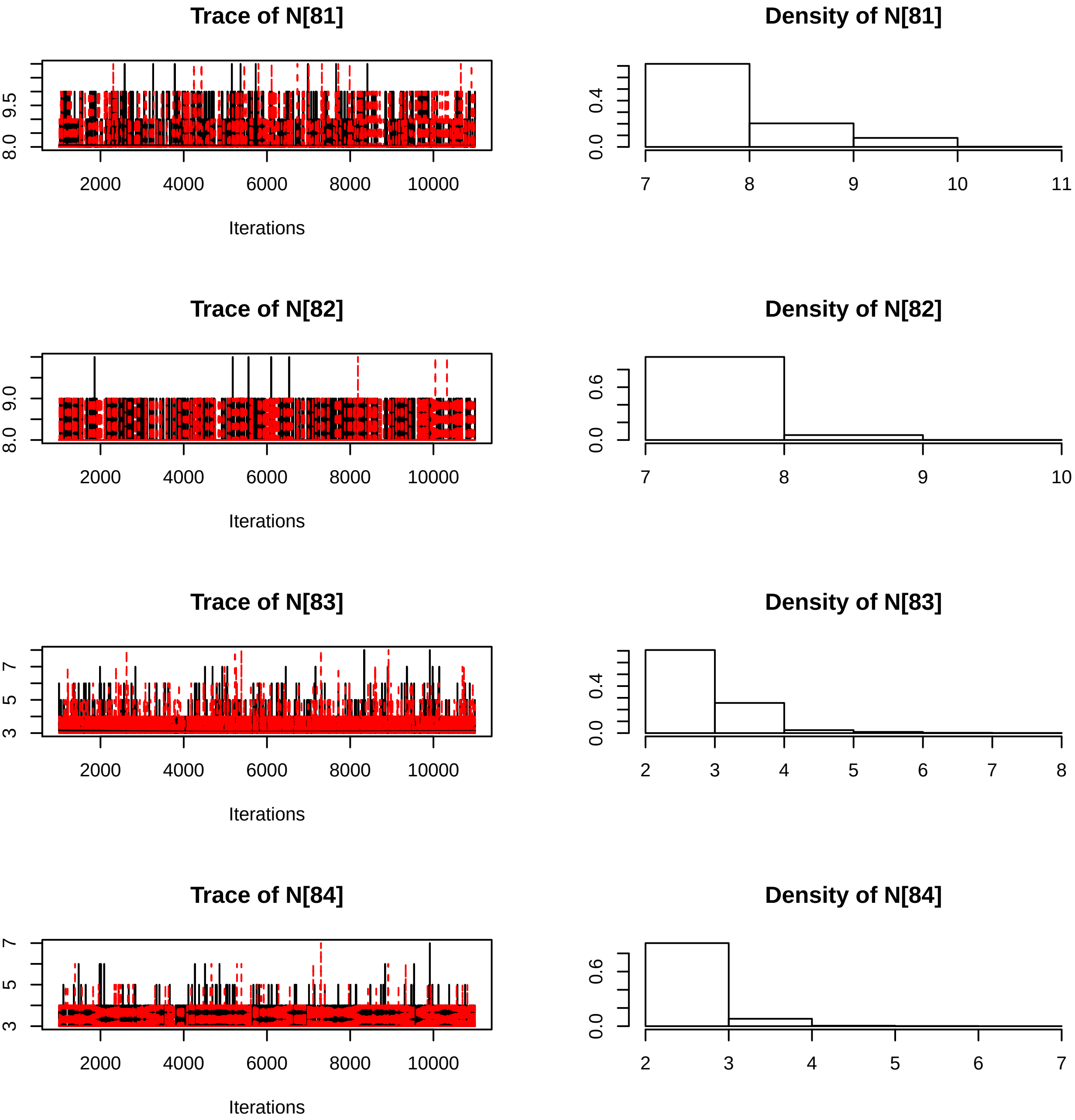
Trace of N[81]
Density of N[81]
Trace of N[82]
Density of N[82]
Trace of N[83]
Density of N[83]
Trace of N[84]
Density of N[84]
Iterations
Iterations
Iterations
Iterations

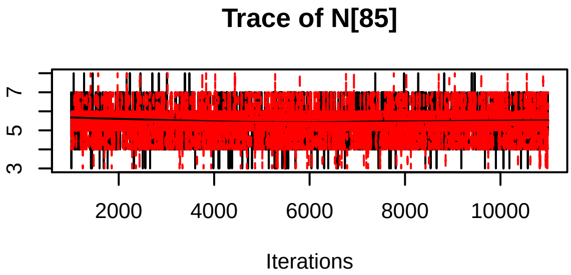
Trace of N[85]
Iterations

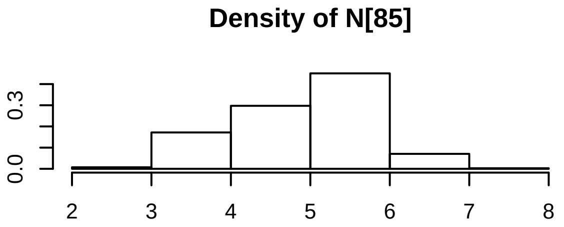
Density of N[85]

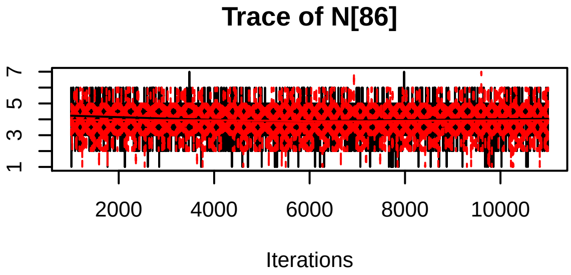
Trace of N[86]
Iterations

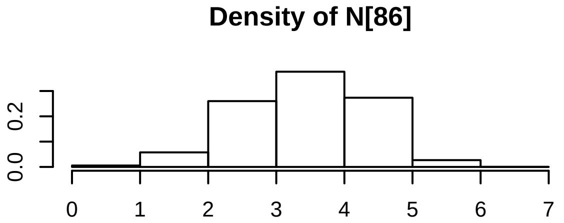
Density of N[86]

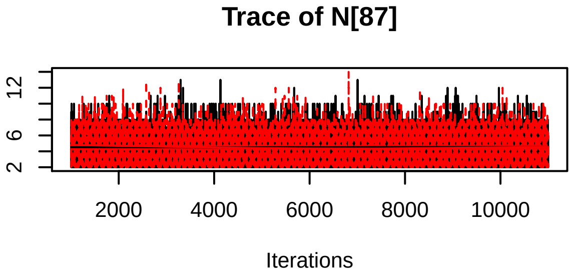
Trace of N[87]
Iterations

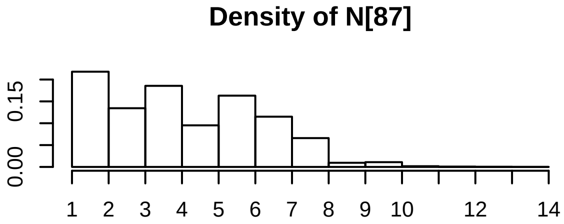
Density of N[87]

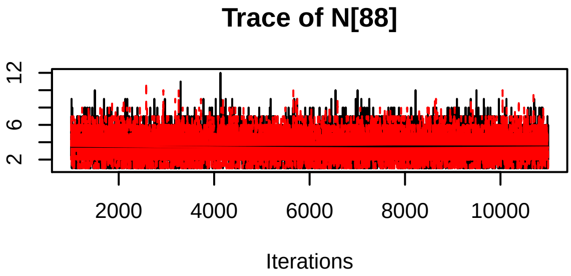
Trace of N[88]
Iterations

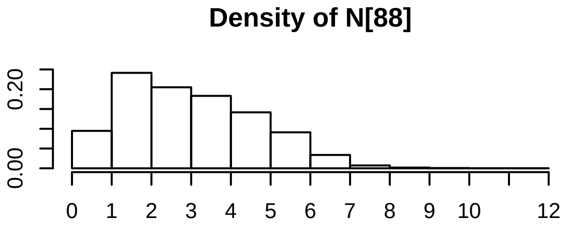
Density of N[88]

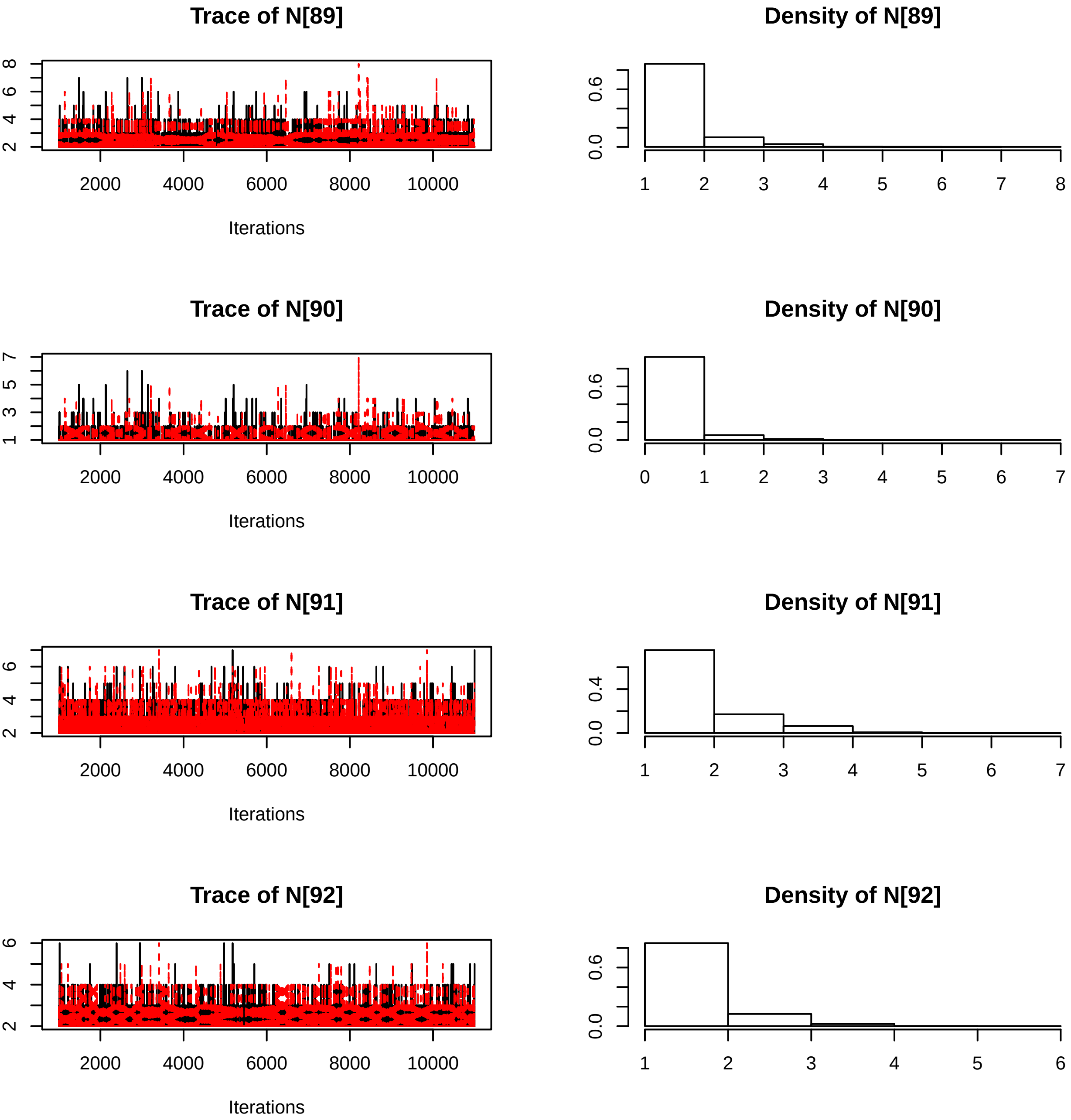
Trace of N[89]
Iterations
Density of N[89]
Trace of N[90]
Iterations
Density of N[90]
Trace of N[91]
Iterations
Density of N[91]
Trace of N[92]
Iterations
Density of N[92]

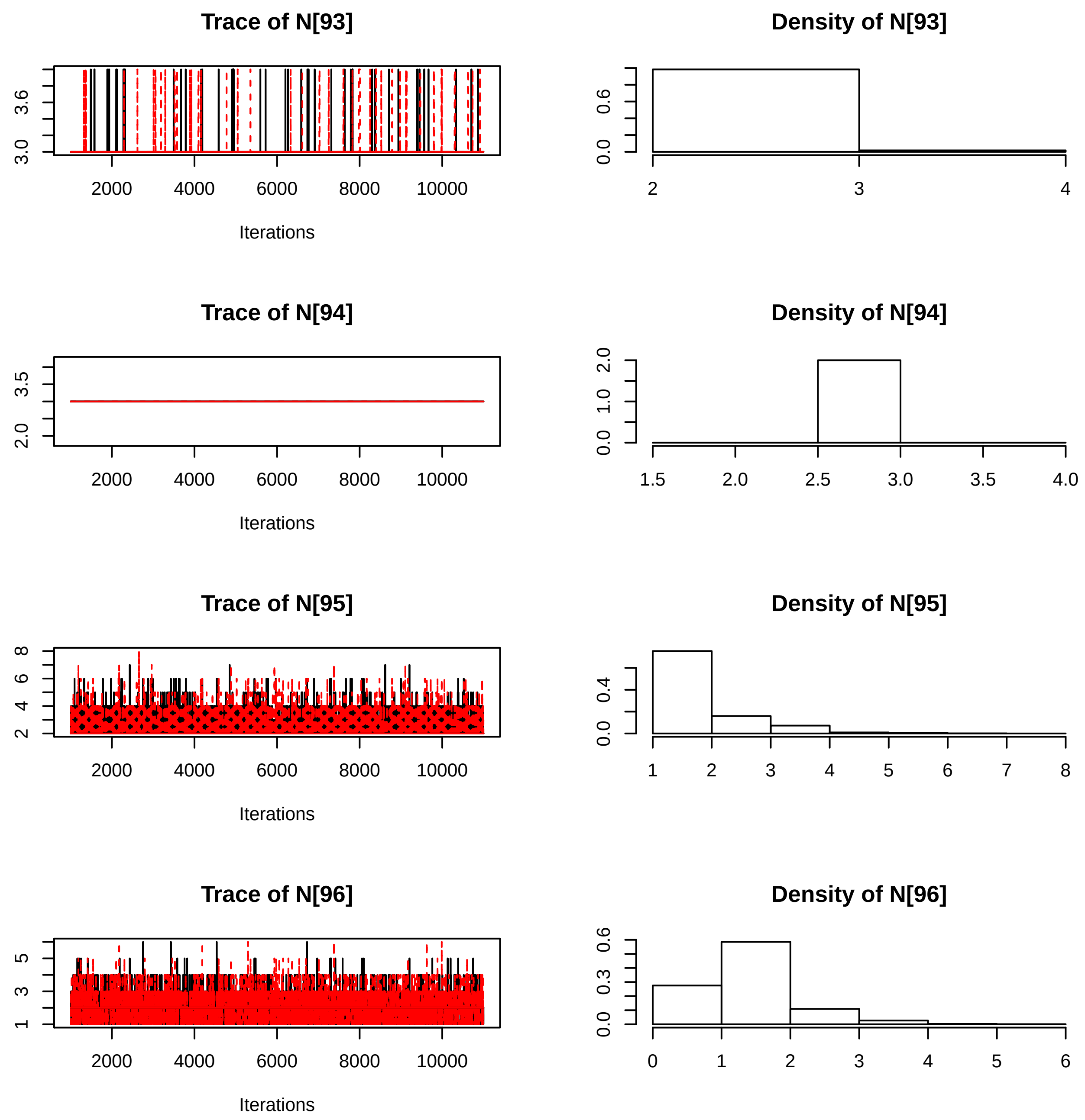

Trace of N[93]
Density of N[93]
Trace of N[94]
Density of N[94]
Trace of N[95]
Density of N[95]
Trace of N[96]
Density of N[96]
Iterations

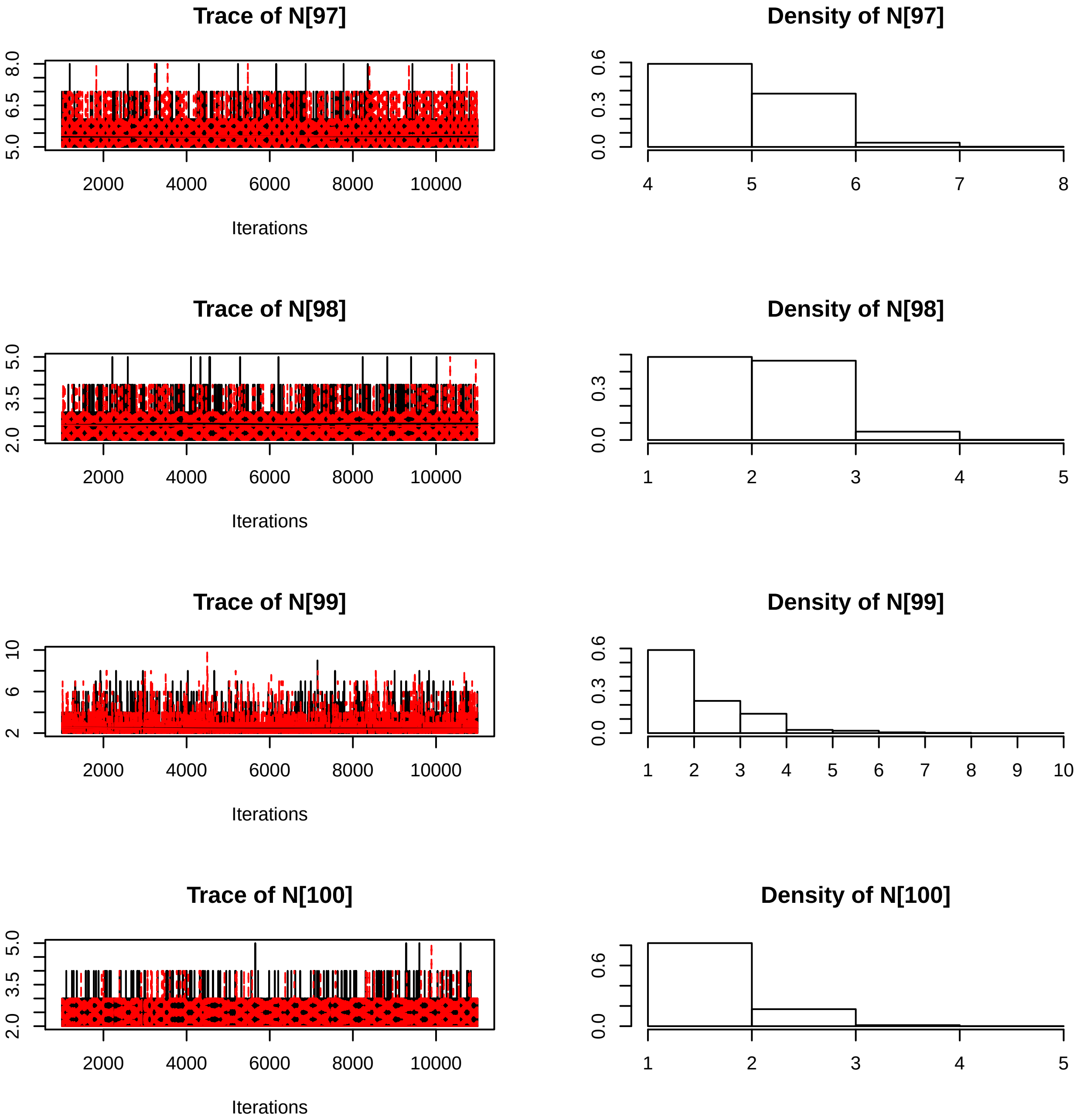

Trace of N[97]
5.0
6.5
8.0
2000
4000
6000
8000
10000
Iterations
Density of N[97]
0.0
0.3
0.6
4
5
6
7
8
Trace of N[98]
2.0
3.5
5.0
2000
4000
6000
8000
10000
Iterations
Density of N[98]
0.0
0.3
1
2
3
4
5
Trace of N[99]
2
6
10
2000
4000
6000
8000
10000
Iterations
Density of N[99]
0.0
0.3
0.6
1
2
3
4
5
6
7
8
9
10
Trace of N[100]
2.0
3.5
5.0
2000
4000
6000
8000
10000
Iterations
Density of N[100]
0.0
0.6
1
2
3
4
5

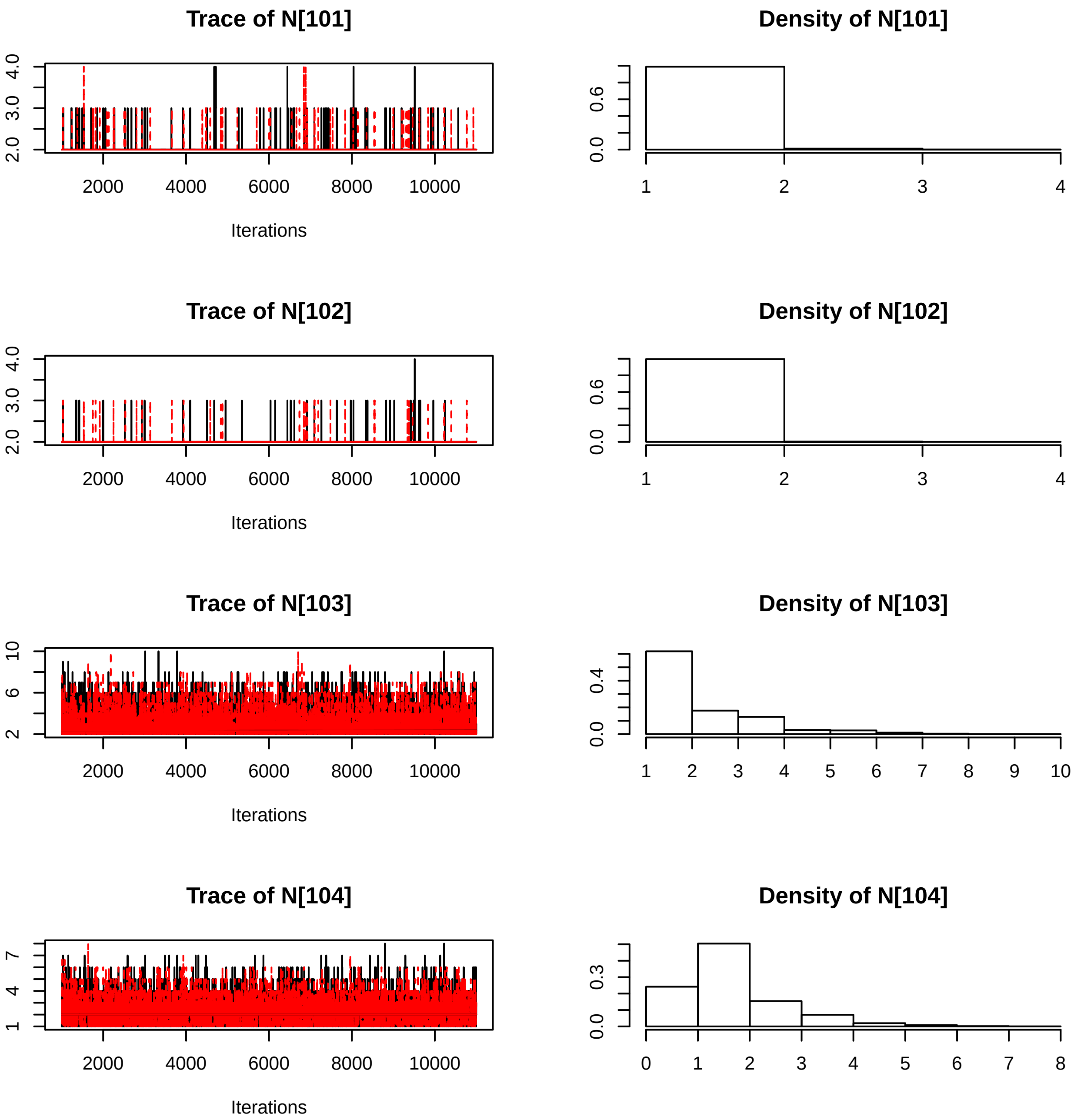
Trace of N[101]
Density of N[101]
Iterations
Trace of N[102]
Density of N[102]
Iterations
Trace of N[103]
Density of N[103]
Iterations
Trace of N[104]
Density of N[104]
Iterations

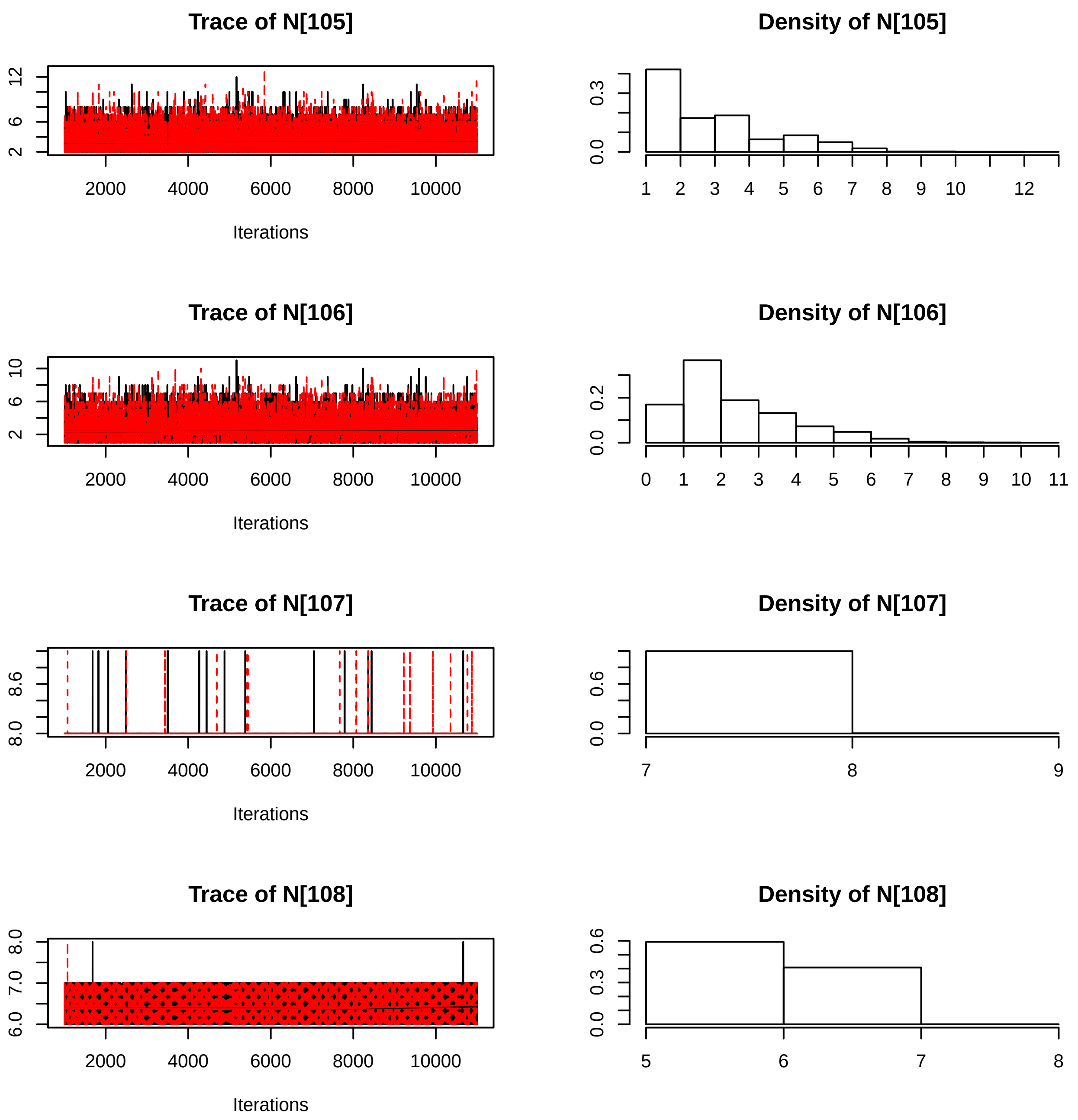
Trace of N[105]
Iterations
Density of N[105]
Trace of N[106]
Iterations
Density of N[106]
Trace of N[107]
Iterations
Density of N[107]
Trace of N[108]
Iterations
Density of N[108]

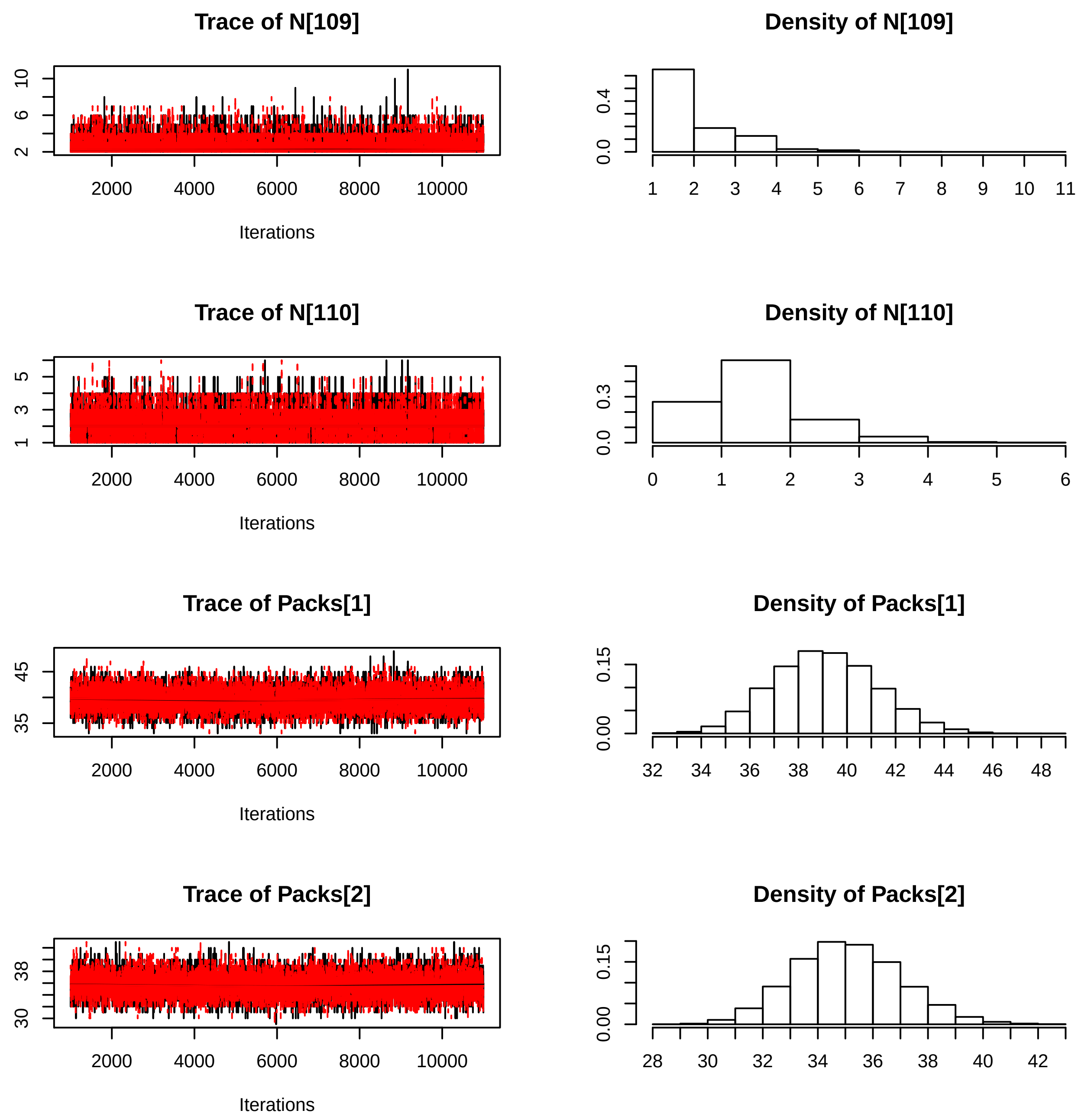
Trace of N[109]
Density of N[109]
Trace of N[110]
Density of N[110]
Trace of Packs[1]
Density of Packs[1]
Trace of Packs[2]
Density of Packs[2]
Iterations
Iterations
Iterations
Iterations

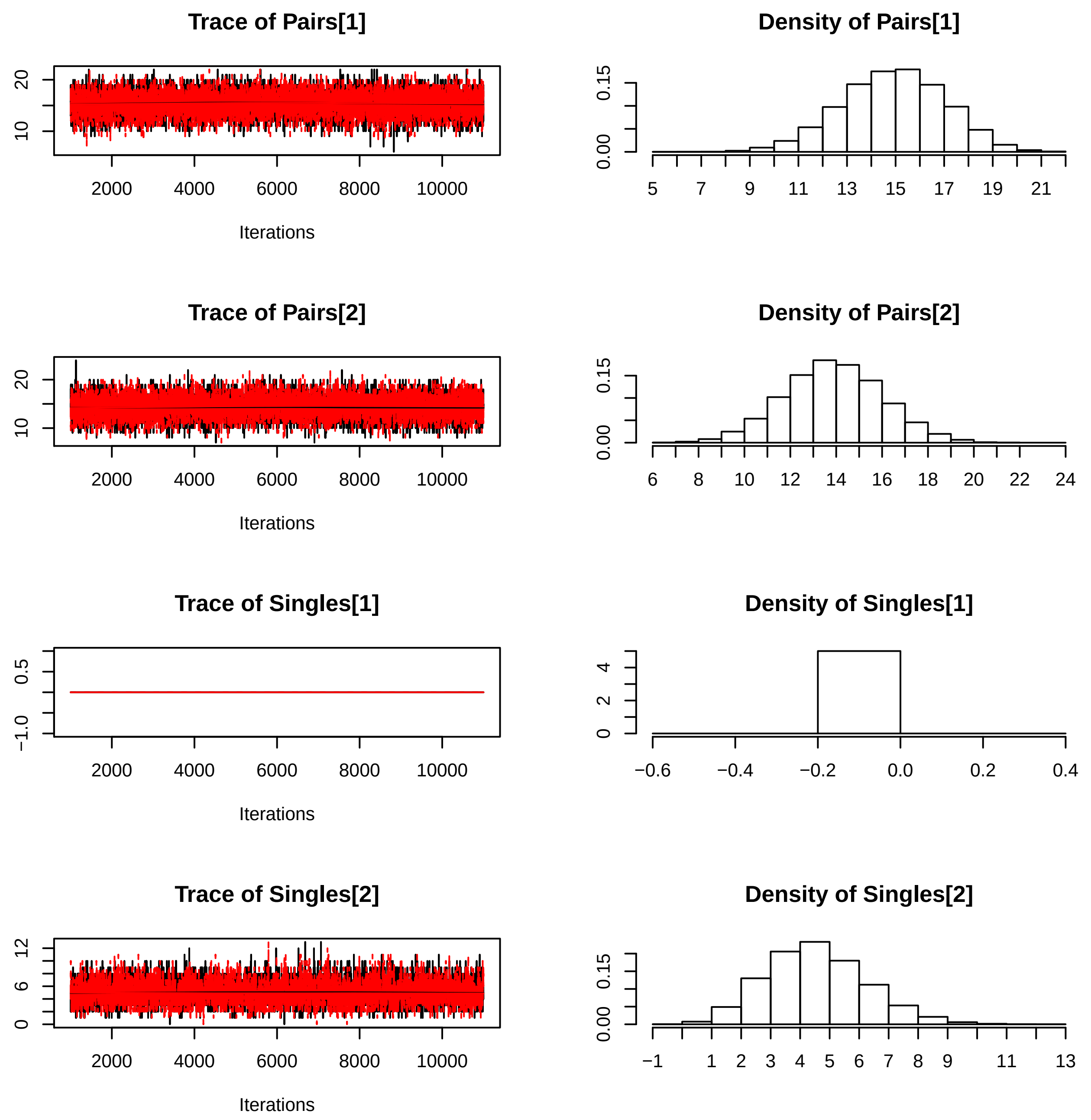
Trace of Pairs[1]
Density of Pairs[1]
Trace of Pairs[2]
Density of Pairs[2]
Trace of Singles[1]
Density of Singles[1]
Trace of Singles[2]
Density of Singles[2]
Iterations
2000 4000 6000 8000 10000
0.00 0.15
5 7 9 11 13 15 17 19 21
6 8 10 12 14 16 18 20 22 24
−1.0 0.5
0 2 4
−0.6 −0.4 −0.2 0.0 0.2 0.4
0 6 12
−1 1 2 3 4 5 6 7 8 9 11 13

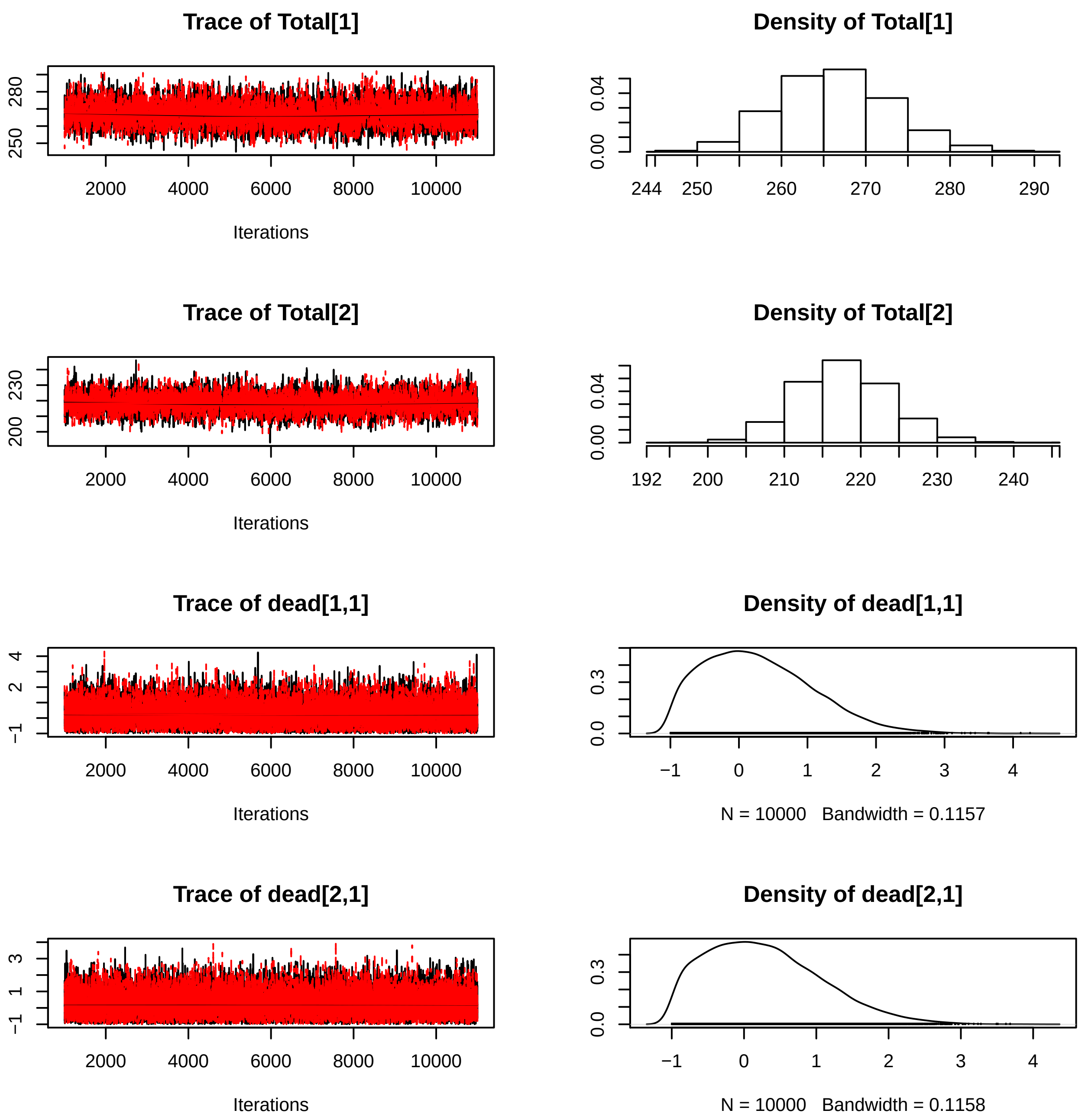
Trace of Total[1]
Iterations
Density of Total[1]
Trace of Total[2]
Iterations
Density of Total[2]
Trace of dead[1,1]
Iterations
Density of dead[1,1]
N = 10000 Bandwidth = 0.1157
Trace of dead[2,1]
Iterations
Density of dead[2,1]
N = 10000 Bandwidth = 0.1158

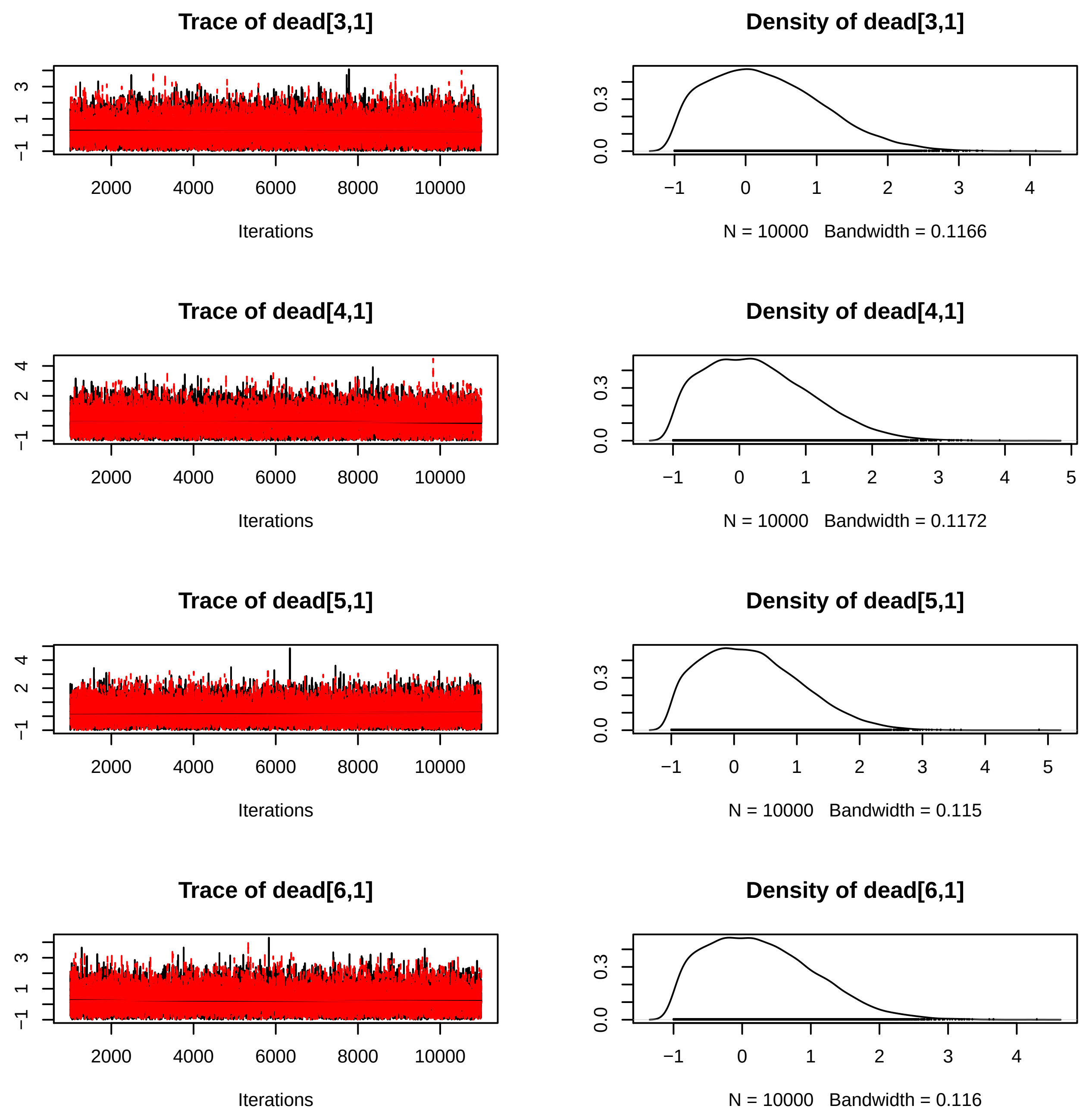
Trace of dead[3,1]
Iterations
Density of dead[3,1]
N = 10000 Bandwidth = 0.1166
Trace of dead[4,1]
Iterations
Density of dead[4,1]
N = 10000 Bandwidth = 0.1172
Trace of dead[5,1]
Iterations
Density of dead[5,1]
N = 10000 Bandwidth = 0.115
Trace of dead[6,1]
Iterations
Density of dead[6,1]
N = 10000 Bandwidth = 0.116

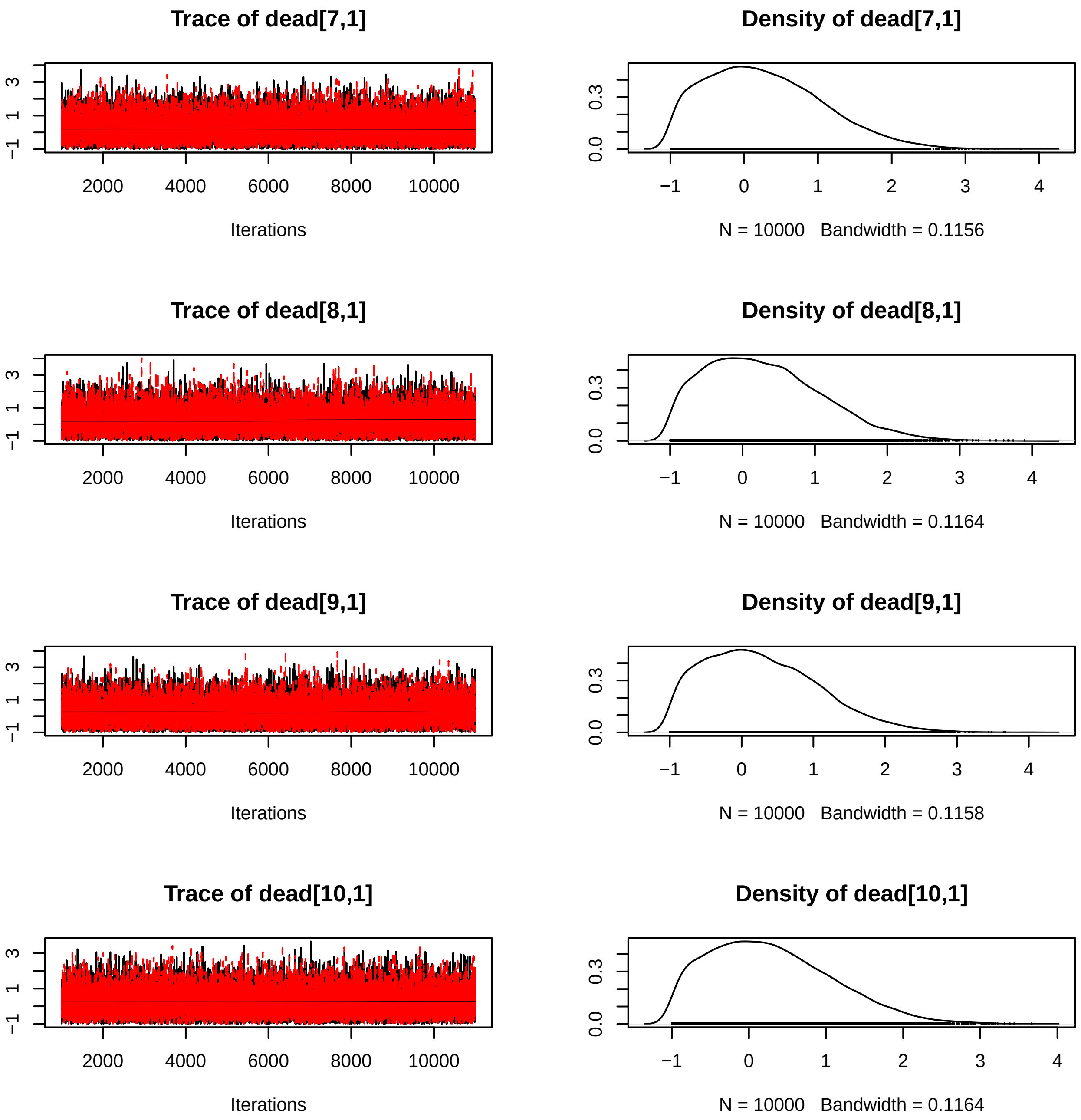
Trace of dead[7,1]
Iterations
Density of dead[7,1]
N = 10000 Bandwidth = 0.1156
Trace of dead[8,1]
Iterations
Density of dead[8,1]
N = 10000 Bandwidth = 0.1164
Trace of dead[9,1]
Iterations
Density of dead[9,1]
N = 10000 Bandwidth = 0.1158
Trace of dead[10,1]
Iterations
Density of dead[10,1]
N = 10000 Bandwidth = 0.1164
2000 4000 6000 8000 10000
−1 0 1 2 3 4
0.0 0.3

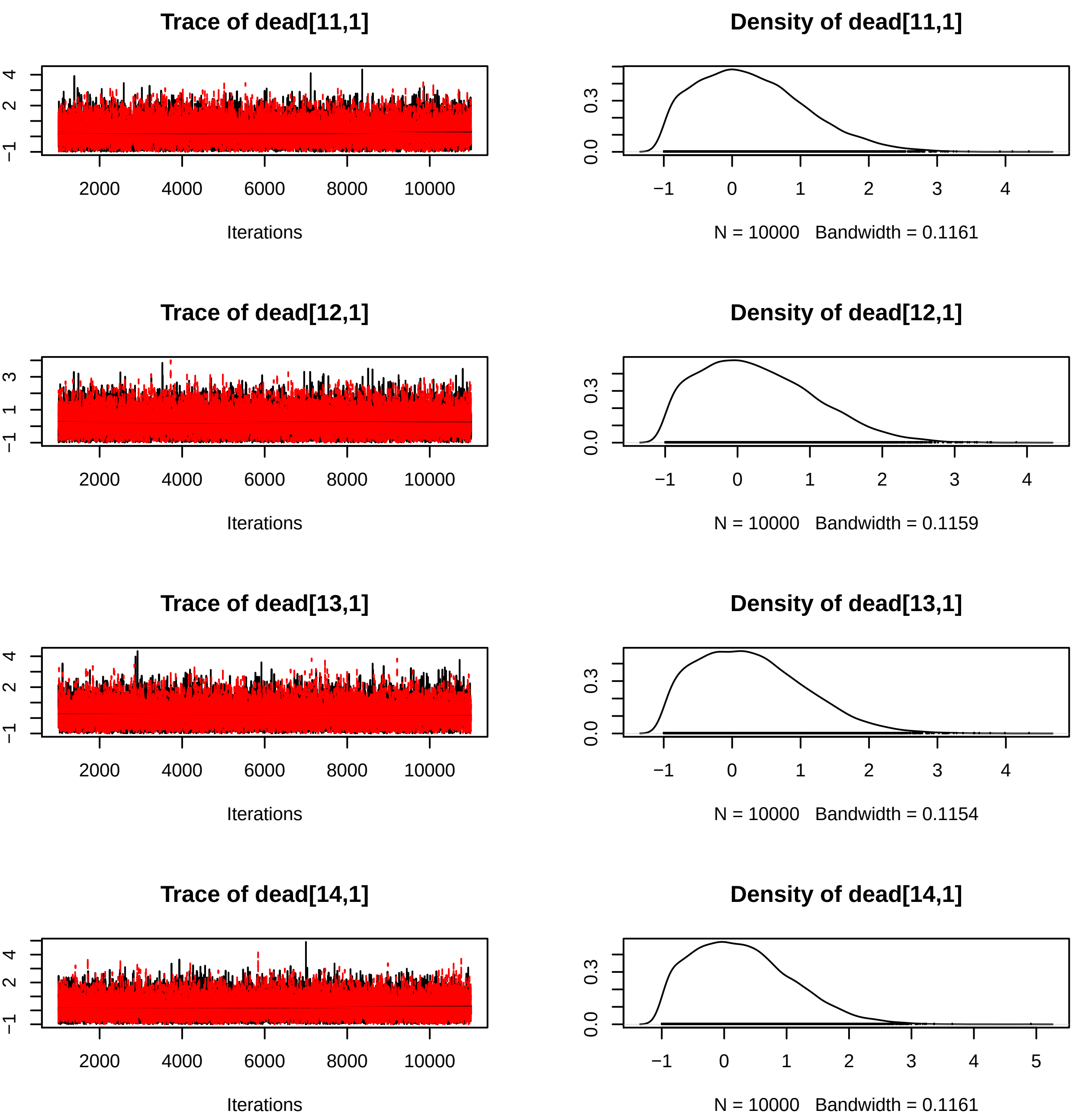
Trace of dead[11,1]
Iterations
Density of dead[11,1]
N = 10000 Bandwidth = 0.1161
Trace of dead[12,1]
Iterations
Density of dead[12,1]
N = 10000 Bandwidth = 0.1159
Trace of dead[13,1]
Iterations
Density of dead[13,1]
N = 10000 Bandwidth = 0.1154
Trace of dead[14,1]
Iterations
Density of dead[14,1]
N = 10000 Bandwidth = 0.1161

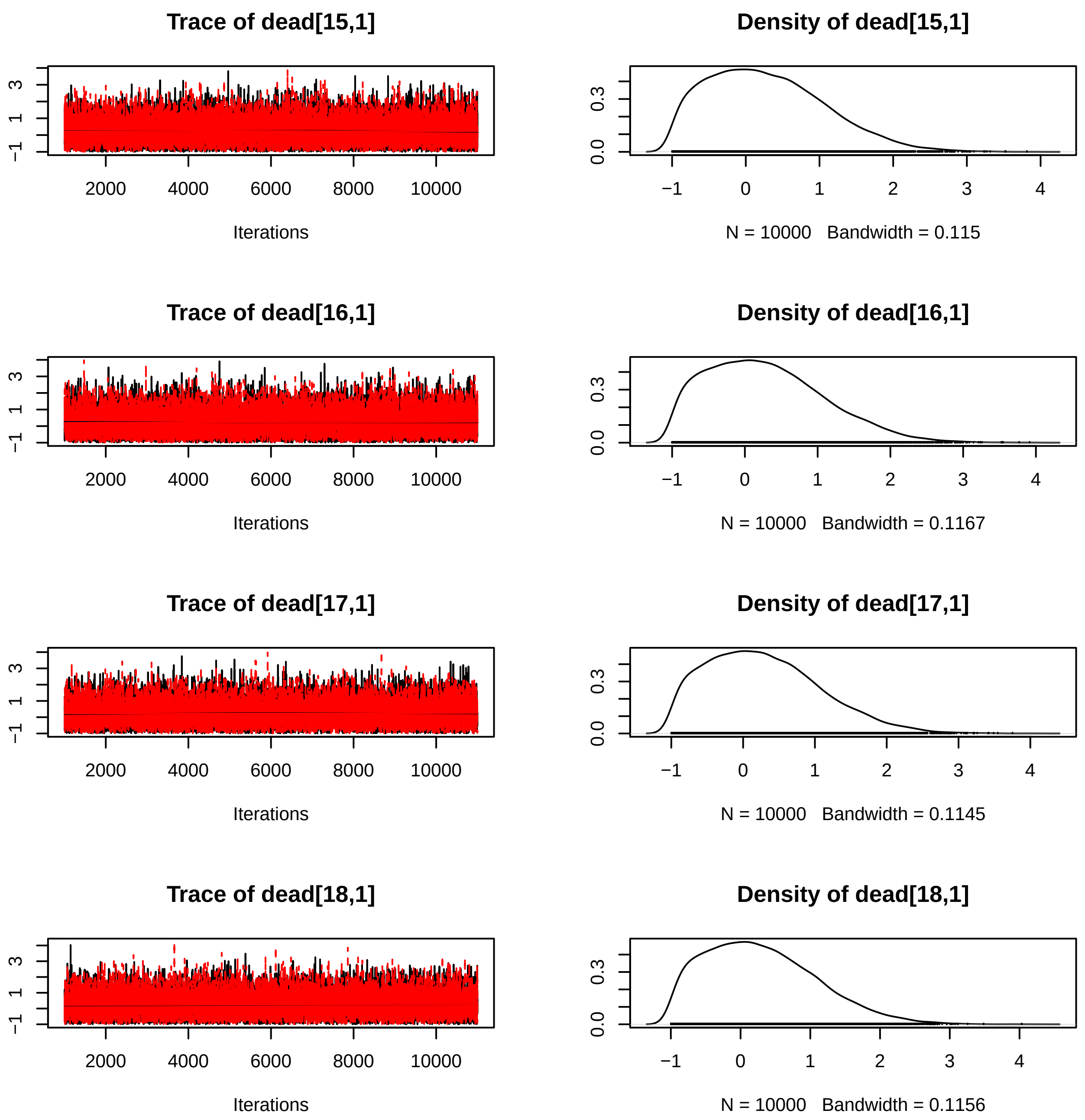
Trace of dead[15,1]
Iterations
Density of dead[15,1]
N = 10000 Bandwidth = 0.115
Trace of dead[16,1]
Iterations
Density of dead[16,1]
N = 10000 Bandwidth = 0.1167
Trace of dead[17,1]
Iterations
Density of dead[17,1]
N = 10000 Bandwidth = 0.1145
Trace of dead[18,1]
Iterations
Density of dead[18,1]
N = 10000 Bandwidth = 0.1156

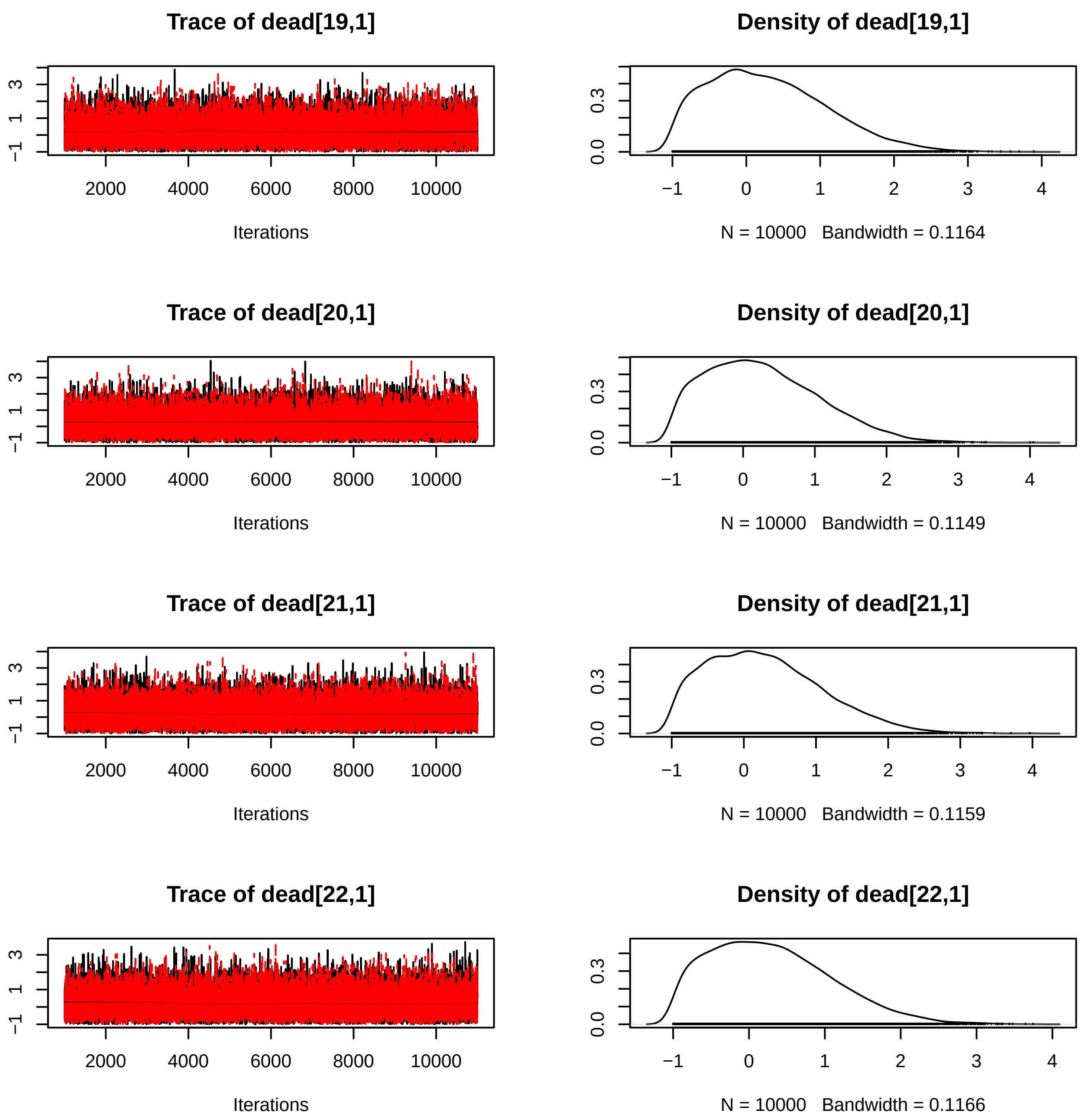

Trace of dead[19,1]
Iterations
Density of dead[19,1]
N = 10000 Bandwidth = 0.1164
Trace of dead[20,1]
Iterations
Density of dead[20,1]
N = 10000 Bandwidth = 0.1149
Trace of dead[21,1]
Iterations
Density of dead[21,1]
N = 10000 Bandwidth = 0.1159
Trace of dead[22,1]
Iterations
Density of dead[22,1]
N = 10000 Bandwidth = 0.1166

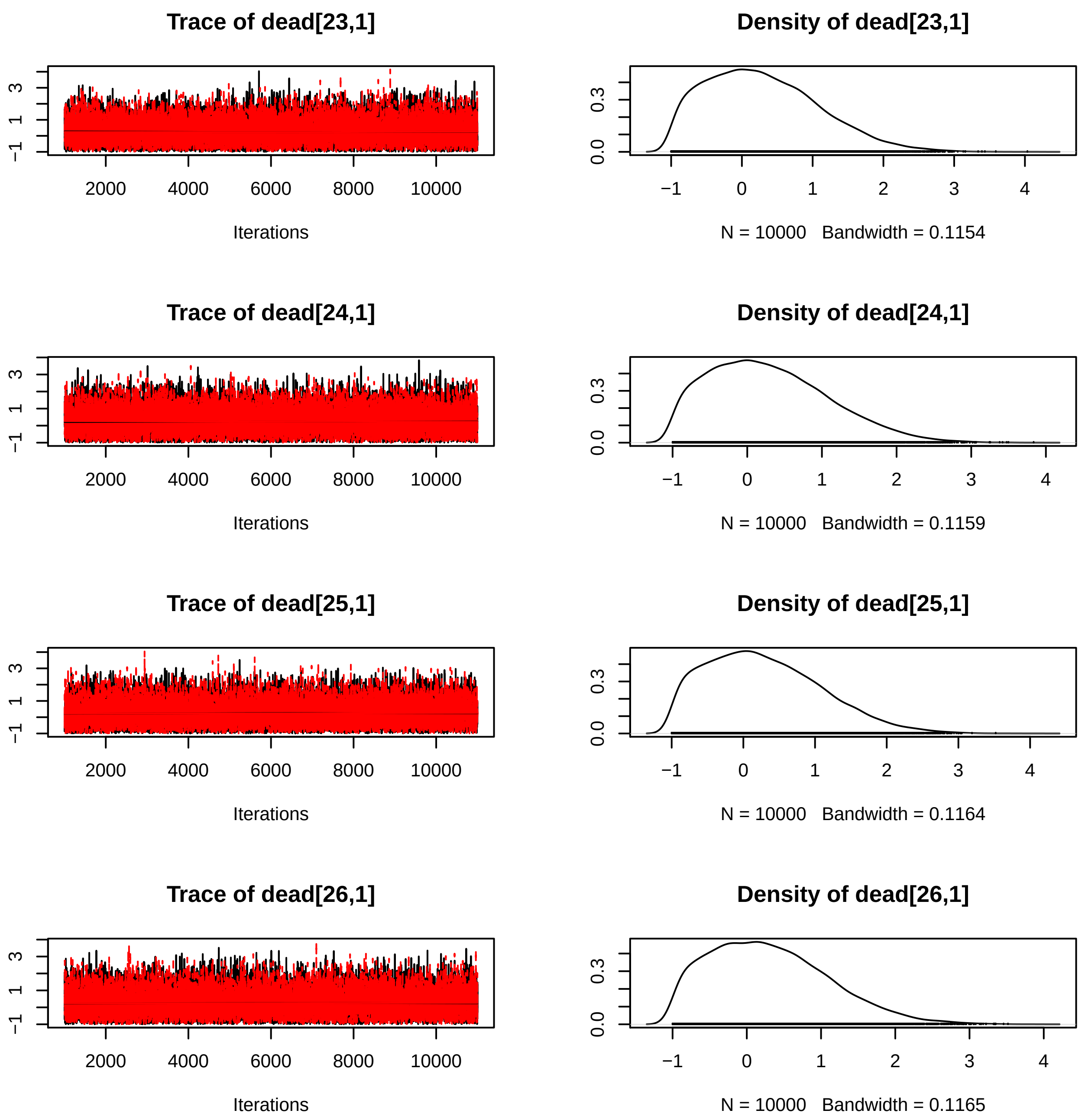
Trace of dead[23,1]
Density of dead[23,1]
−1
1
3
0.0
0.3
2000
4000
6000
8000
10000
Iterations
−1
0
1
2
3
4
N = 10000 Bandwidth = 0.1154
Trace of dead[24,1]
Density of dead[24,1]
Iterations
N = 10000 Bandwidth = 0.1159
Trace of dead[25,1]
Density of dead[25,1]
Iterations
N = 10000 Bandwidth = 0.1164
Trace of dead[26,1]
Density of dead[26,1]
Iterations
N = 10000 Bandwidth = 0.1165

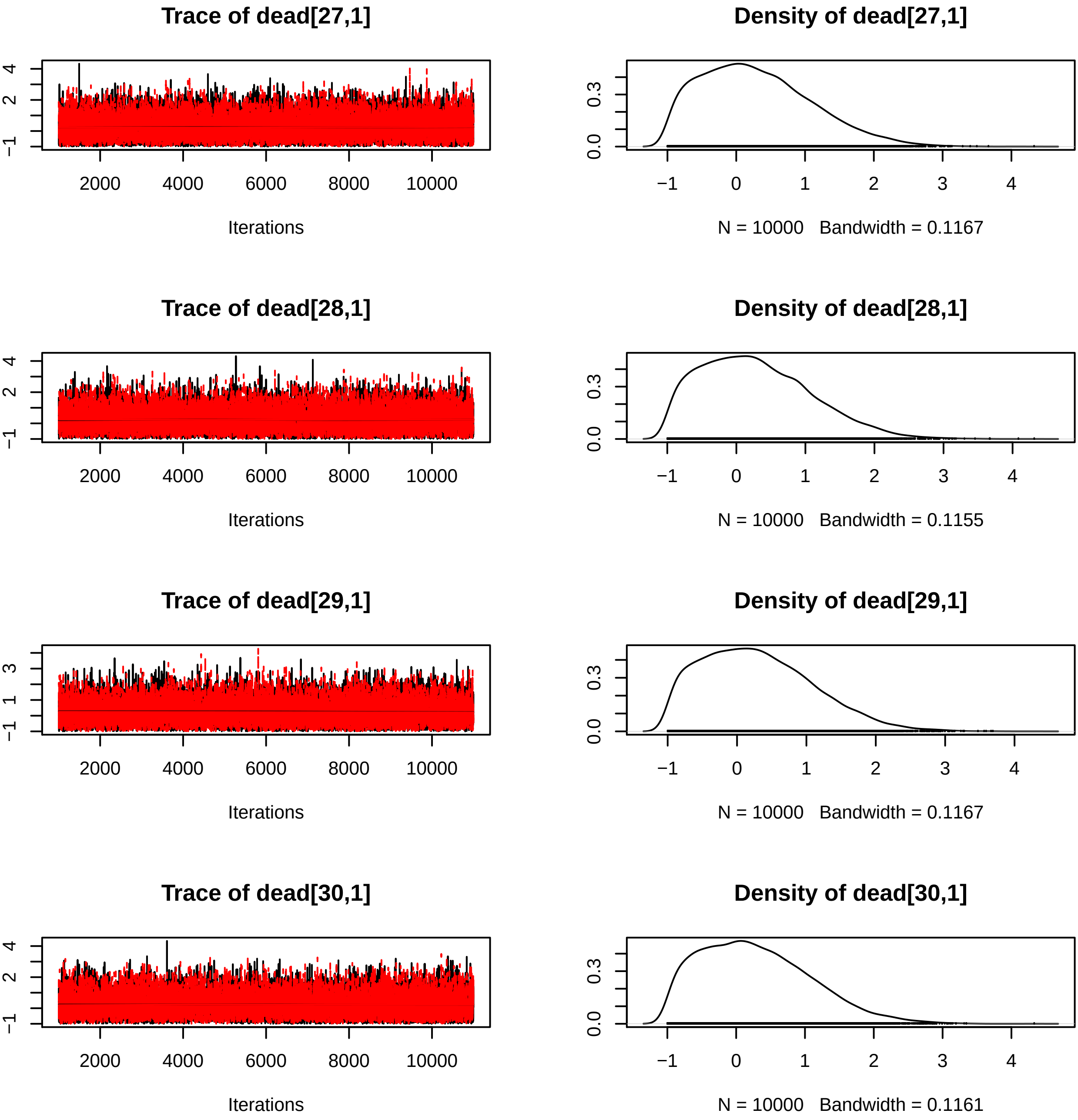
Trace of dead[27,1]
Iterations
Density of dead[27,1]
N = 10000 Bandwidth = 0.1167
Trace of dead[28,1]
Iterations
Density of dead[28,1]
N = 10000 Bandwidth = 0.1155
Trace of dead[29,1]
Iterations
Density of dead[29,1]
N = 10000 Bandwidth = 0.1167
Trace of dead[30,1]
Iterations
Density of dead[30,1]
N = 10000 Bandwidth = 0.1161

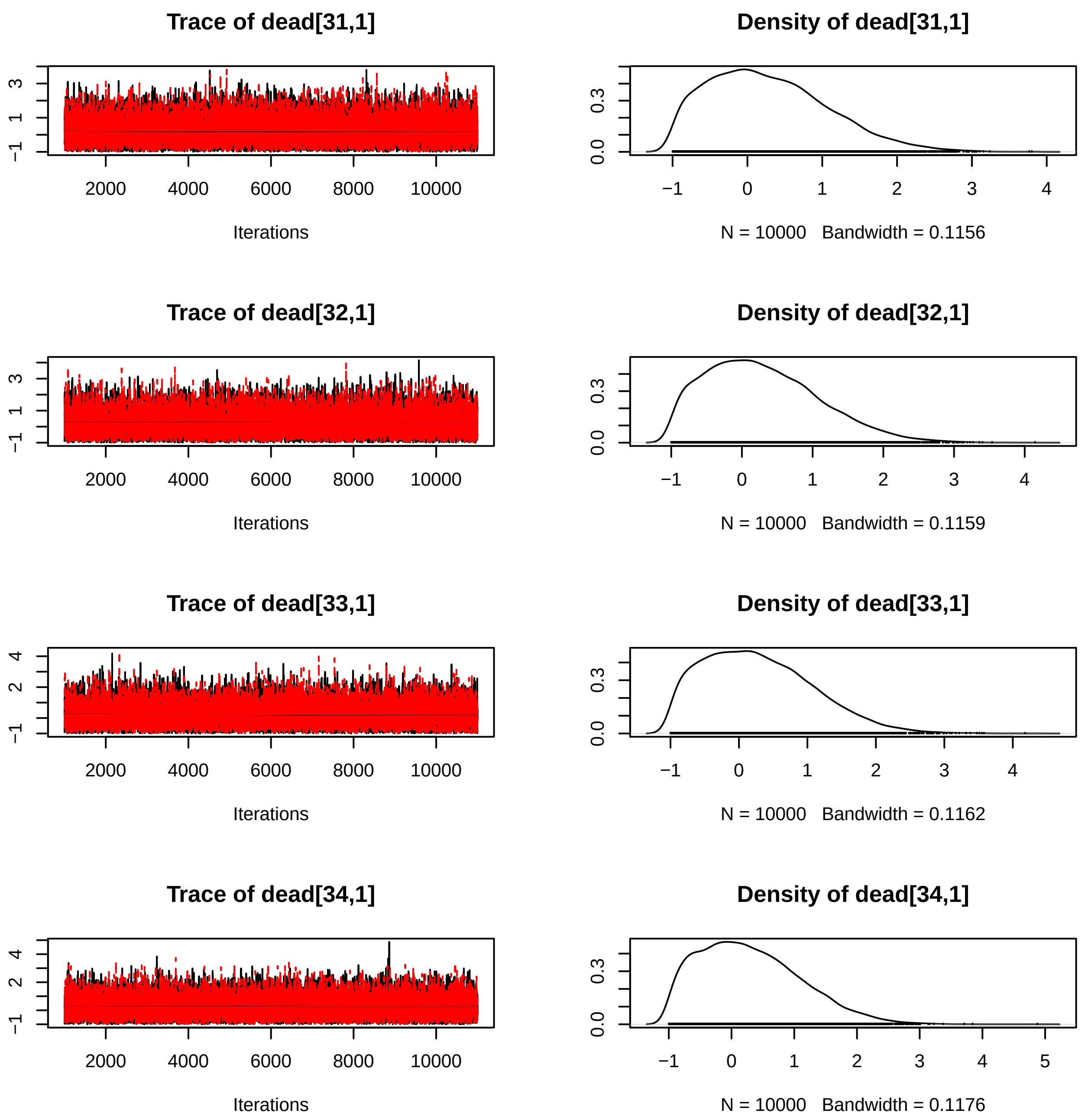

Trace of dead[31,1]
Iterations
Density of dead[31,1]
N = 10000 Bandwidth = 0.1156
Trace of dead[32,1]
Iterations
Density of dead[32,1]
N = 10000 Bandwidth = 0.1159
Trace of dead[33,1]
Iterations
Density of dead[33,1]
N = 10000 Bandwidth = 0.1162
Trace of dead[34,1]
Iterations
Density of dead[34,1]
N = 10000 Bandwidth = 0.1176

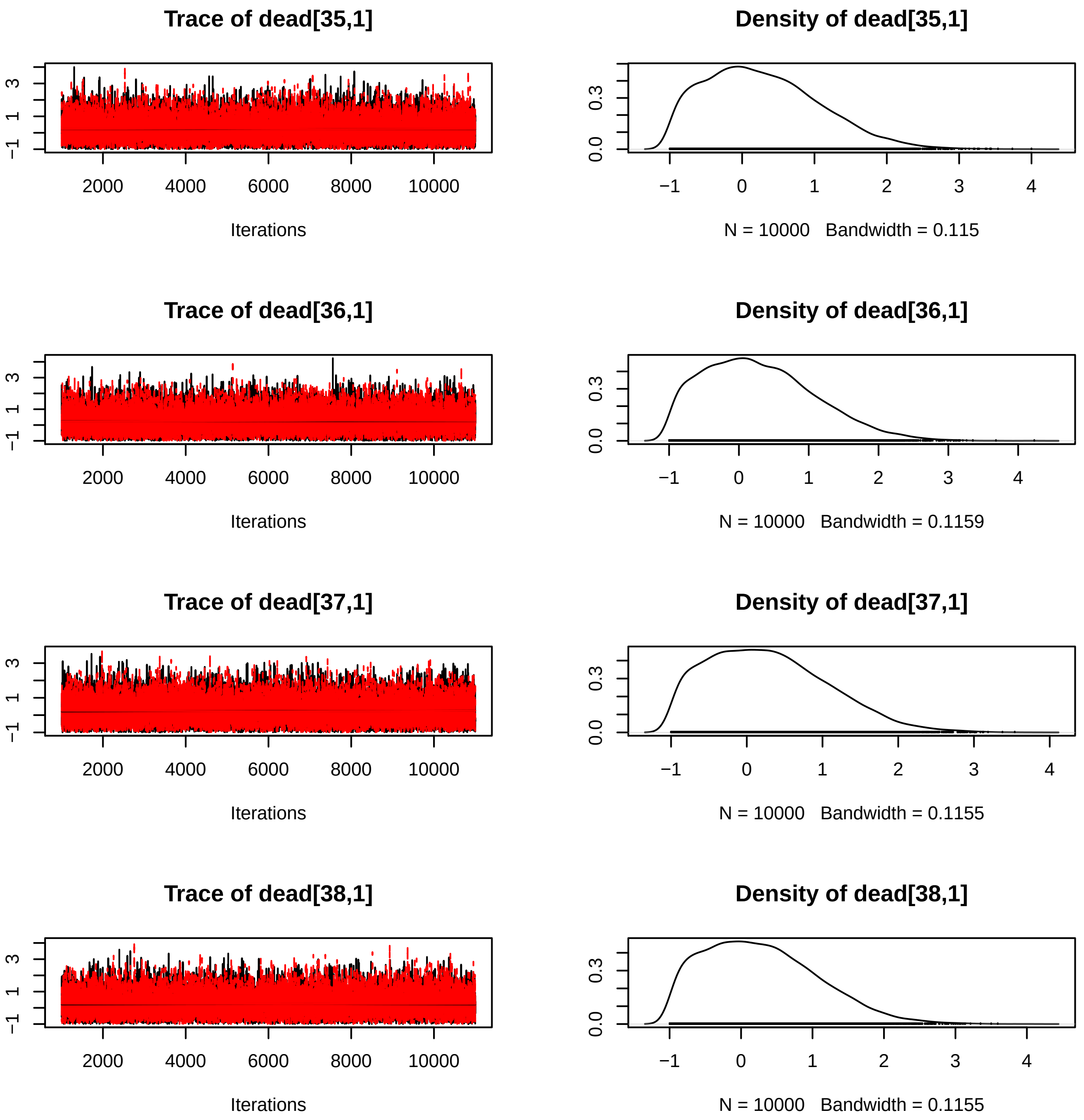

Trace of dead[35,1]
Iterations
Density of dead[35,1]
N = 10000 Bandwidth = 0.115
Trace of dead[36,1]
Iterations
Density of dead[36,1]
N = 10000 Bandwidth = 0.1159
Trace of dead[37,1]
Iterations
Density of dead[37,1]
N = 10000 Bandwidth = 0.1155
Trace of dead[38,1]
Iterations
Density of dead[38,1]
N = 10000 Bandwidth = 0.1155

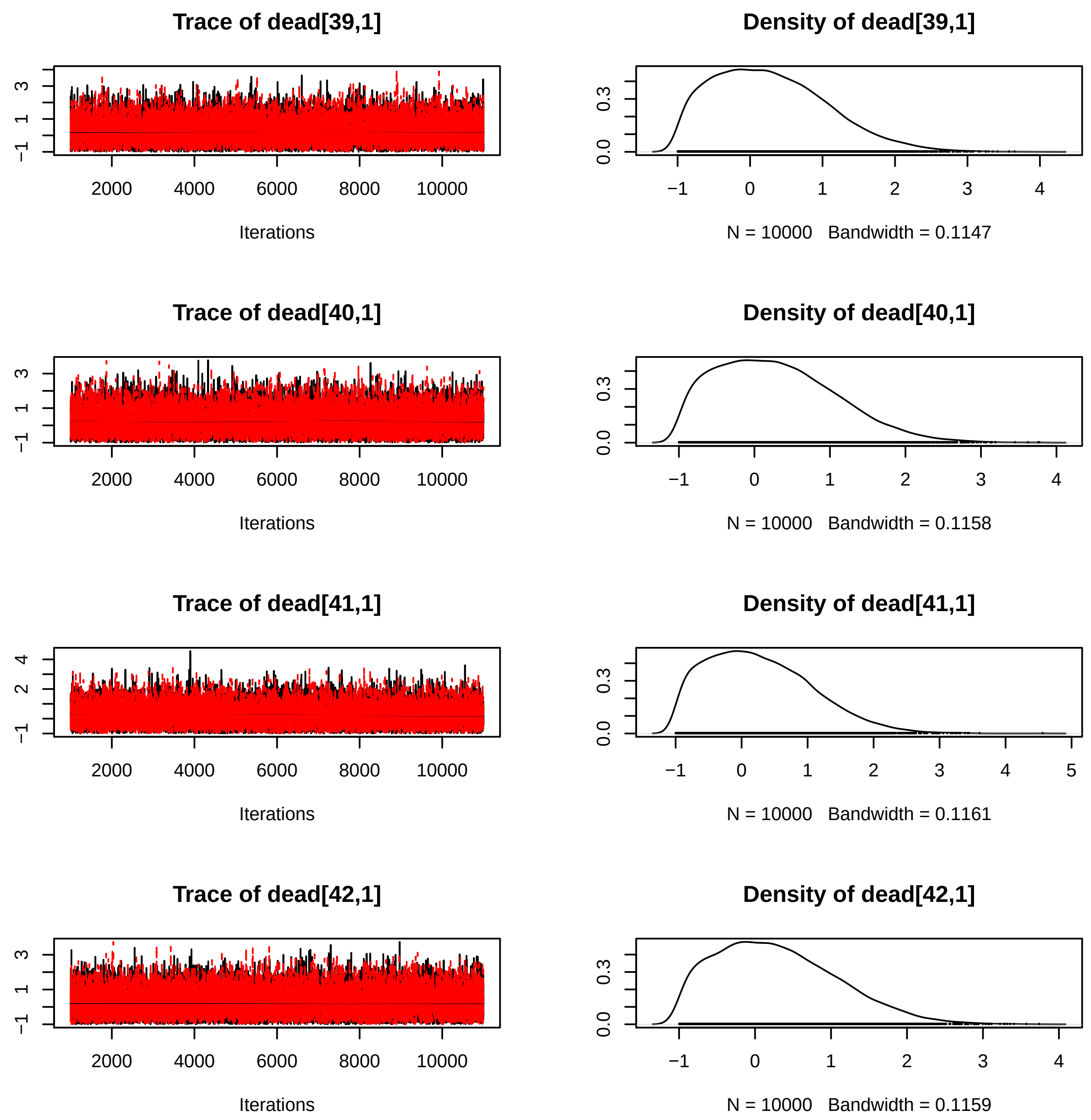
Trace of dead[39,1]
Density of dead[39,1]
Iterations
N = 10000 Bandwidth = 0.1147
Trace of dead[40,1]
Density of dead[40,1]
Iterations
N = 10000 Bandwidth = 0.1158
Trace of dead[41,1]
Density of dead[41,1]
Iterations
N = 10000 Bandwidth = 0.1161
Trace of dead[42,1]
Density of dead[42,1]
Iterations
N = 10000 Bandwidth = 0.1159

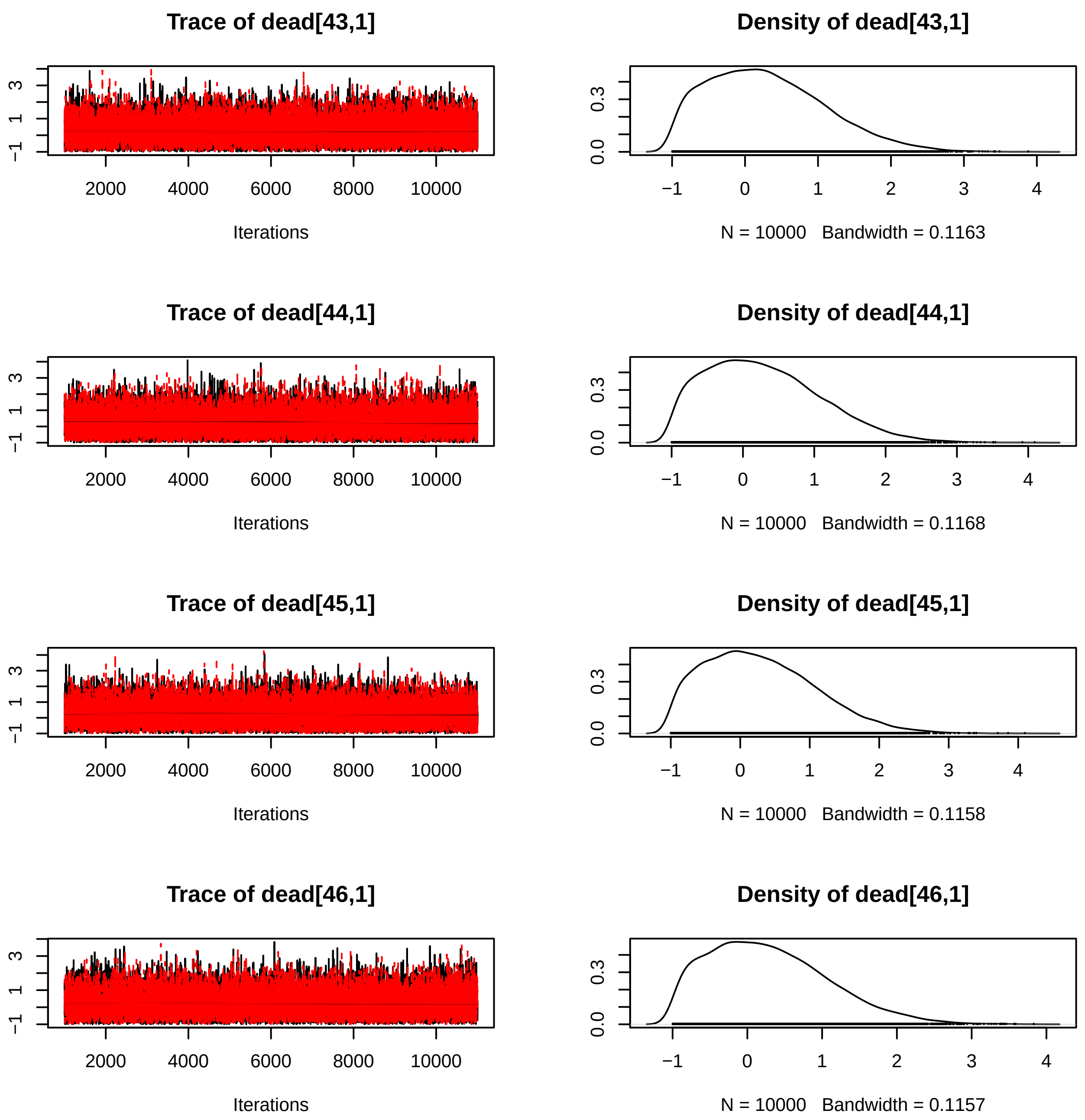
Trace of dead[43,1]
Iterations
Density of dead[43,1]
N = 10000 Bandwidth = 0.1163
Trace of dead[44,1]
Iterations
Density of dead[44,1]
N = 10000 Bandwidth = 0.1168
Trace of dead[45,1]
Iterations
Density of dead[45,1]
N = 10000 Bandwidth = 0.1158
Trace of dead[46,1]
Iterations
Density of dead[46,1]
N = 10000 Bandwidth = 0.1157

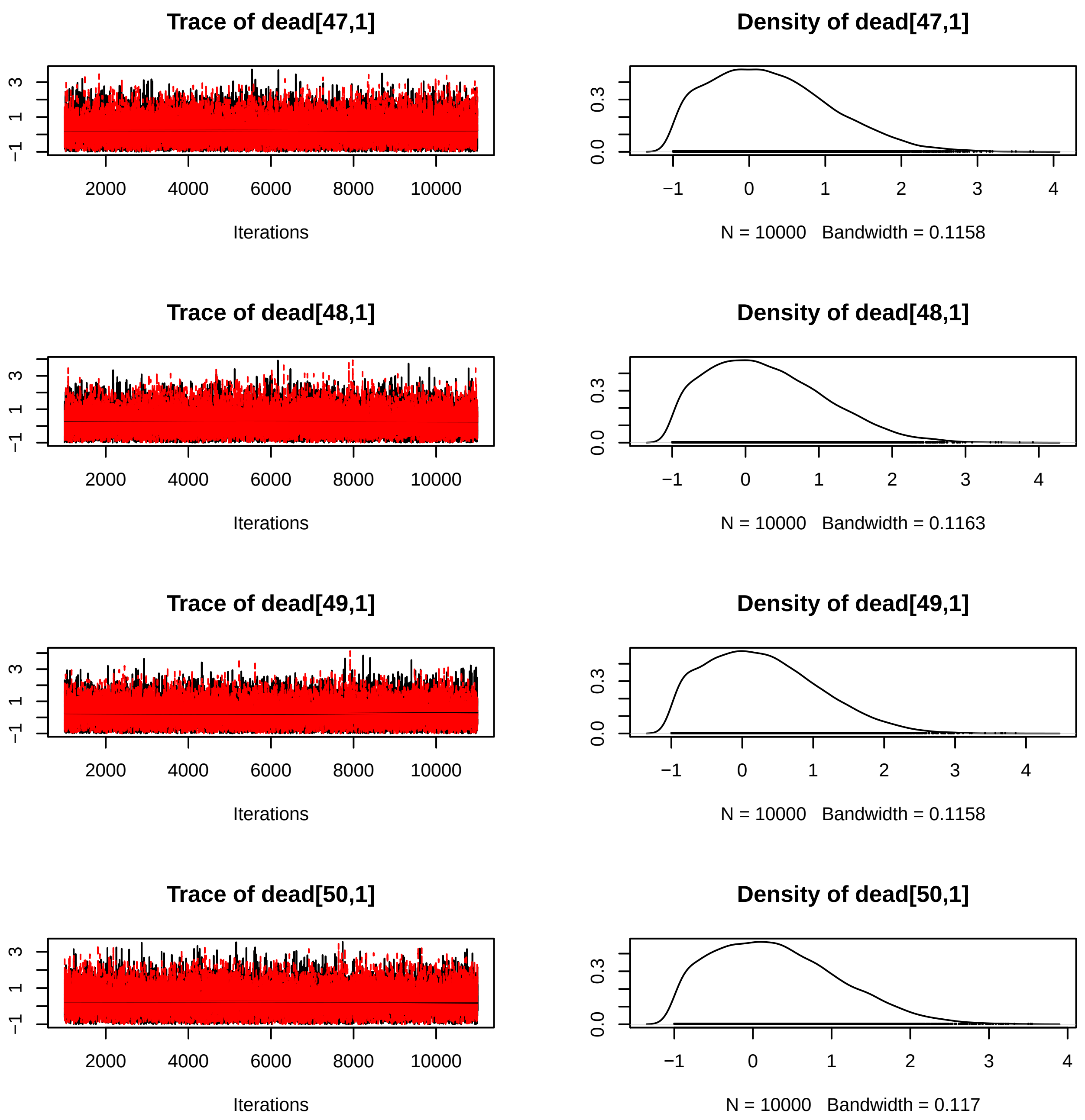
Trace of dead[47,1]
Iterations
Density of dead[47,1]
N = 10000 Bandwidth = 0.1158
Trace of dead[48,1]
Iterations
Density of dead[48,1]
N = 10000 Bandwidth = 0.1163
Trace of dead[49,1]
Iterations
Density of dead[49,1]
N = 10000 Bandwidth = 0.1158
Trace of dead[50,1]
Iterations
Density of dead[50,1]
N = 10000 Bandwidth = 0.117

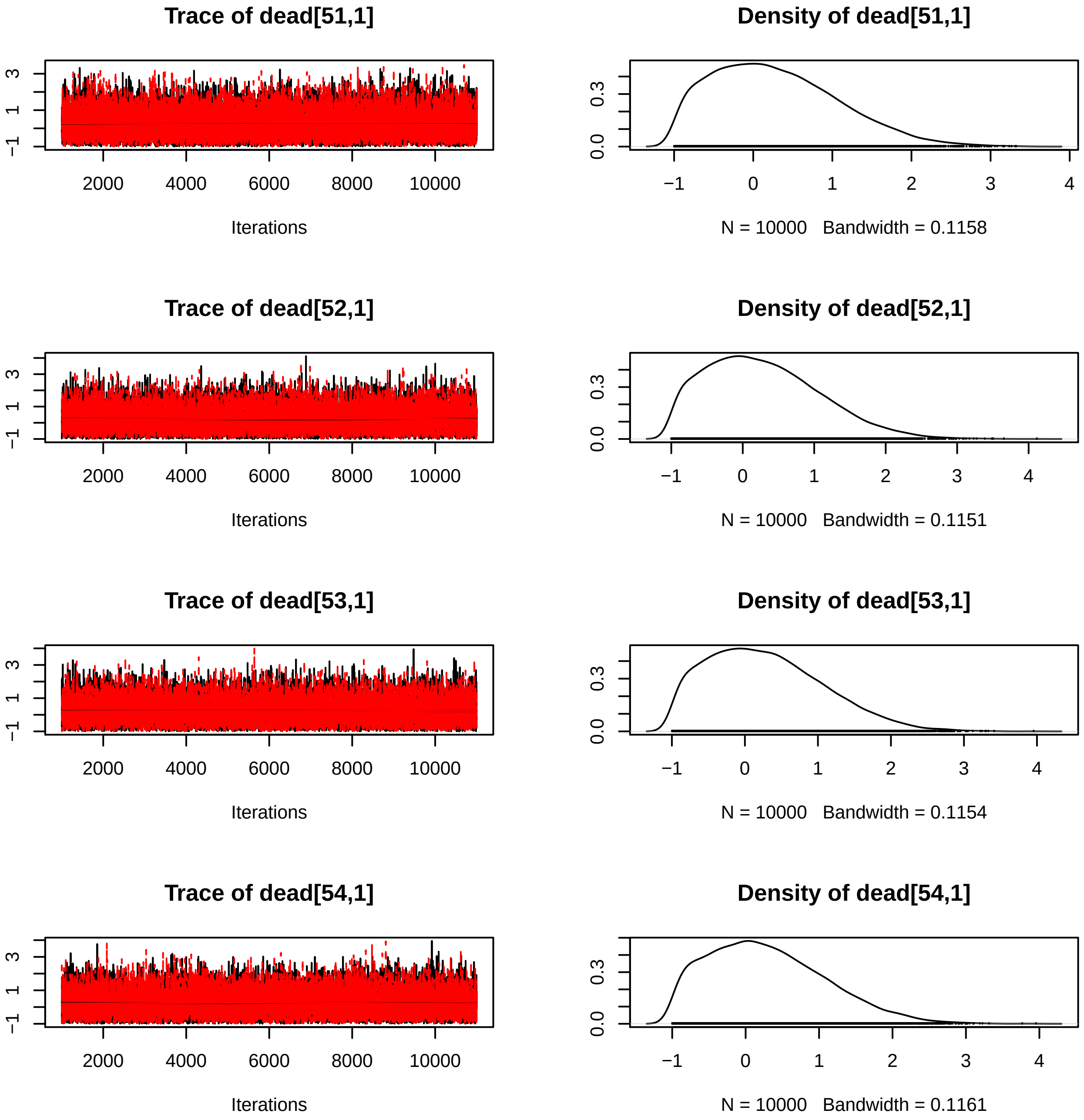
Trace of dead[51,1]
Iterations
Density of dead[51,1]
N = 10000   Bandwidth = 0.1158
Trace of dead[52,1]
Iterations
Density of dead[52,1]
N = 10000   Bandwidth = 0.1151
Trace of dead[53,1]
Iterations
Density of dead[53,1]
N = 10000   Bandwidth = 0.1154
Trace of dead[54,1]
Iterations
Density of dead[54,1]
N = 10000   Bandwidth = 0.1161

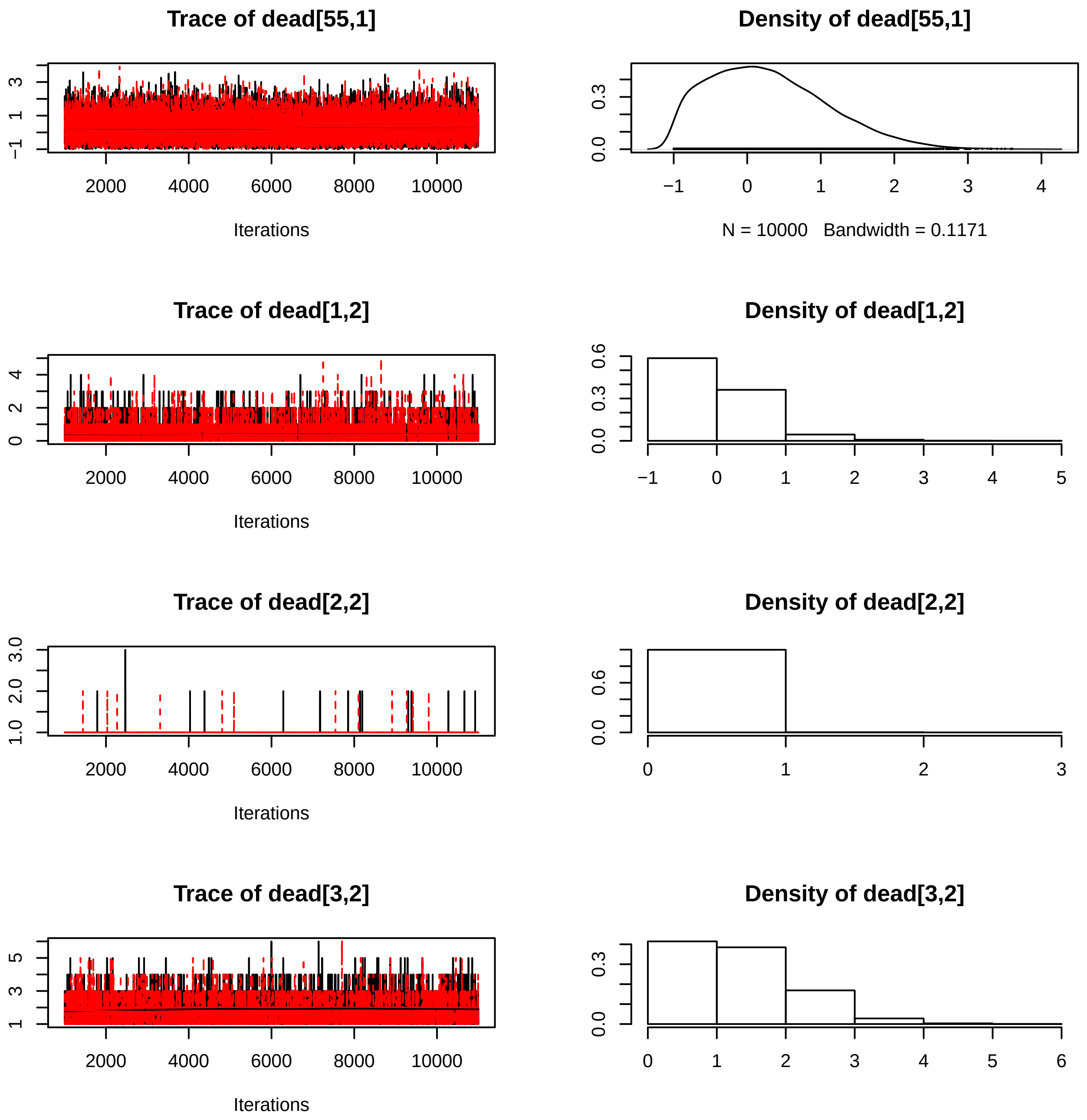
Trace of dead[55,1]
Iterations
Density of dead[55,1]
N = 10000 Bandwidth = 0.1171
Trace of dead[1,2]
Iterations
Density of dead[1,2]
Trace of dead[2,2]
Iterations
Density of dead[2,2]
Trace of dead[3,2]
Iterations
Density of dead[3,2]

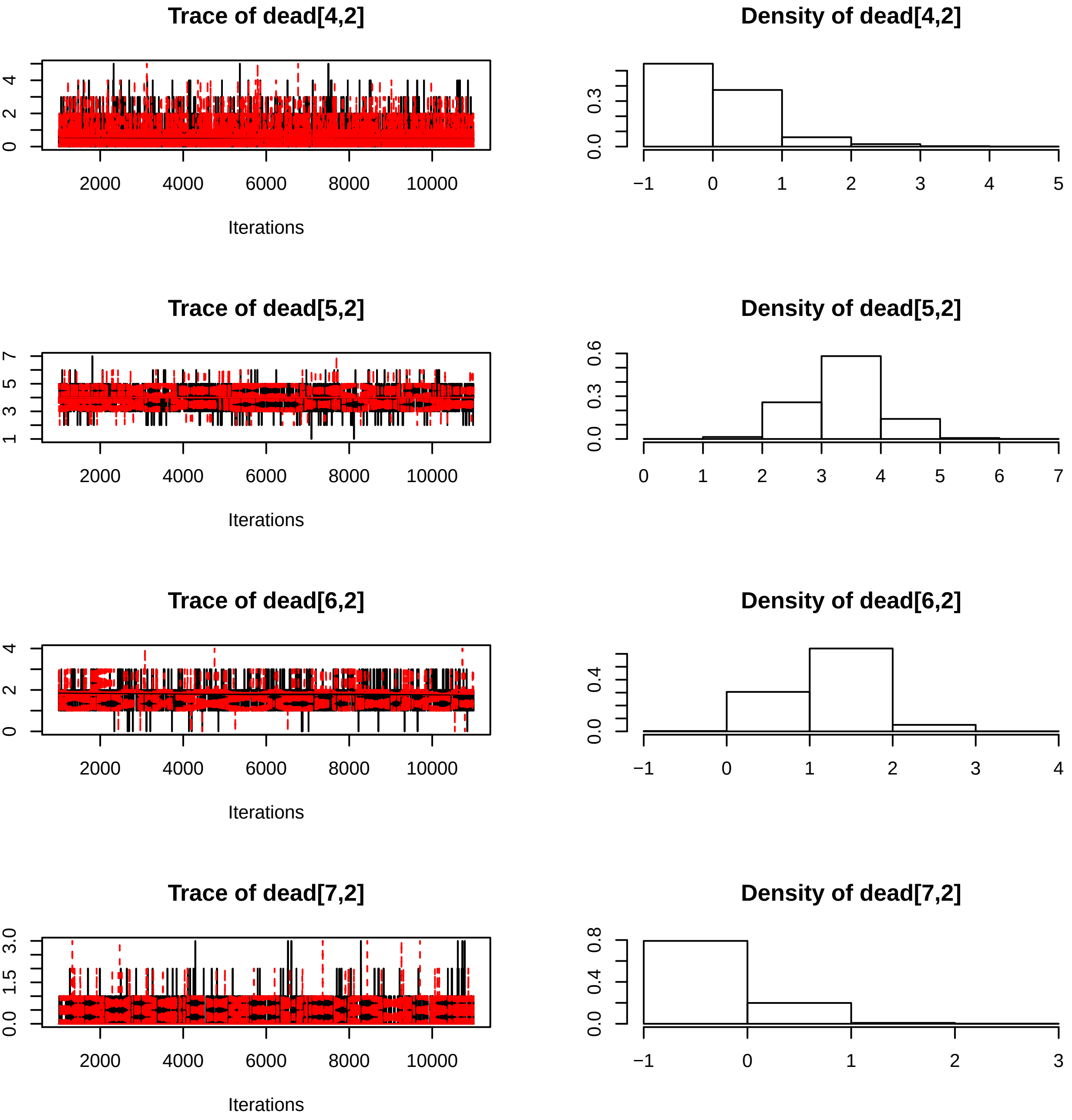
Trace of dead[4,2]
Density of dead[4,2]
Trace of dead[5,2]
Density of dead[5,2]
Trace of dead[6,2]
Density of dead[6,2]
Trace of dead[7,2]
Density of dead[7,2]
Iterations
2000
4000
6000
8000
10000

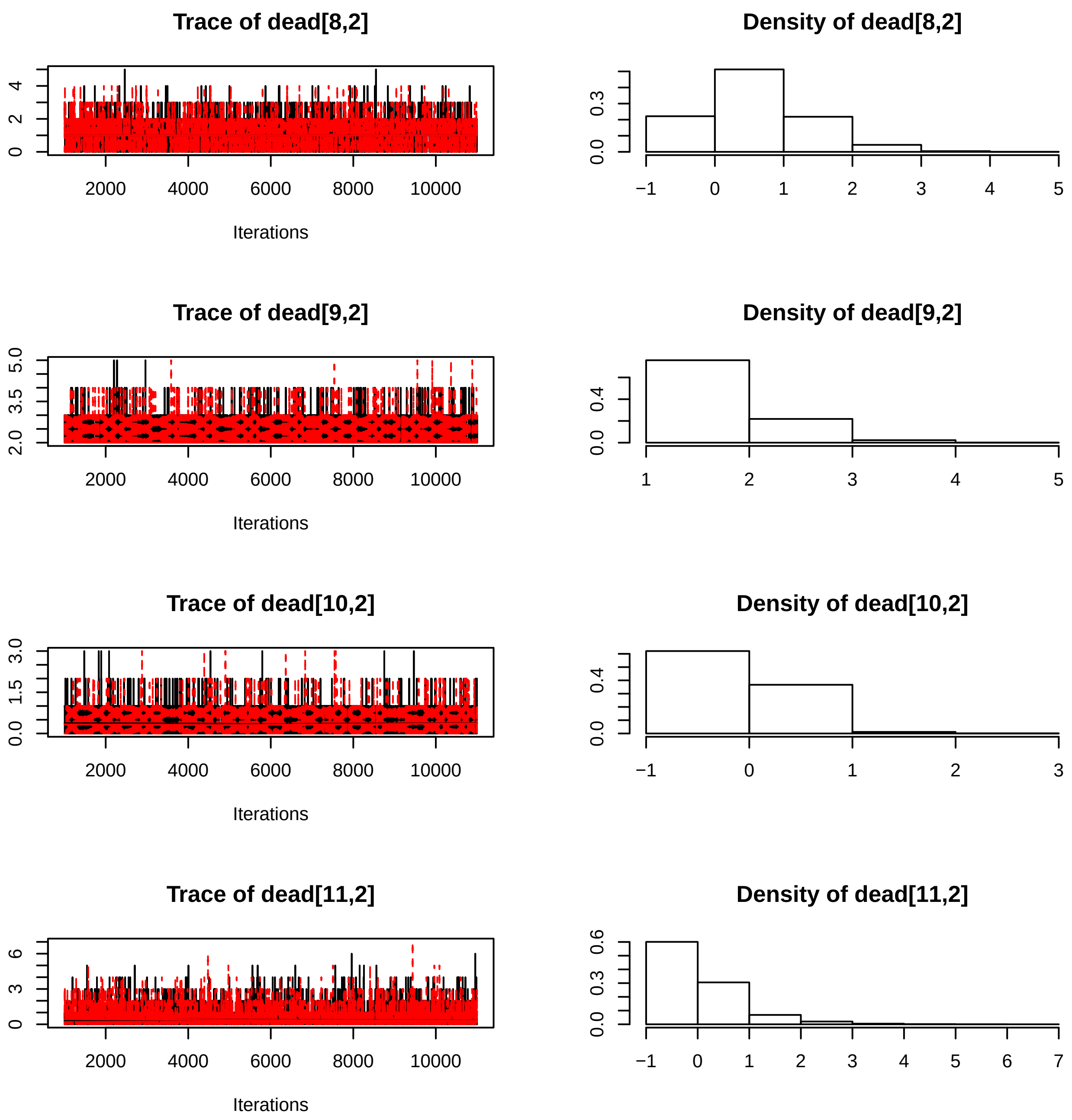

Trace of dead[8,2]
Density of dead[8,2]
Trace of dead[9,2]
Density of dead[9,2]
Trace of dead[10,2]
Density of dead[10,2]
Trace of dead[11,2]
Density of dead[11,2]
Iterations

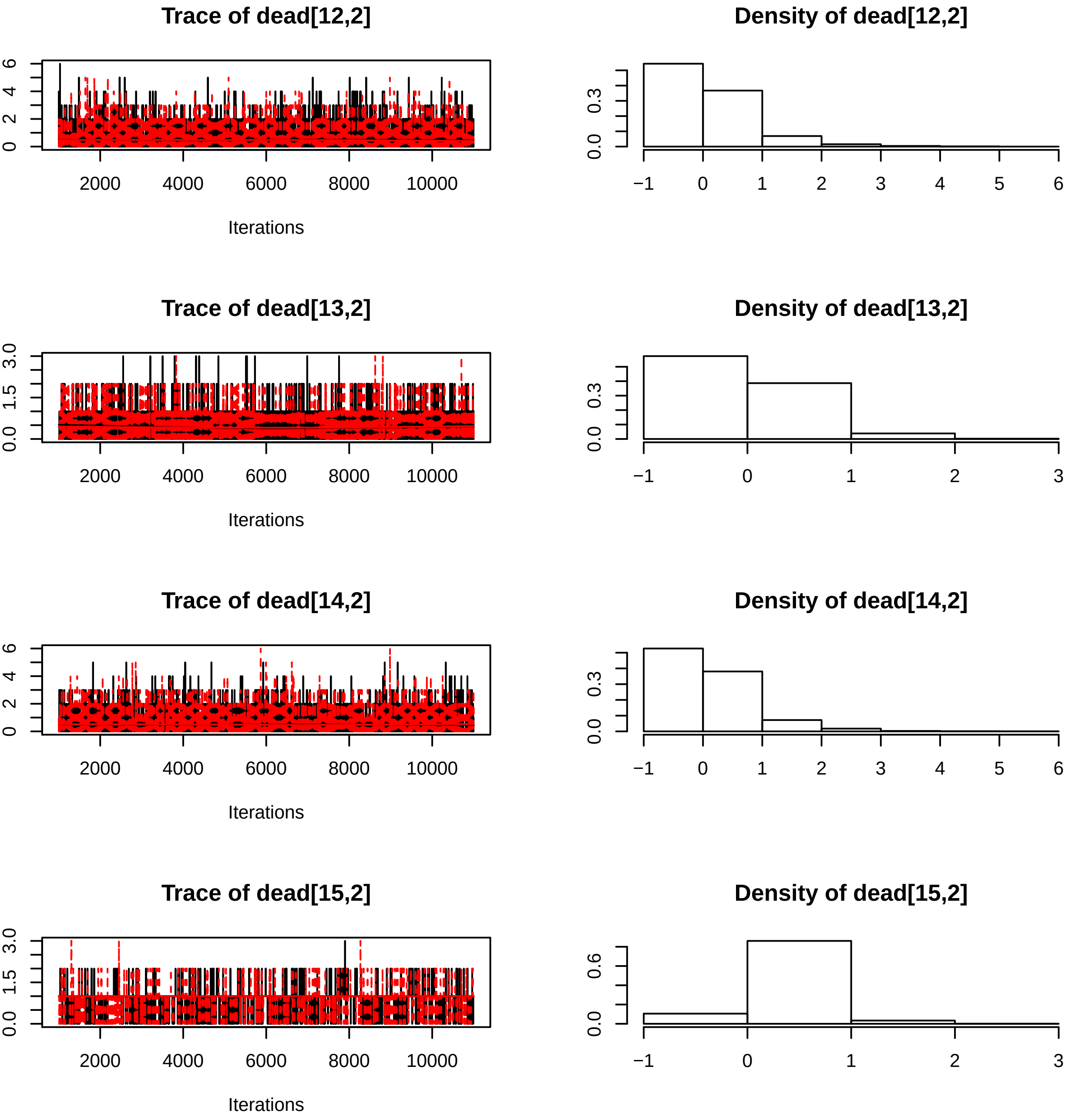
Trace of dead[12,2]
Density of dead[12,2]
Trace of dead[13,2]
Density of dead[13,2]
Trace of dead[14,2]
Density of dead[14,2]
Trace of dead[15,2]
Density of dead[15,2]
Iterations
2000
4000
6000
8000
10000

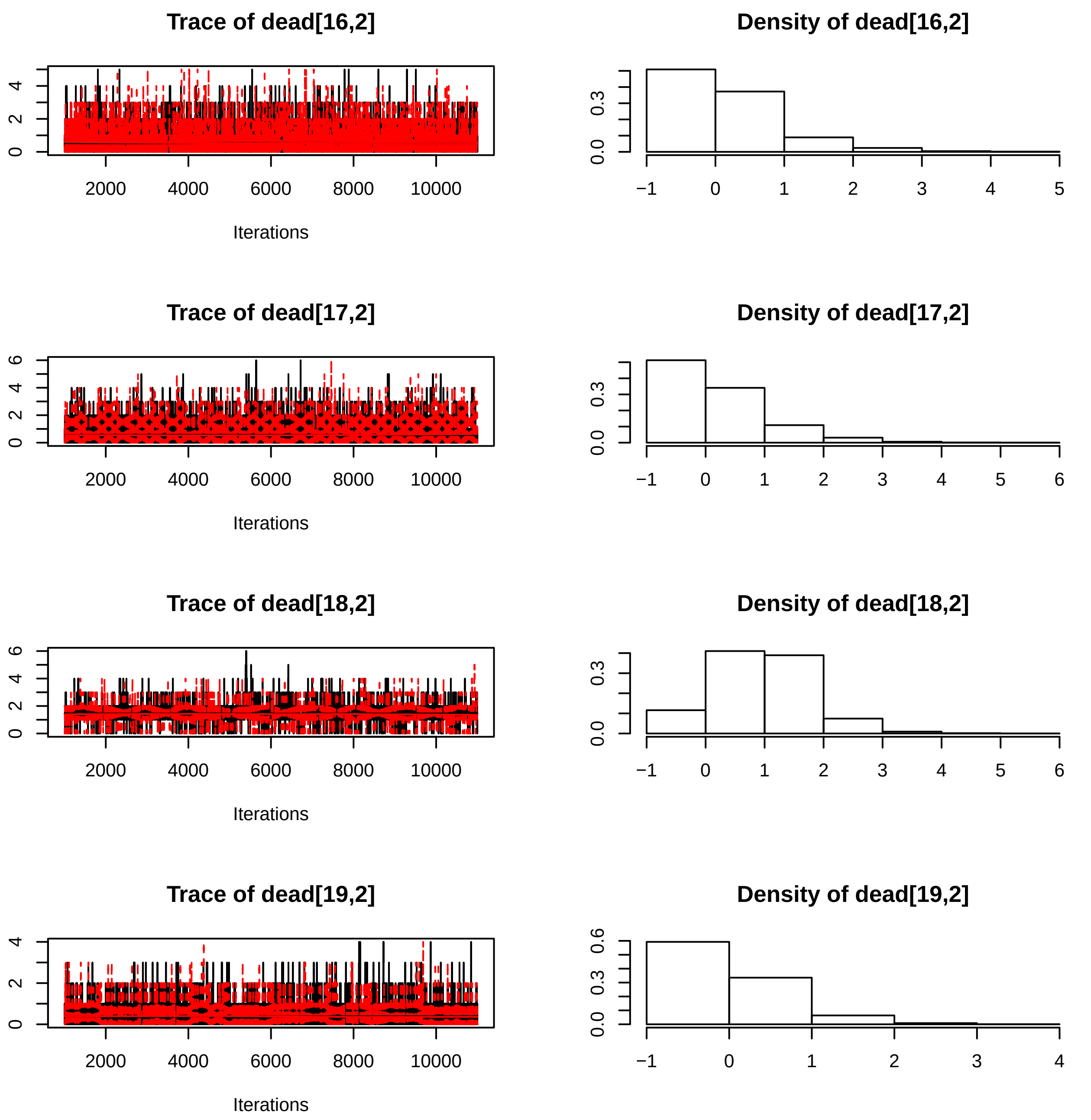
Trace of dead[16,2]
Iterations
Density of dead[16,2]
Trace of dead[17,2]
Iterations
Density of dead[17,2]
Trace of dead[18,2]
Iterations
Density of dead[18,2]
Trace of dead[19,2]
Iterations
Density of dead[19,2]

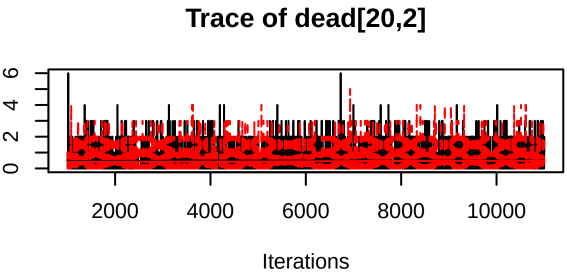
Trace of dead[20,2]
0
2
4
6
2000
4000
6000
8000
10000
Iterations

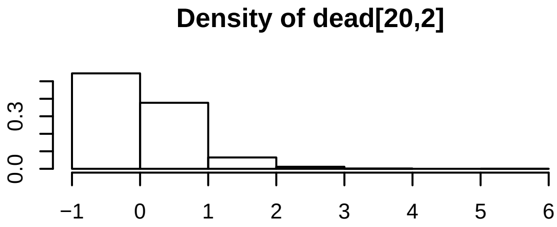
Density of dead[20,2]
0.0
0.3
−1
0
1
2
3
4
5
6

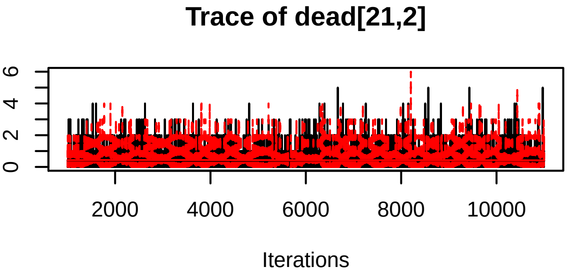
Trace of dead[21,2]
0
2
4
6
2000
4000
6000
8000
10000
Iterations

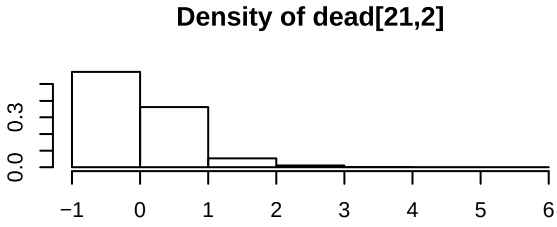
Density of dead[21,2]
0.0
0.3
−1
0
1
2
3
4
5
6

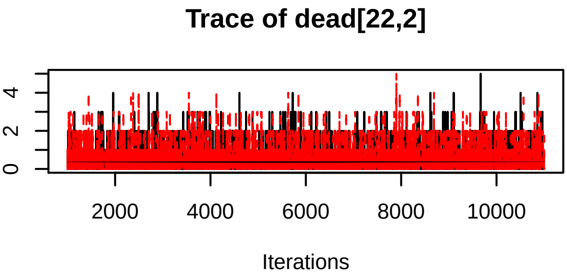
Trace of dead[22,2]
0
2
4
2000
4000
6000
8000
10000
Iterations

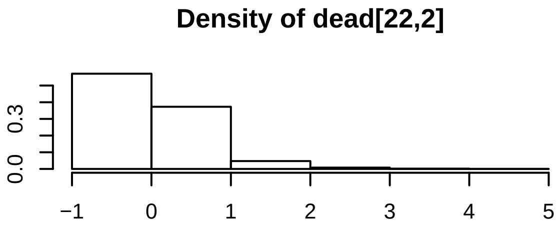
Density of dead[22,2]
0.0
0.3
−1
0
1
2
3
4
5

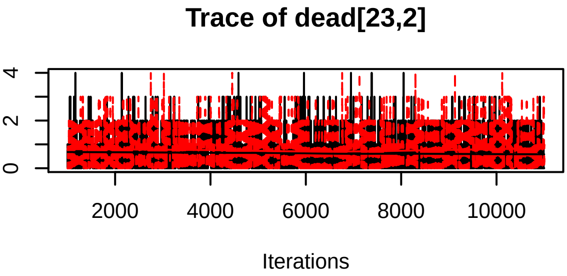
Trace of dead[23,2]
0
2
4
2000
4000
6000
8000
10000
Iterations

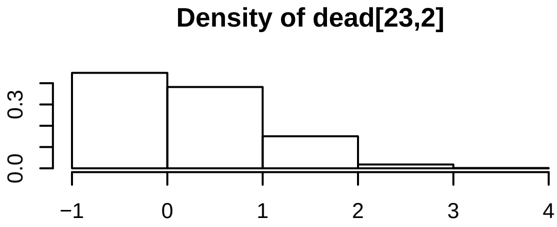
Density of dead[23,2]
0.0
0.3
−1
0
1
2
3
4

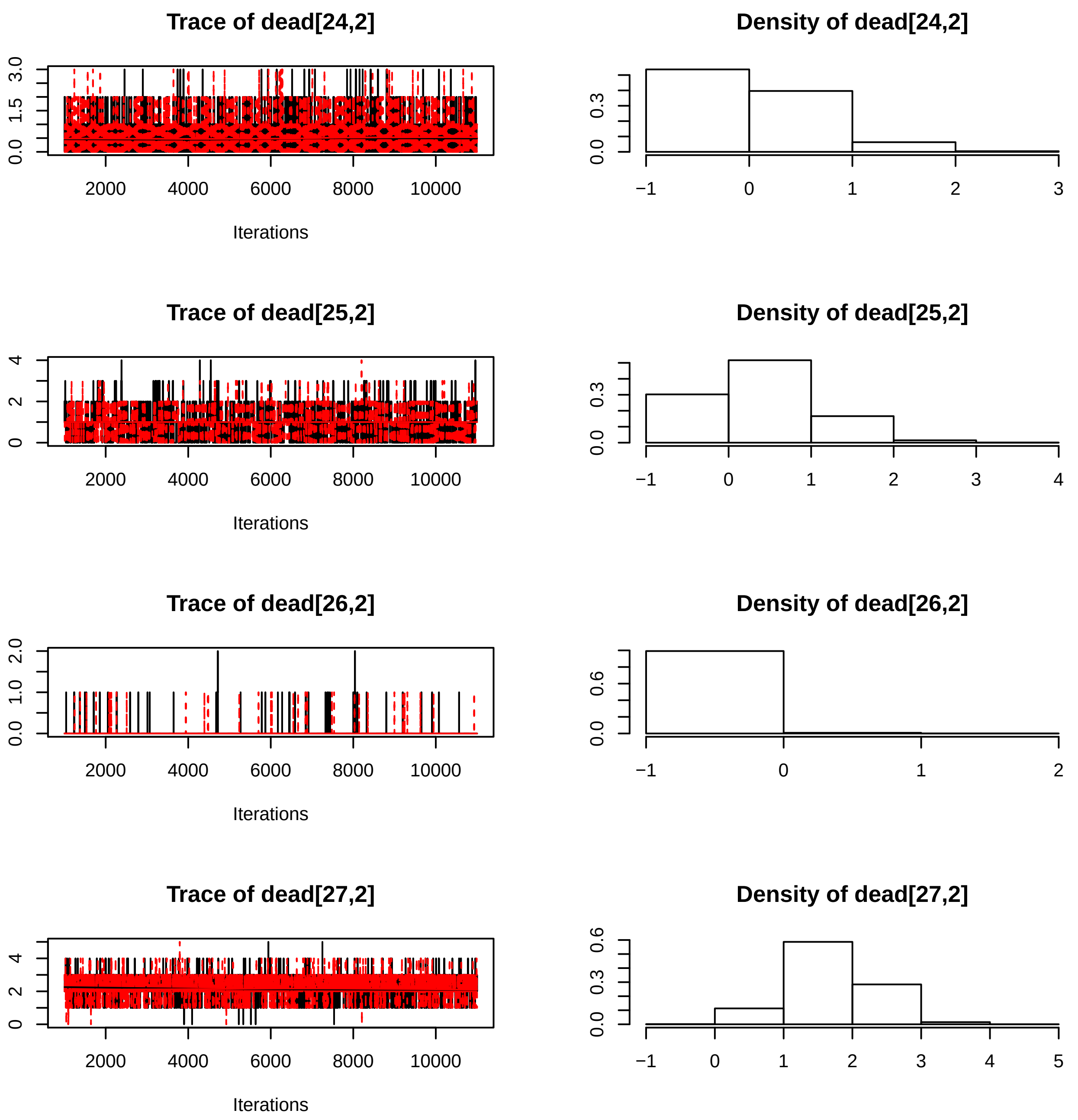
Trace of dead[24,2]
Density of dead[24,2]
Trace of dead[25,2]
Density of dead[25,2]
Trace of dead[26,2]
Density of dead[26,2]
Trace of dead[27,2]
Density of dead[27,2]
Iterations
Iterations
Iterations
Iterations
2000 4000 6000 8000 10000

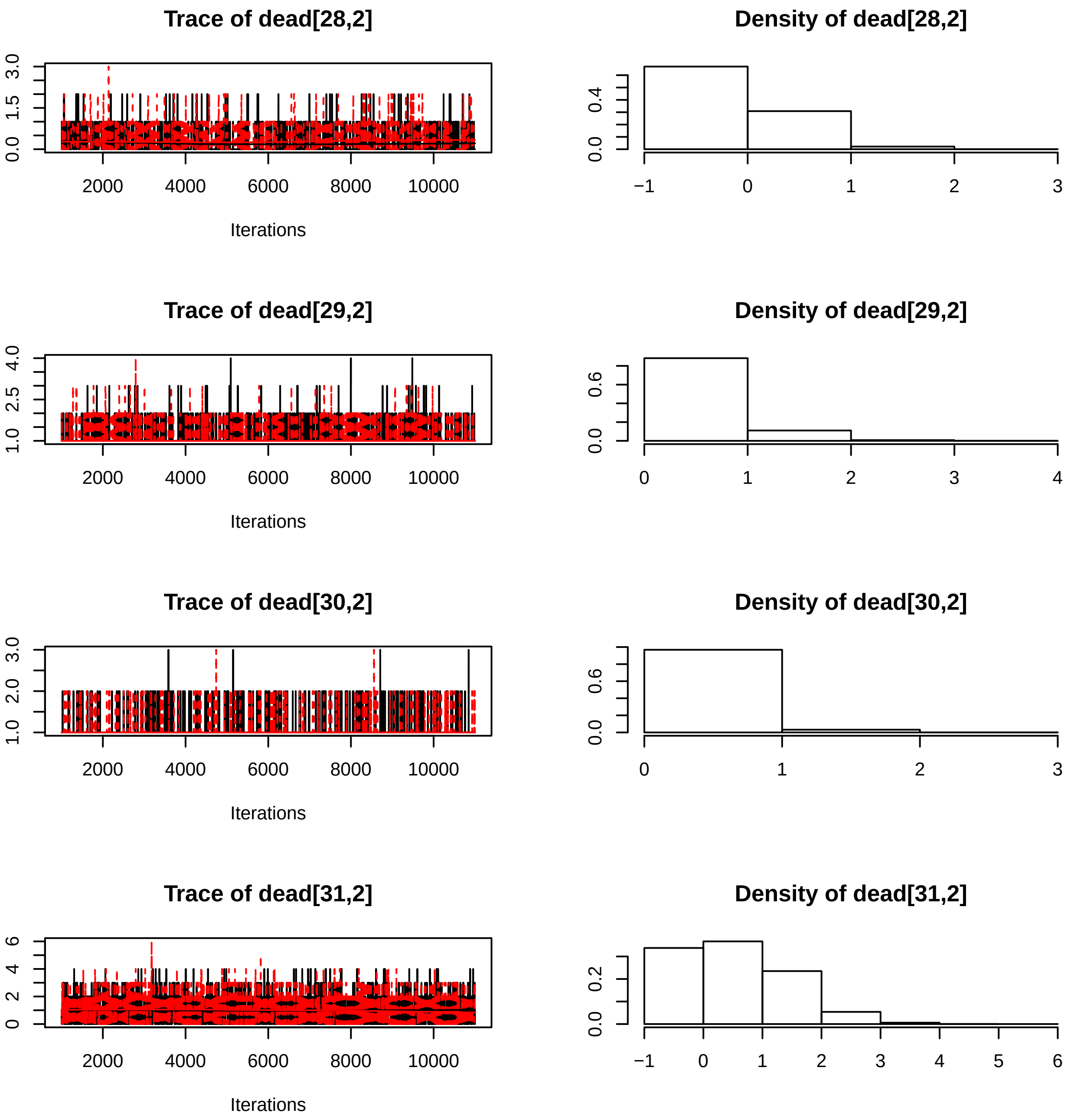
Trace of dead[28,2]
Iterations
Density of dead[28,2]
Trace of dead[29,2]
Iterations
Density of dead[29,2]
Trace of dead[30,2]
Iterations
Density of dead[30,2]
Trace of dead[31,2]
Iterations
Density of dead[31,2]
2000 4000 6000 8000 10000

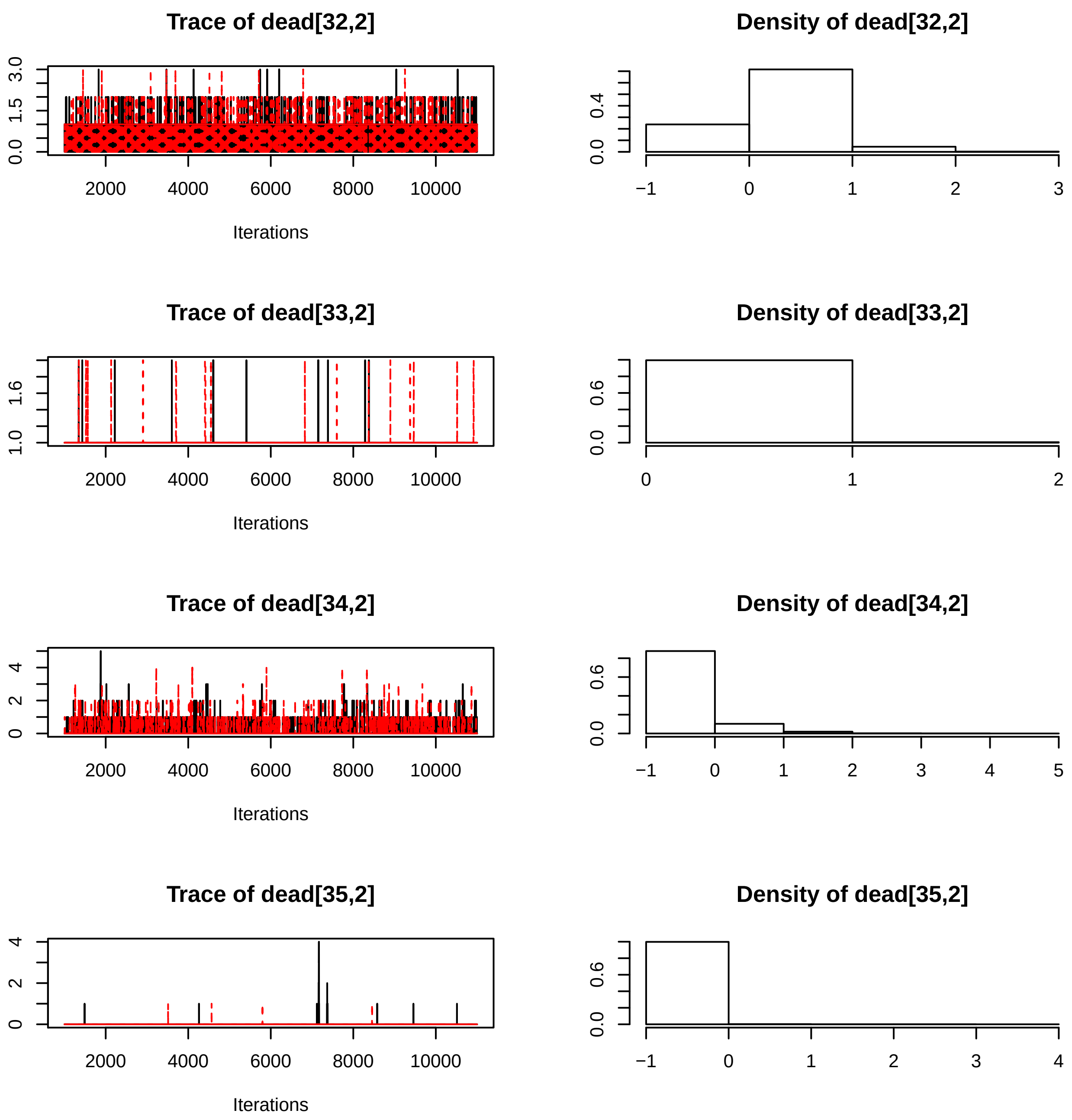
Trace of dead[32,2]
Density of dead[32,2]
Trace of dead[33,2]
Density of dead[33,2]
Trace of dead[34,2]
Density of dead[34,2]
Trace of dead[35,2]
Density of dead[35,2]
Iterations
2000
4000
6000
8000
10000
0.0
1.5
3.0
0.4
1.0
1.6
0.6
0
2
4
−1
1
3
5

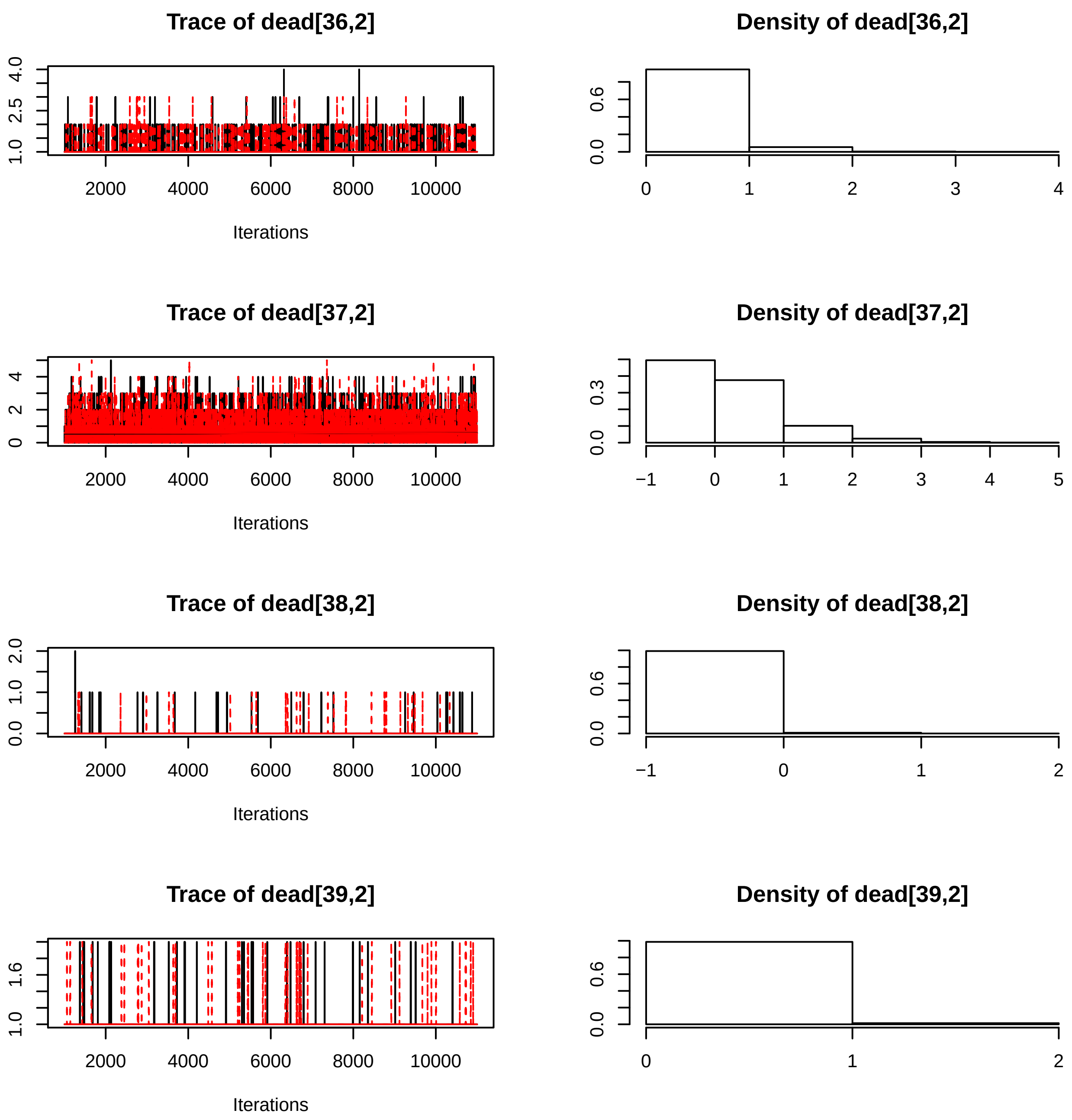

Trace of dead[36,2]
Density of dead[36,2]
Iterations
Trace of dead[37,2]
Density of dead[37,2]
Iterations
Trace of dead[38,2]
Density of dead[38,2]
Iterations
Trace of dead[39,2]
Density of dead[39,2]
Iterations
2000
4000
6000
8000
10000

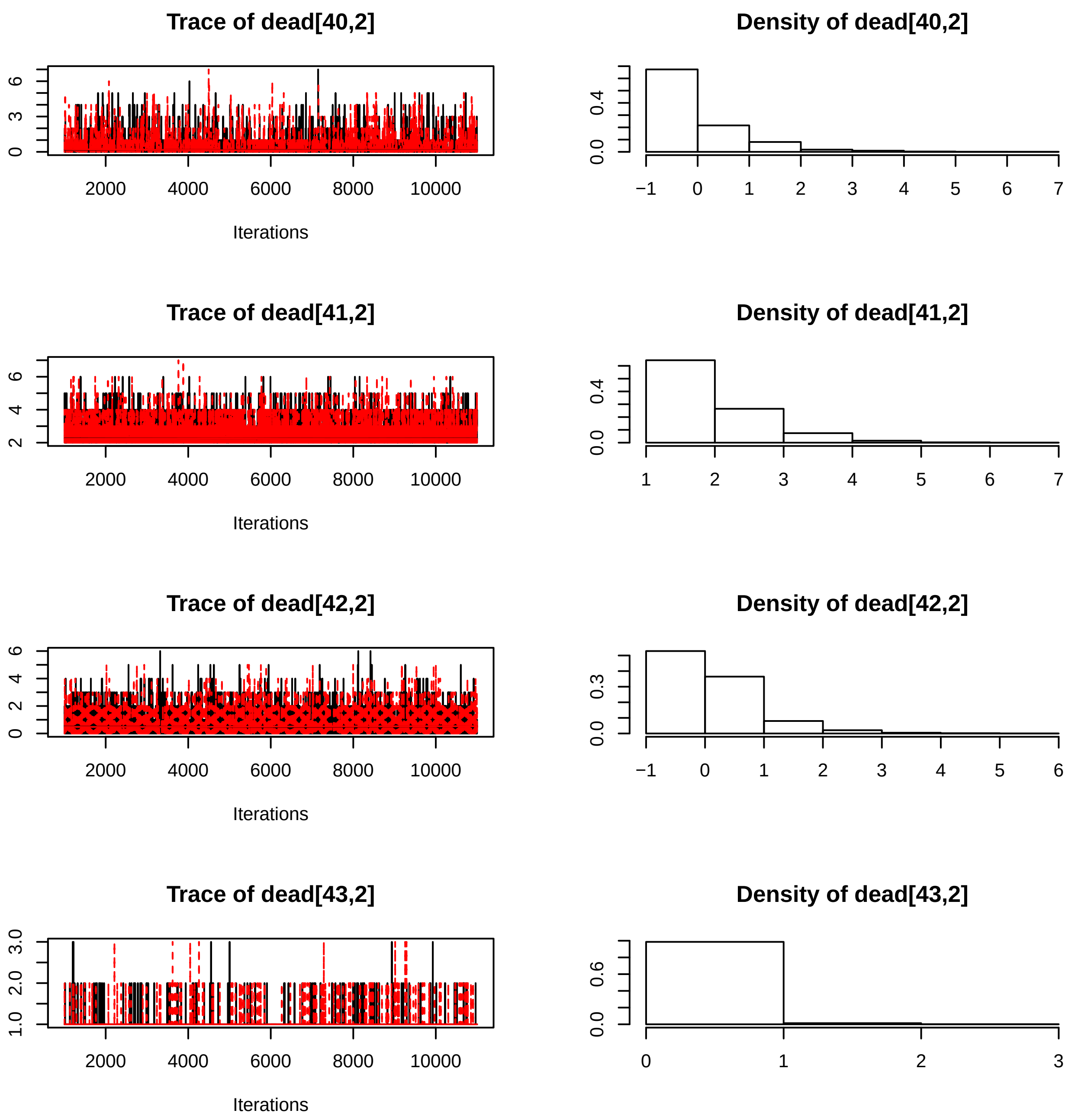
Trace of dead[40,2]
Density of dead[40,2]
0
3
6
2000
4000
6000
8000
10000
Iterations
0.0
0.4
−1
0
1
2
3
4
5
6
7
Trace of dead[41,2]
Density of dead[41,2]
2
4
6
2000
4000
6000
8000
10000
Iterations
0.0
0.4
1
2
3
4
5
6
7
Trace of dead[42,2]
Density of dead[42,2]
0
2
4
6
2000
4000
6000
8000
10000
Iterations
0.0
0.3
−1
0
1
2
3
4
5
6
Trace of dead[43,2]
Density of dead[43,2]
1.0
2.0
3.0
2000
4000
6000
8000
10000
Iterations
0.0
0.6
0
1
2
3

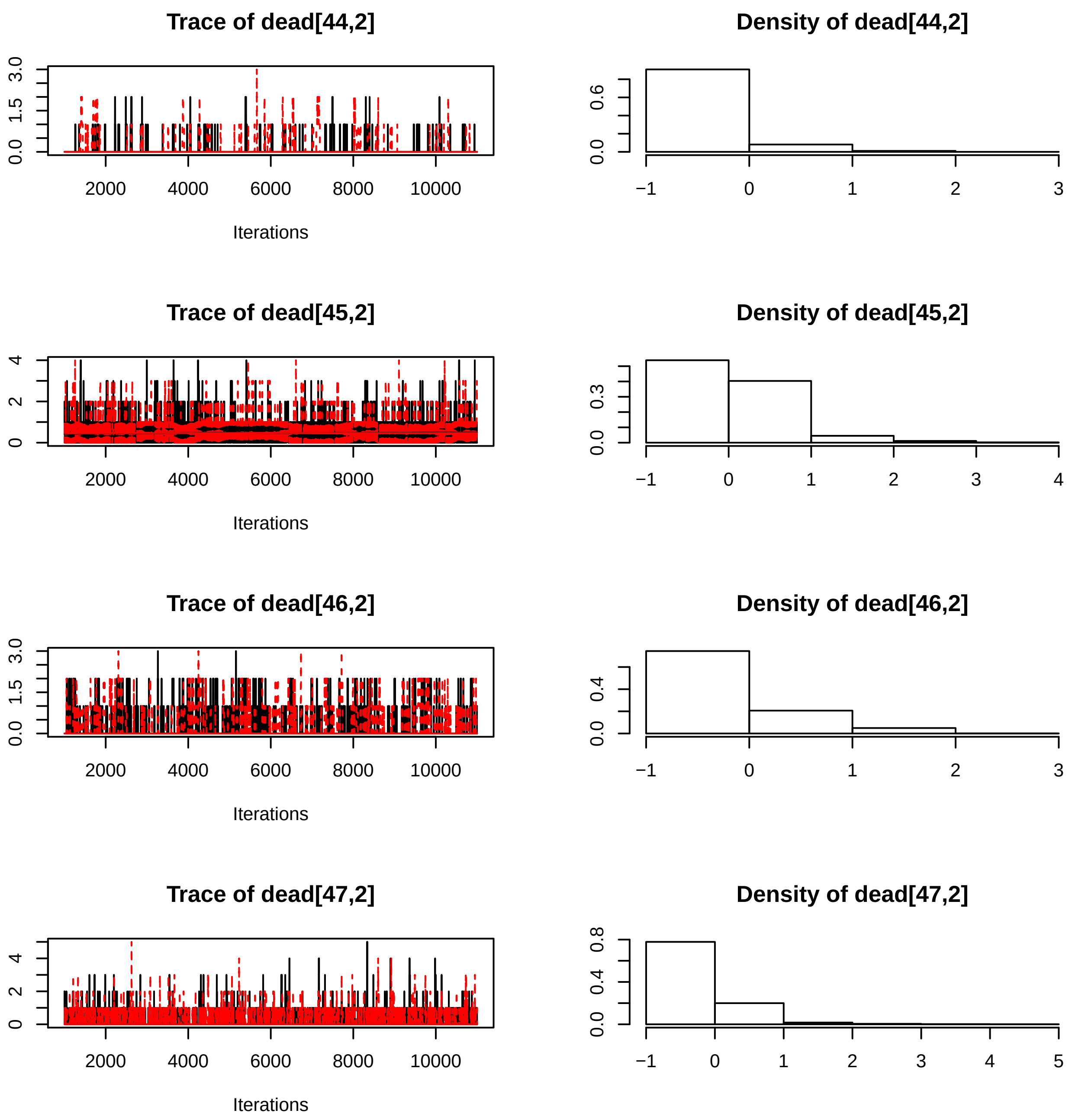
Trace of dead[44,2]
Density of dead[44,2]
Iterations
Trace of dead[45,2]
Density of dead[45,2]
Iterations
Trace of dead[46,2]
Density of dead[46,2]
Iterations
Trace of dead[47,2]
Density of dead[47,2]
Iterations

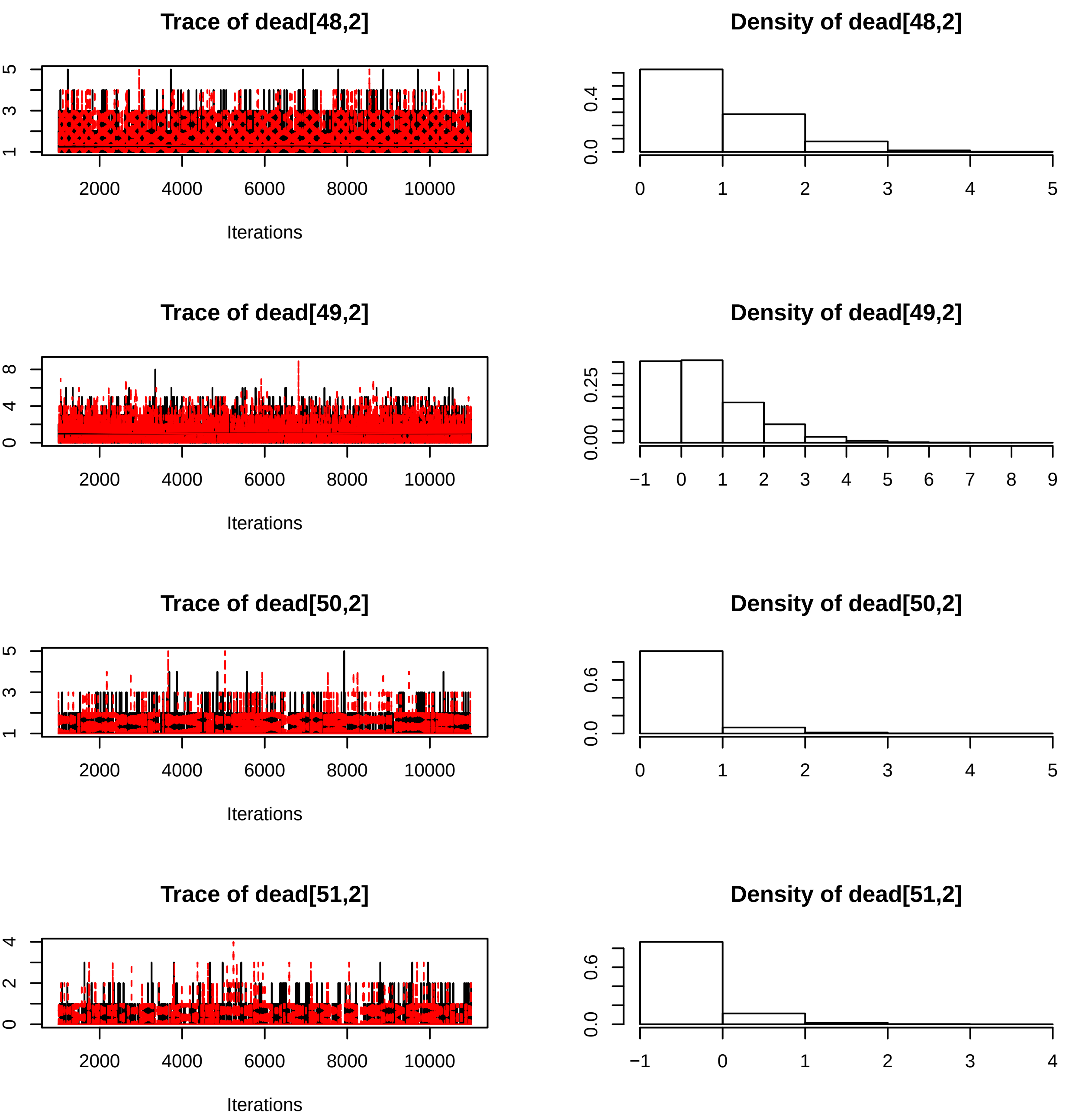

Trace of dead[48,2]
Iterations
Density of dead[48,2]
Trace of dead[49,2]
Iterations
Density of dead[49,2]
Trace of dead[50,2]
Iterations
Density of dead[50,2]
Trace of dead[51,2]
Iterations
Density of dead[51,2]

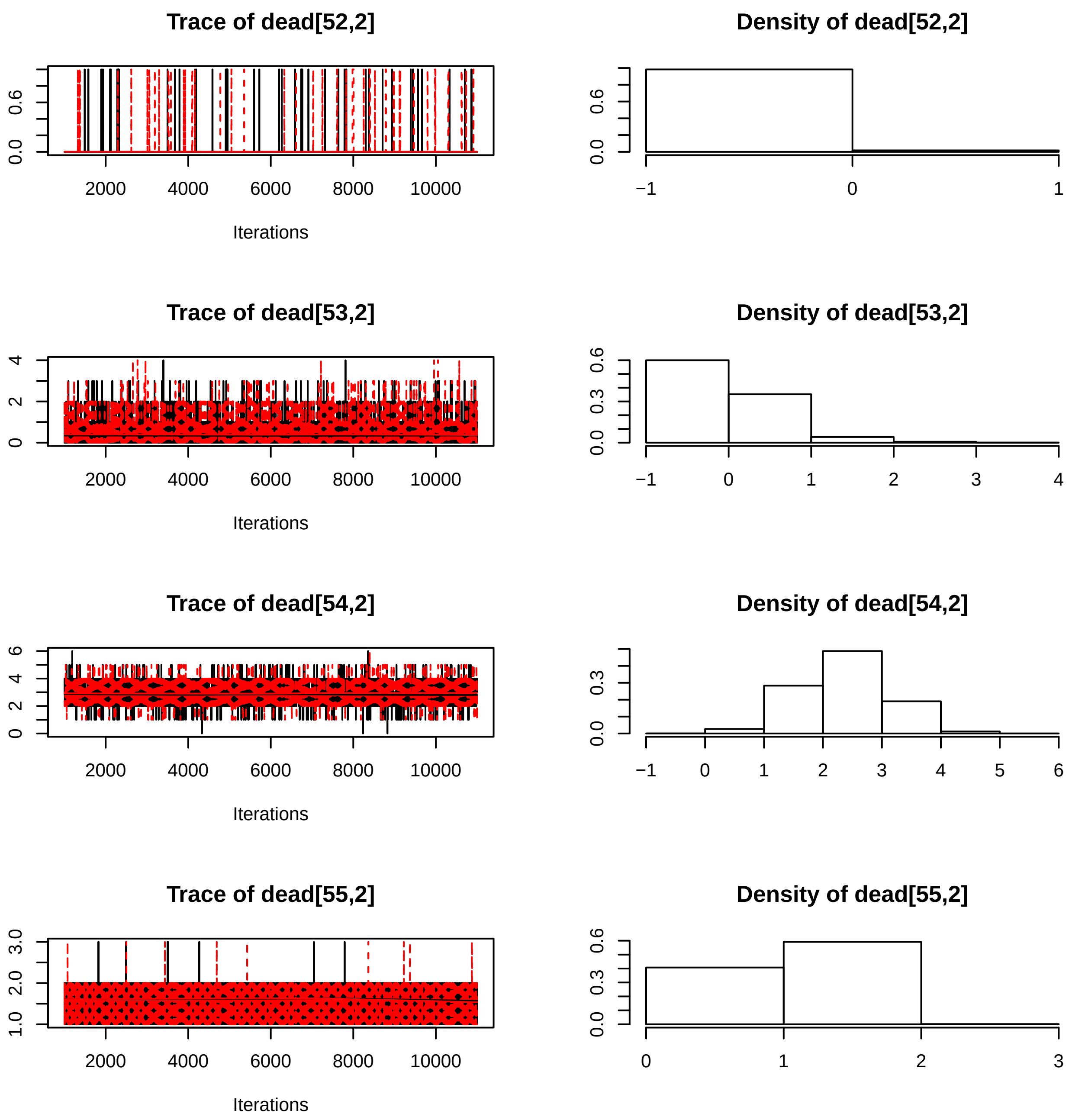

Trace of dead[52,2]
Iterations
Density of dead[52,2]
Trace of dead[53,2]
Iterations
Density of dead[53,2]
Trace of dead[54,2]
Iterations
Density of dead[54,2]
Trace of dead[55,2]
Iterations
Density of dead[55,2]

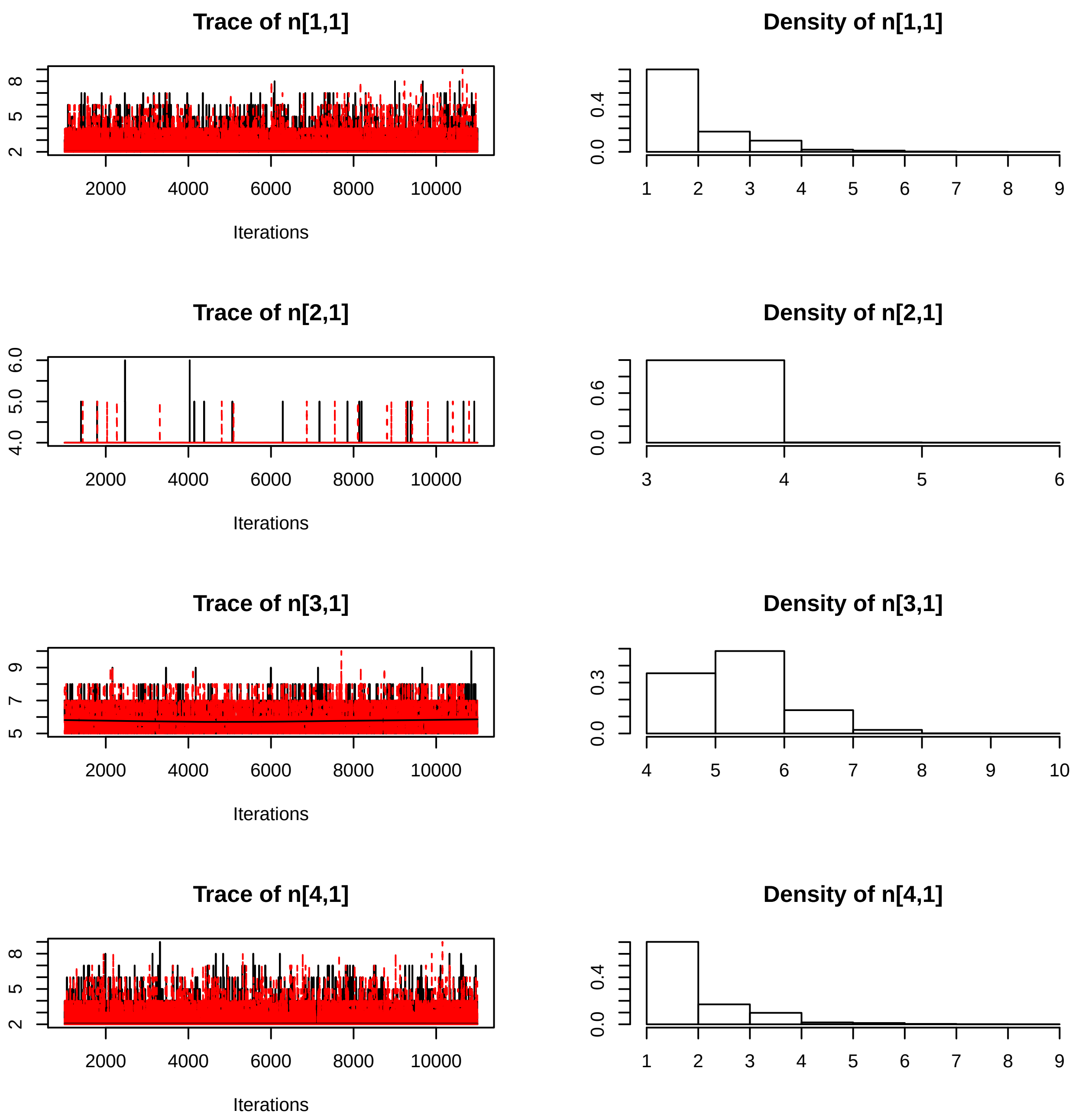
Trace of n[1,1]
Density of n[1,1]
Trace of n[2,1]
Density of n[2,1]
Trace of n[3,1]
Density of n[3,1]
Trace of n[4,1]
Density of n[4,1]
Iterations
2000
4000
6000
8000
10000

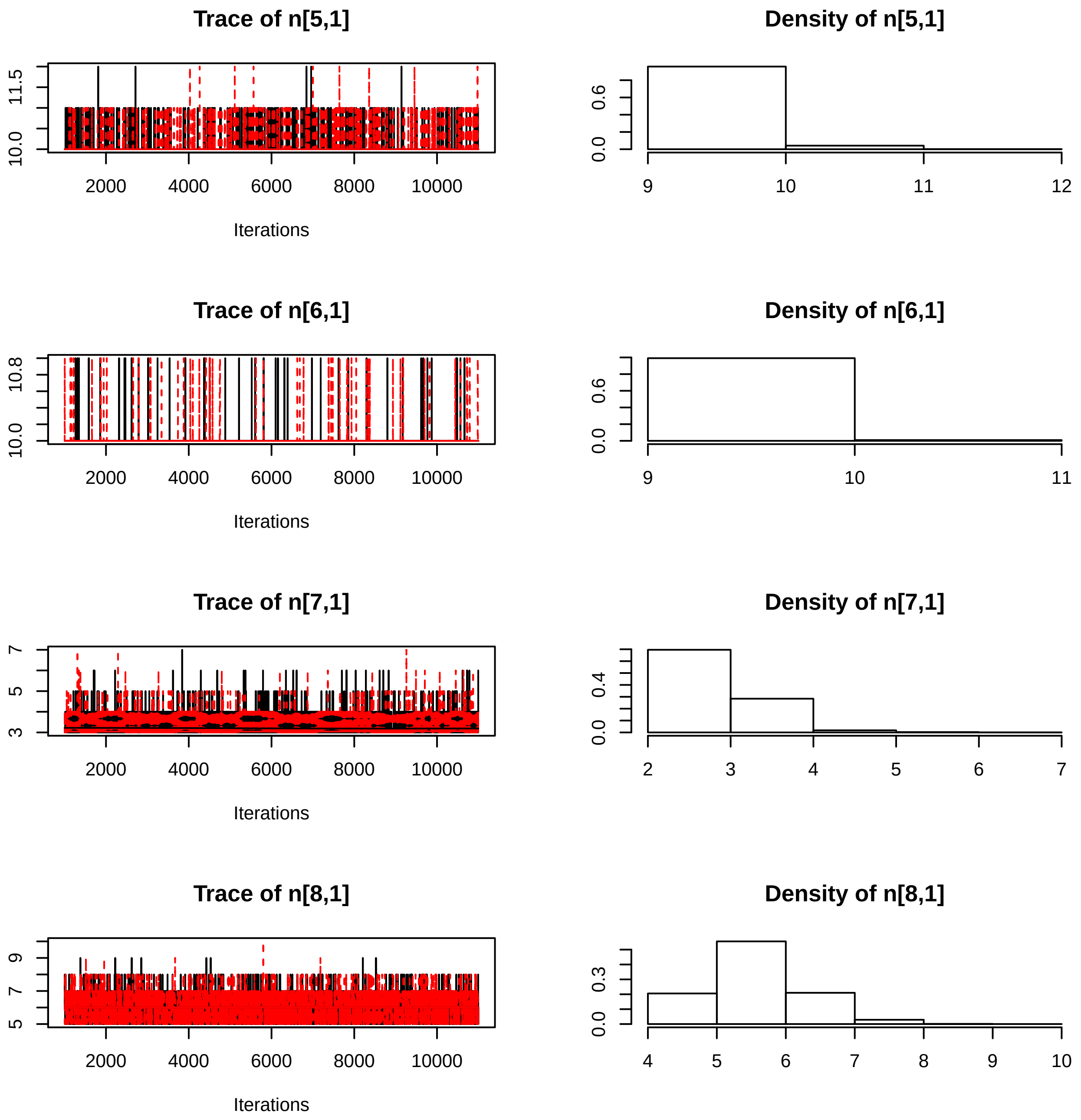
Trace of n[5,1]
11.5
10.0
2000
4000
6000
8000
10000
Iterations
Density of n[5,1]
0.6
0.0
9
10
11
12
Trace of n[6,1]
10.8
10.0
2000
4000
6000
8000
10000
Iterations
Density of n[6,1]
0.6
0.0
9
10
11
Trace of n[7,1]
7
5
3
2000
4000
6000
8000
10000
Iterations
Density of n[7,1]
0.4
0.0
2
3
4
5
6
7
Trace of n[8,1]
9
7
5
2000
4000
6000
8000
10000
Iterations
Density of n[8,1]
0.3
0.0
4
5
6
7
8
9
10

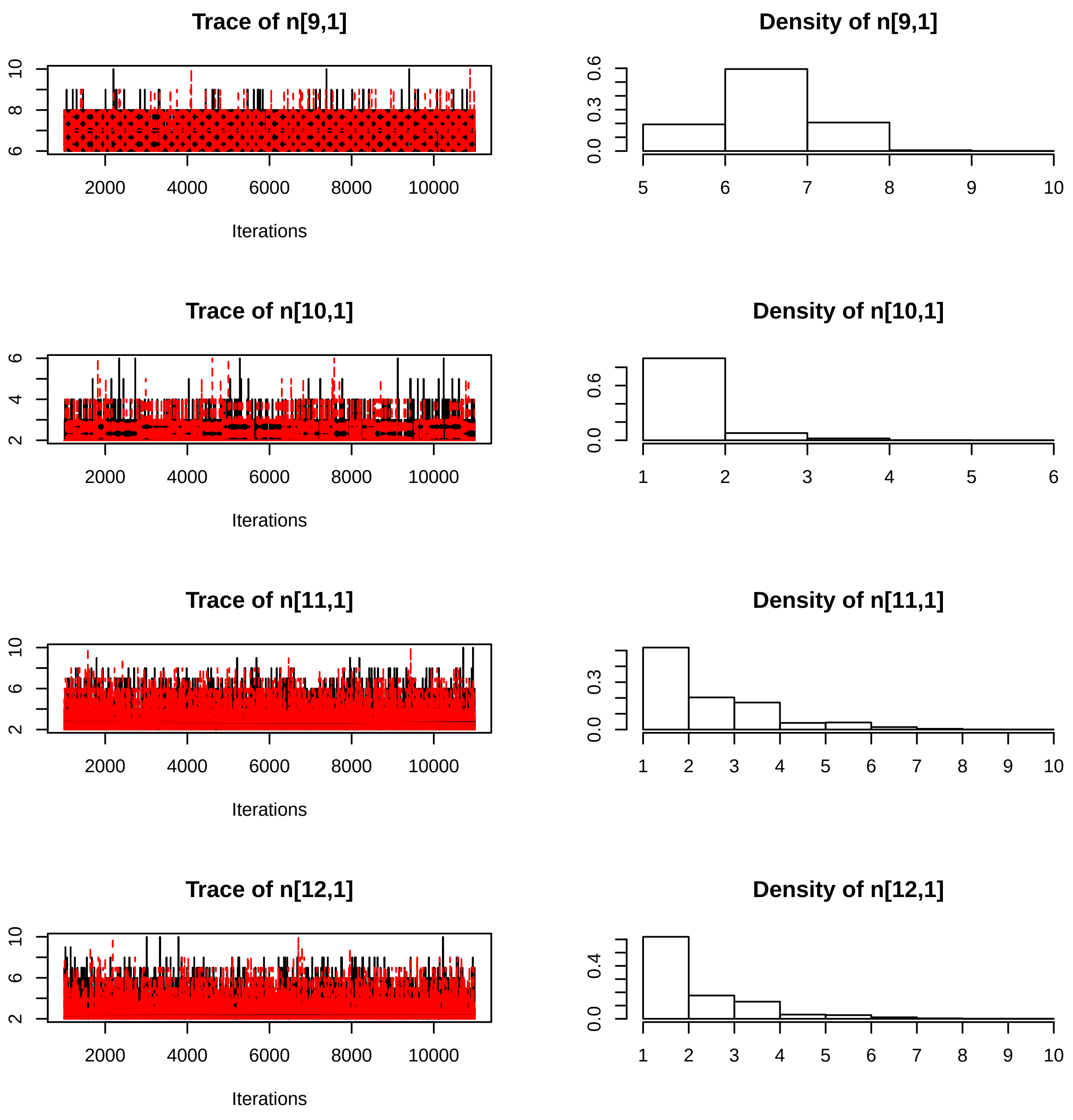
Trace of n[9,1]
Density of n[9,1]
Trace of n[10,1]
Density of n[10,1]
Trace of n[11,1]
Density of n[11,1]
Trace of n[12,1]
Density of n[12,1]
Iterations

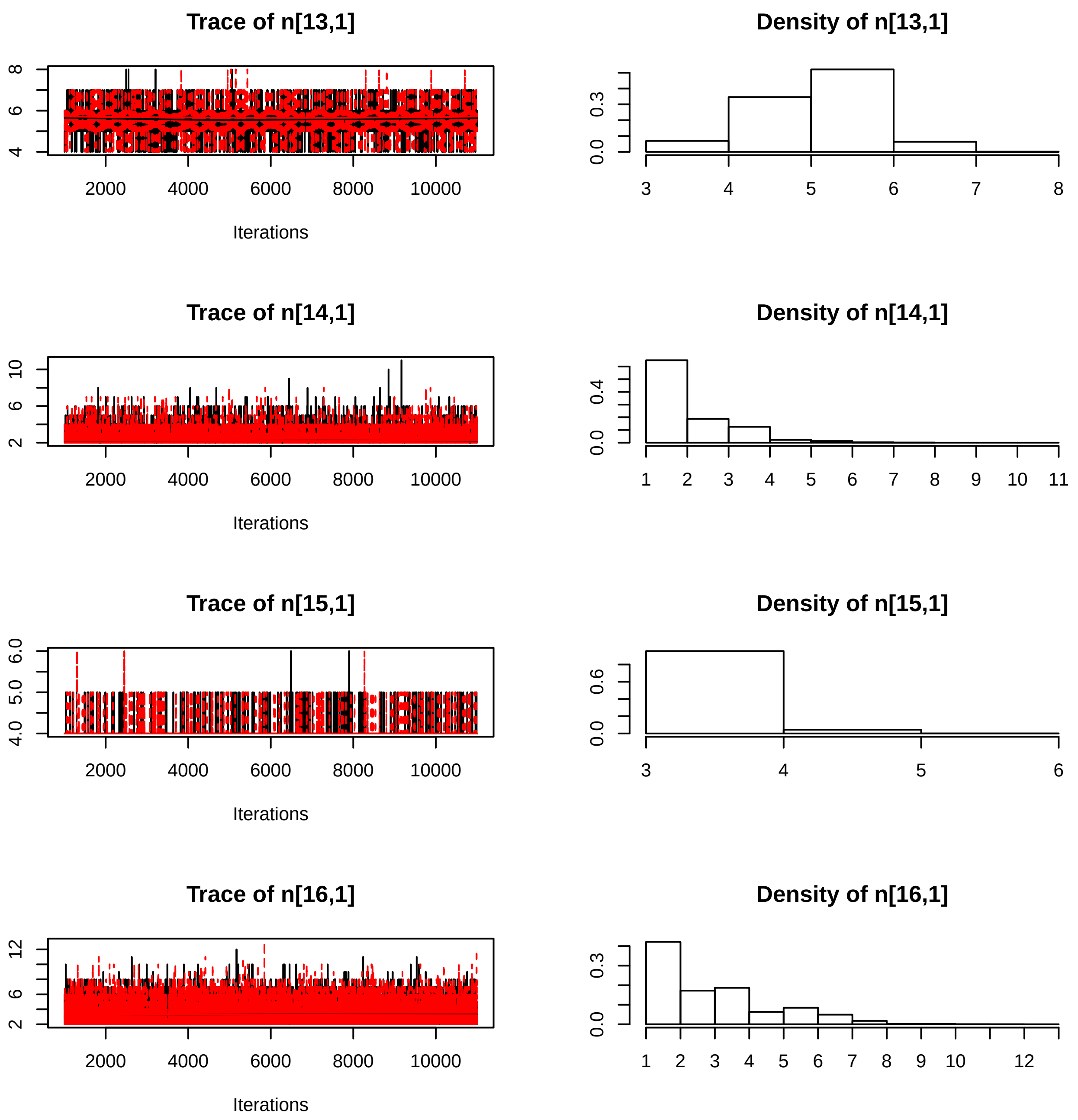
Trace of n[13,1]
Density of n[13,1]
Iterations
Trace of n[14,1]
Density of n[14,1]
Iterations
Trace of n[15,1]
Density of n[15,1]
Iterations
Trace of n[16,1]
Density of n[16,1]
Iterations

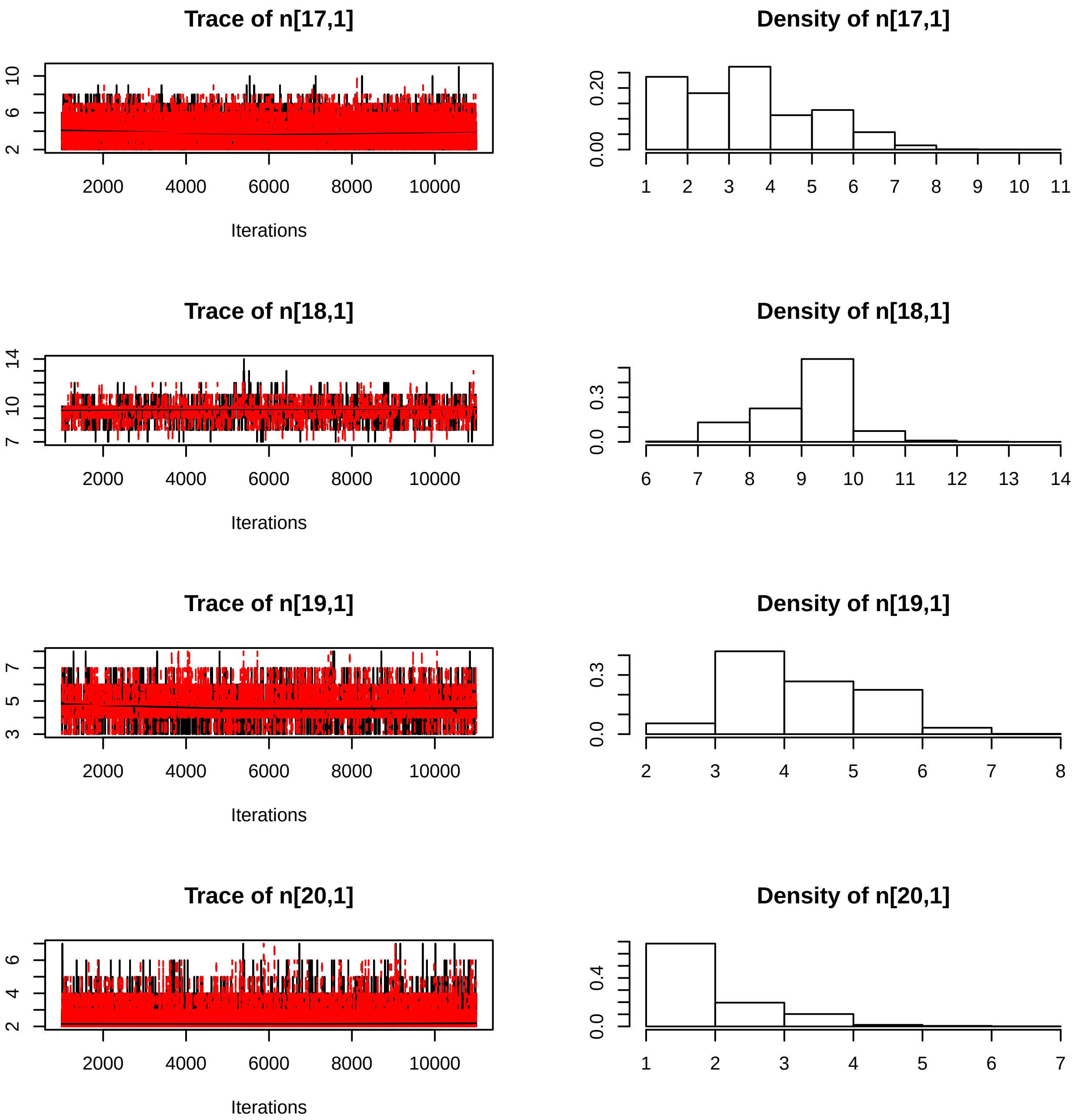
Trace of n[17,1]
Density of n[17,1]
Iterations
Trace of n[18,1]
Density of n[18,1]
Iterations
Trace of n[19,1]
Density of n[19,1]
Iterations
Trace of n[20,1]
Density of n[20,1]
Iterations

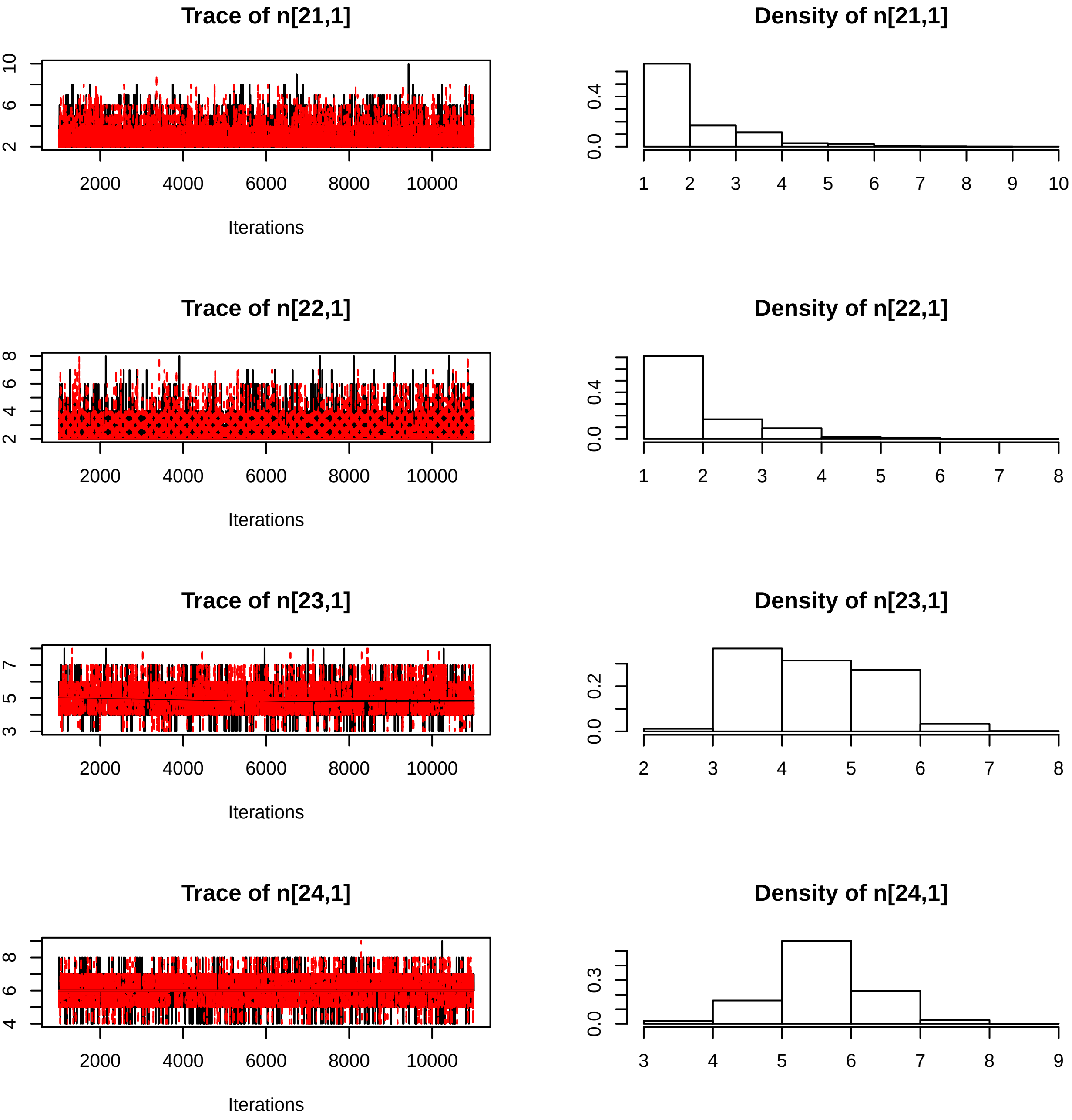

Trace of n[21,1]
Iterations
Density of n[21,1]
Trace of n[22,1]
Iterations
Density of n[22,1]
Trace of n[23,1]
Iterations
Density of n[23,1]
Trace of n[24,1]
Iterations
Density of n[24,1]

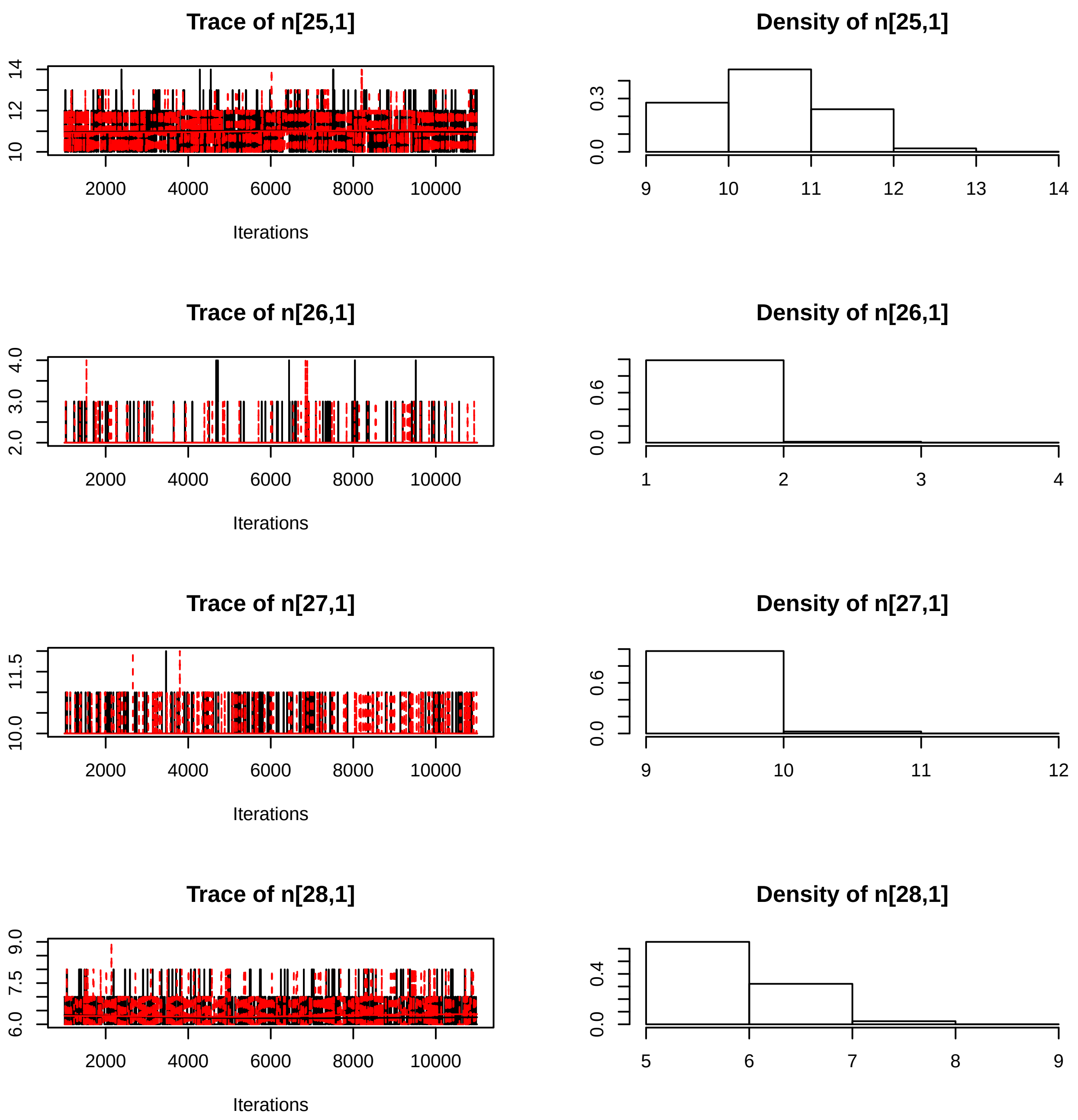
Trace of n[25,1]
Density of n[25,1]
10
12
14
2000
4000
6000
8000
10000
Iterations
0.0
0.3
9
10
11
12
13
14
Trace of n[26,1]
Density of n[26,1]
2.0
3.0
4.0
2000
4000
6000
8000
10000
Iterations
0.0
0.6
1
2
3
4
Trace of n[27,1]
Density of n[27,1]
10.0
11.5
2000
4000
6000
8000
10000
Iterations
0.0
0.6
9
10
11
12
Trace of n[28,1]
Density of n[28,1]
6.0
7.5
9.0
2000
4000
6000
8000
10000
Iterations
0.0
0.4
5
6
7
8
9

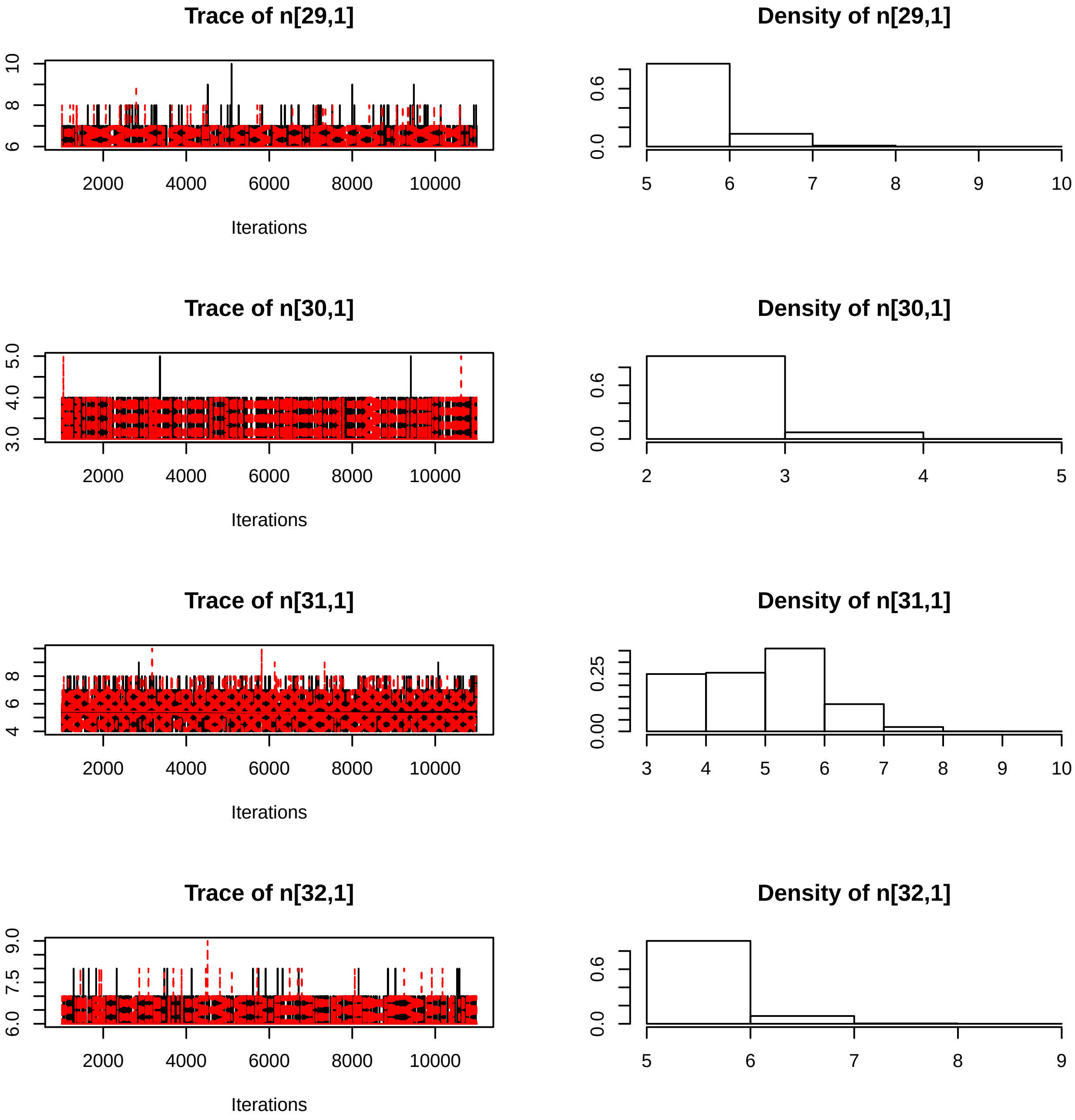

Trace of n[29,1]
Iterations
Density of n[29,1]
Trace of n[30,1]
Iterations
Density of n[30,1]
Trace of n[31,1]
Iterations
Density of n[31,1]
Trace of n[32,1]
Iterations
Density of n[32,1]

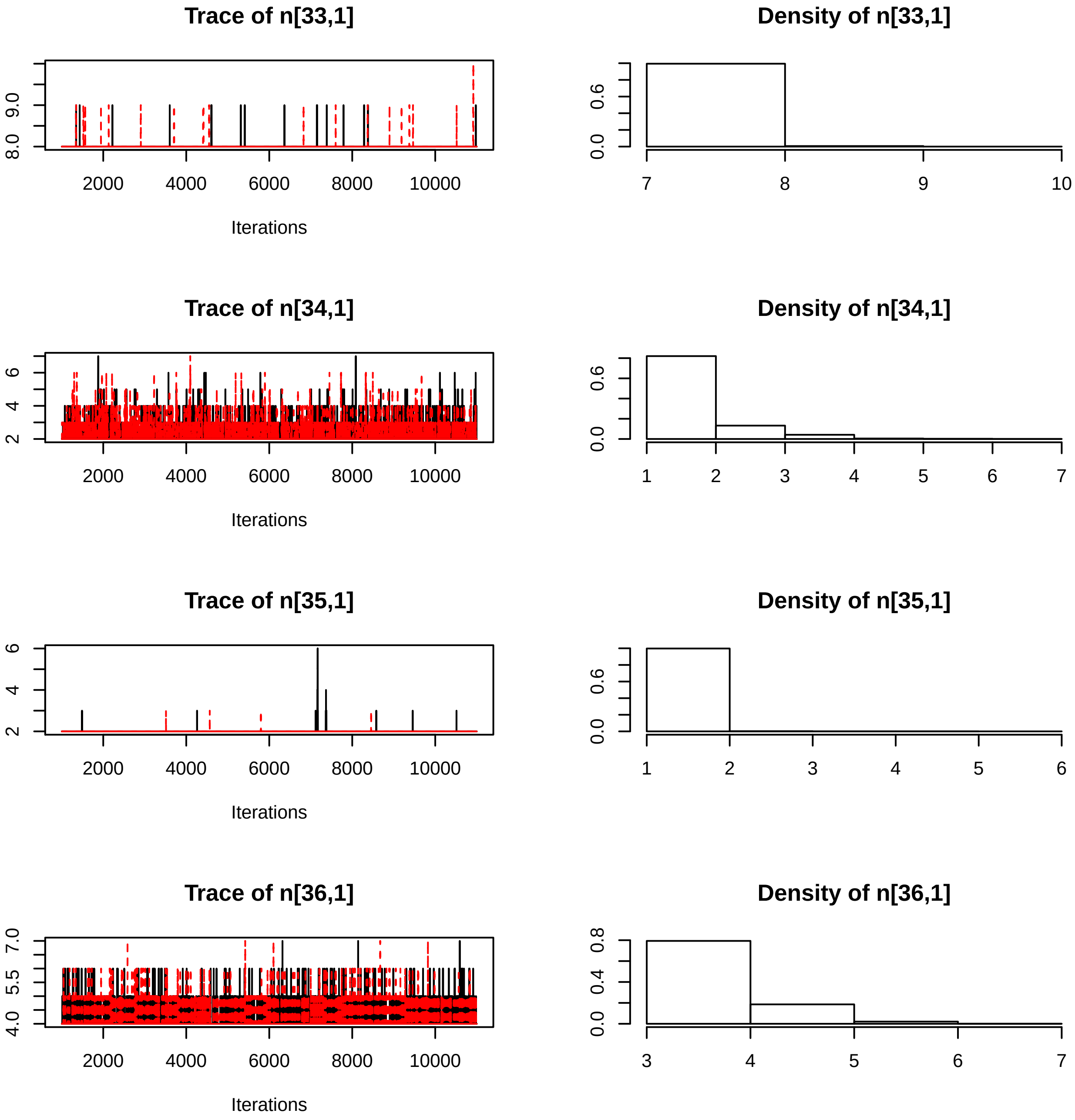
Trace of n[33,1]
Density of n[33,1]
Trace of n[34,1]
Density of n[34,1]
Trace of n[35,1]
Density of n[35,1]
Trace of n[36,1]
Density of n[36,1]
Iterations

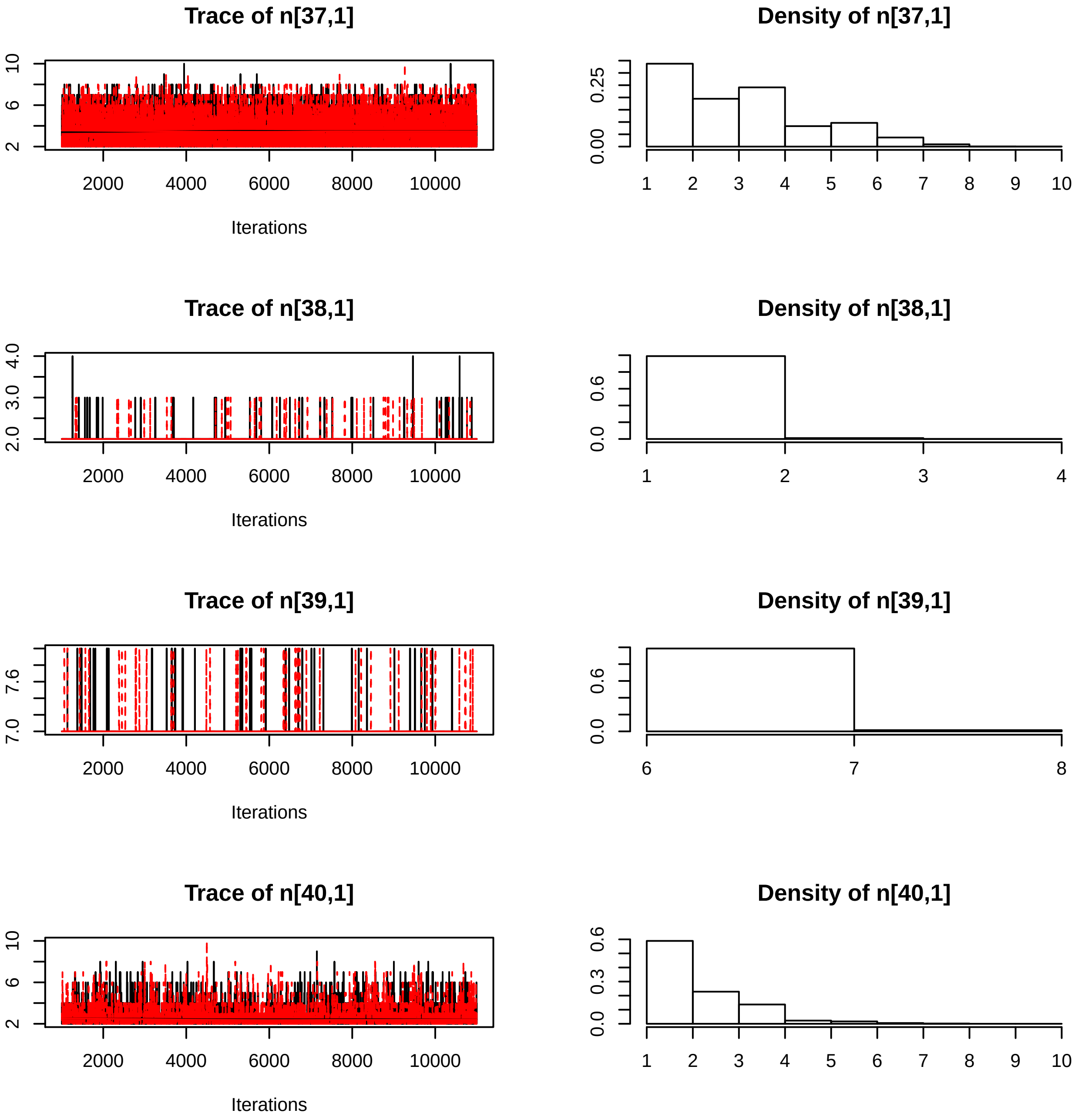
Trace of n[37,1]
Density of n[37,1]
Iterations
Trace of n[38,1]
Density of n[38,1]
Iterations
Trace of n[39,1]
Density of n[39,1]
Iterations
Trace of n[40,1]
Density of n[40,1]
Iterations

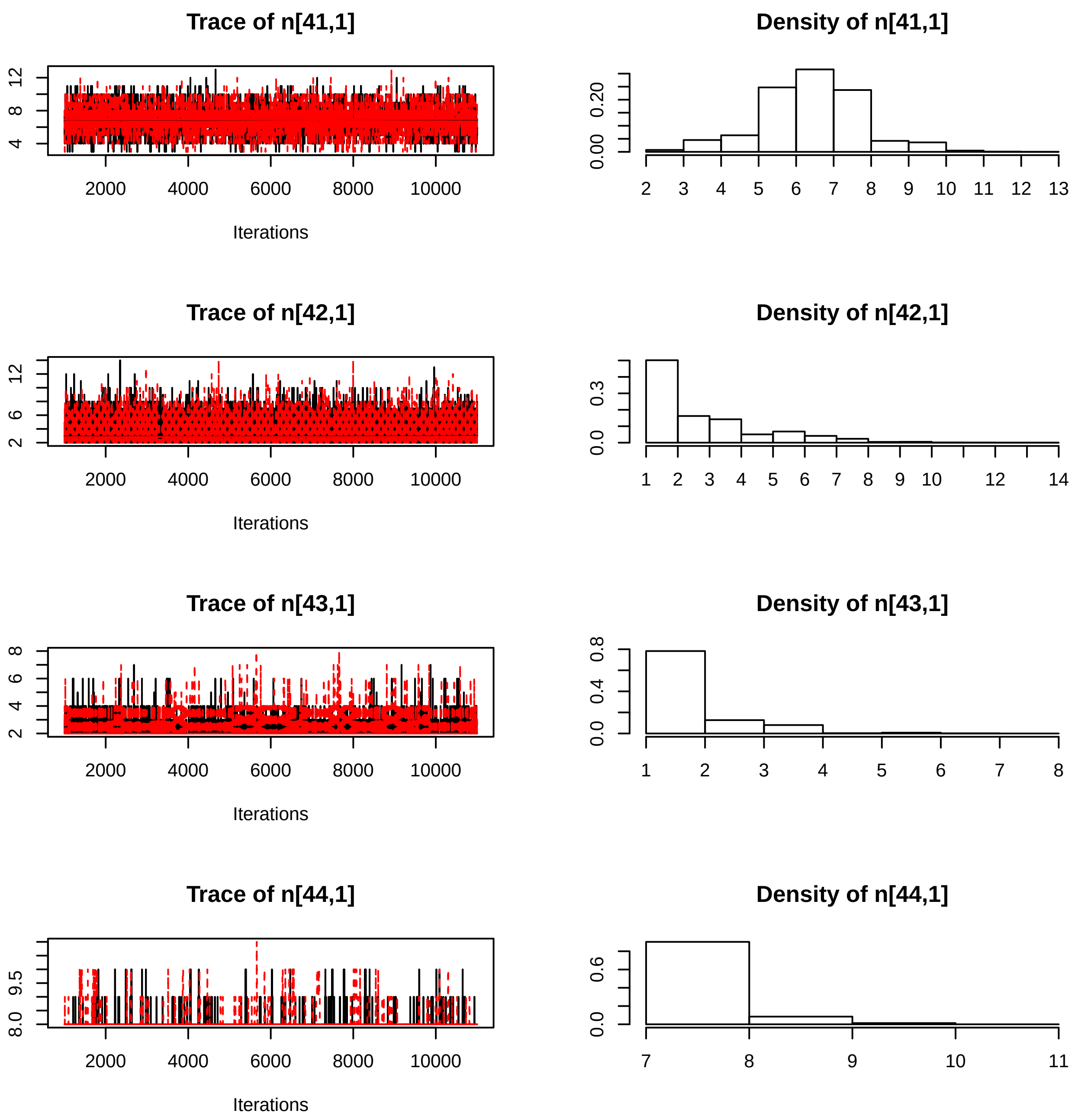
Trace of n[41,1]
Density of n[41,1]
Trace of n[42,1]
Density of n[42,1]
Trace of n[43,1]
Density of n[43,1]
Trace of n[44,1]
Density of n[44,1]
Iterations
Iterations
Iterations
Iterations

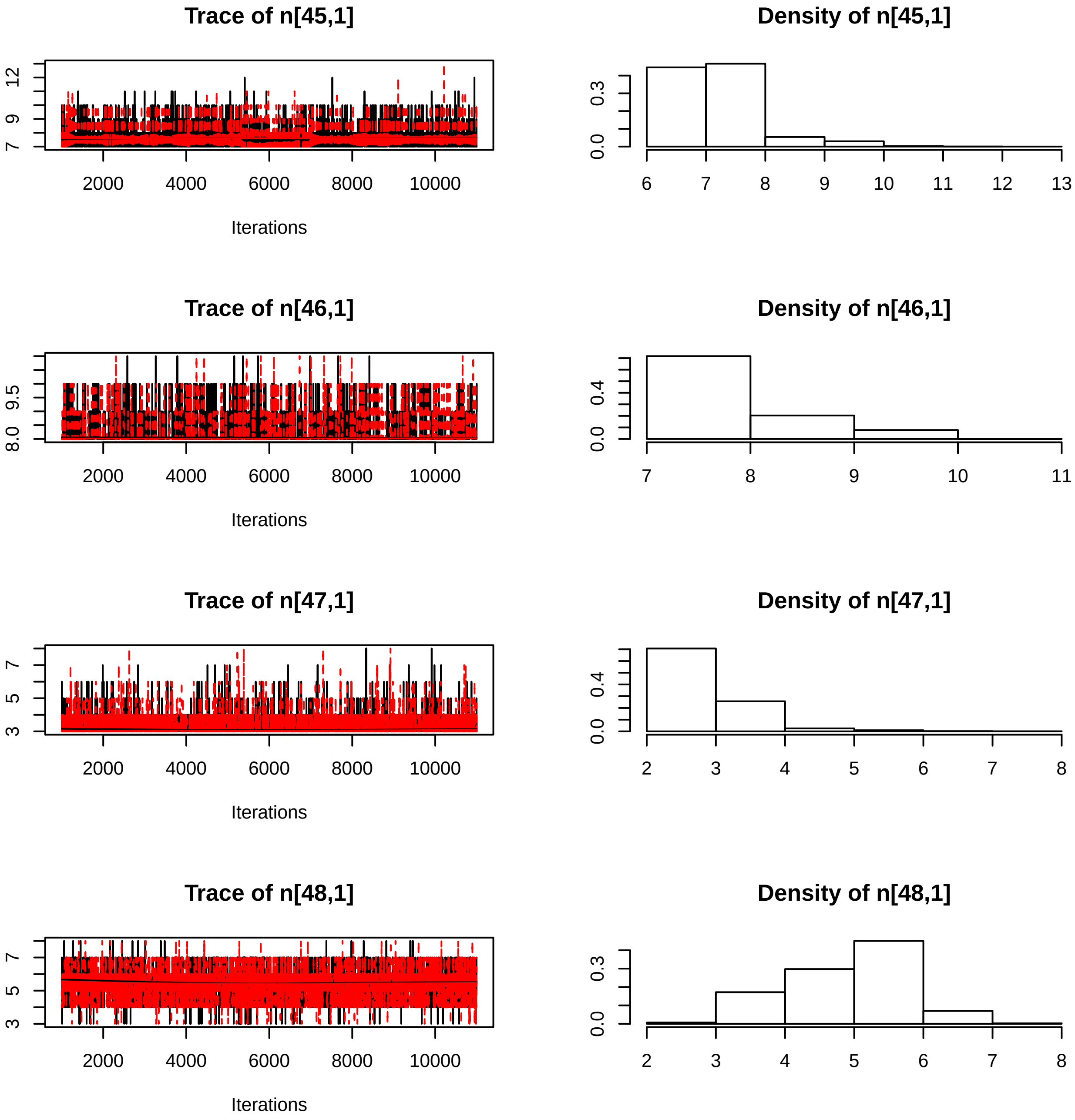

Trace of n[45,1]
12
9
7
2000
4000
6000
8000
10000
Iterations
Density of n[45,1]
0.3
0.0
6
7
8
9
10
11
12
13
Trace of n[46,1]
9.5
8.0
2000
4000
6000
8000
10000
Iterations
Density of n[46,1]
0.4
0.0
7
8
9
10
11
Trace of n[47,1]
7
5
3
2000
4000
6000
8000
10000
Iterations
Density of n[47,1]
0.4
0.0
2
3
4
5
6
7
8
Trace of n[48,1]
7
5
3
2000
4000
6000
8000
10000
Iterations
Density of n[48,1]
0.3
0.0
2
3
4
5
6
7
8

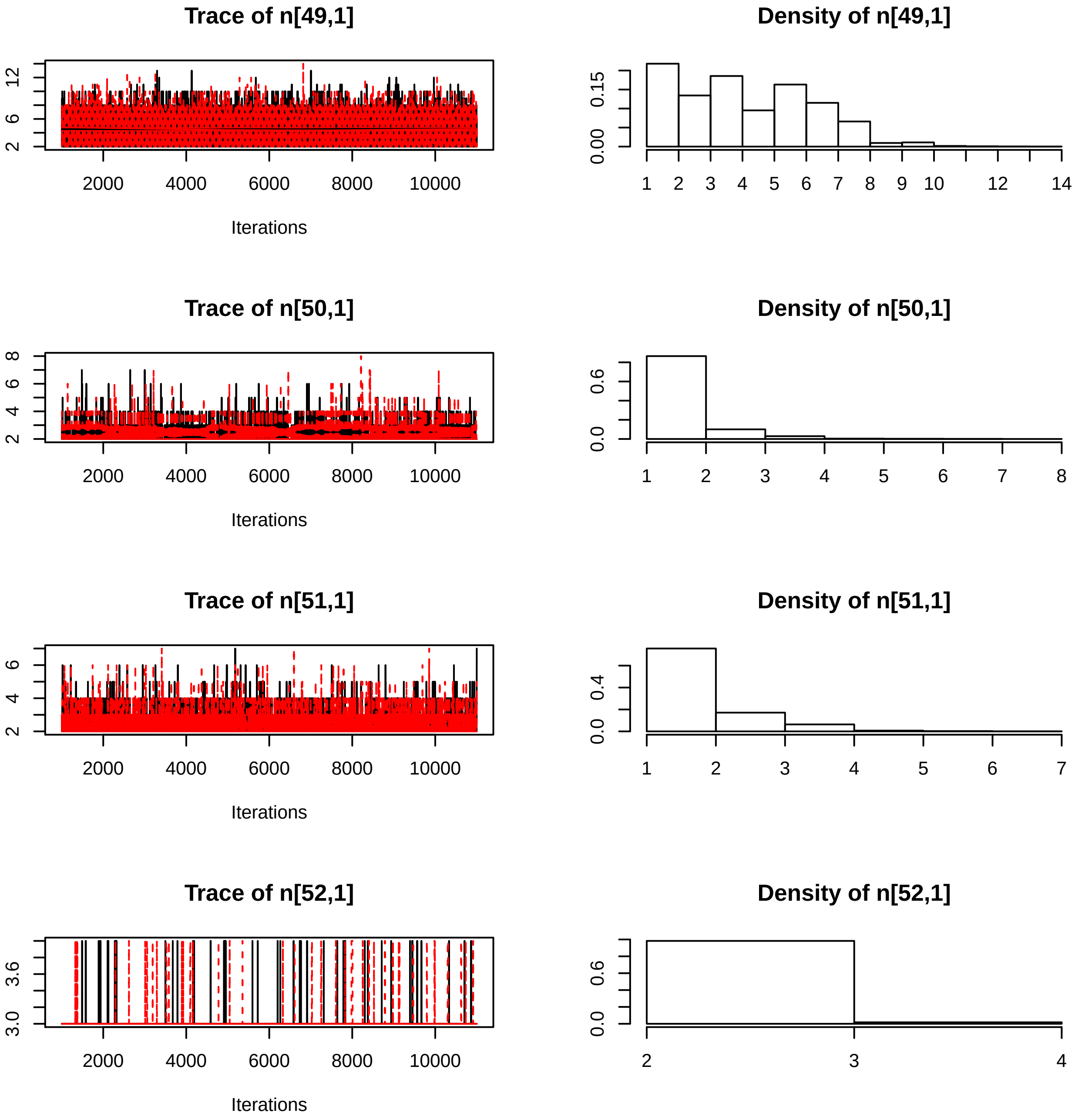

Trace of n[49,1]
Density of n[49,1]
Trace of n[50,1]
Density of n[50,1]
Trace of n[51,1]
Density of n[51,1]
Trace of n[52,1]
Density of n[52,1]
Iterations
2000 4000 6000 8000 10000

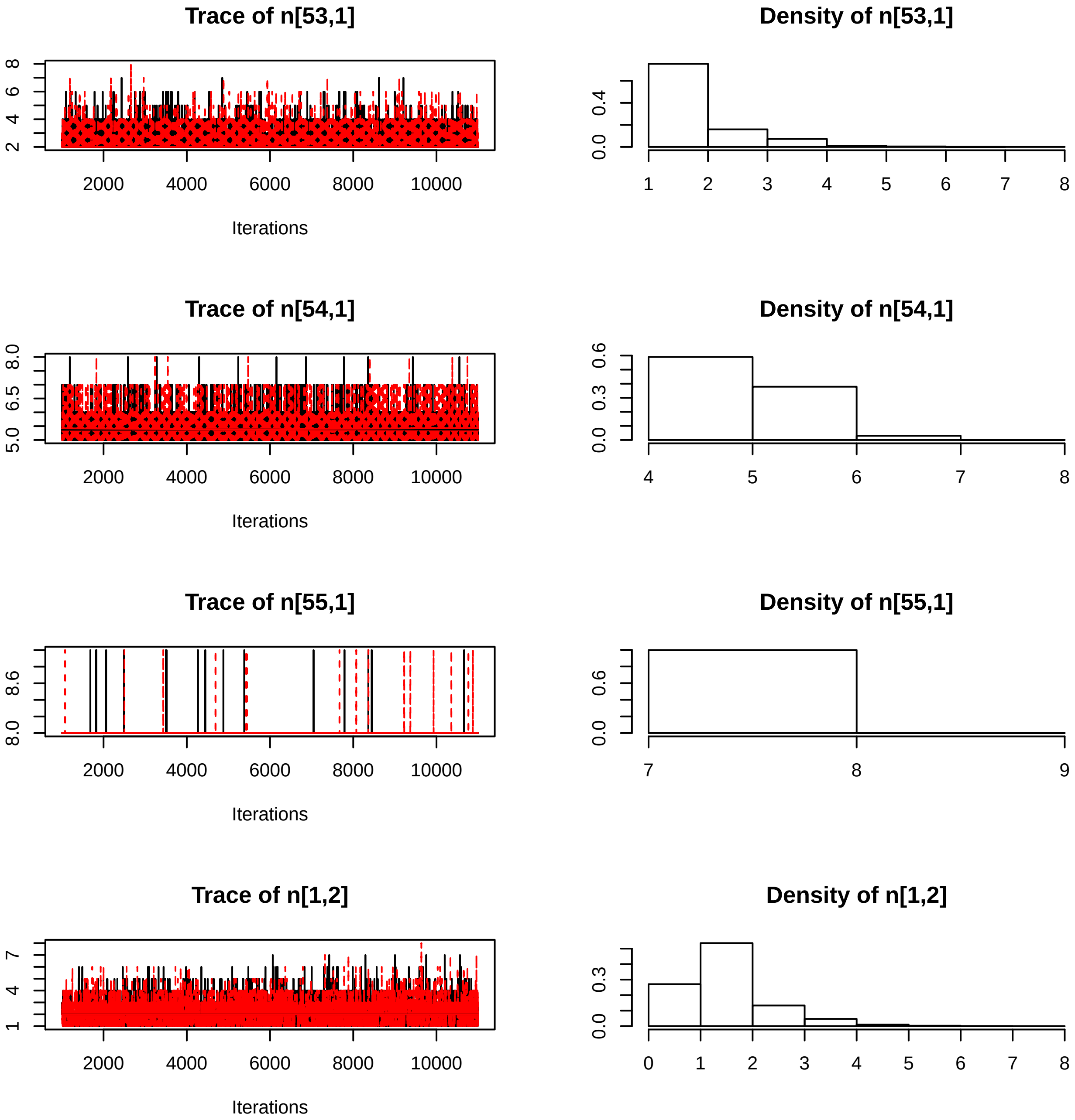
Trace of n[53,1]
Iterations
Density of n[53,1]
Trace of n[54,1]
Iterations
Density of n[54,1]
Trace of n[55,1]
Iterations
Density of n[55,1]
Trace of n[1,2]
Iterations
Density of n[1,2]
2000
4000
6000
8000
10000

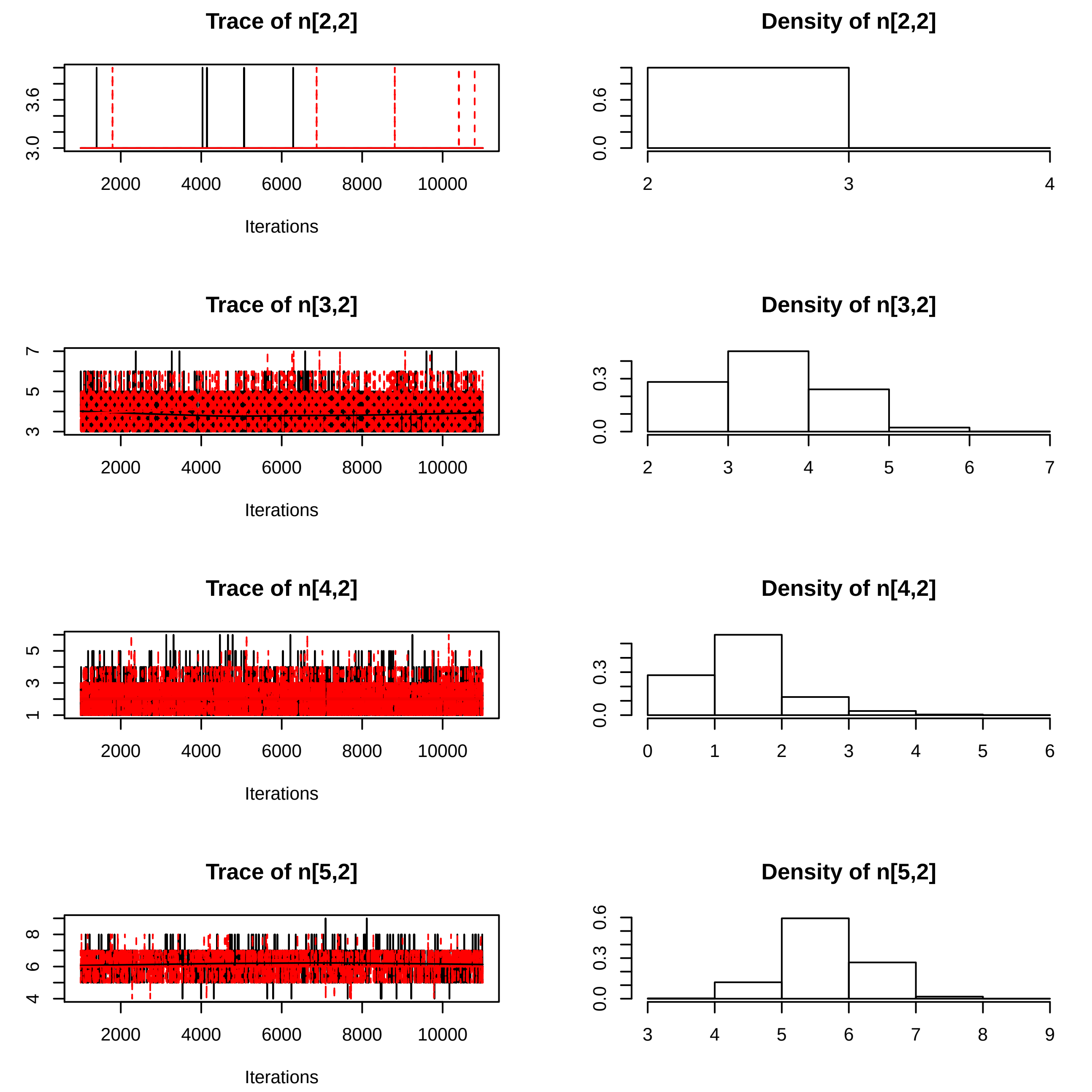

Trace of n[2,2]
3.0
3.6
2000
4000
6000
8000
10000
Iterations
Density of n[2,2]
0.0
0.6
2
3
4
Trace of n[3,2]
3
5
7
2000
4000
6000
8000
10000
Iterations
Density of n[3,2]
0.0
0.3
2
3
4
5
6
7
Trace of n[4,2]
1
3
5
2000
4000
6000
8000
10000
Iterations
Density of n[4,2]
0.0
0.3
0
1
2
3
4
5
6
Trace of n[5,2]
4
6
8
2000
4000
6000
8000
10000
Iterations
Density of n[5,2]
0.0
0.3
0.6
3
4
5
6
7
8
9

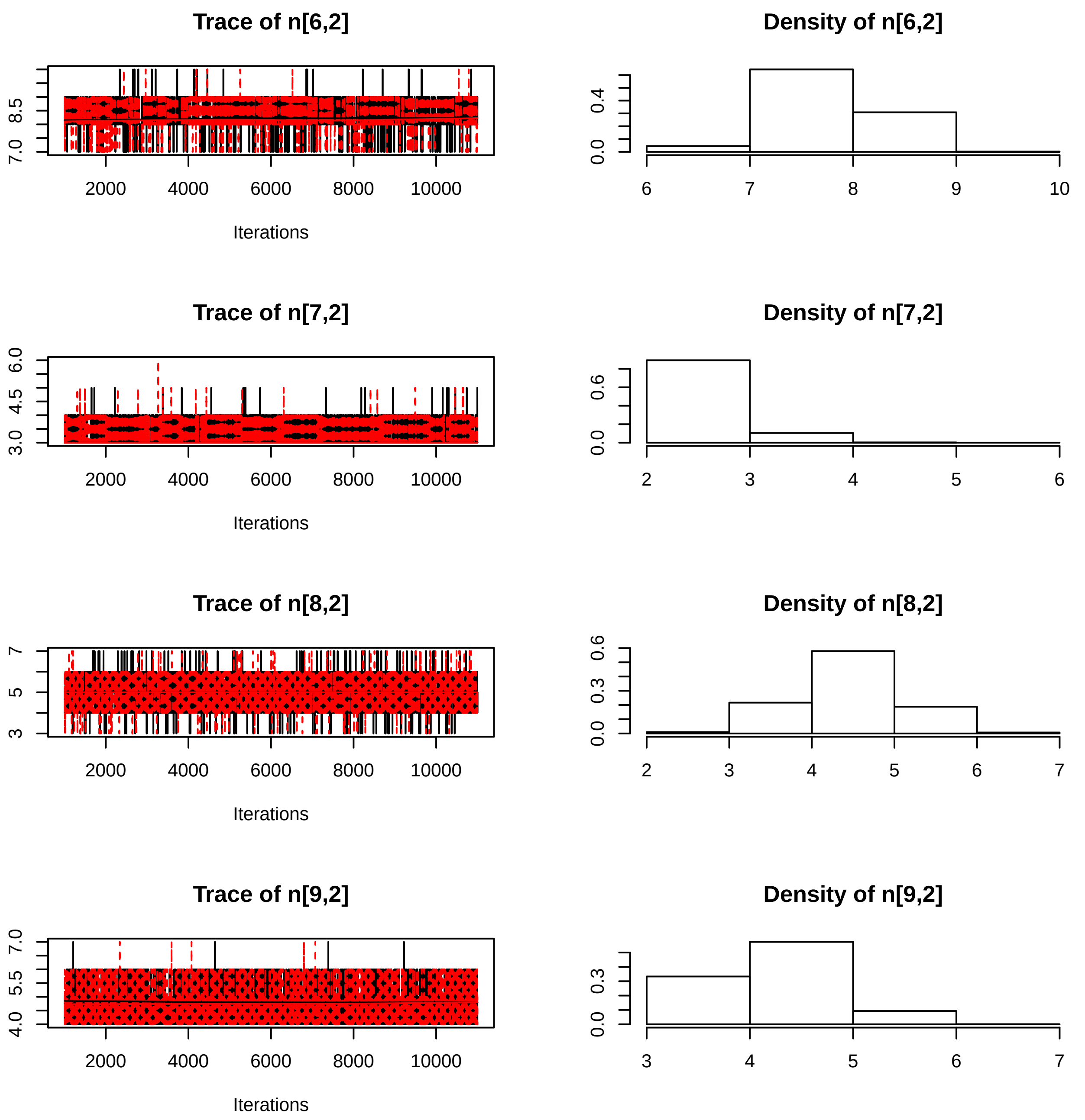

Trace of n[6,2]
8.5
7.0
2000
4000
6000
8000
10000
Iterations
Density of n[6,2]
0.4
0.0
6
7
8
9
10
Trace of n[7,2]
6.0
4.5
3.0
2000
4000
6000
8000
10000
Iterations
Density of n[7,2]
0.6
0.0
2
3
4
5
6
Trace of n[8,2]
7
5
3
2000
4000
6000
8000
10000
Iterations
Density of n[8,2]
0.6
0.3
0.0
2
3
4
5
6
7
Trace of n[9,2]
7.0
5.5
4.0
2000
4000
6000
8000
10000
Iterations
Density of n[9,2]
0.3
0.0
3
4
5
6
7

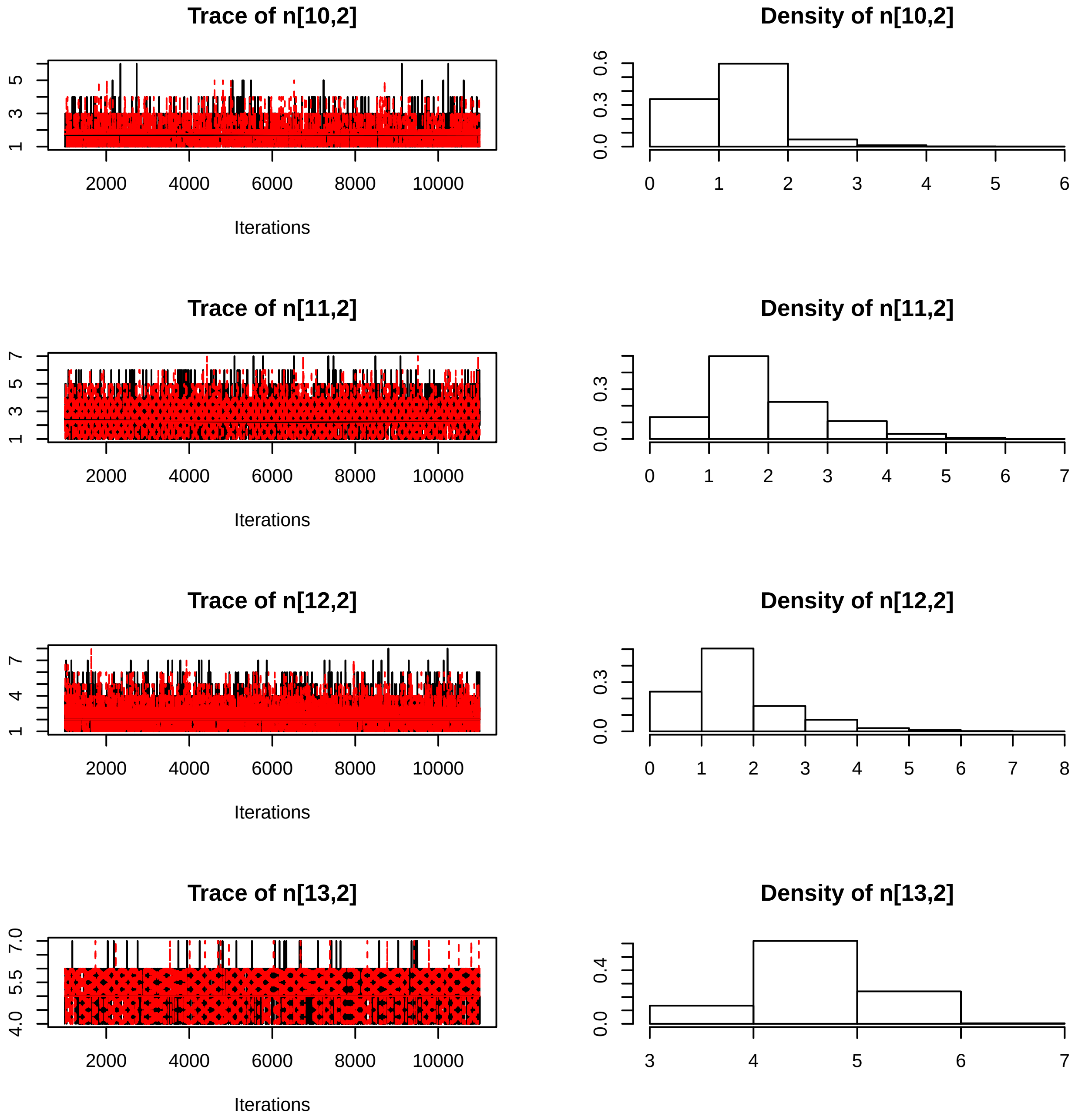
Trace of n[10,2]
Iterations
Density of n[10,2]
Trace of n[11,2]
Iterations
Density of n[11,2]
Trace of n[12,2]
Iterations
Density of n[12,2]
Trace of n[13,2]
Iterations
Density of n[13,2]

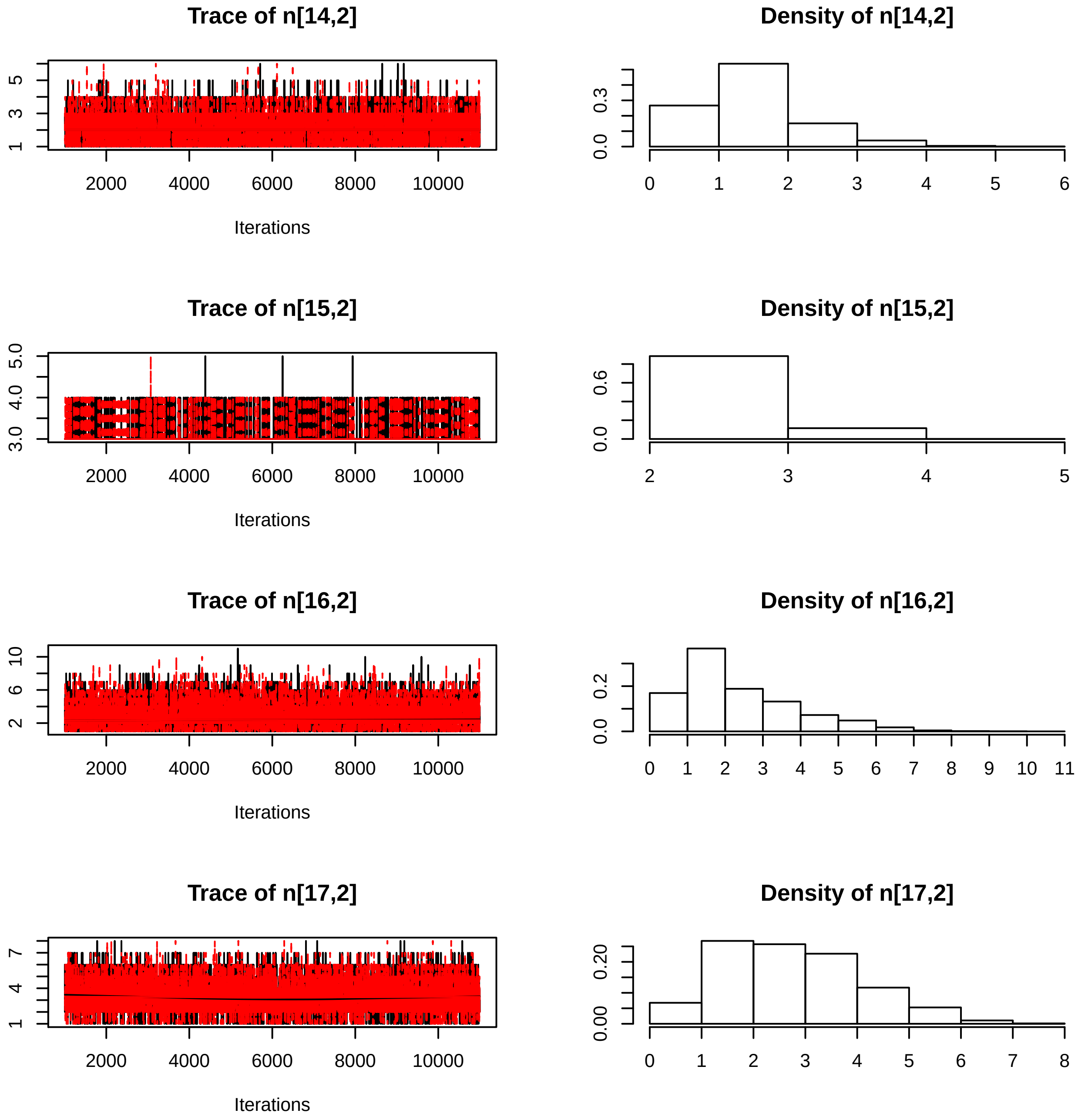
Trace of n[14,2]
Iterations
Density of n[14,2]
Trace of n[15,2]
Iterations
Density of n[15,2]
Trace of n[16,2]
Iterations
Density of n[16,2]
Trace of n[17,2]
Iterations
Density of n[17,2]

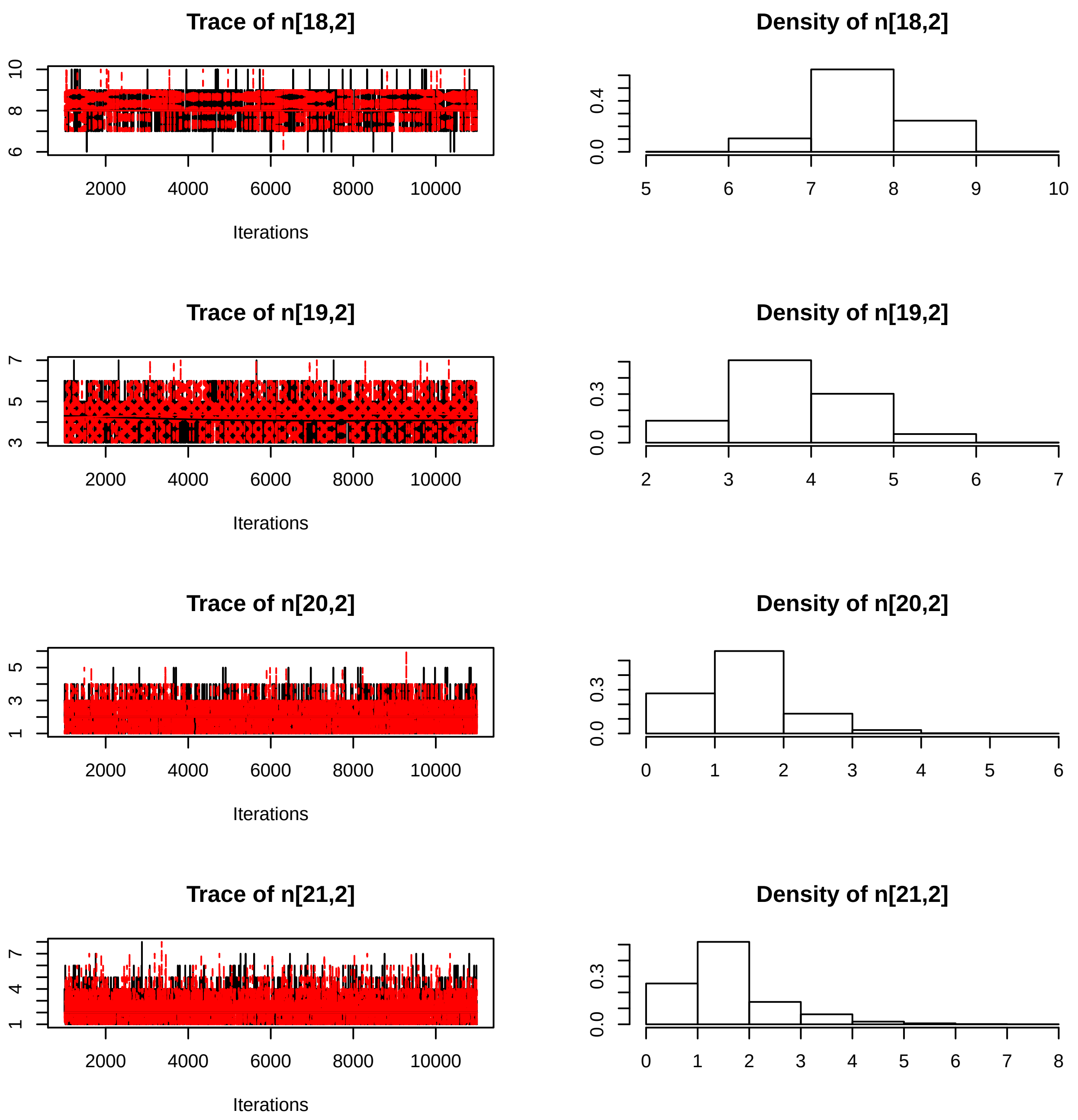
Trace of n[18,2]
Density of n[18,2]
Trace of n[19,2]
Density of n[19,2]
Trace of n[20,2]
Density of n[20,2]
Trace of n[21,2]
Density of n[21,2]
Iterations
2000
4000
6000
8000
10000

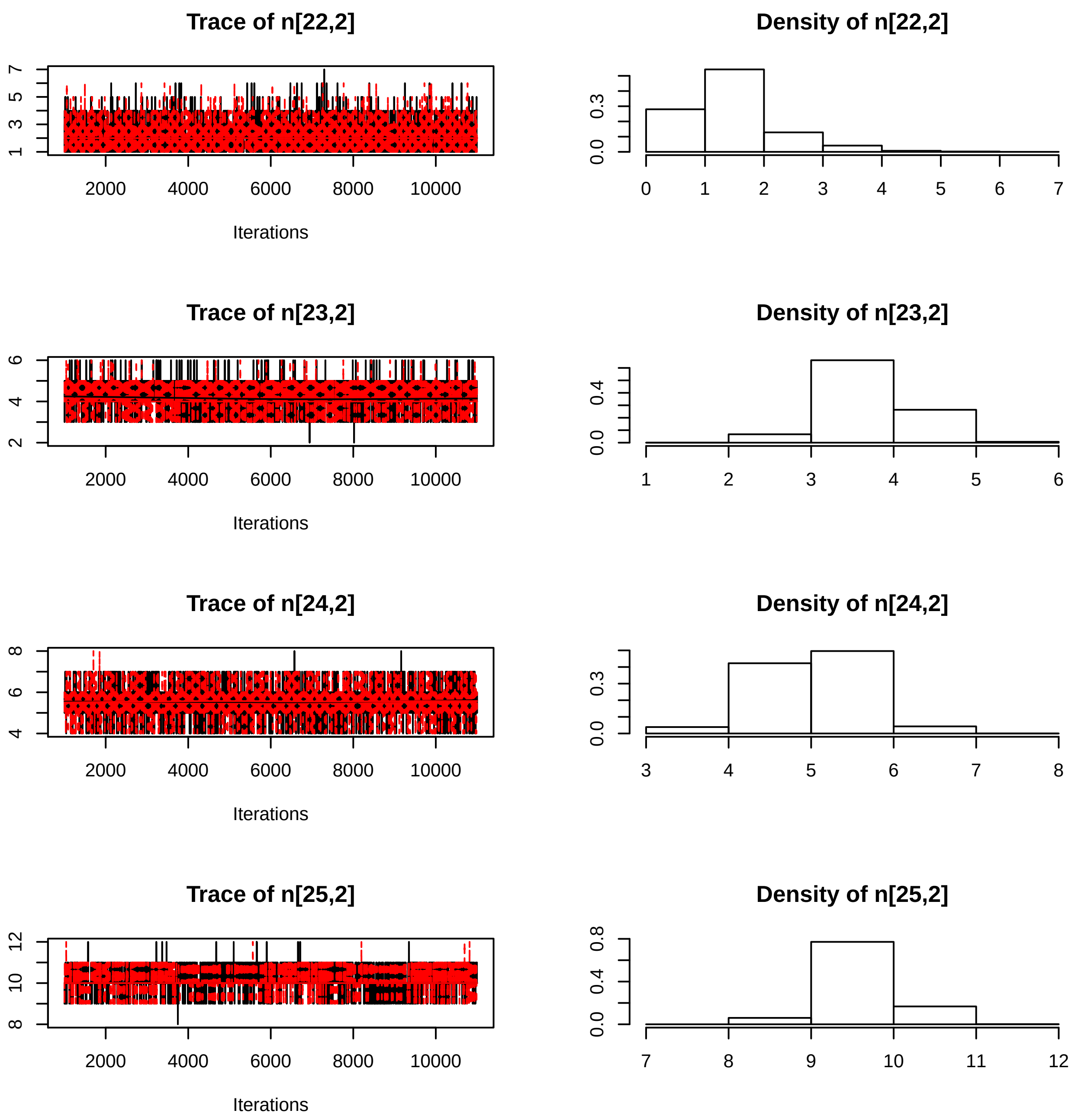
Trace of n[22,2]
Iterations
Density of n[22,2]
Trace of n[23,2]
Iterations
Density of n[23,2]
Trace of n[24,2]
Iterations
Density of n[24,2]
Trace of n[25,2]
Iterations
Density of n[25,2]
2000
4000
6000
8000
10000

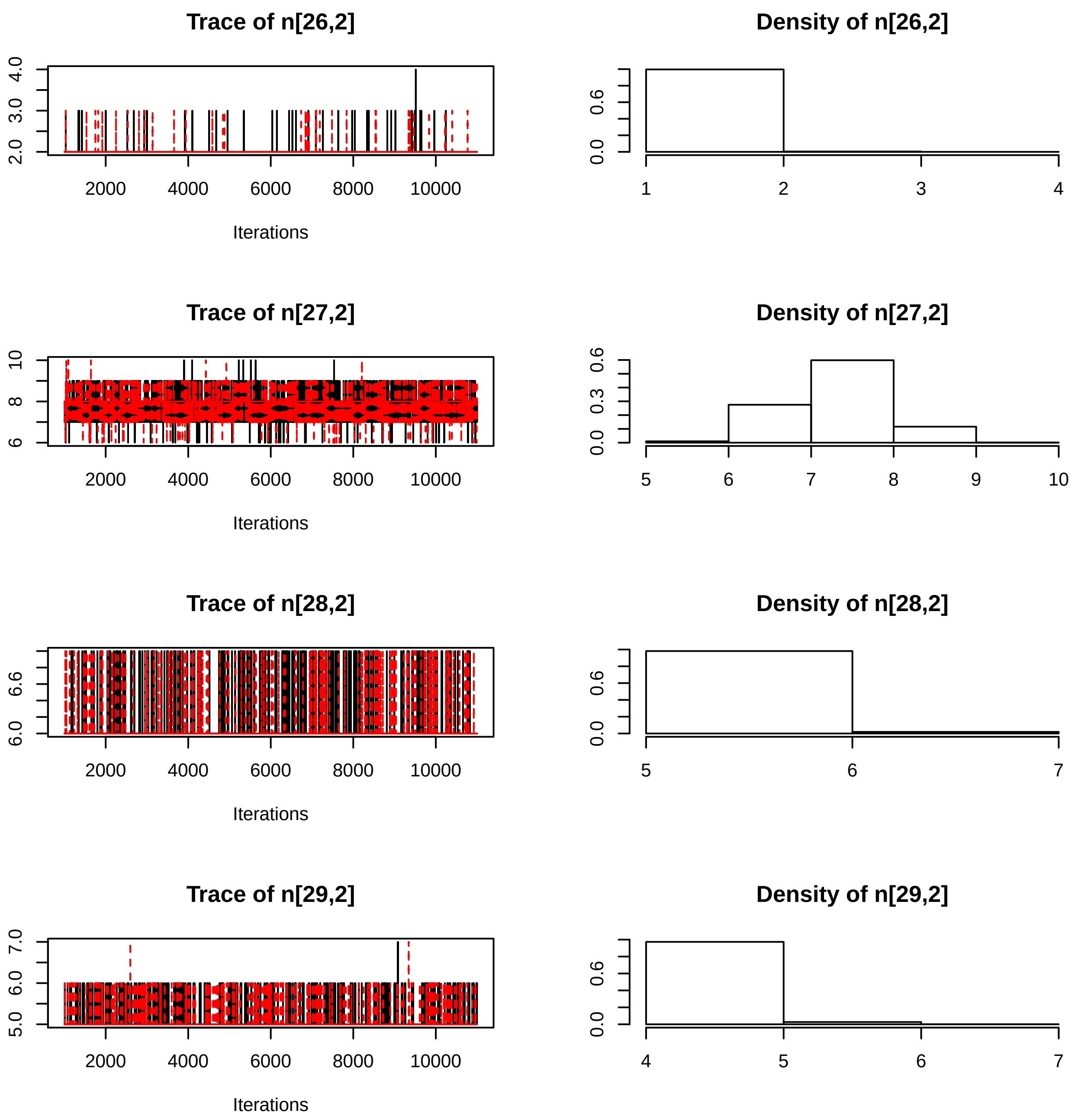

Trace of n[26,2]
2.0
3.0
4.0
2000
4000
6000
8000
10000
Iterations
Density of n[26,2]
0.0
0.6
1
2
3
4
Trace of n[27,2]
6
8
10
2000
4000
6000
8000
10000
Iterations
Density of n[27,2]
0.0
0.3
0.6
5
6
7
8
9
10
Trace of n[28,2]
6.0
6.6
2000
4000
6000
8000
10000
Iterations
Density of n[28,2]
0.0
0.6
5
6
7
Trace of n[29,2]
5.0
6.0
7.0
2000
4000
6000
8000
10000
Iterations
Density of n[29,2]
0.0
0.6
4
5
6
7

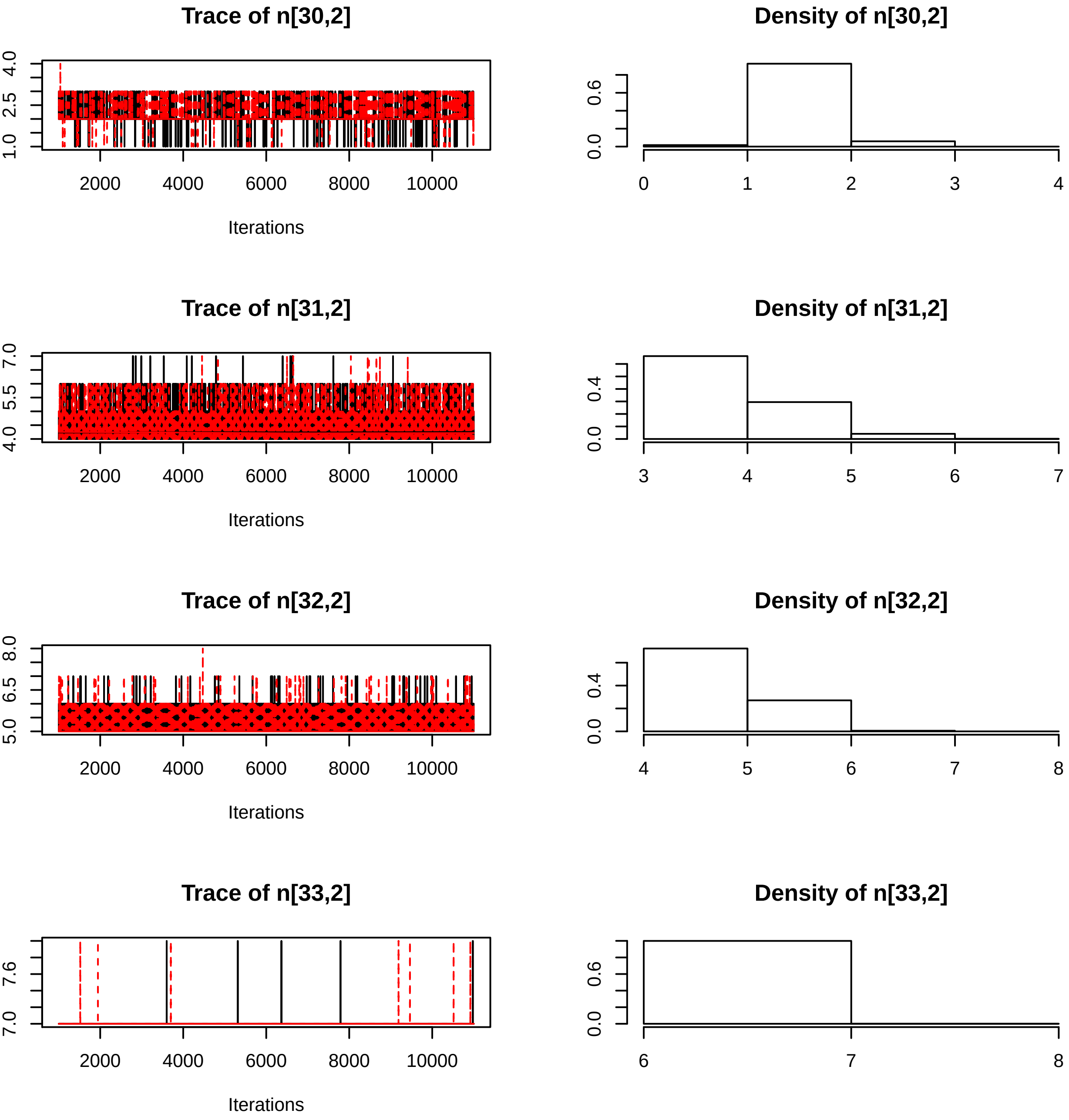
Trace of n[30,2]
Iterations
Density of n[30,2]
Trace of n[31,2]
Iterations
Density of n[31,2]
Trace of n[32,2]
Iterations
Density of n[32,2]
Trace of n[33,2]
Iterations
Density of n[33,2]

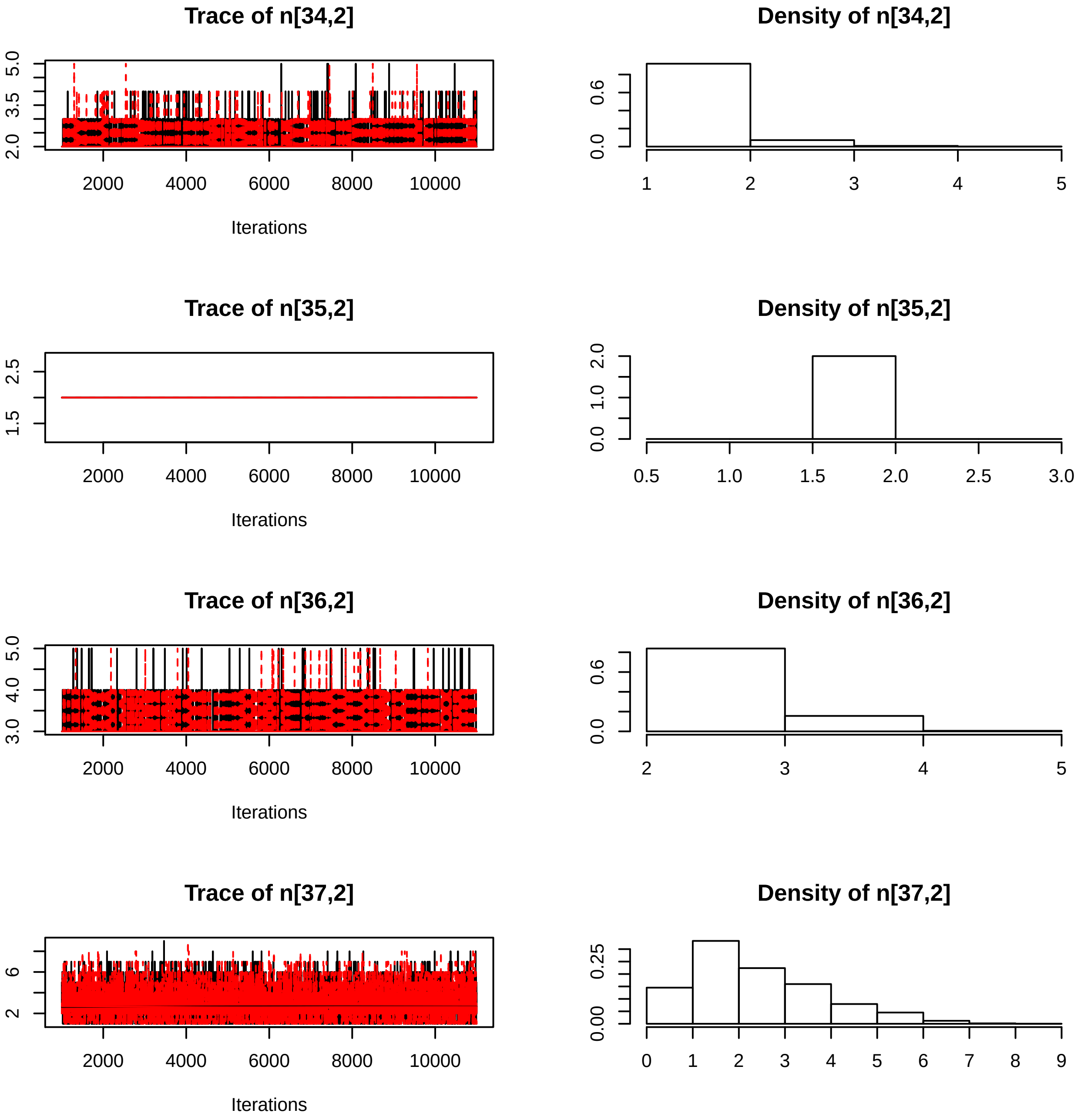

Trace of n[34,2]
5.0
3.5
2.0
2000
4000
6000
8000
10000
Iterations
Density of n[34,2]
0.6
0.0
1
2
3
4
5
Trace of n[35,2]
2.5
1.5
2000
4000
6000
8000
10000
Iterations
Density of n[35,2]
2.0
1.0
0.0
0.5
1.0
1.5
2.0
2.5
3.0
Trace of n[36,2]
5.0
4.0
3.0
2000
4000
6000
8000
10000
Iterations
Density of n[36,2]
0.6
0.0
2
3
4
5
Trace of n[37,2]
6
2
2000
4000
6000
8000
10000
Iterations
Density of n[37,2]
0.25
0.00
0
1
2
3
4
5
6
7
8
9

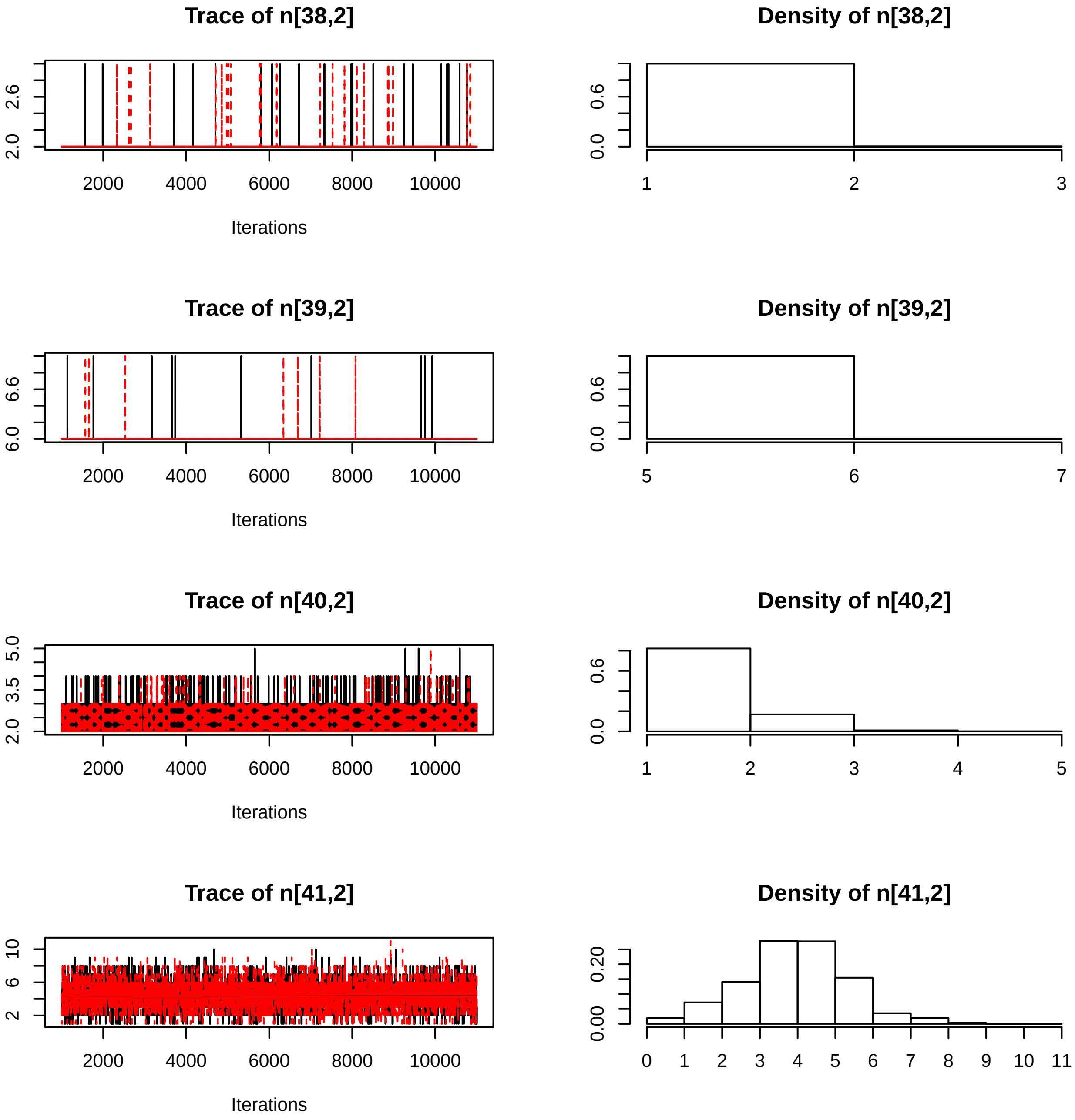

Trace of n[38,2]
Density of n[38,2]
Trace of n[39,2]
Density of n[39,2]
Trace of n[40,2]
Density of n[40,2]
Trace of n[41,2]
Density of n[41,2]
Iterations

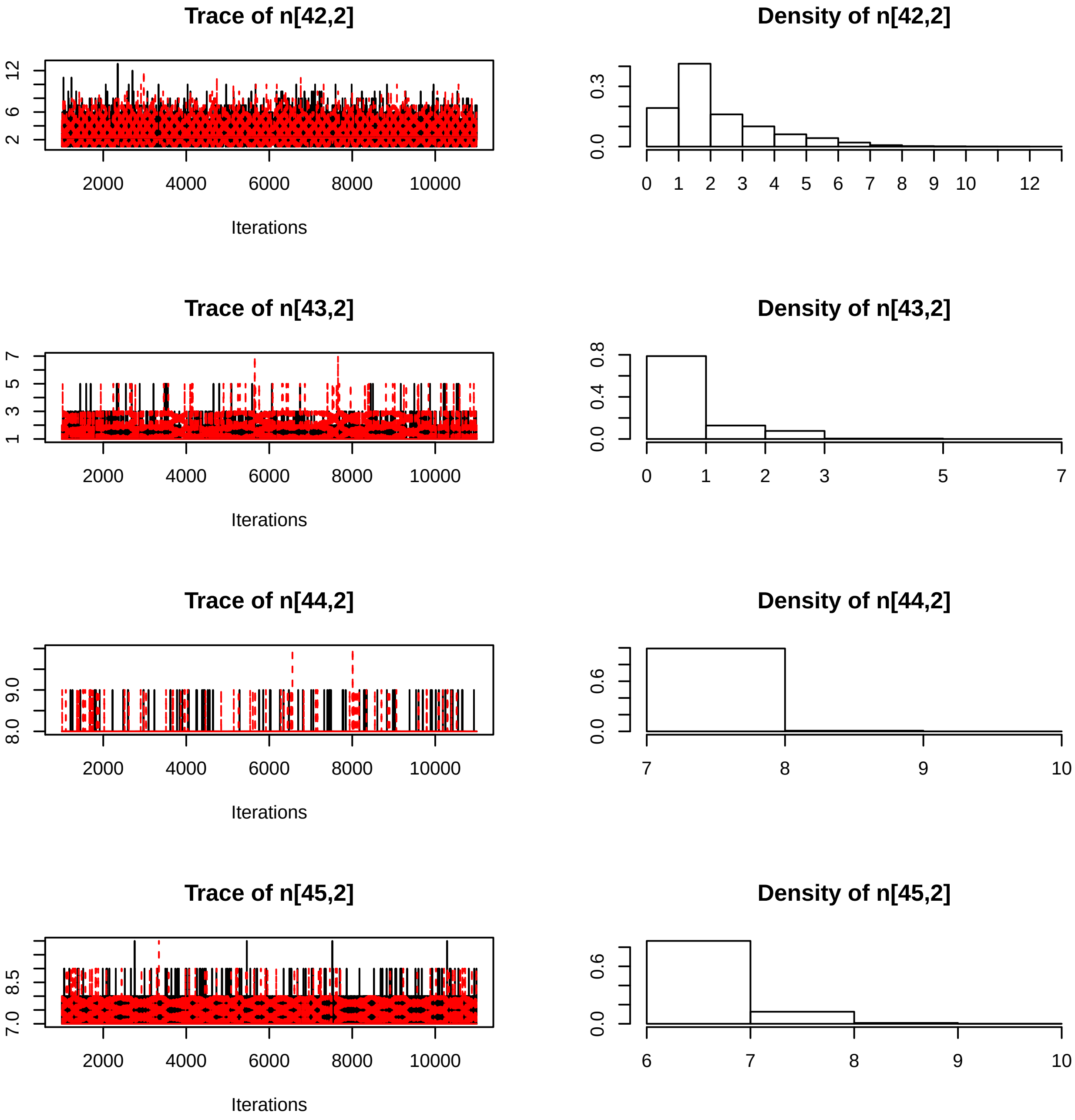
Trace of n[42,2]
Density of n[42,2]
Trace of n[43,2]
Density of n[43,2]
Trace of n[44,2]
Density of n[44,2]
Trace of n[45,2]
Density of n[45,2]
Iterations
2000
4000
6000
8000
10000
2
6
12
0.0
0.3
0 1 2 3 4 5 6 7 8 9 10 12
1
3
5
7
0.0
0.4
0.8
0 1 2 3 5 7
8.0
9.0
0.0
0.6
7 8 9 10
7.0
8.5
6 7 8 9 10

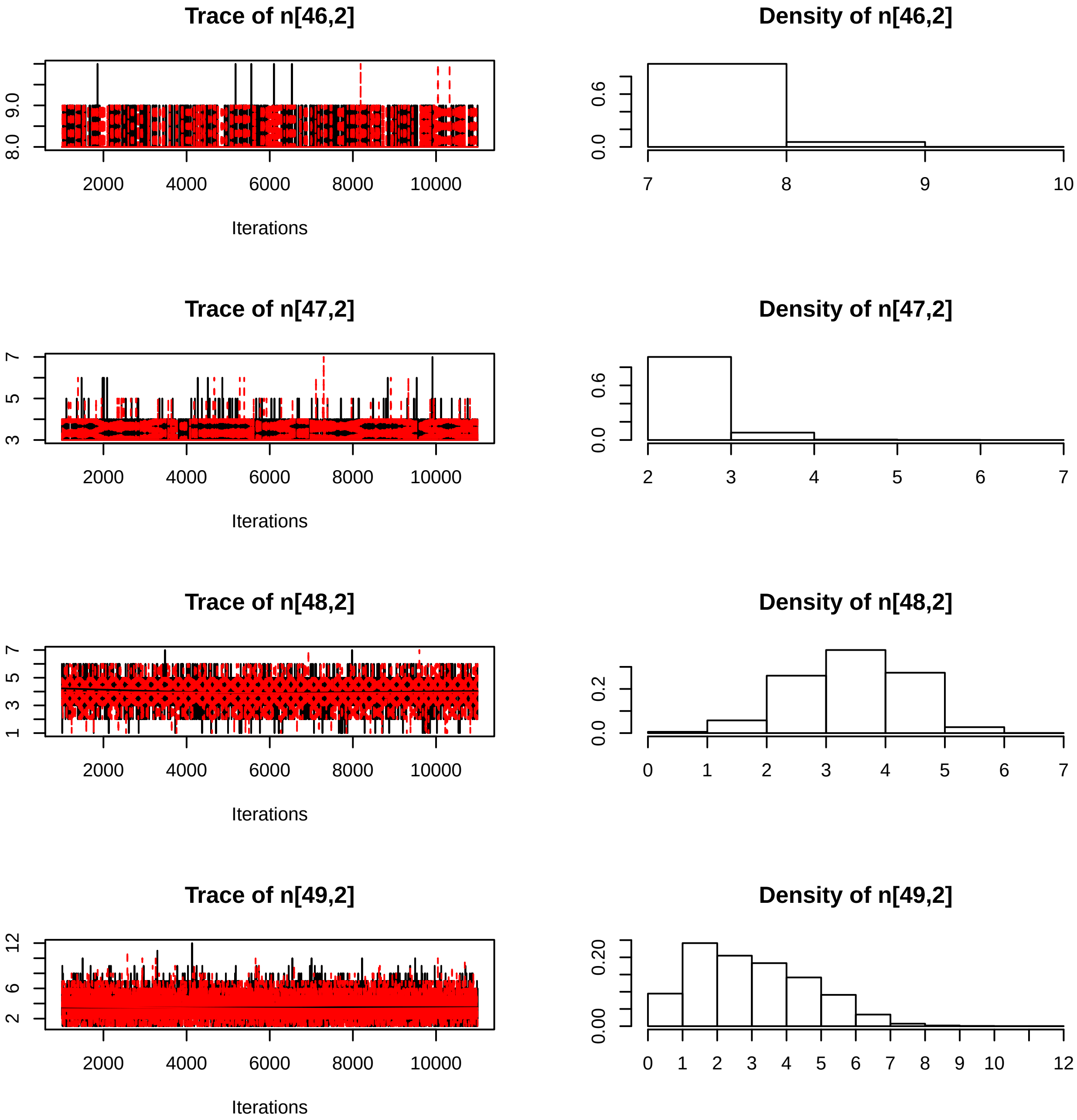

Trace of n[46,2]
Iterations
Density of n[46,2]
Trace of n[47,2]
Iterations
Density of n[47,2]
Trace of n[48,2]
Iterations
Density of n[48,2]
Trace of n[49,2]
Iterations
Density of n[49,2]

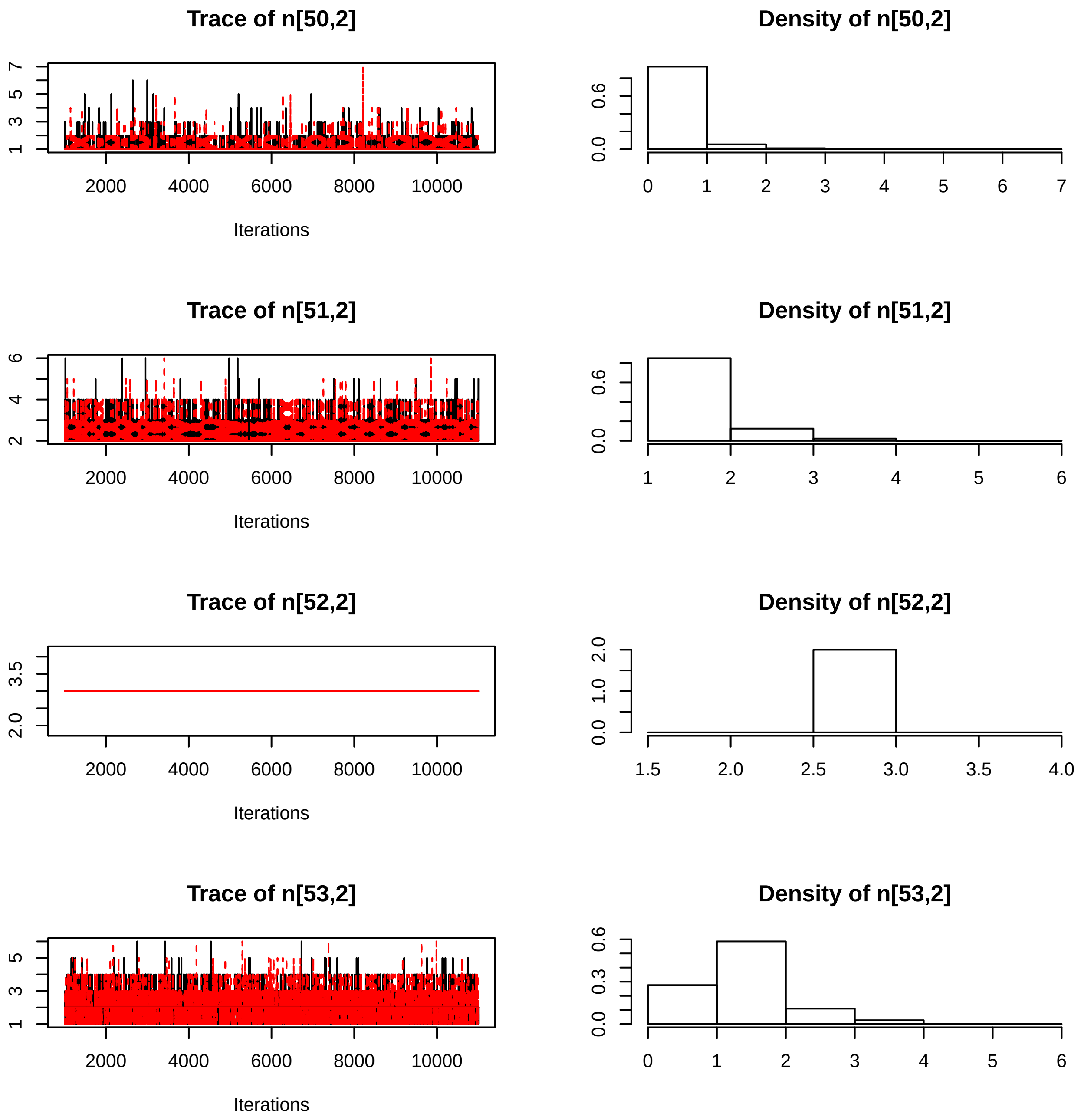
Trace of n[50,2]
Iterations
Density of n[50,2]
Trace of n[51,2]
Iterations
Density of n[51,2]
Trace of n[52,2]
Iterations
Density of n[52,2]
Trace of n[53,2]
Iterations
Density of n[53,2]

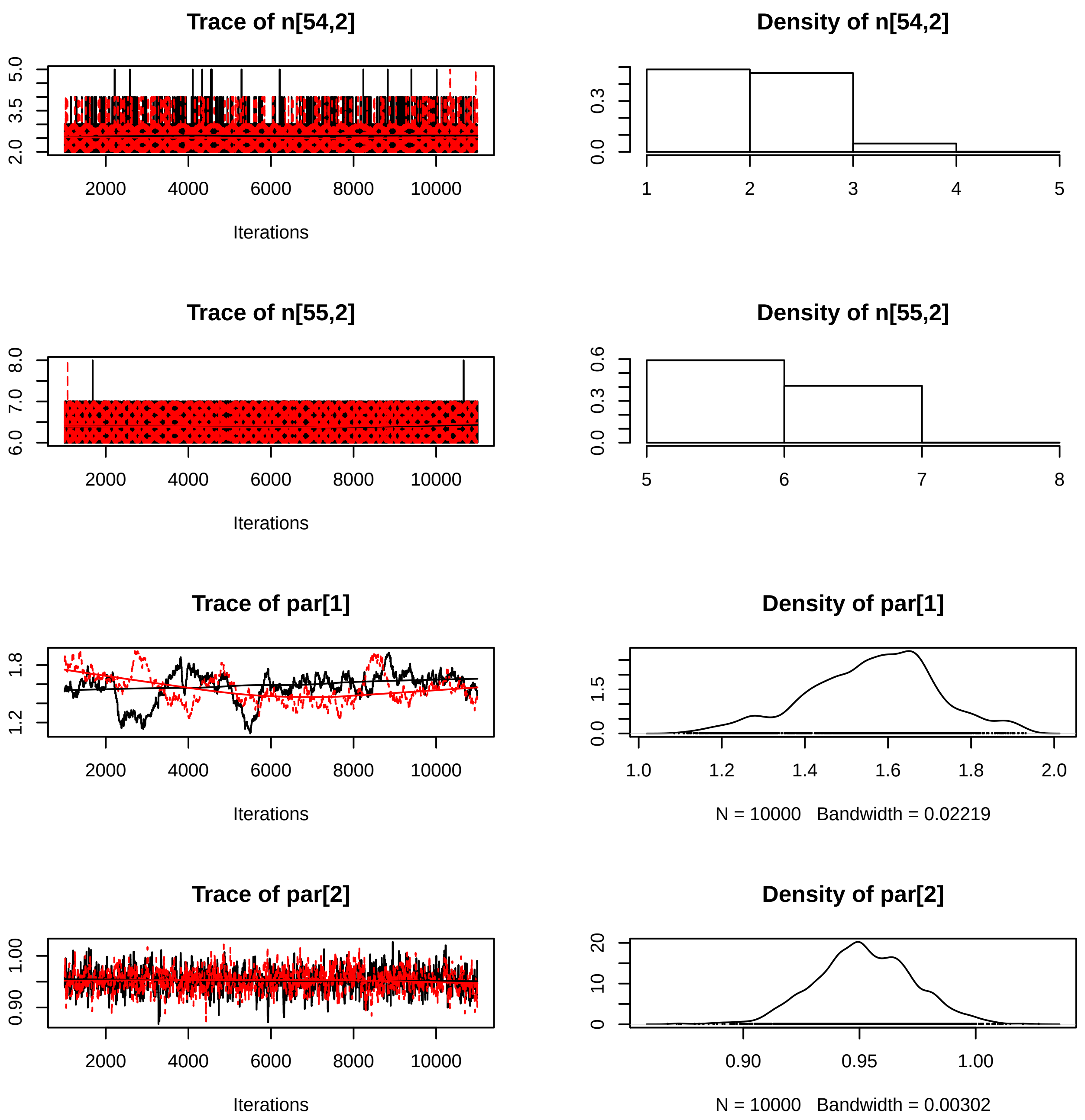

Trace of n[54,2]
Iterations
Density of n[54,2]
Trace of n[55,2]
Iterations
Density of n[55,2]
Trace of par[1]
Iterations
Density of par[1]
N = 10000 Bandwidth = 0.02219
Trace of par[2]
Iterations
Density of par[2]
N = 10000 Bandwidth = 0.00302

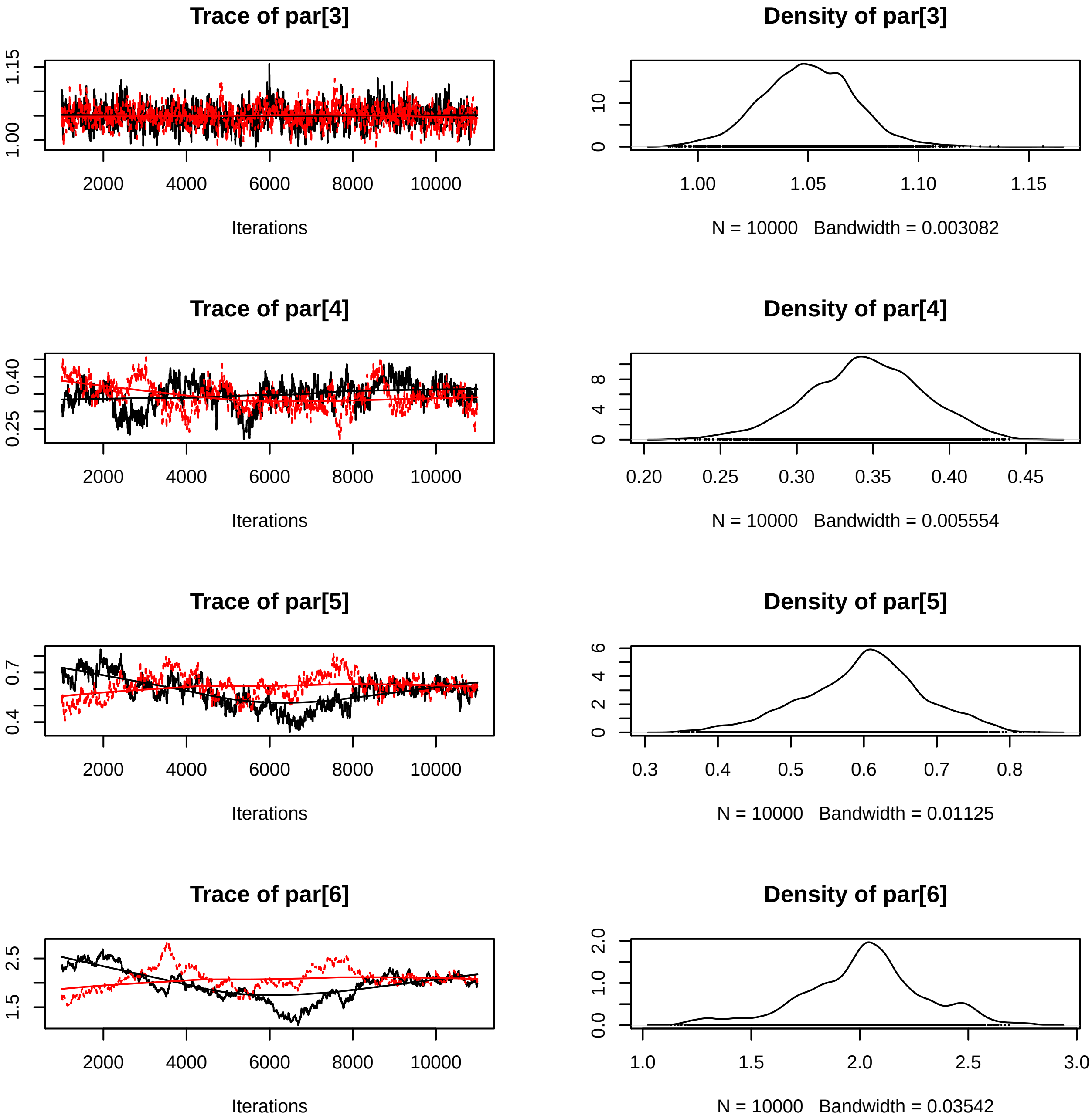

Trace of par[3]
Iterations
Density of par[3]
N = 10000 Bandwidth = 0.003082
Trace of par[4]
Iterations
Density of par[4]
N = 10000 Bandwidth = 0.005554
Trace of par[5]
Iterations
Density of par[5]
N = 10000 Bandwidth = 0.01125
Trace of par[6]
Iterations
Density of par[6]
N = 10000 Bandwidth = 0.03542

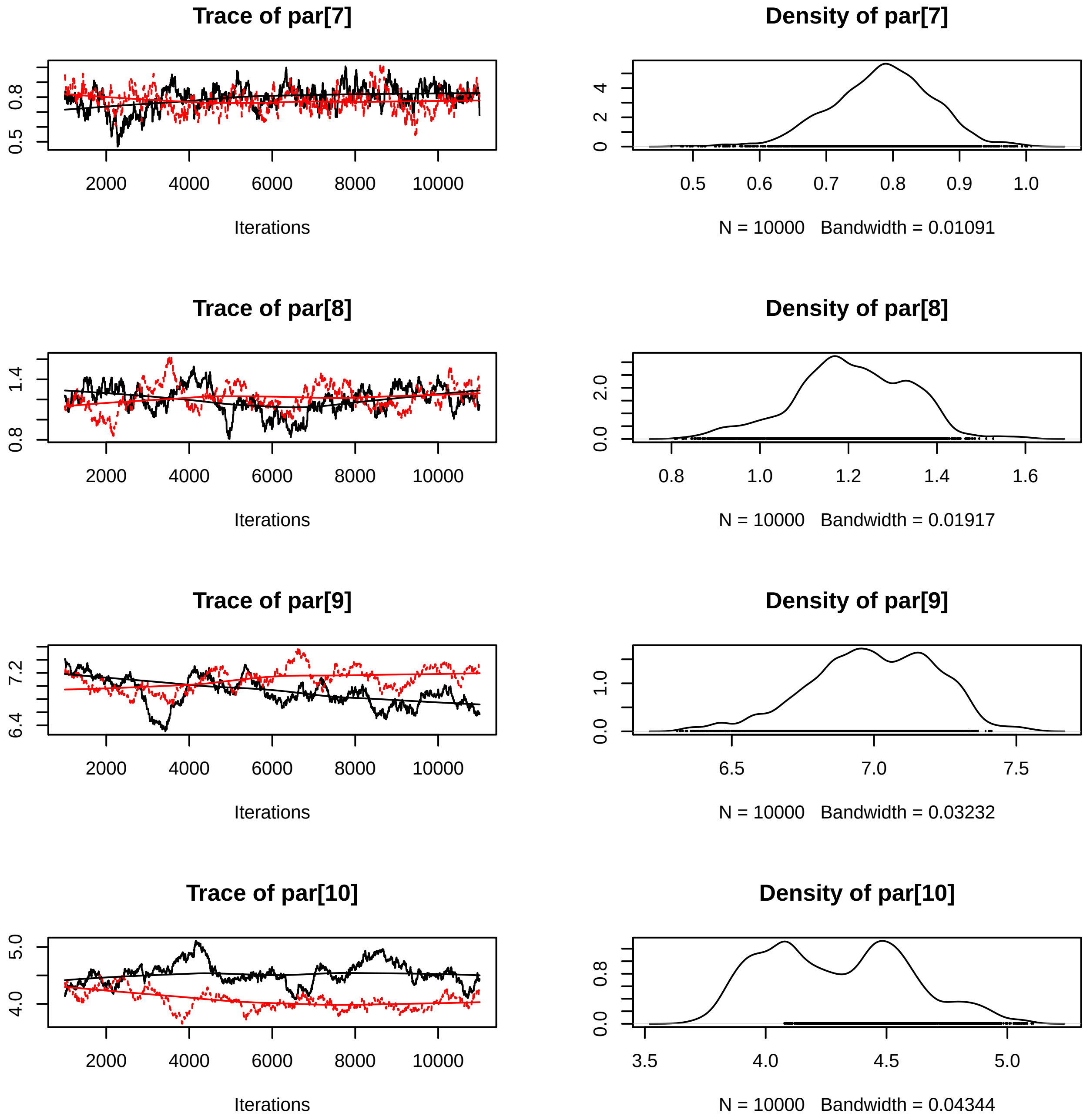
Trace of par[7]
Iterations
Density of par[7]
N = 10000 Bandwidth = 0.01091
Trace of par[8]
Iterations
Density of par[8]
N = 10000 Bandwidth = 0.01917
Trace of par[9]
Iterations
Density of par[9]
N = 10000 Bandwidth = 0.03232
Trace of par[10]
Iterations
Density of par[10]
N = 10000 Bandwidth = 0.04344

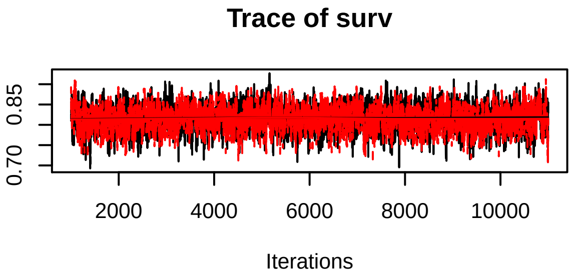
Trace of surv
0.70
0.85
2000
4000
6000
8000
10000
Iterations

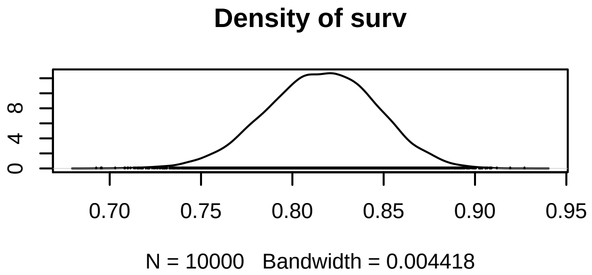
Density of surv
0
4
8
0.70
0.75
0.80
0.85
0.90
0.95
N = 10000   Bandwidth = 0.004418
