## Supplementary figures and images for "Assessment of the residential Finnish wolf population combines DNA captures, citizen observations and mortality data using a Bayesian state-space model"

### Nov_ENG_Revonlahti.png

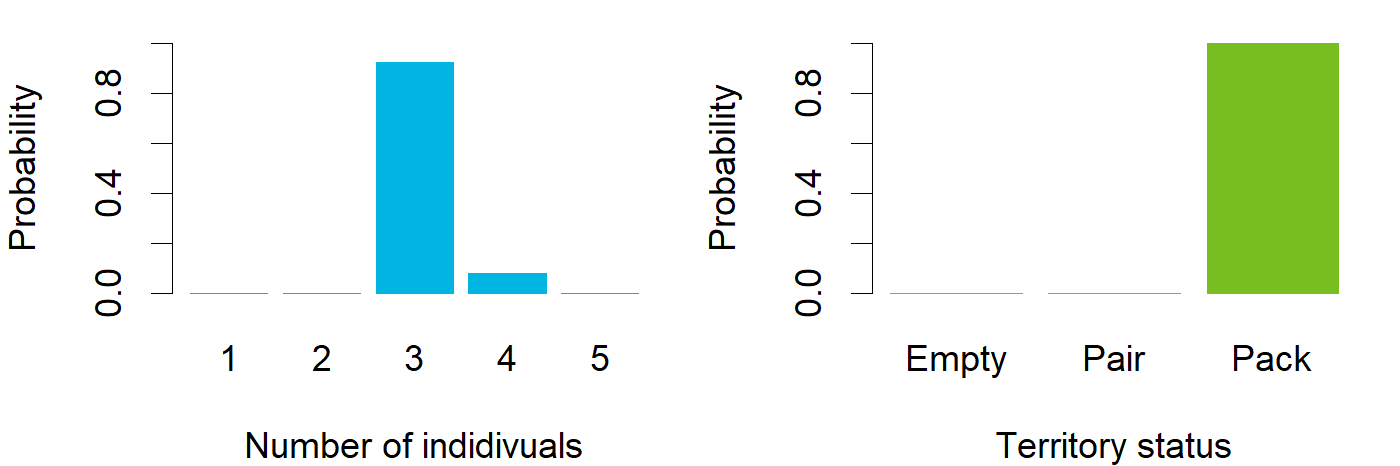

### Nov_ENG_Saramo.png

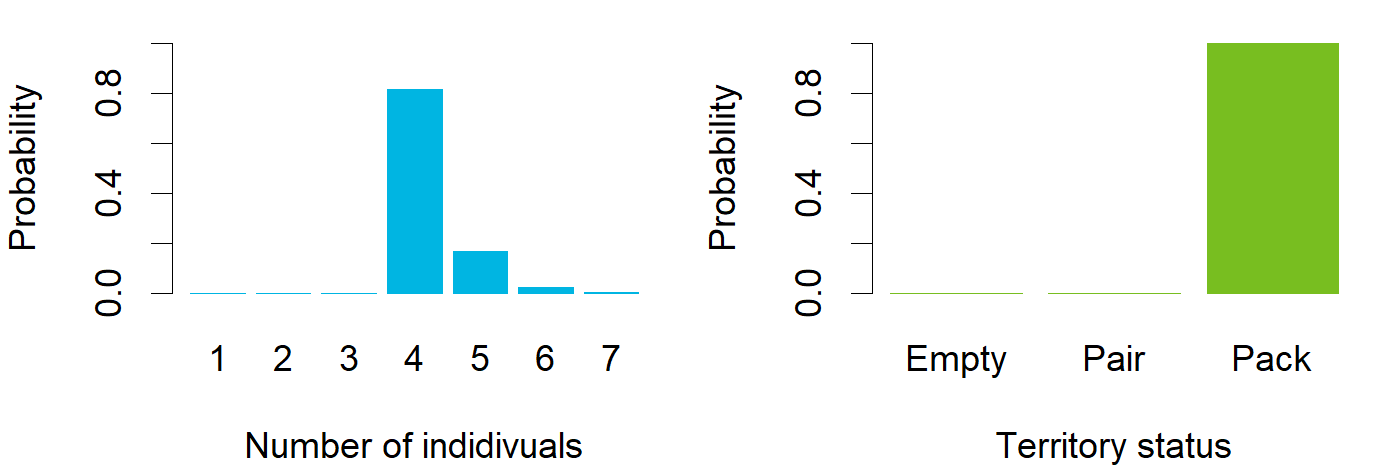

### Nov_ENG_Saunajärvi.png

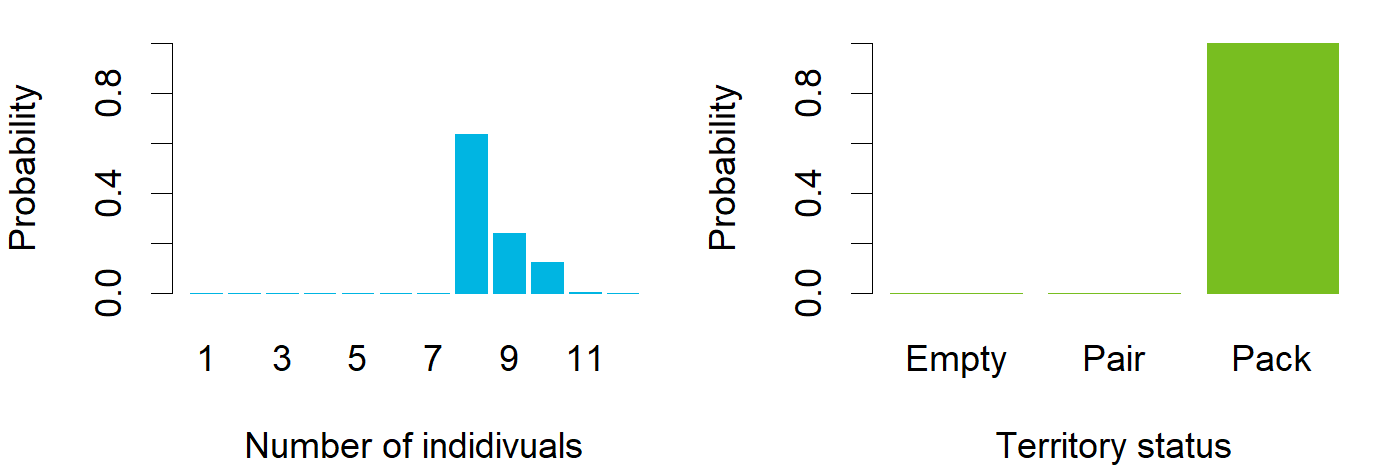

### Nov_ENG_Snappertuna.png

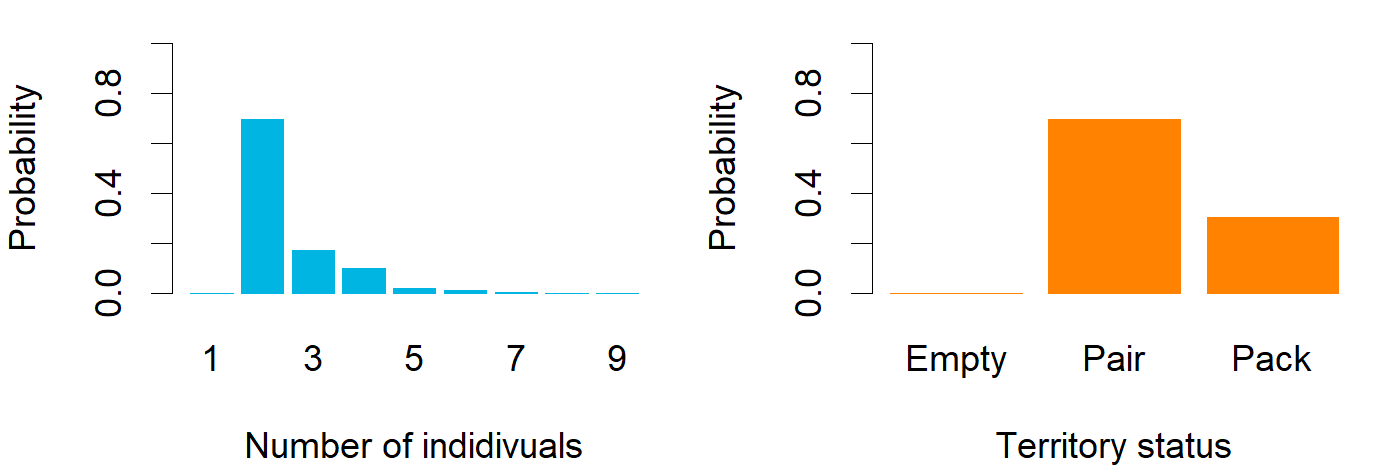

### Nov_ENG_Somerniemi.png

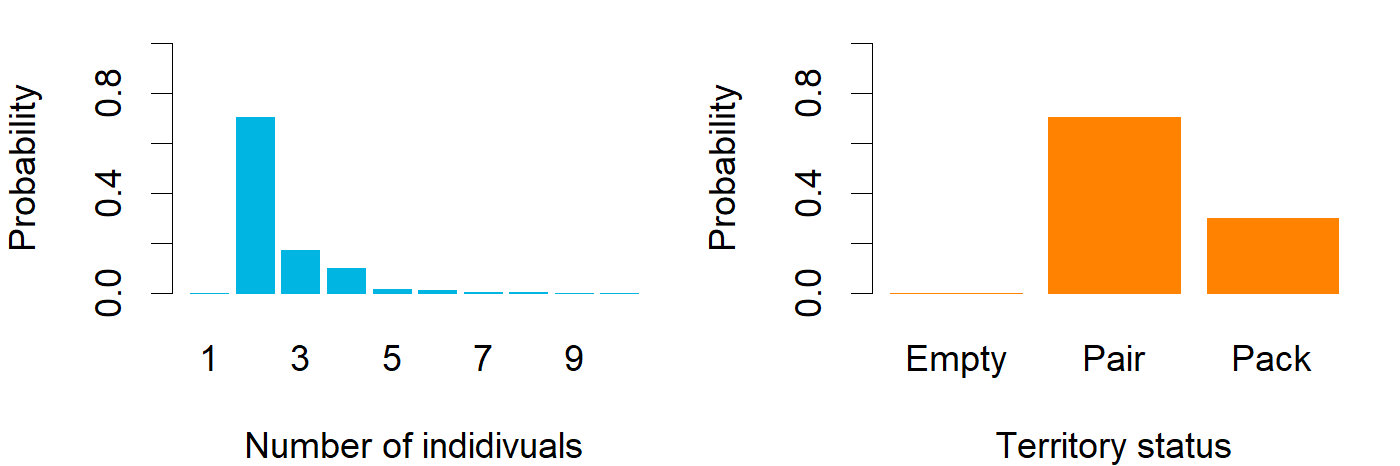

### Nov_ENG_territorytotals.png

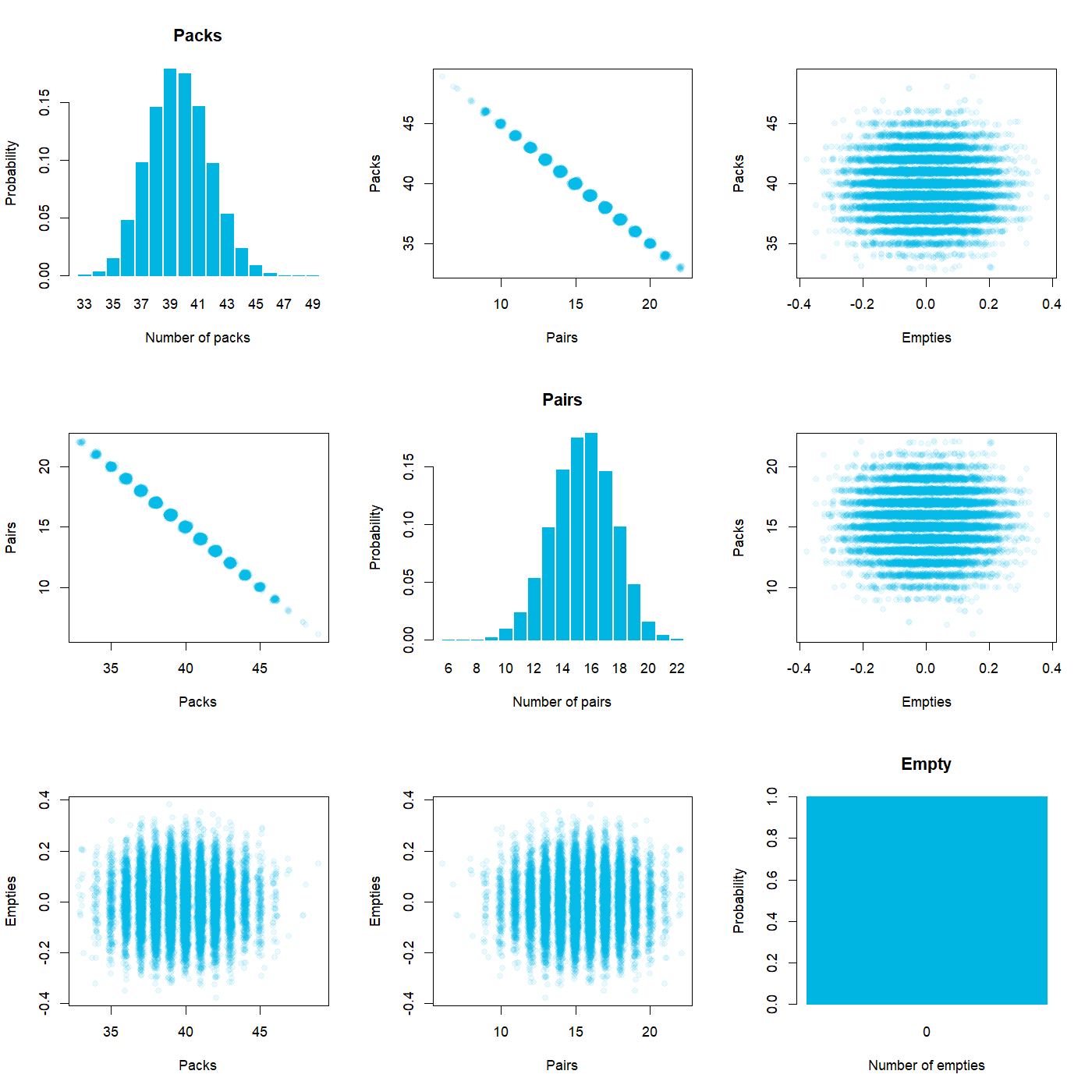

### Nov_ENG_Toholampi.png

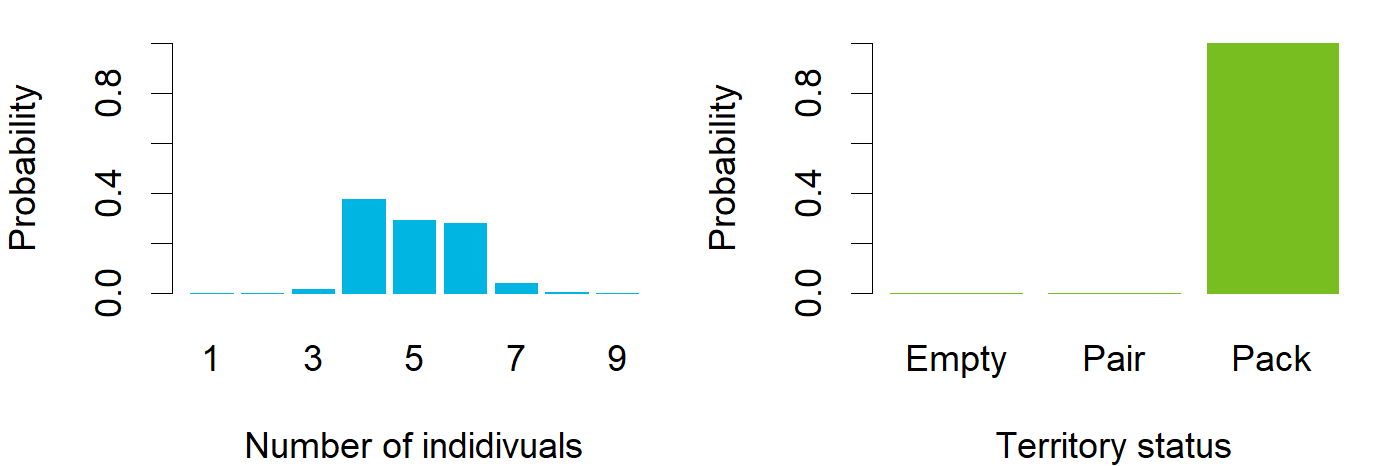

### Nov_ENG_Toijala.png

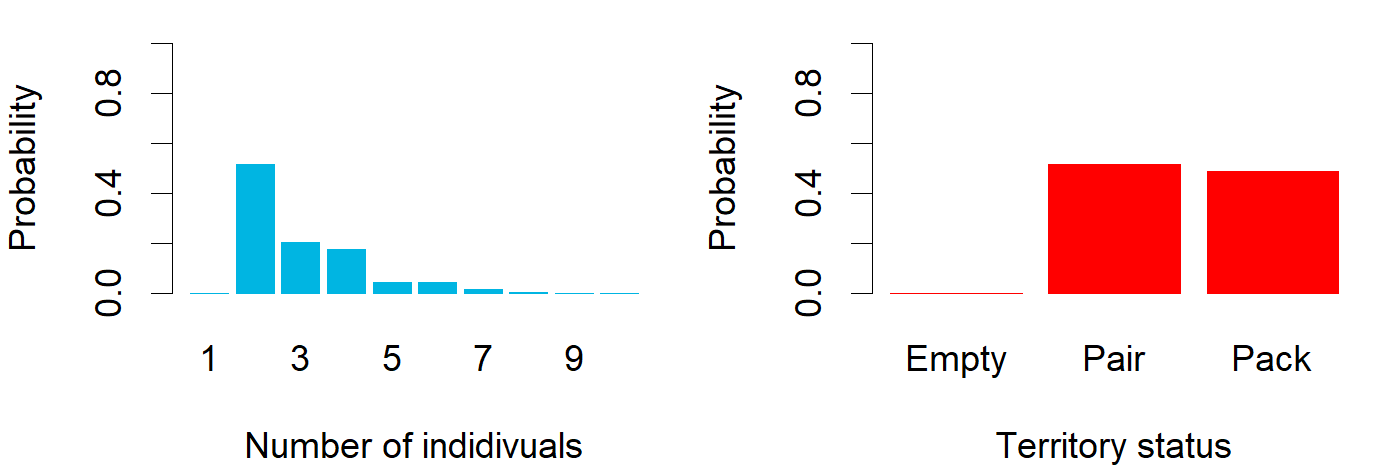

### Nov_ENG_Utajärvi.png

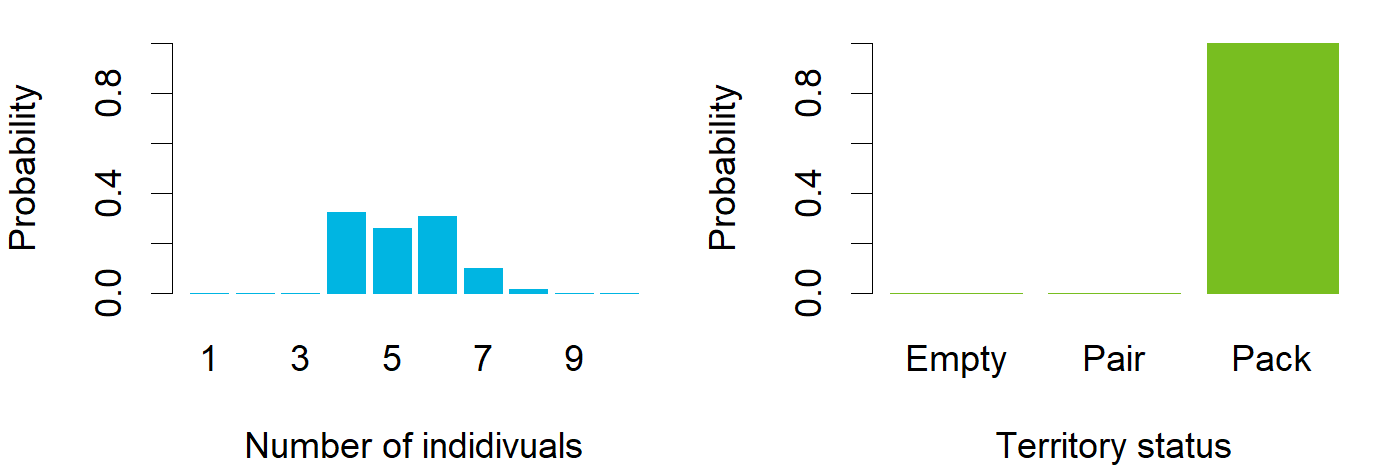

### Nov_ENG_Uura.png

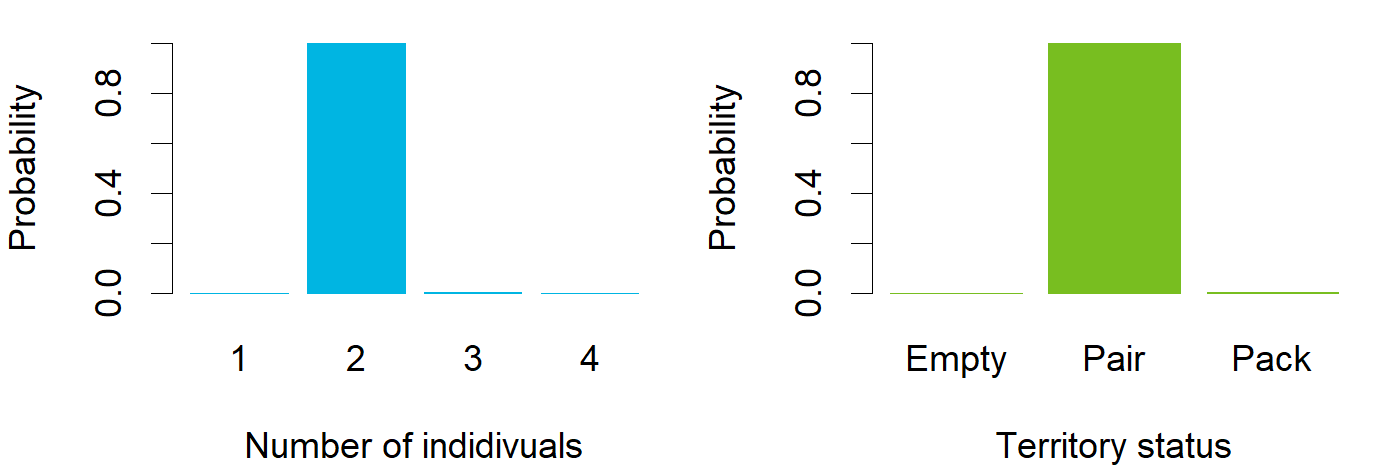

### Nov_ENG_Vartius.png

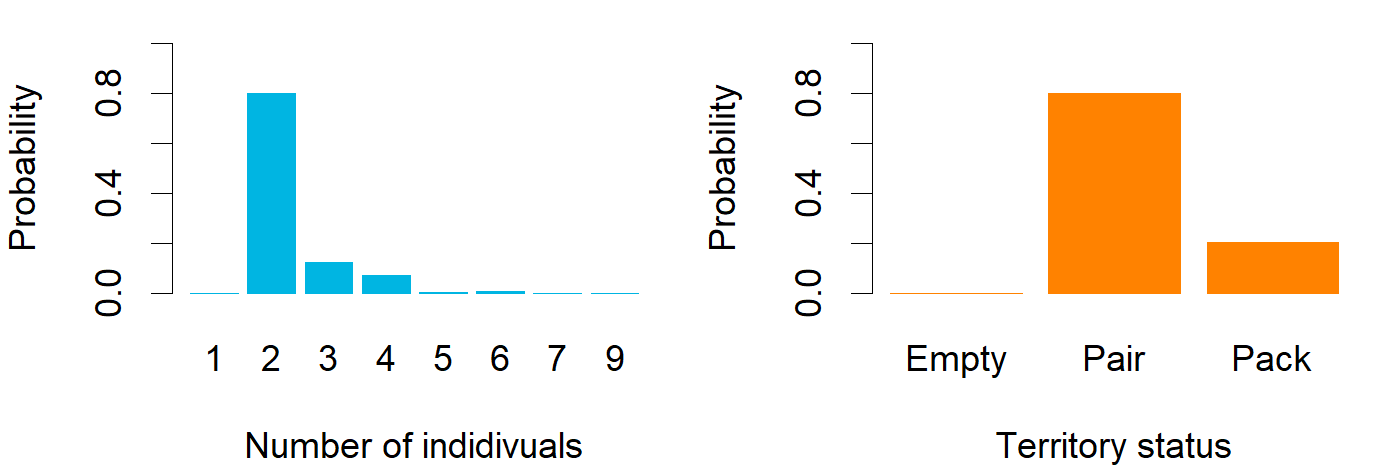

### Nov_ENG_Vuonisjärvi.png

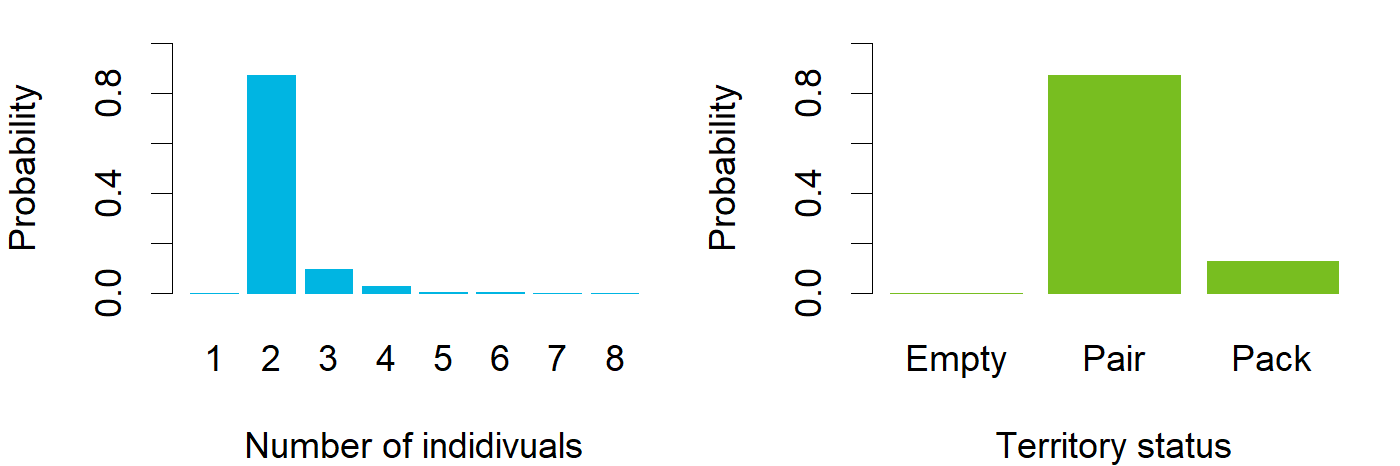

### Nov_ENG_Värtsilä.png

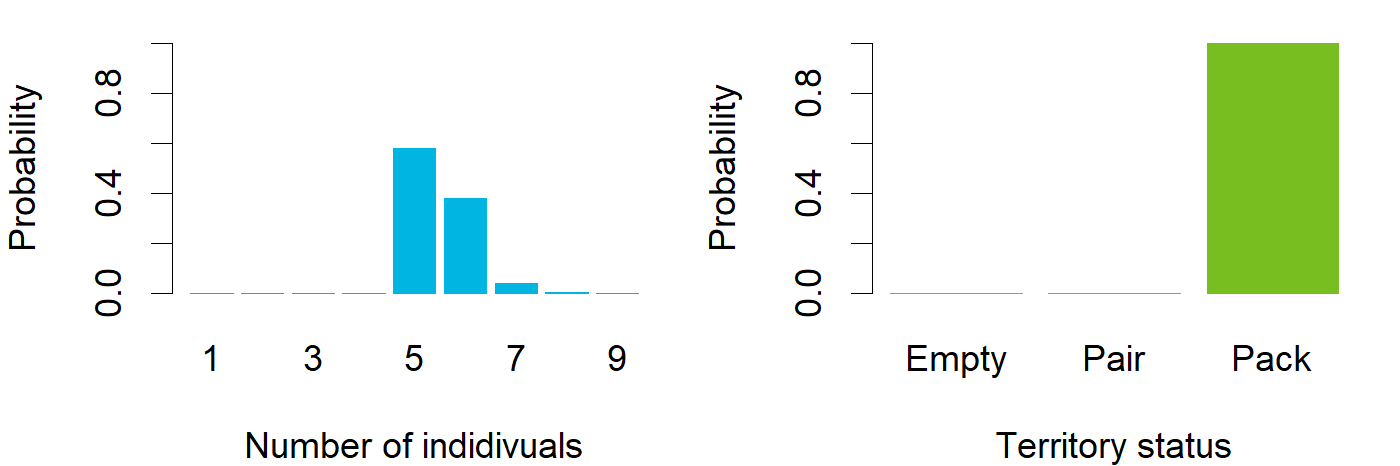

### packsize_vs_pair_obs.png

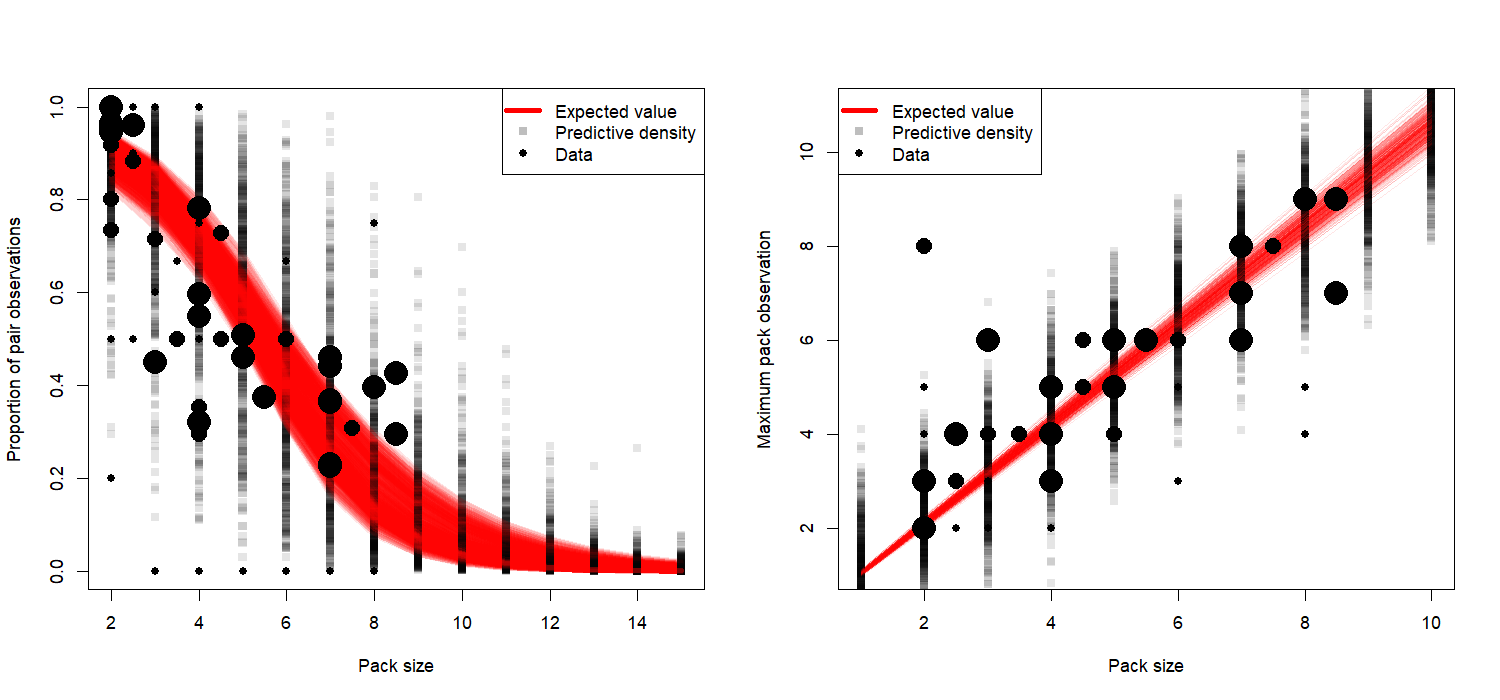
