## Supplementary figures and images for "Assessment of the residential Finnish wolf population combines DNA captures, citizen observations and mortality data using a Bayesian state-space model"

### expert_comparison.png

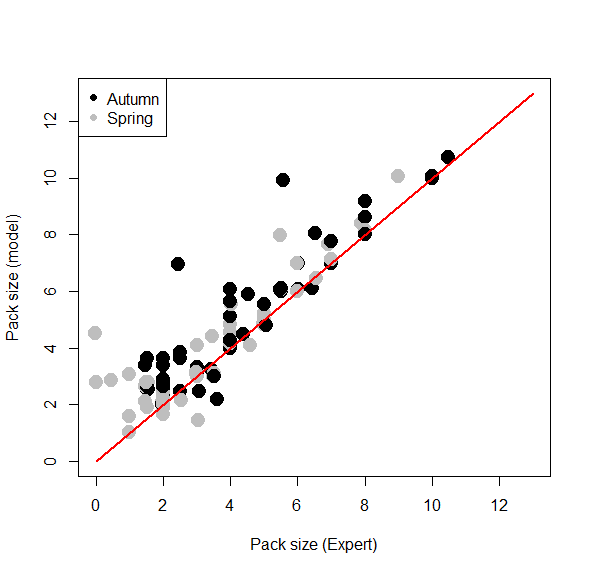

### March_ENG_Eurajoki.png

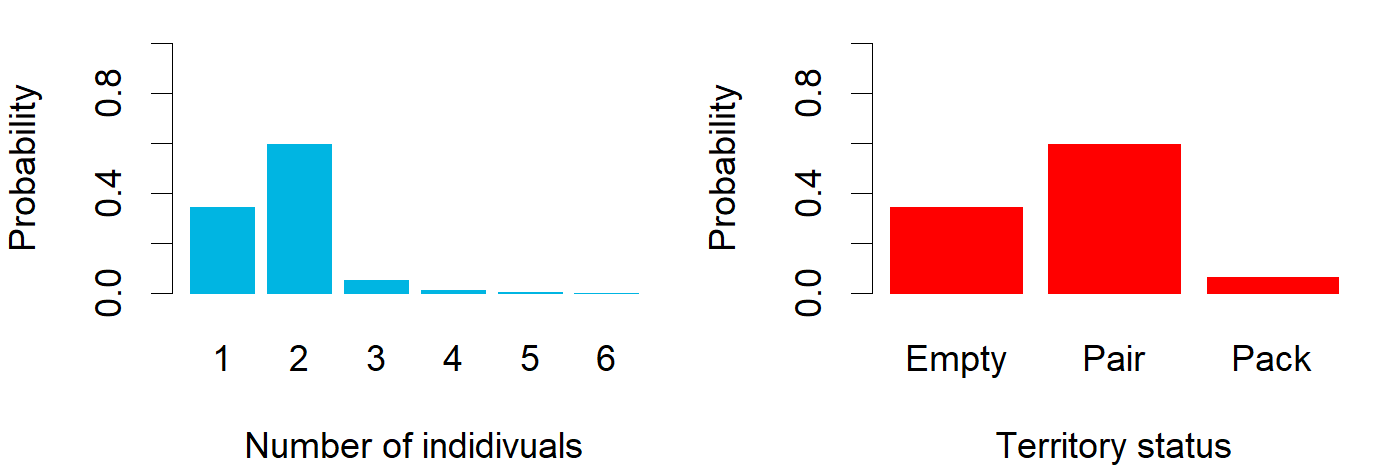

### March_ENG_Haapajärvi.png

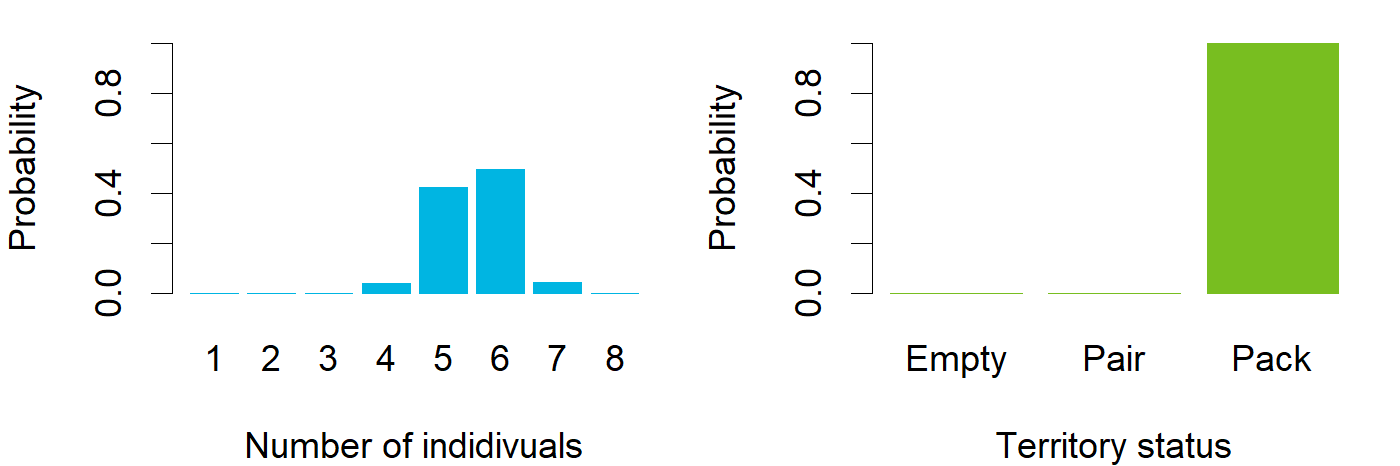

### March_ENG_Halivaara.png

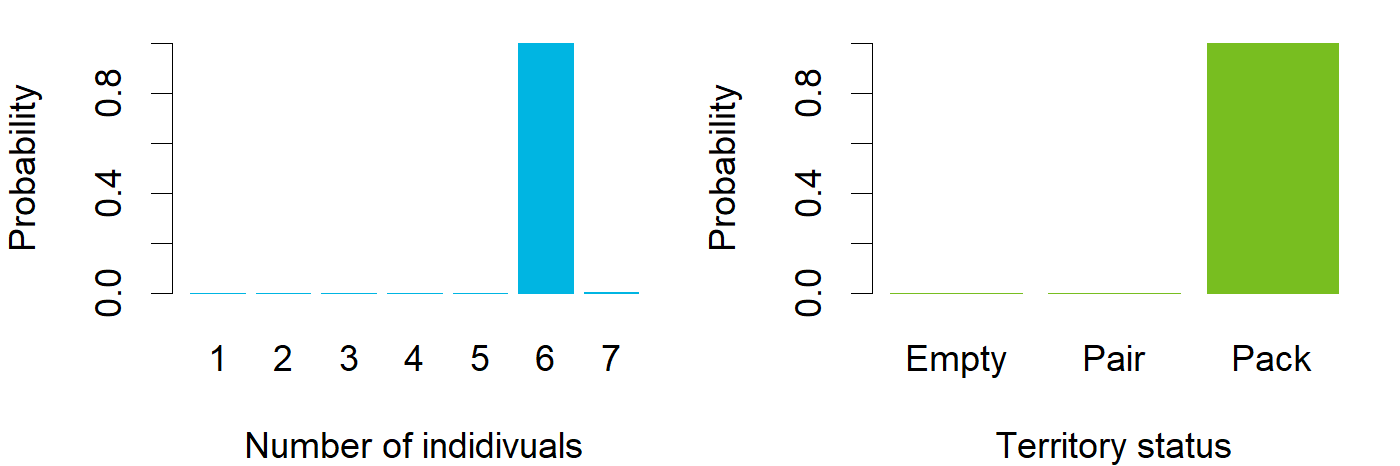

### March_ENG_Honkajoki.png

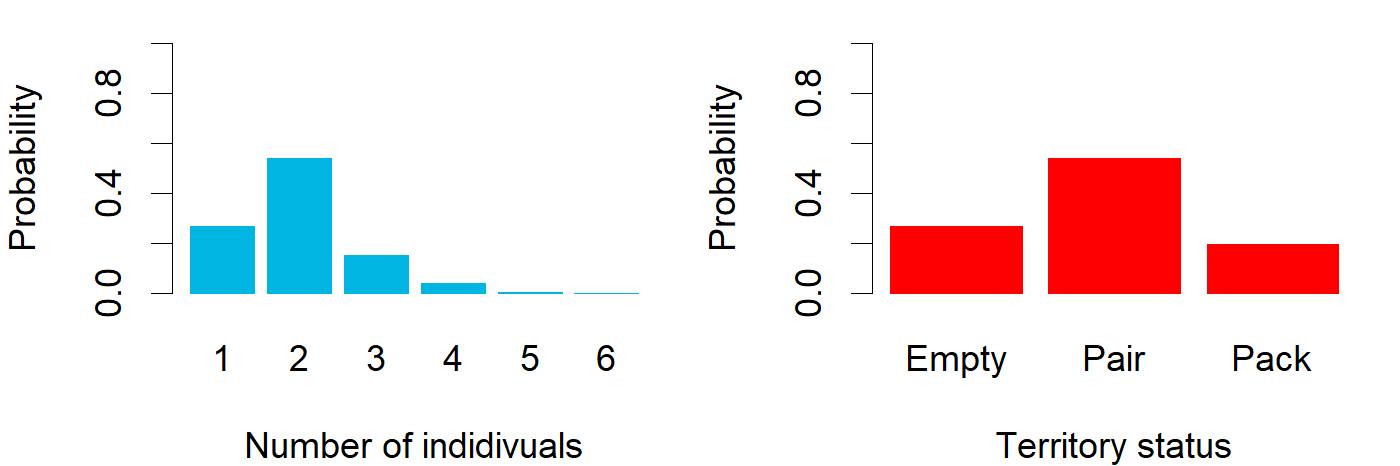

### March_ENG_Höljäkkä.png

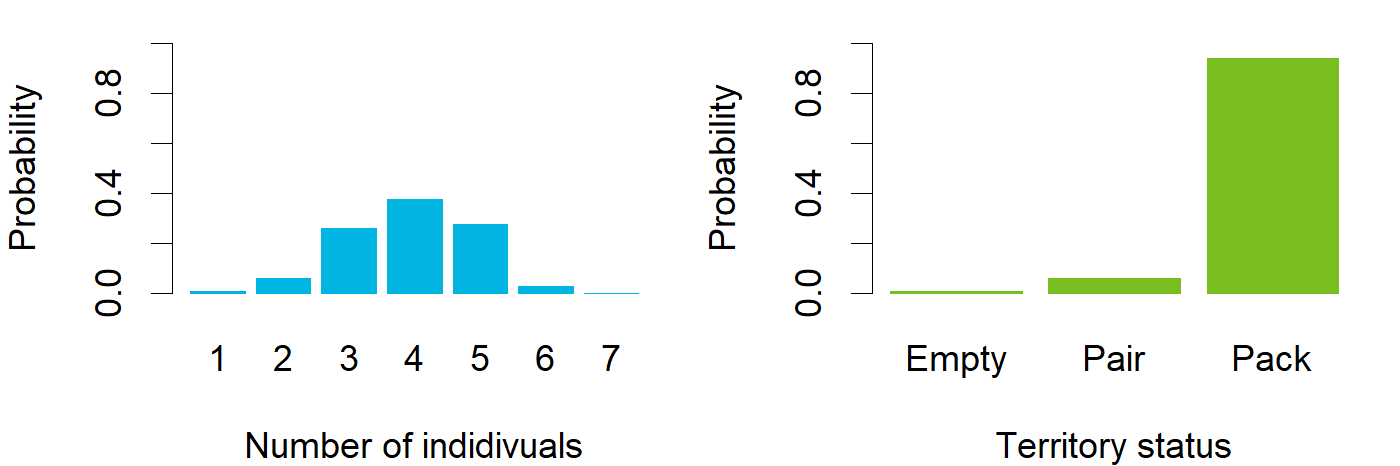

### March_ENG_Ihode.png

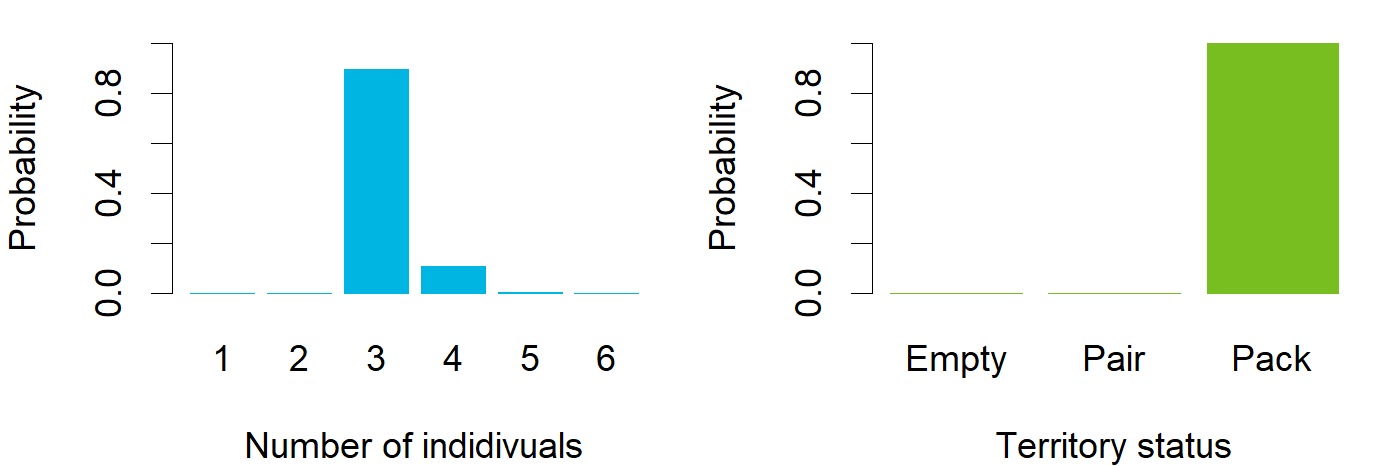

### March_ENG_Inari.png

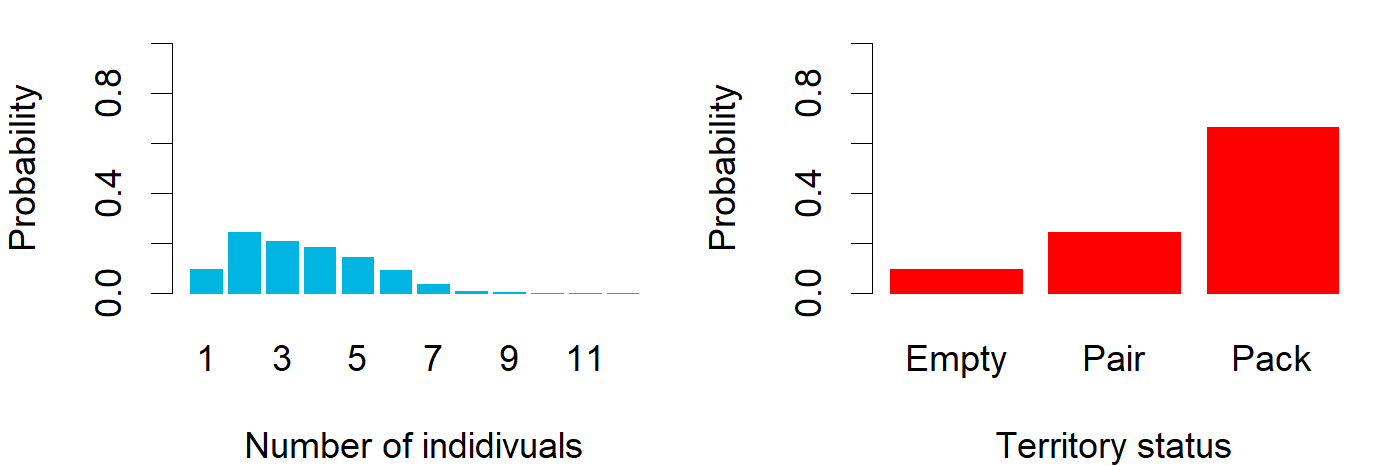

### March_ENG_Isojoki.png

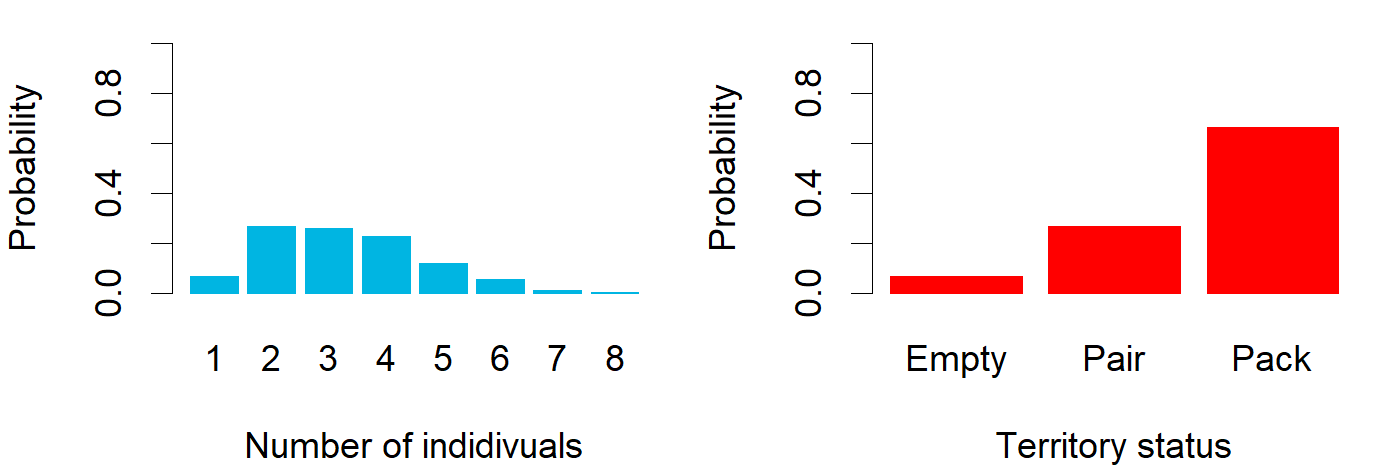

### March_ENG_Jeppo.png

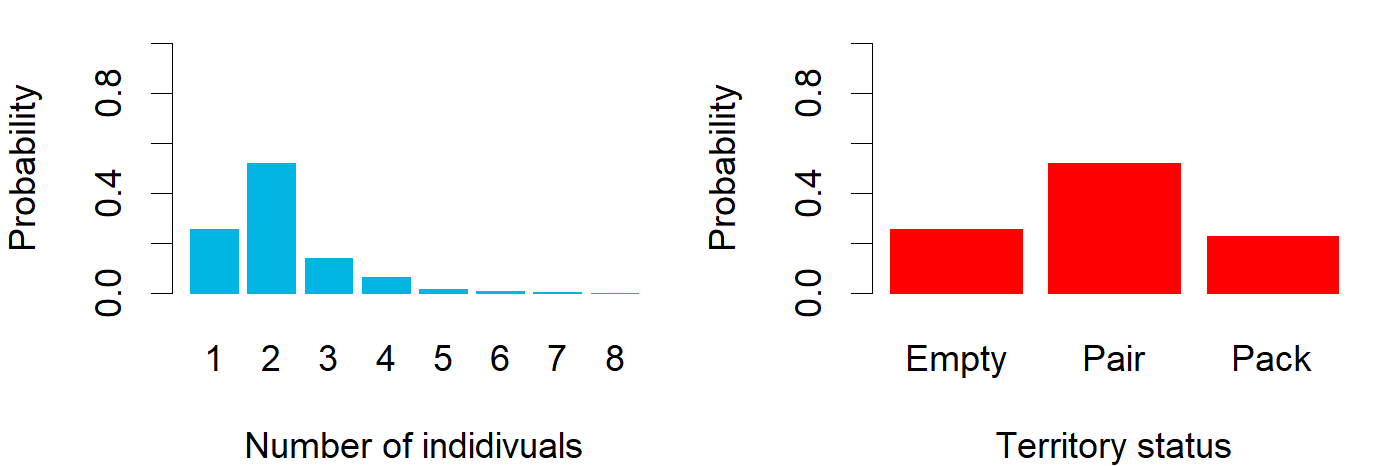

### March_ENG_Juntti.png

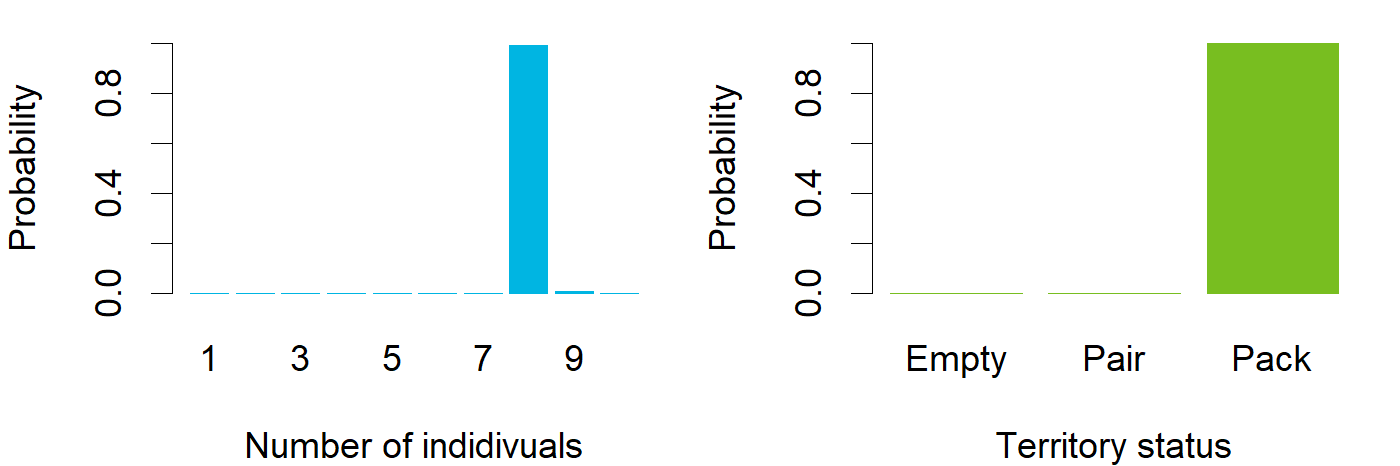

### March_ENG_Jurva.png

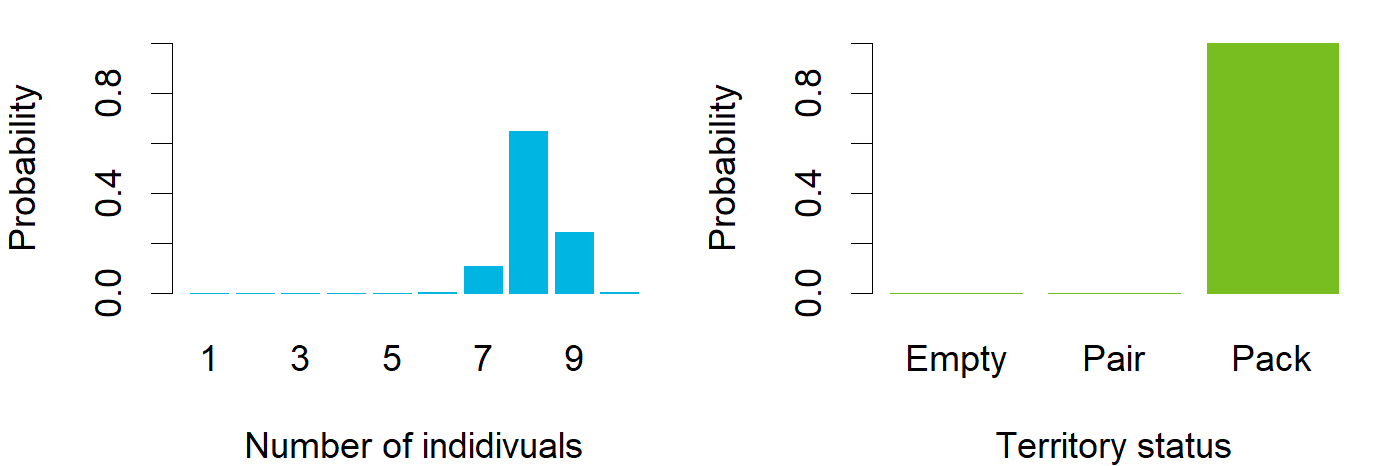

### March_ENG_Juupajoki.png

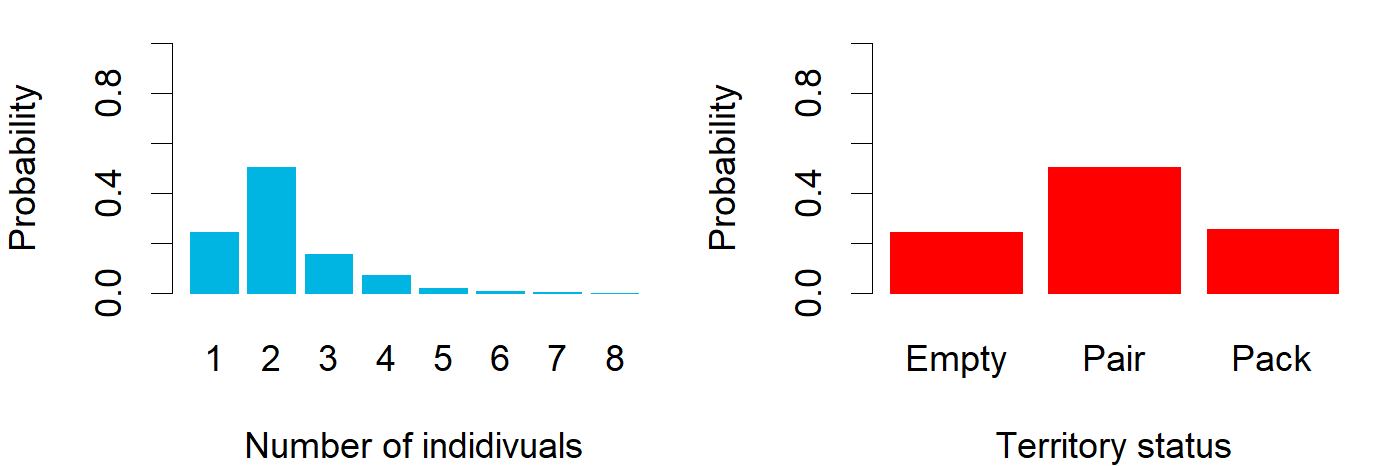

### March_ENG_Kaivola.png

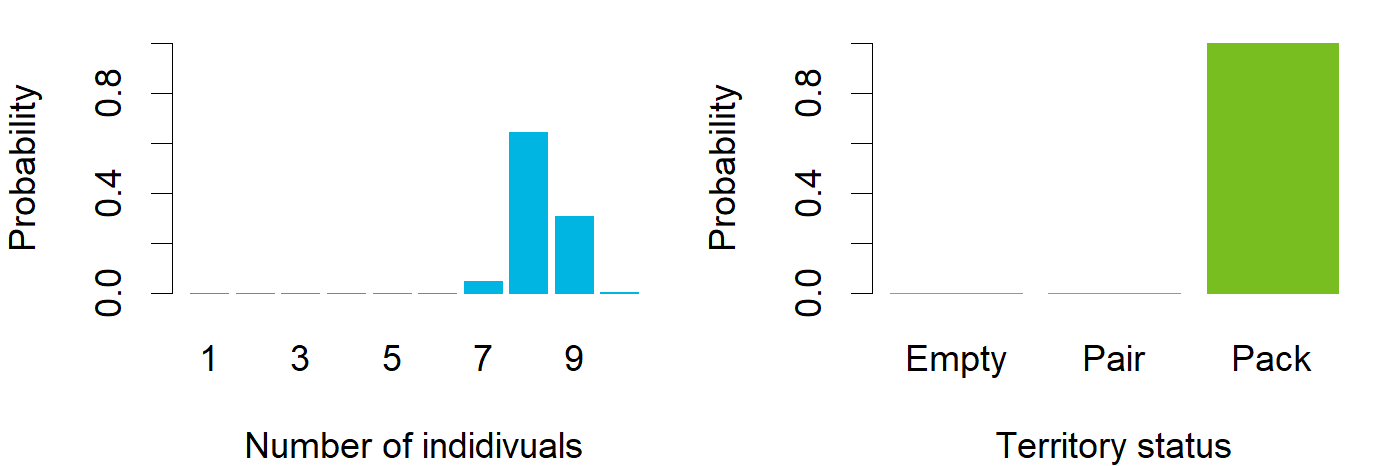

### March_ENG_Kallioluoma.png

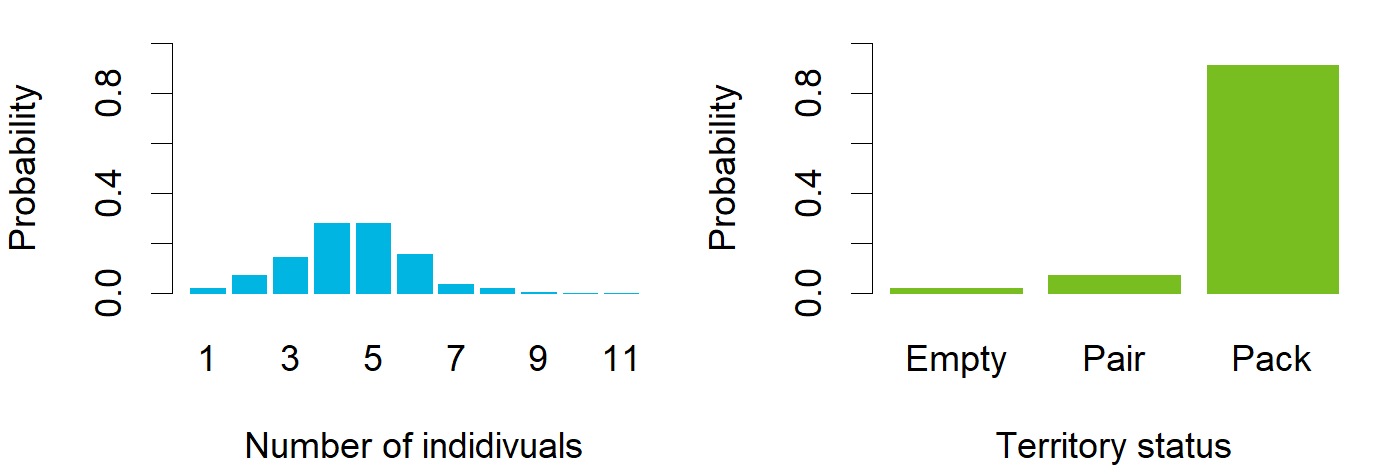

### March_ENG_Kankaanpää.png

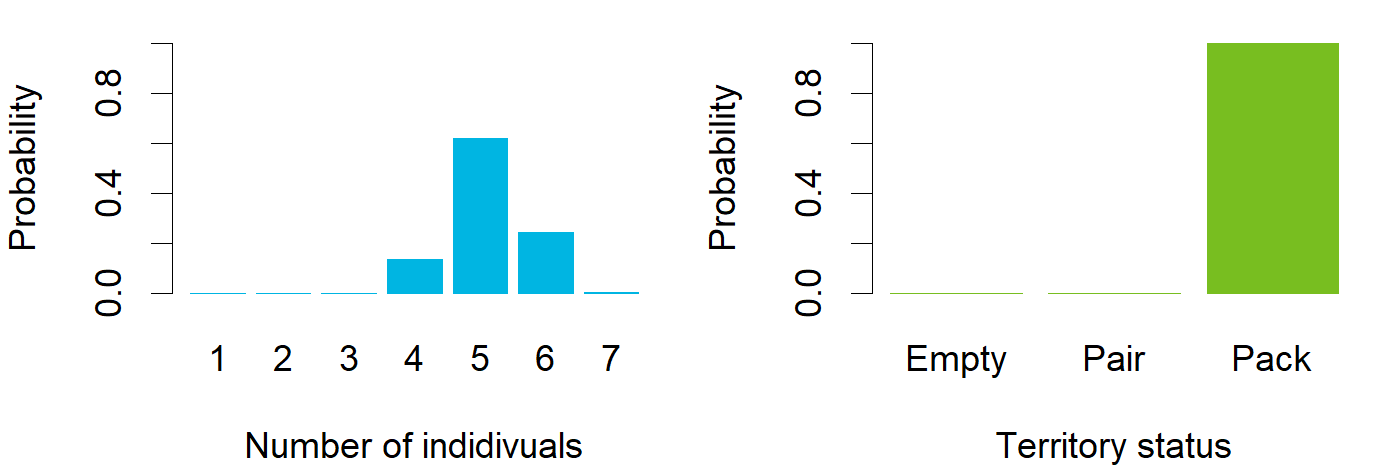

### March_ENG_Karvia.png

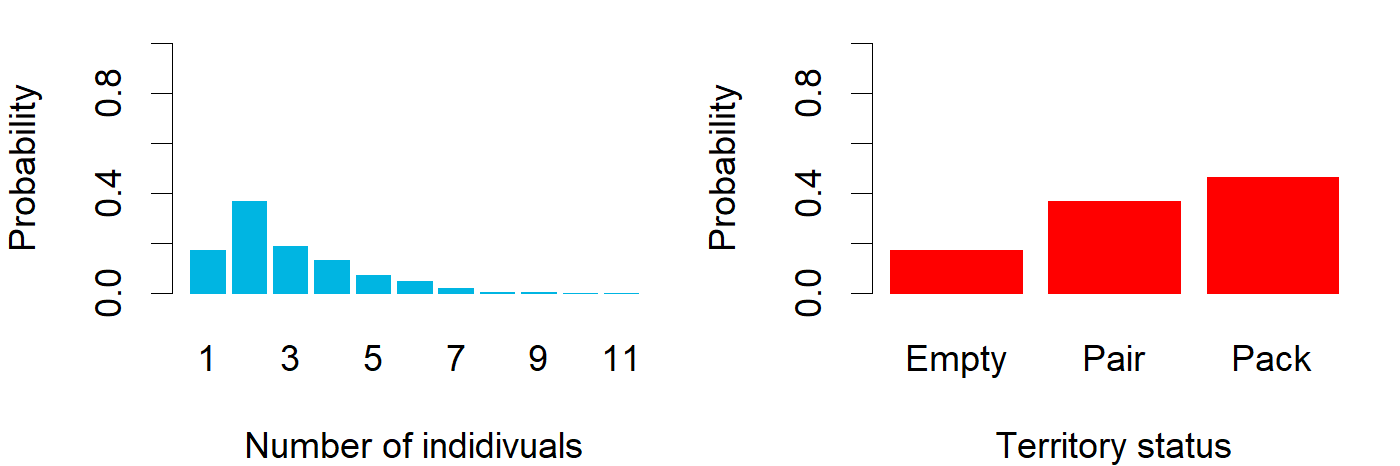

### March_ENG_Kelsimä.png

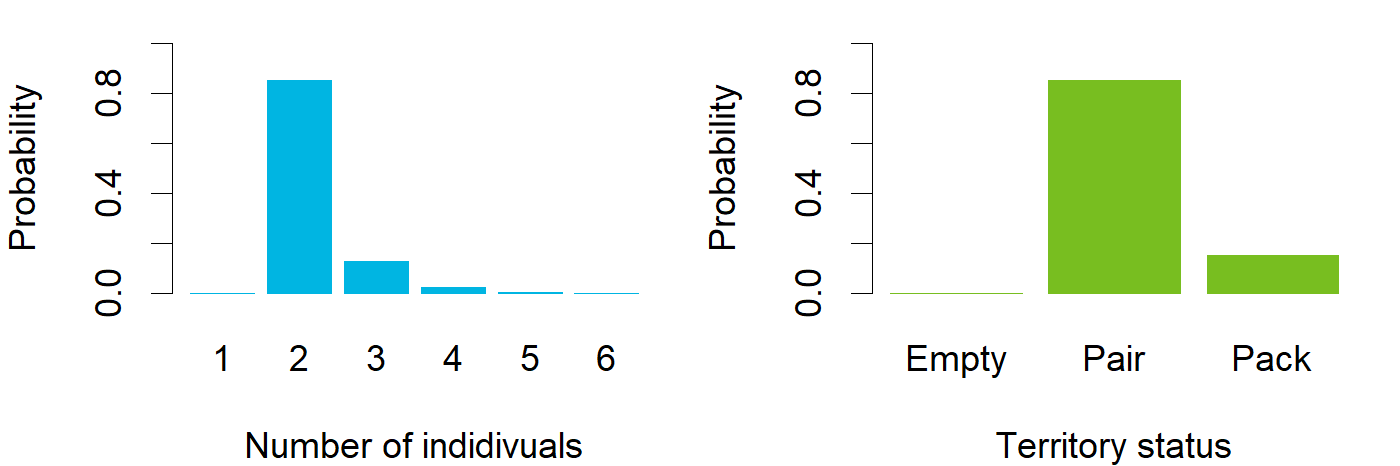

### March_ENG_Kemiö.png

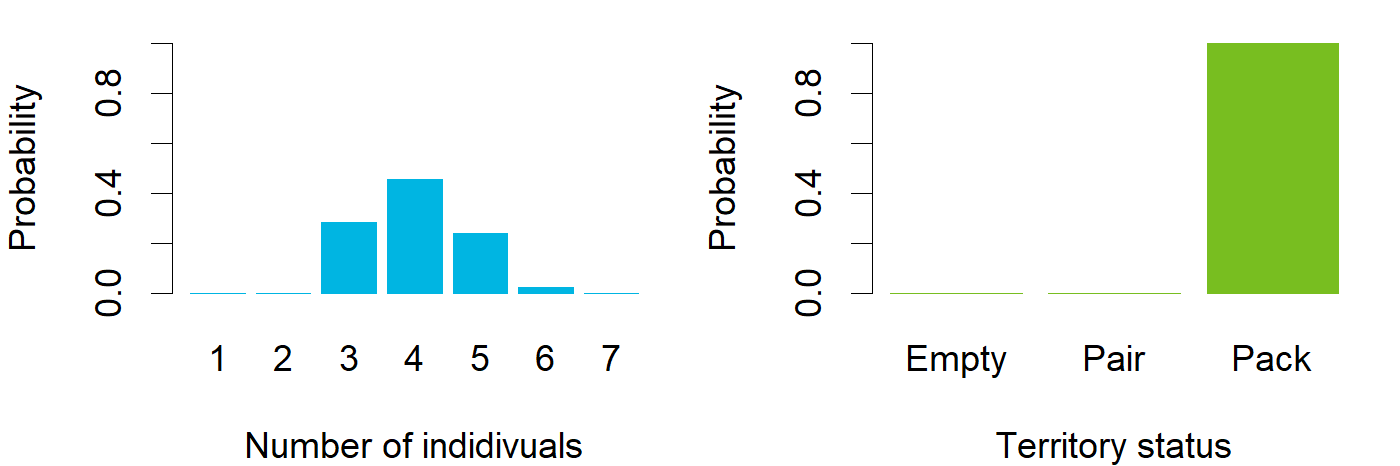

### March_ENG_Kiuruvesi.png

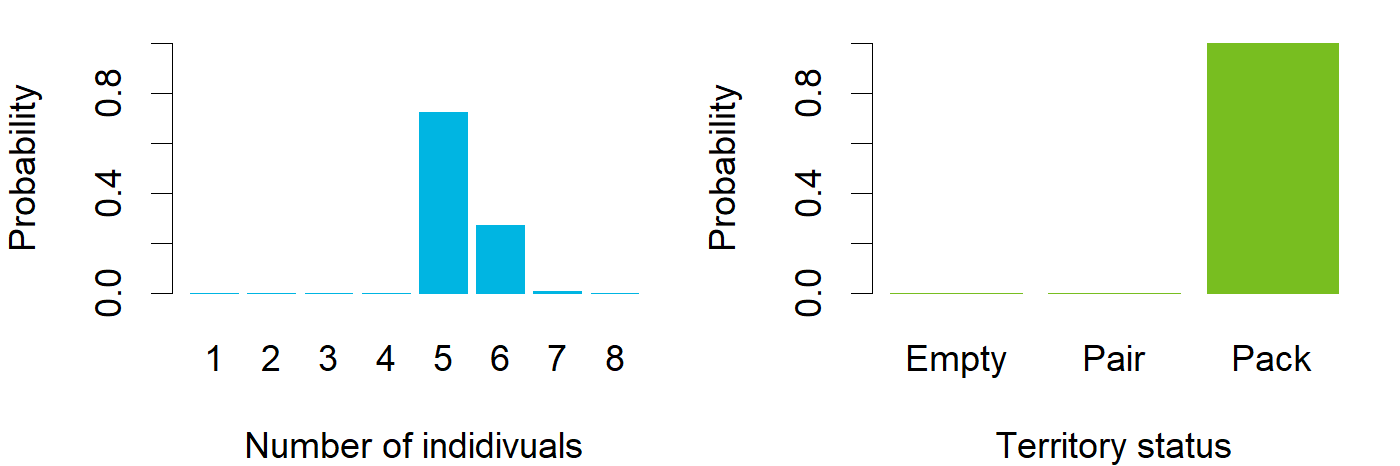

### March_ENG_Kivikiekki.png

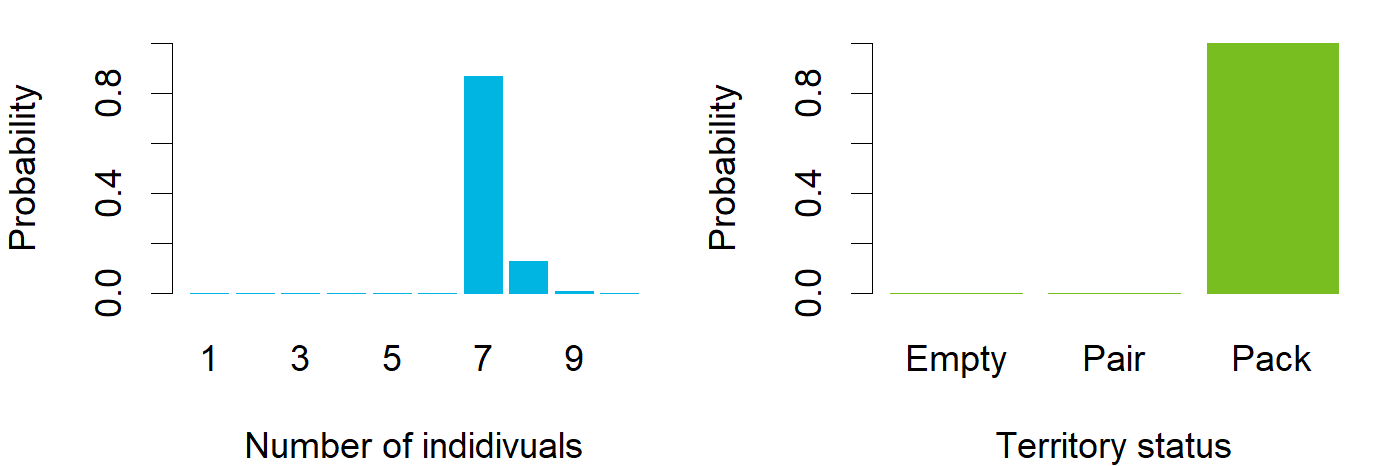

### March_ENG_Kivivaara.png

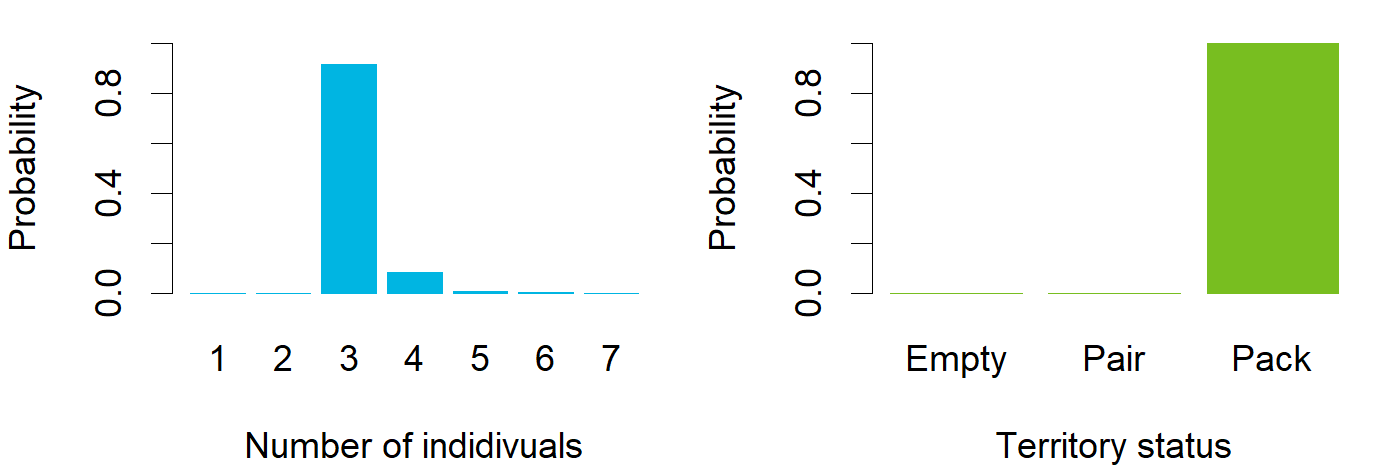

### March_ENG_Kovero.png

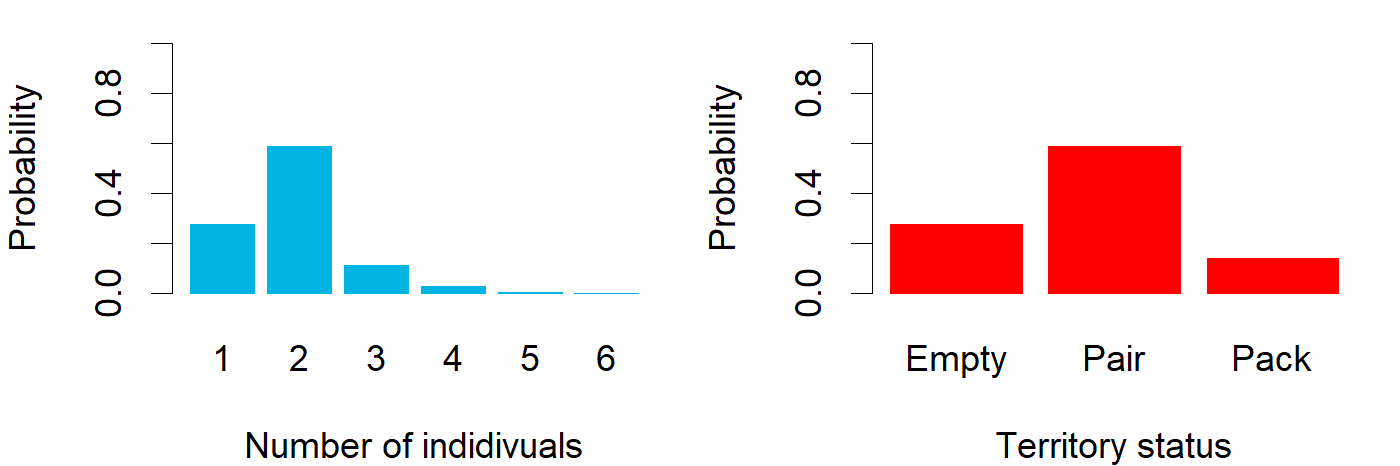

### March_ENG_Köyliö.png

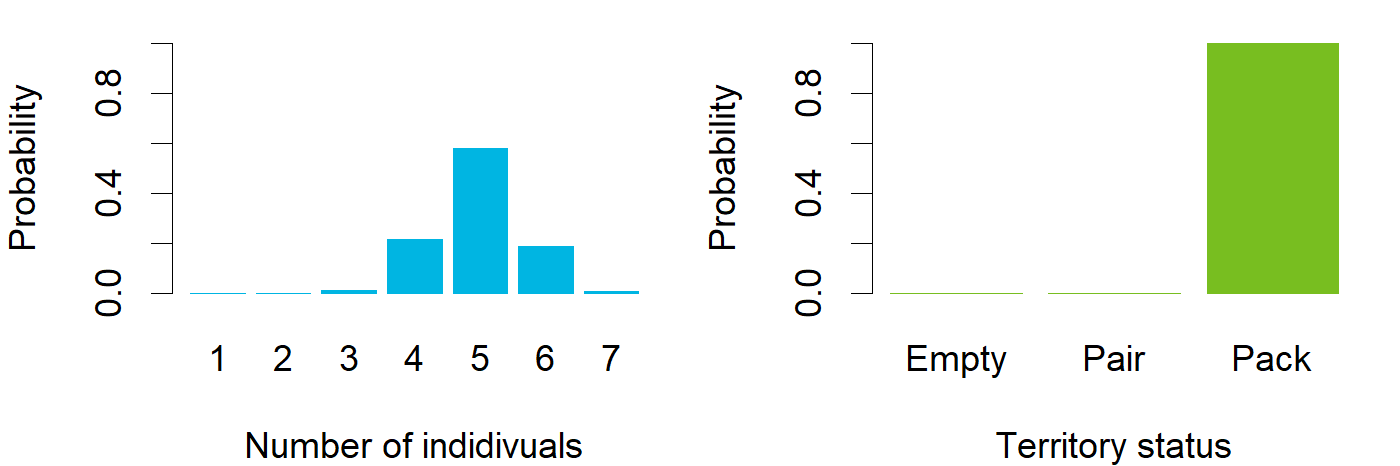

### March_ENG_Laakajärvi.png

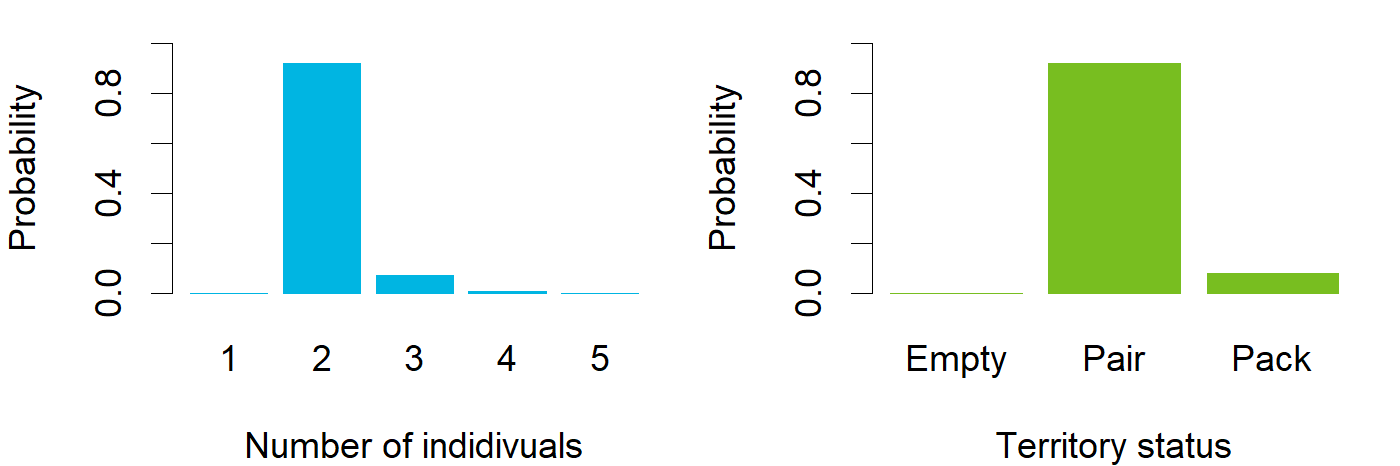

### March_ENG_Laihia.png

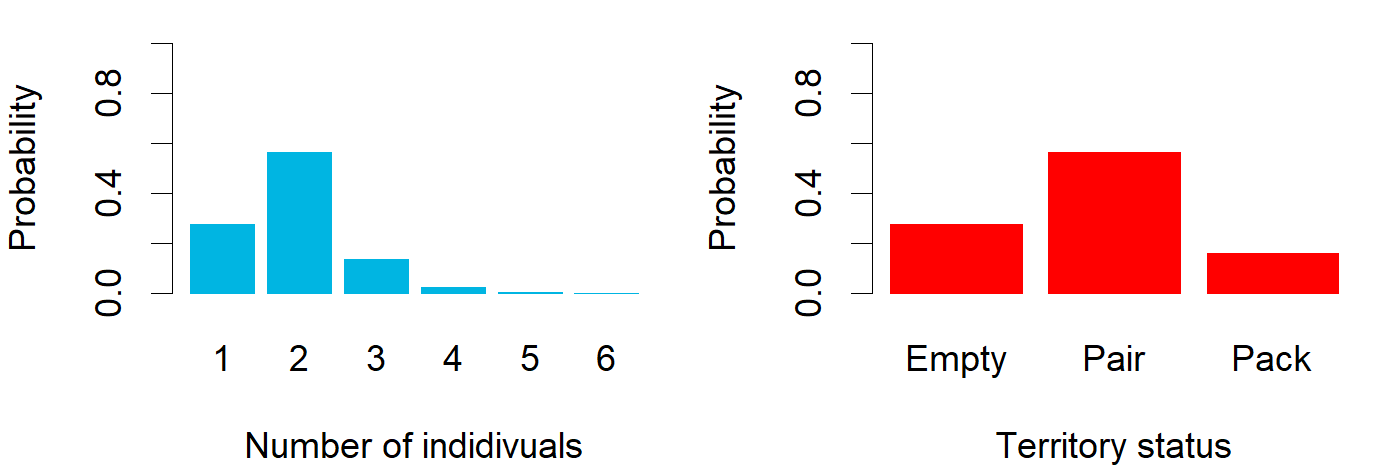

### March_ENG_Lauhavuori.png

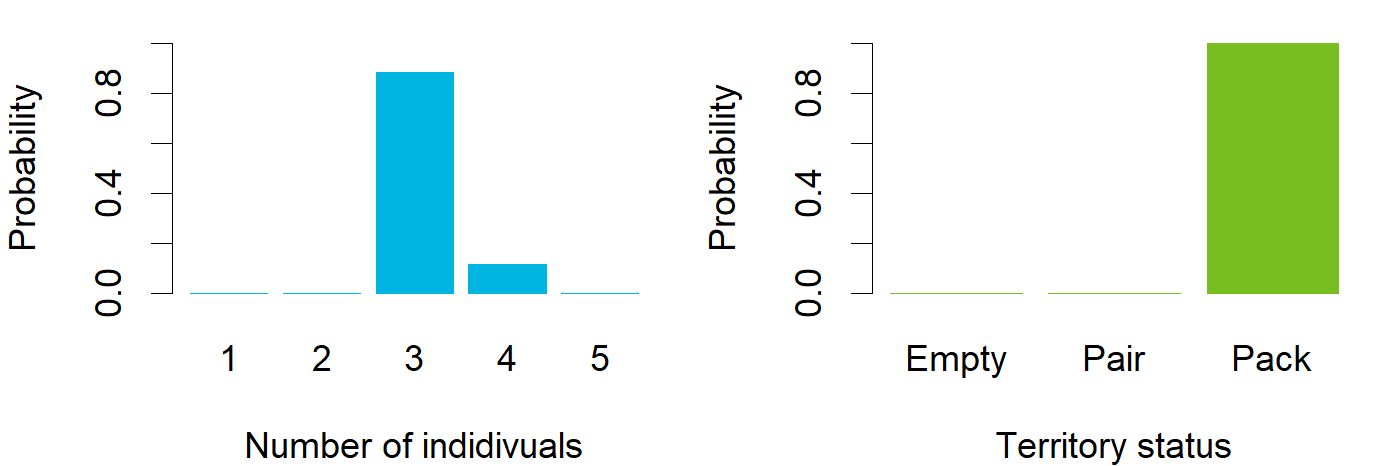

### March_ENG_Leppävirta.png

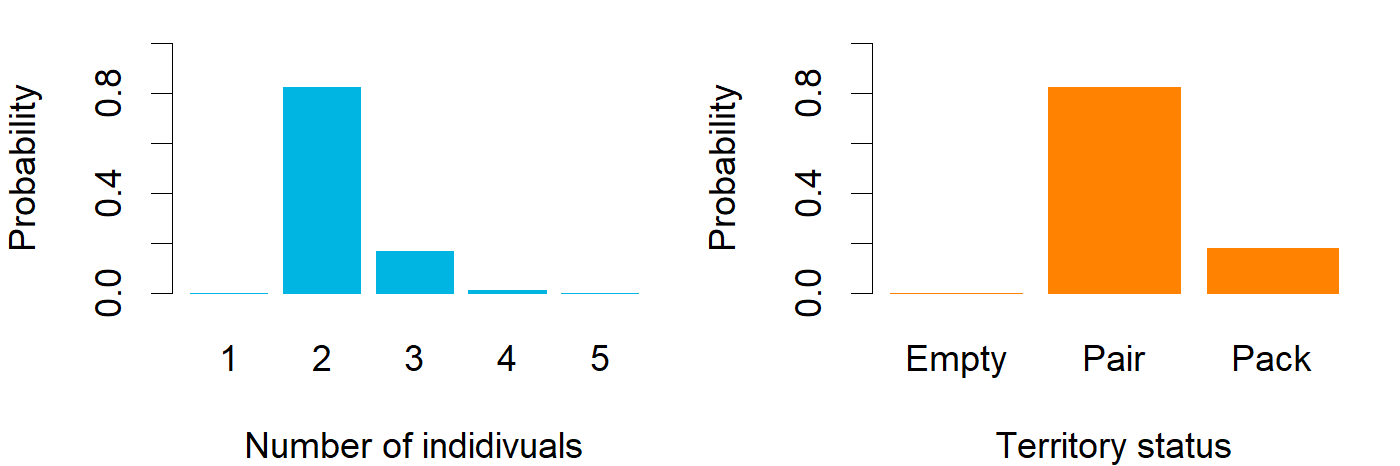

### March_ENG_Marttinen.png

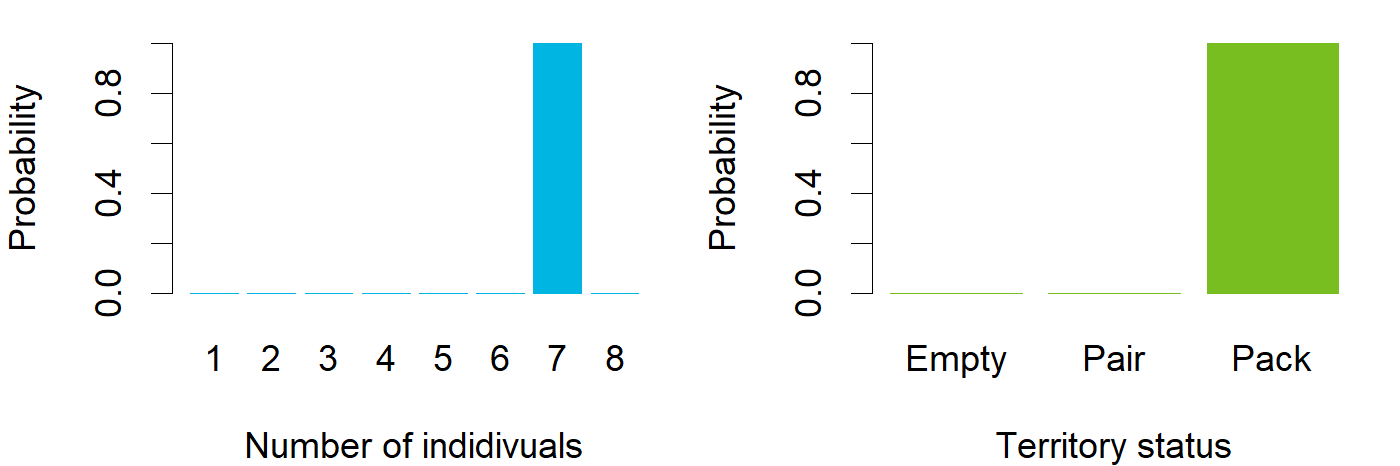

### March_ENG_Mynämäki.png

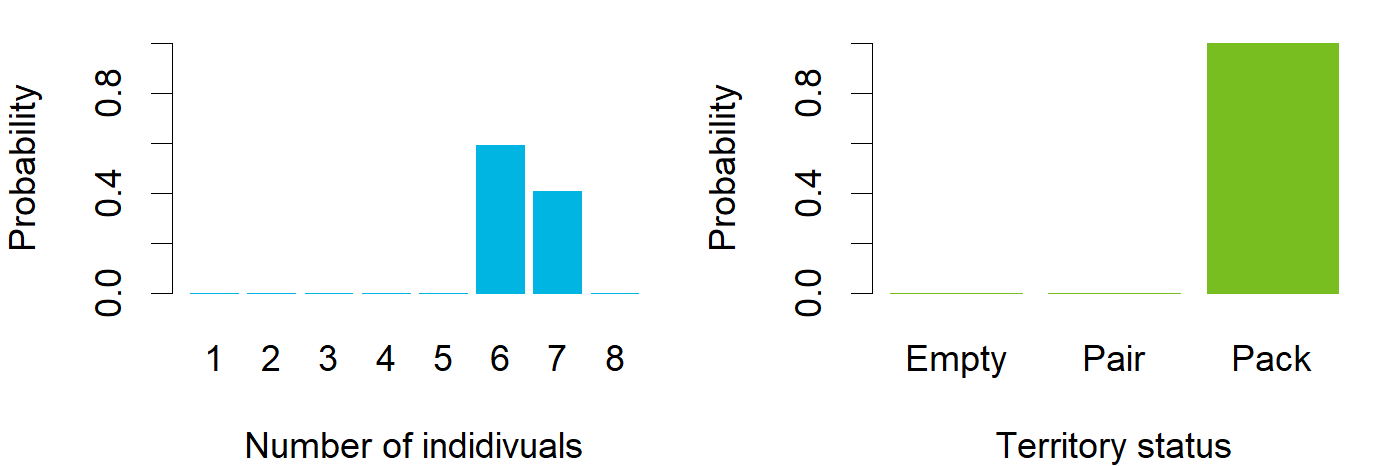

### March_ENG_Möhkö.png

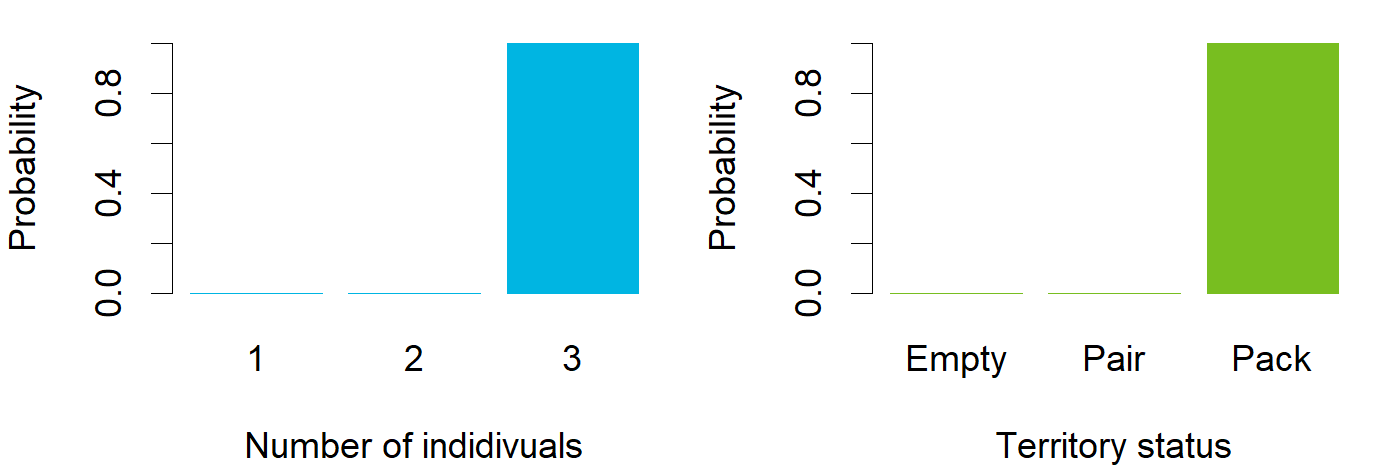

### March_ENG_Nivala.png

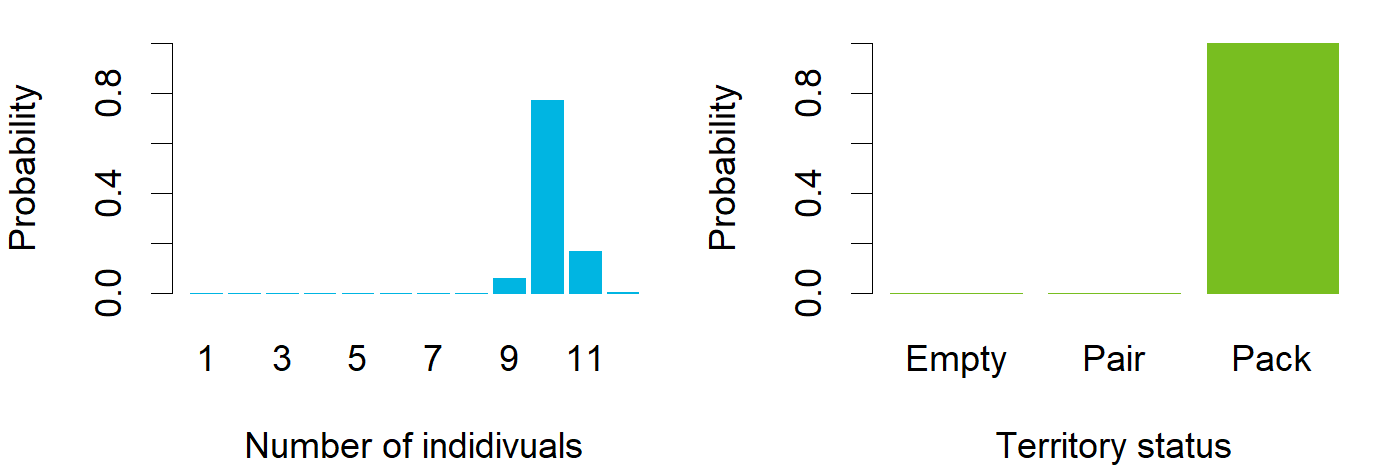

### March_ENG_Närvijoki.png

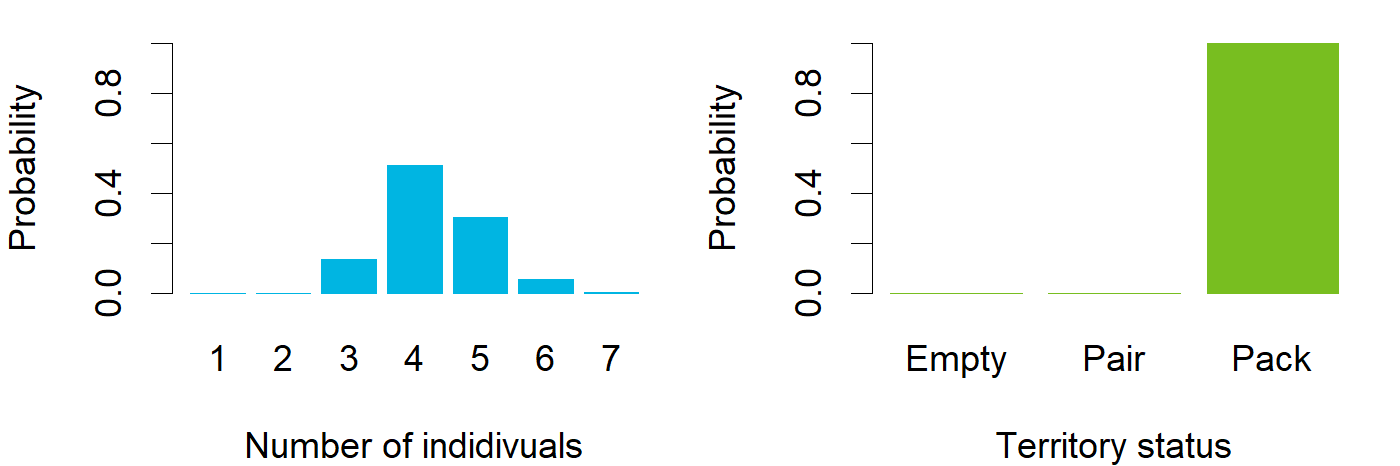

### March_ENG_Panja.png

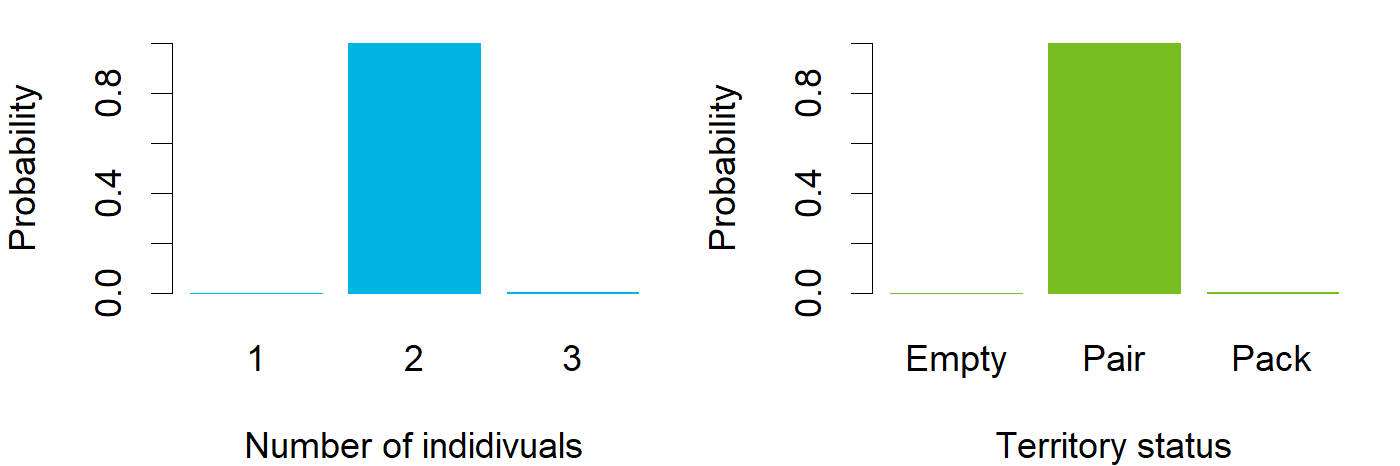

### March_ENG_Perho.png

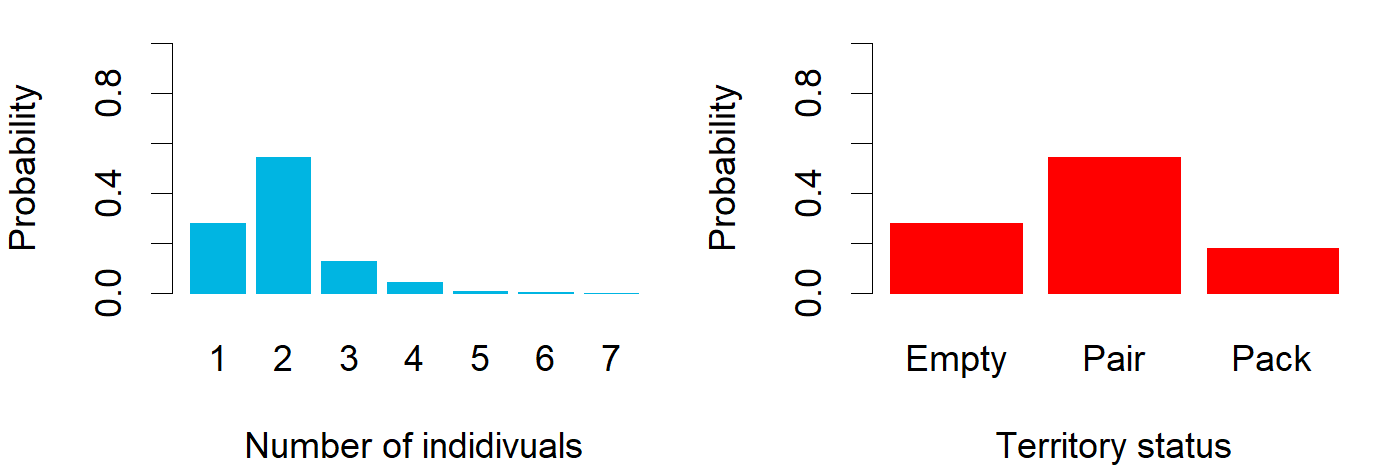

### March_ENG_Peurajärvi.png

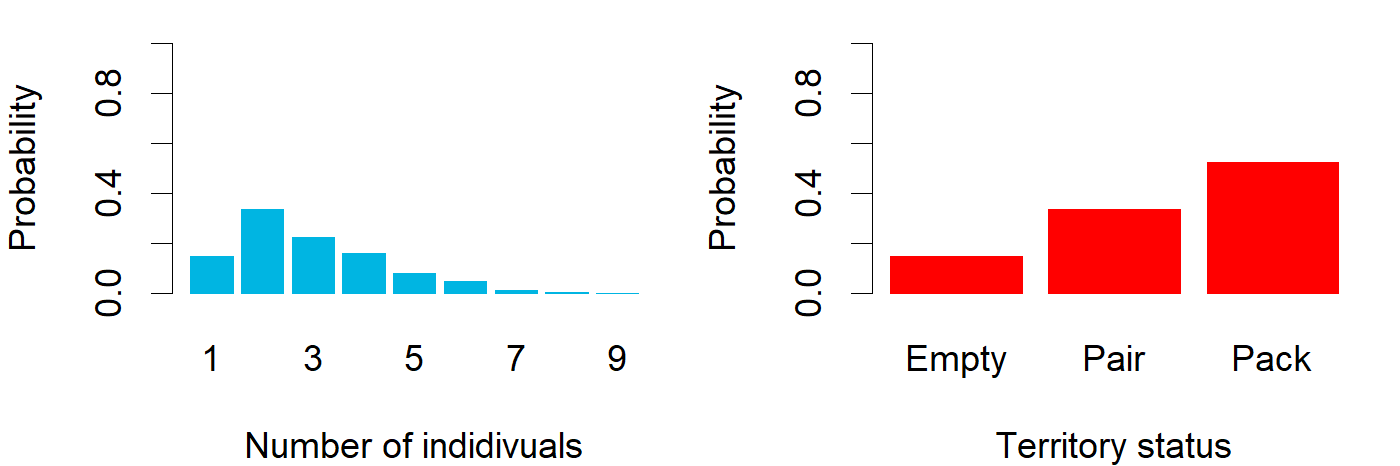

### March_ENG_Pirttivaara.png

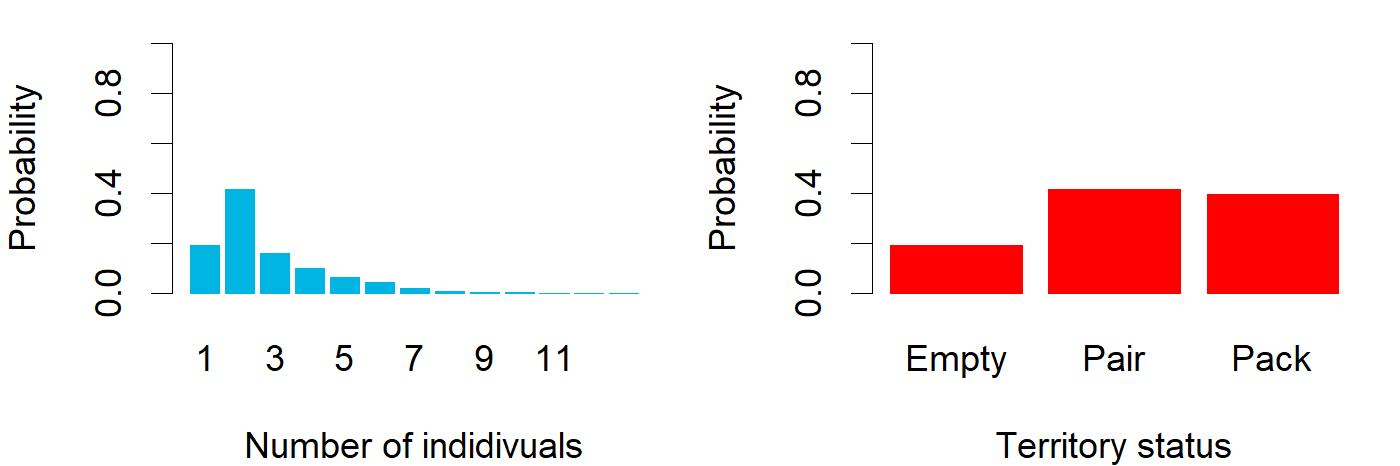

### March_ENG_Pulkkila.png

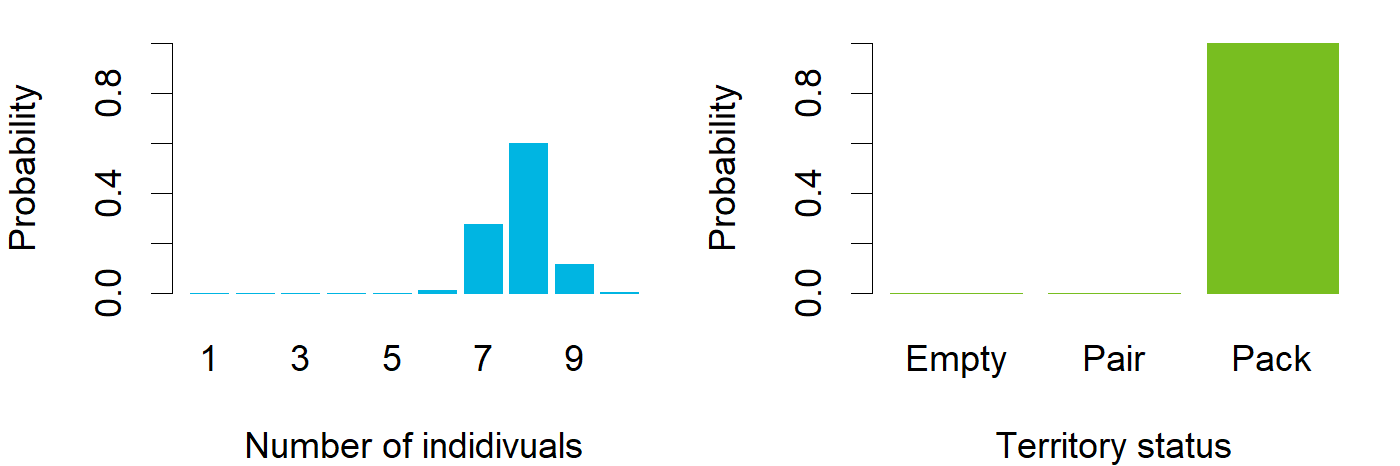

### March_ENG_Punkalaidun.png

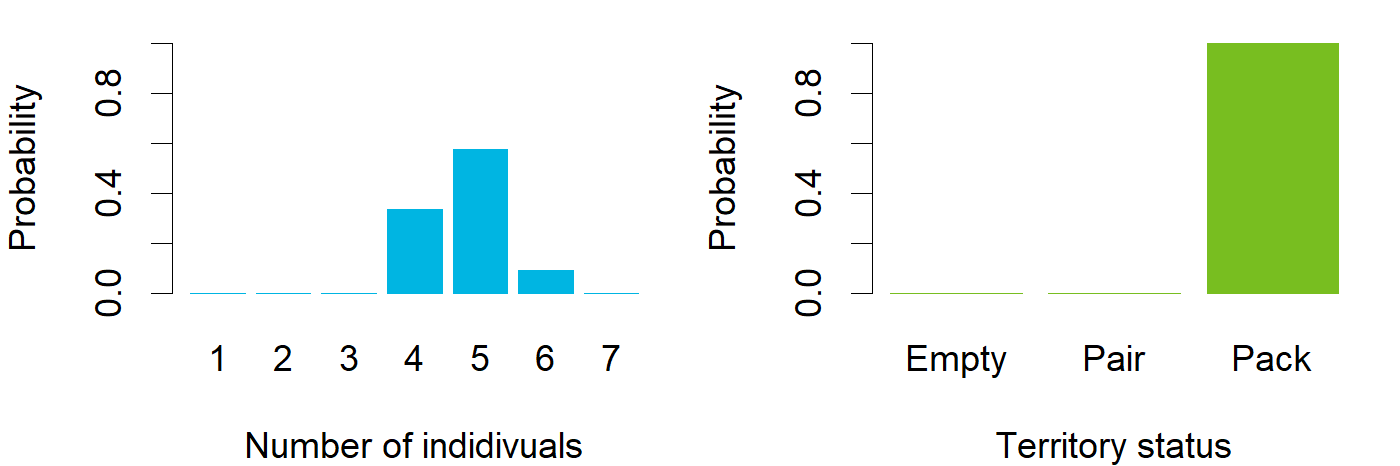

### March_ENG_Pyhäjoki.png

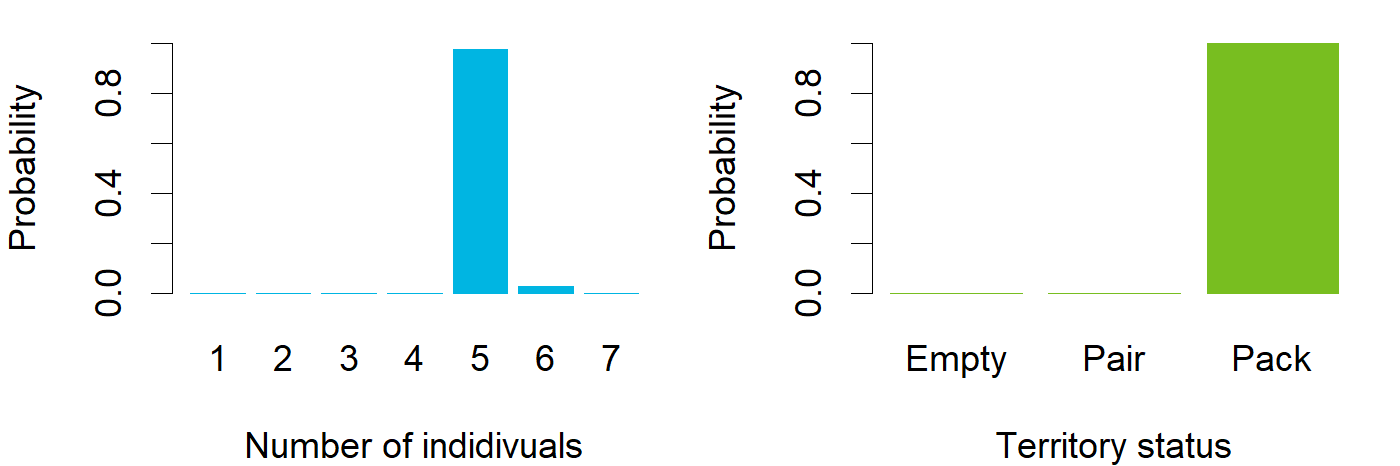

### March_ENG_Pöytyä.png

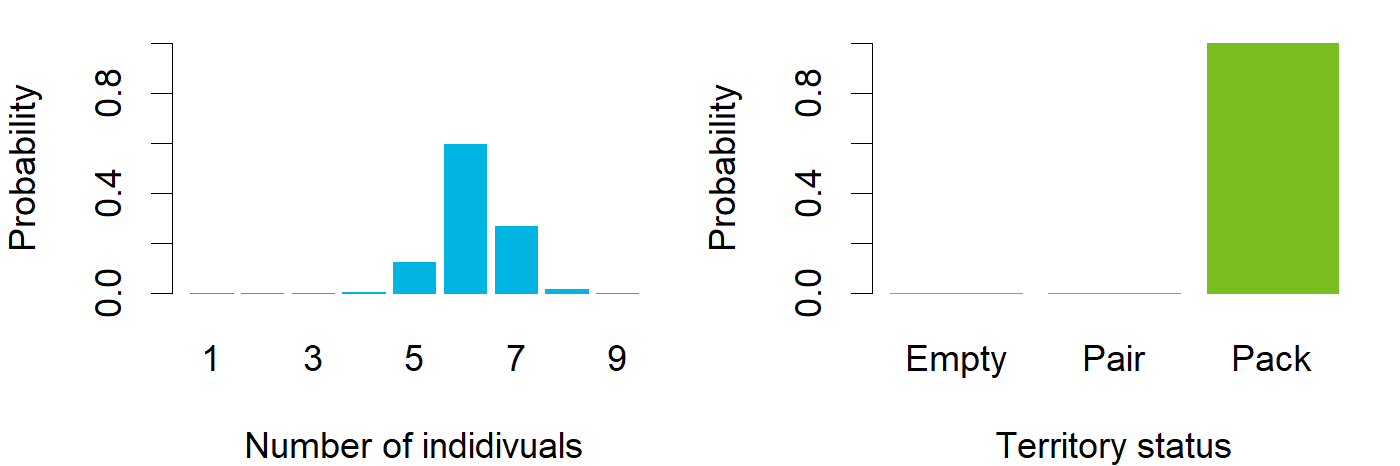

### March_ENG_Raasepori.png

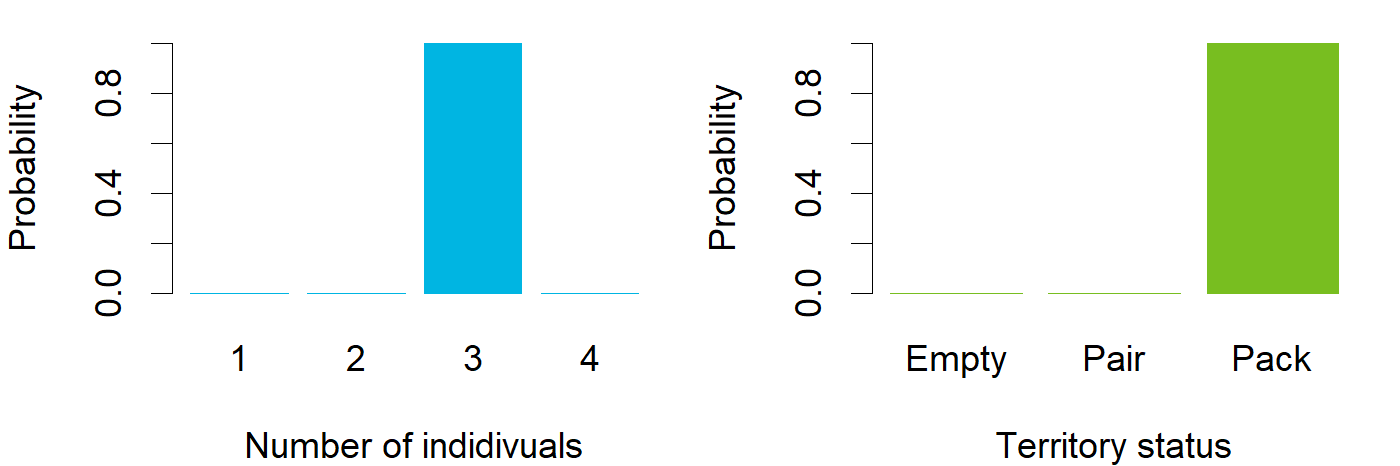

### March_ENG_Rantsila.png

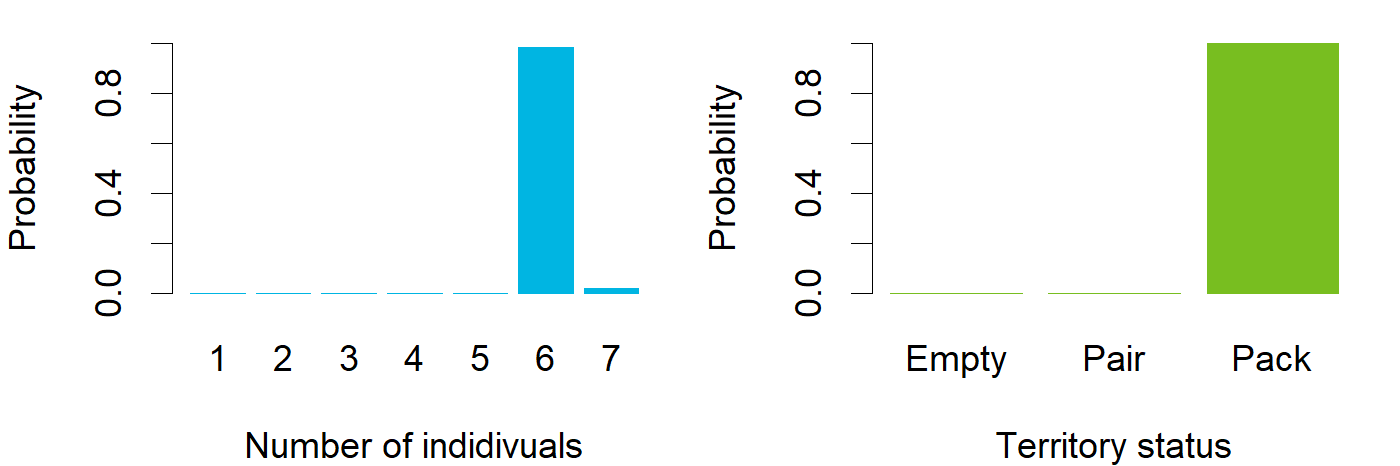

### March_ENG_Revonlahti.png

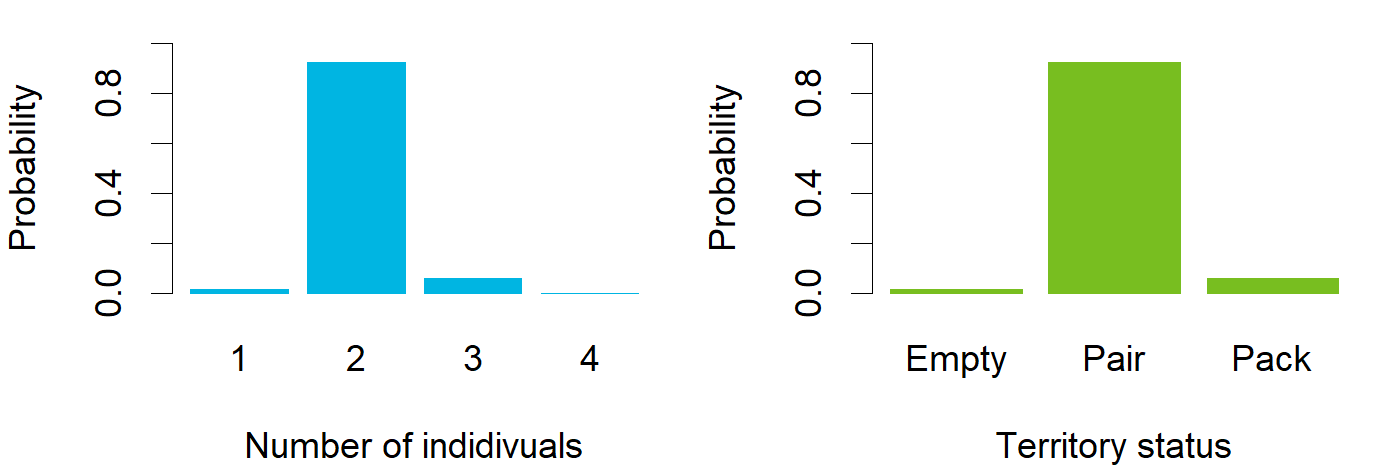

### March_ENG_Rytky.png

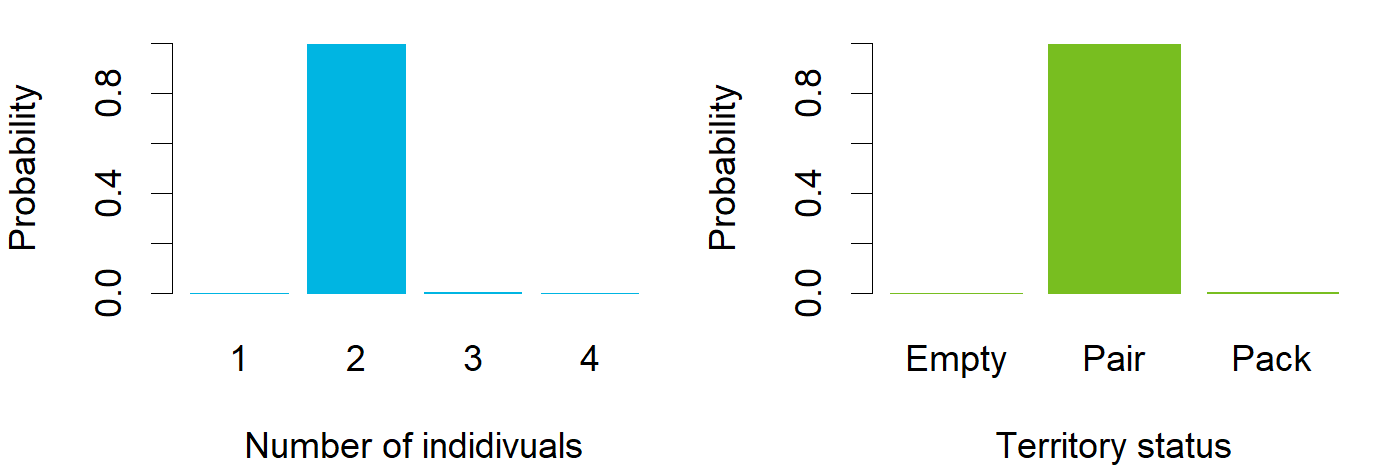

### March_ENG_Saramo.png

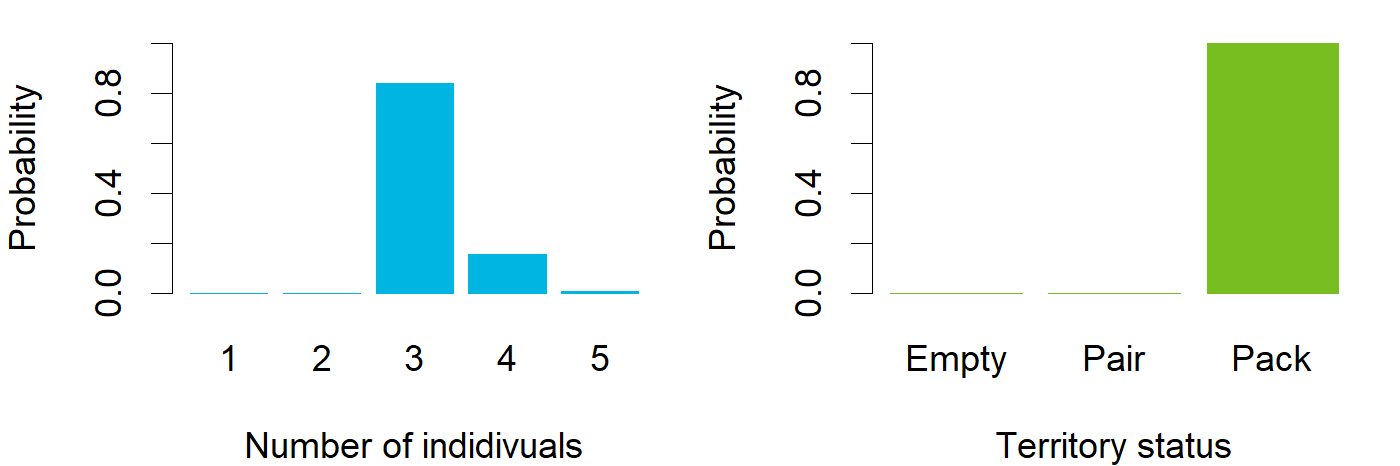

### March_ENG_Saunajärvi.png

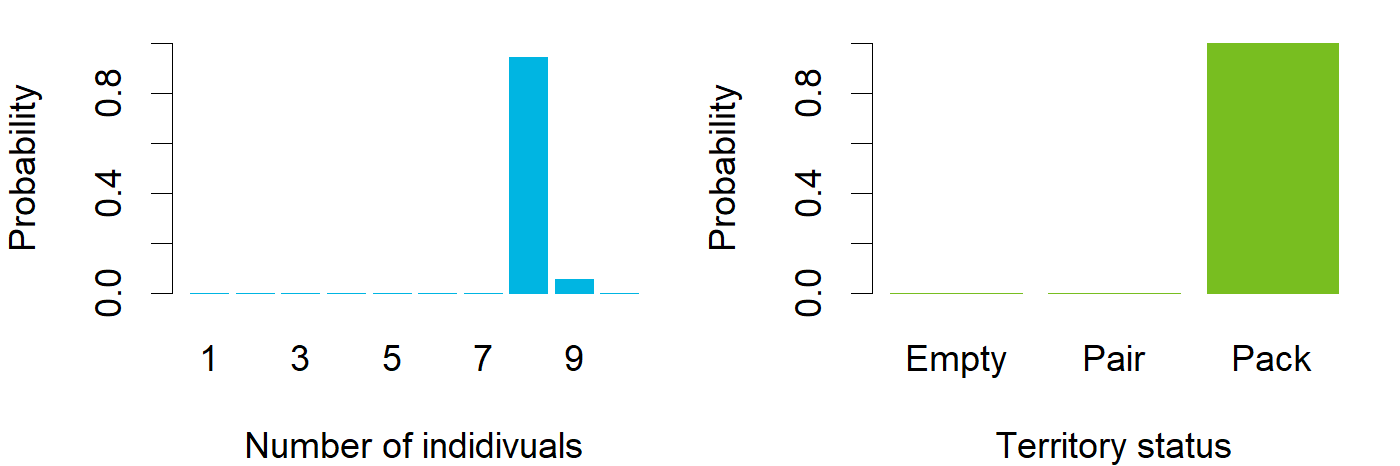

### March_ENG_Snappertuna.png

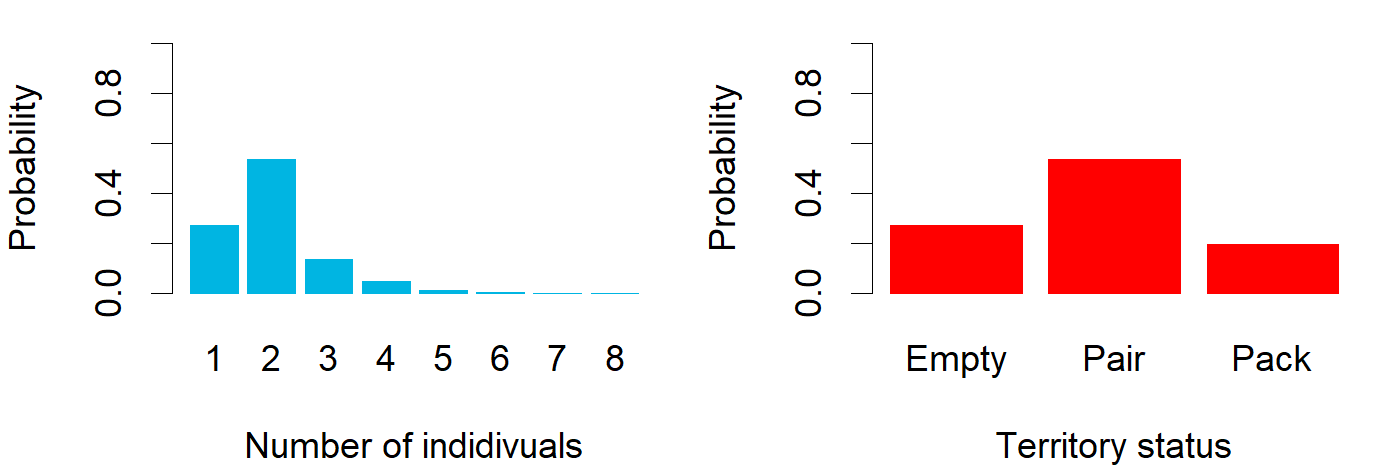

### March_ENG_Somerniemi.png

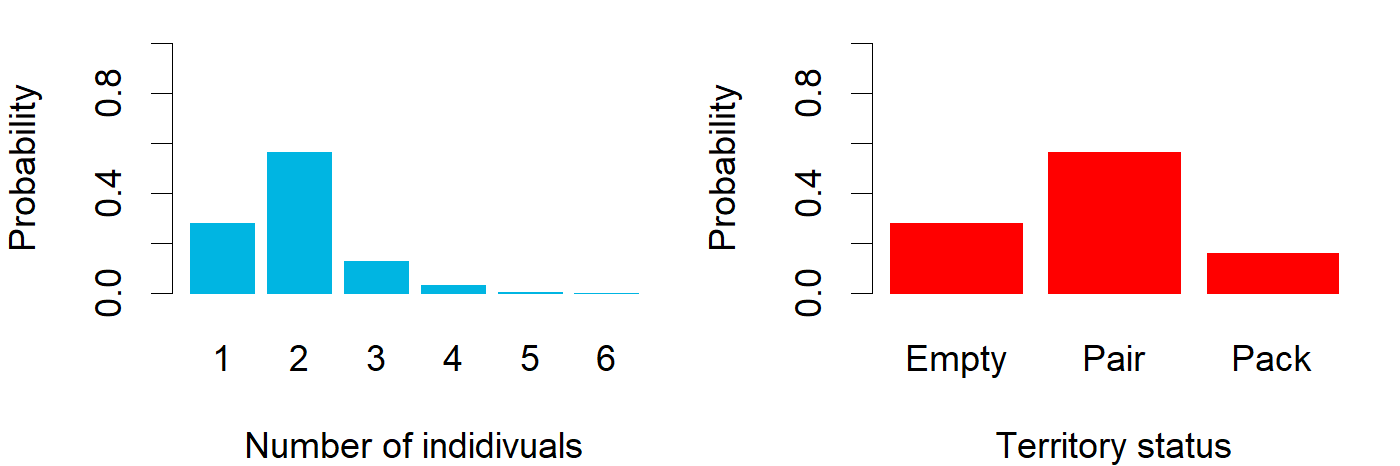

### March_ENG_survival.png

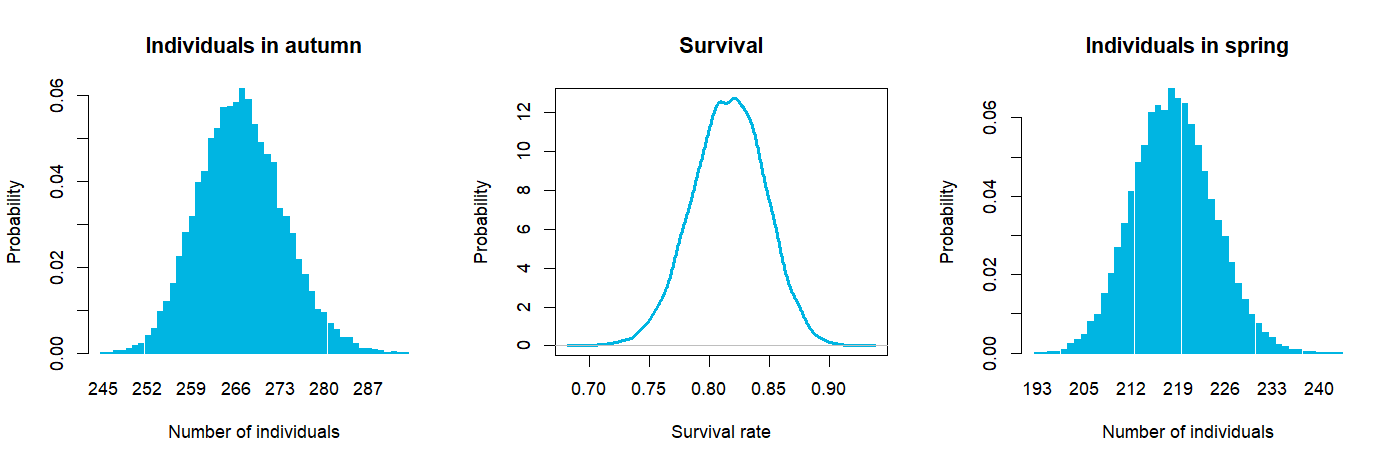

### March_ENG_territorytotals.png

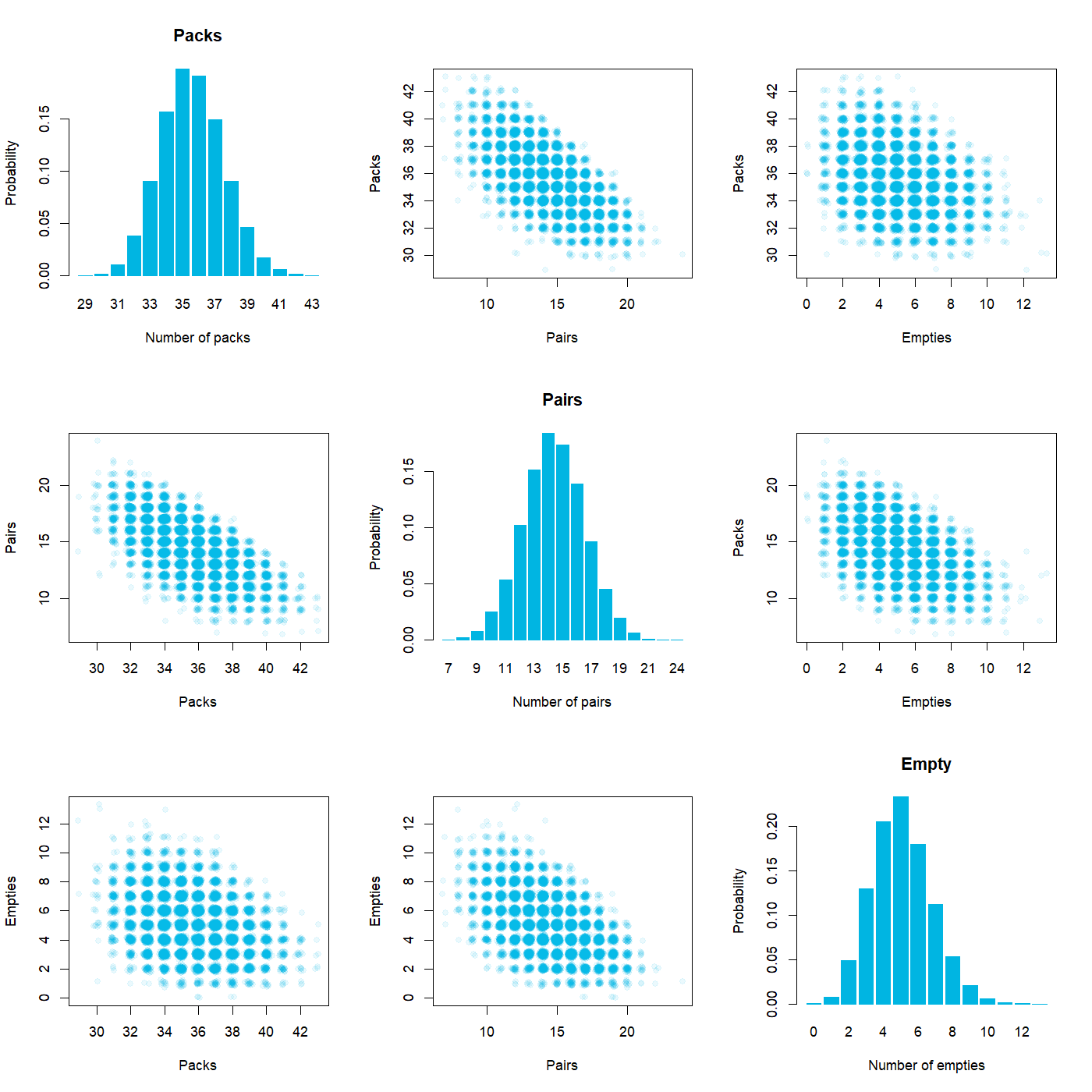

### March_ENG_Toholampi.png

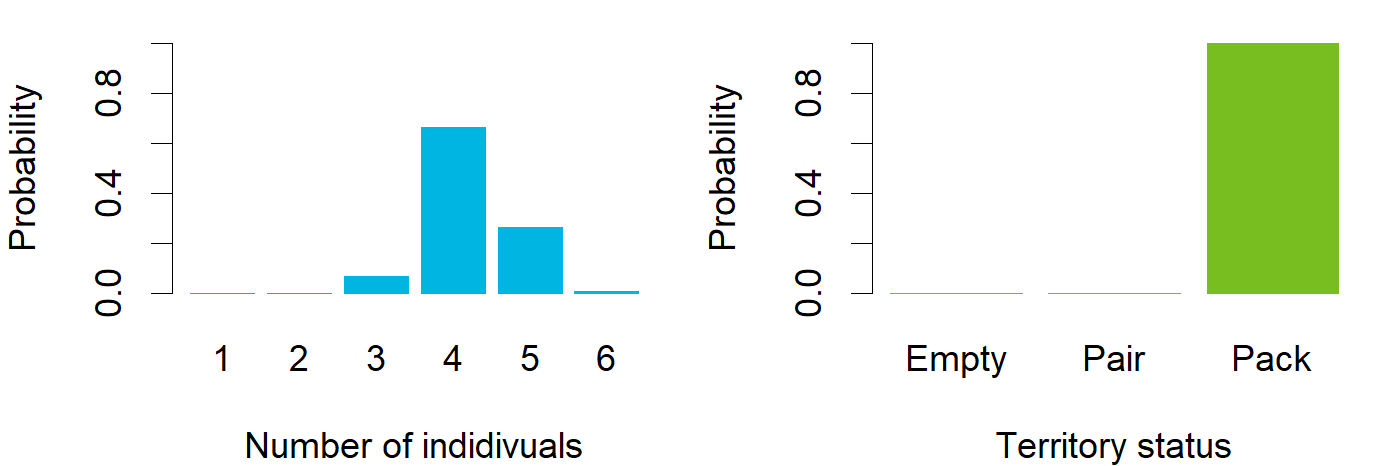

### March_ENG_Toijala.png

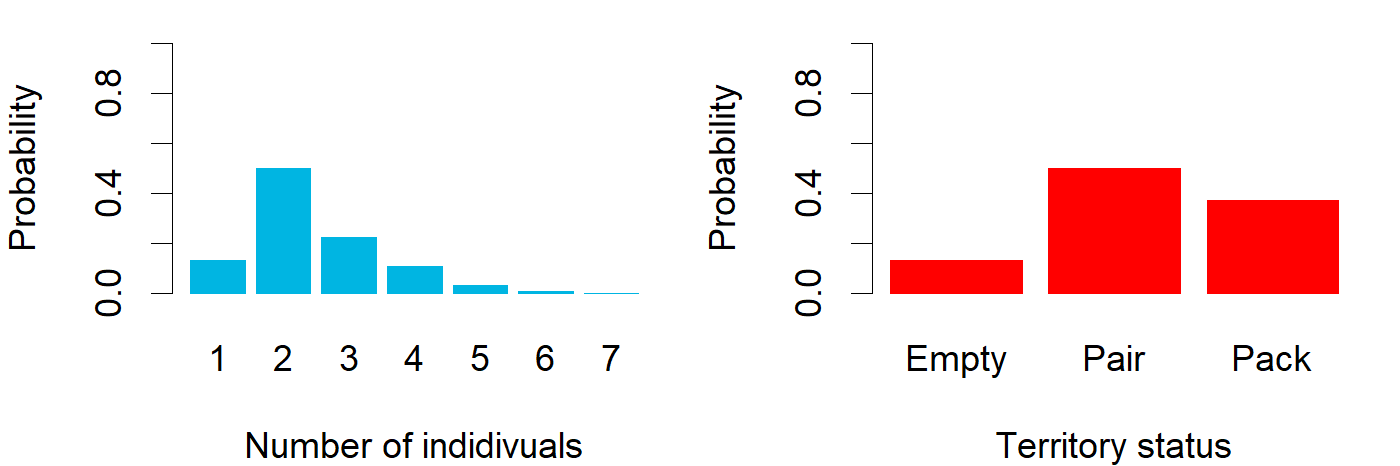

### March_ENG_Utajärvi.png

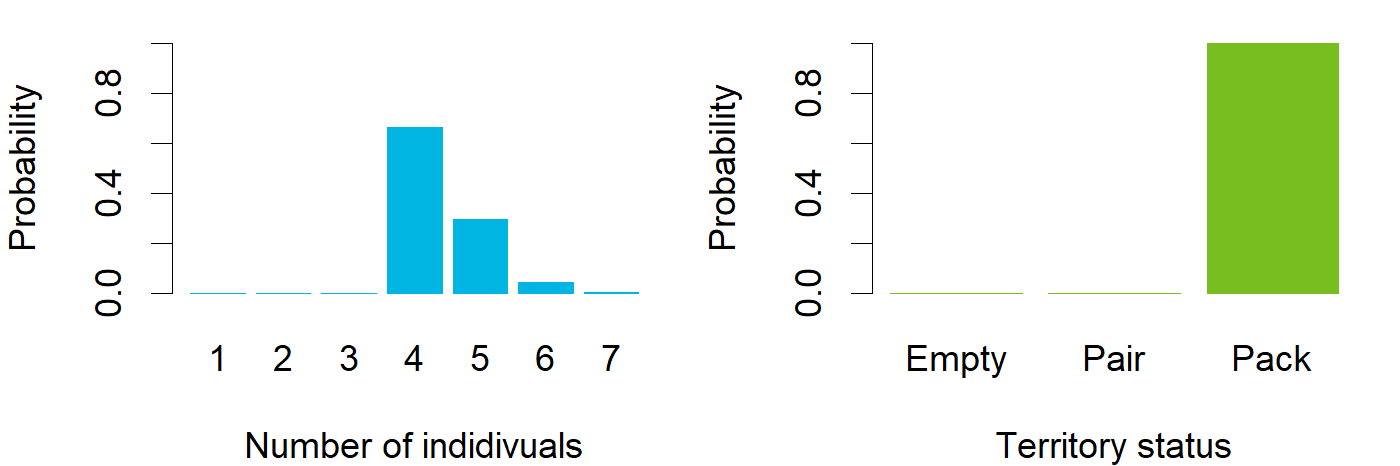

### March_ENG_Uura.png

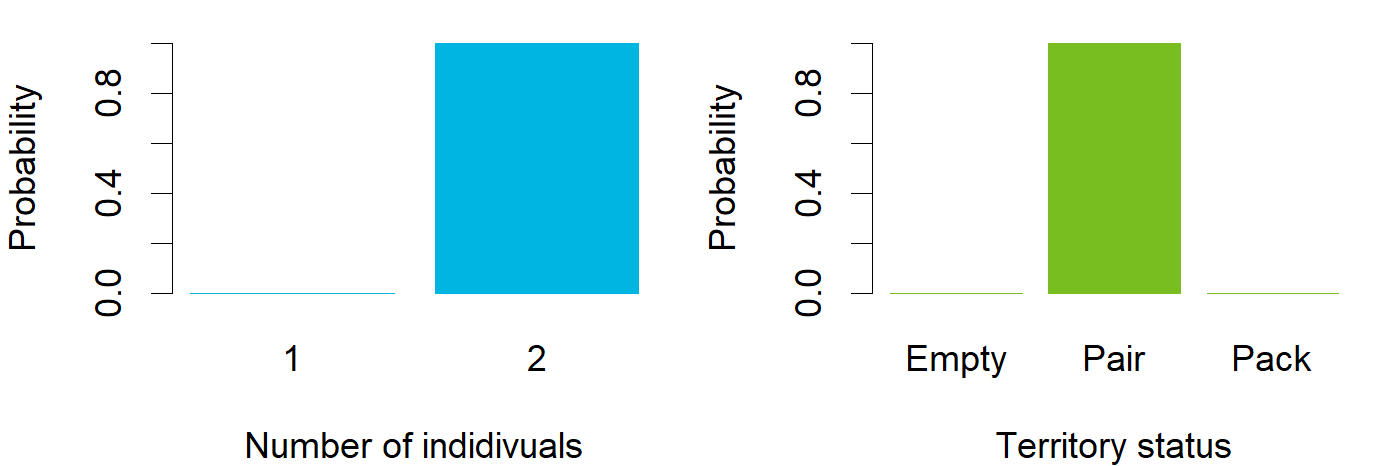

### March_ENG_Vartius.png

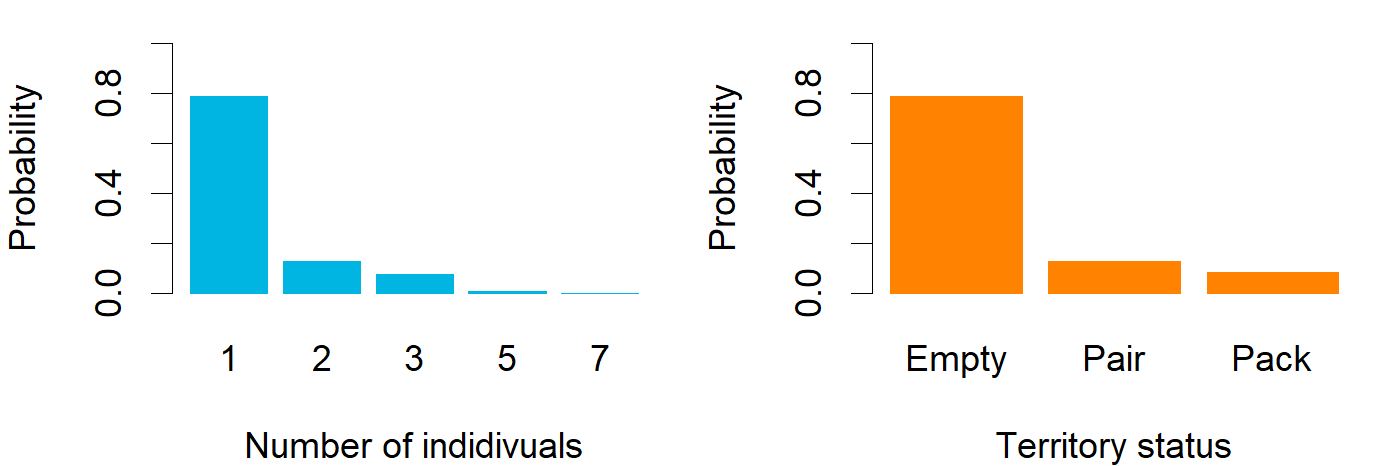

### March_ENG_Vuonisjärvi.png

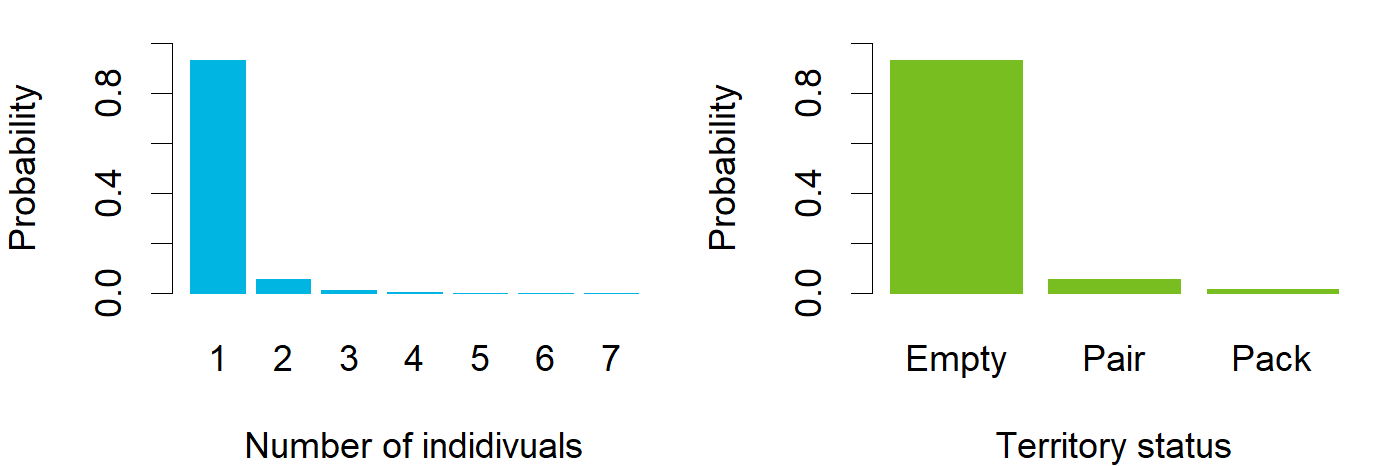

### March_ENG_Värtsilä.png

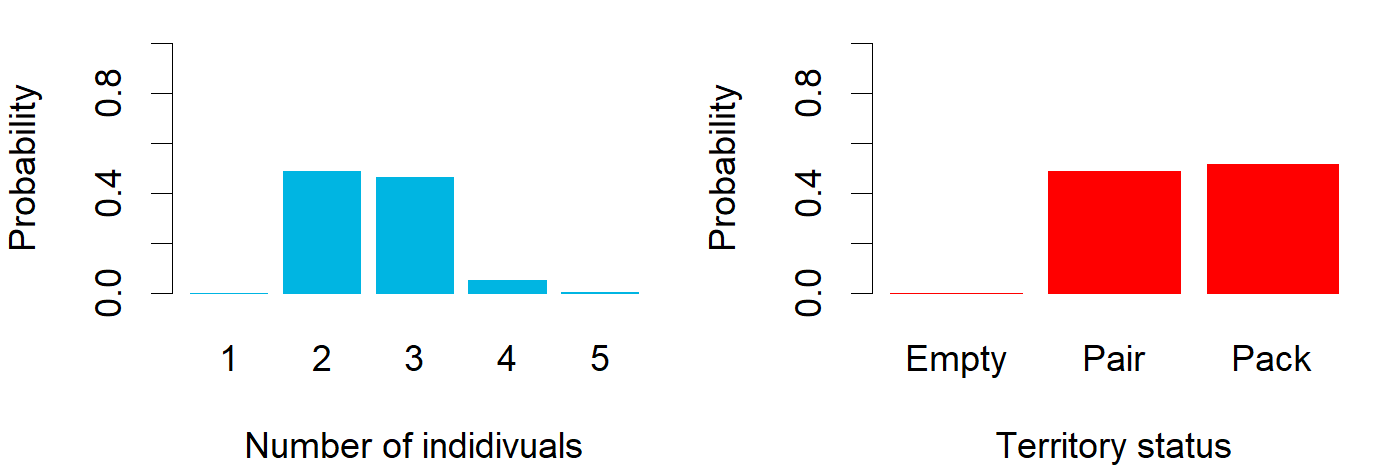

### Nov_ENG_Eurajoki.png

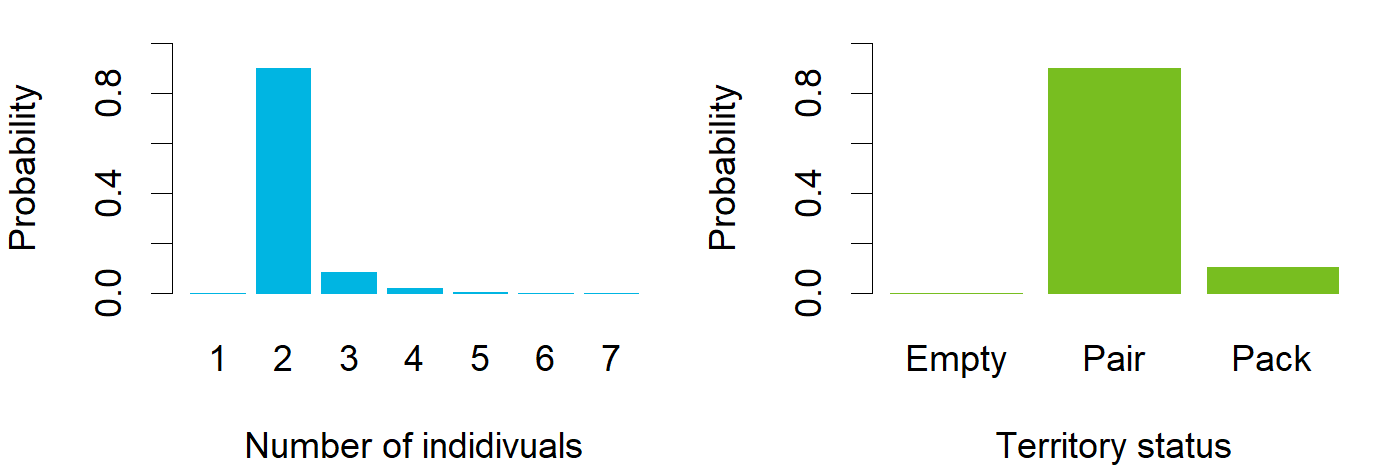

### Nov_ENG_Haapajärvi.png

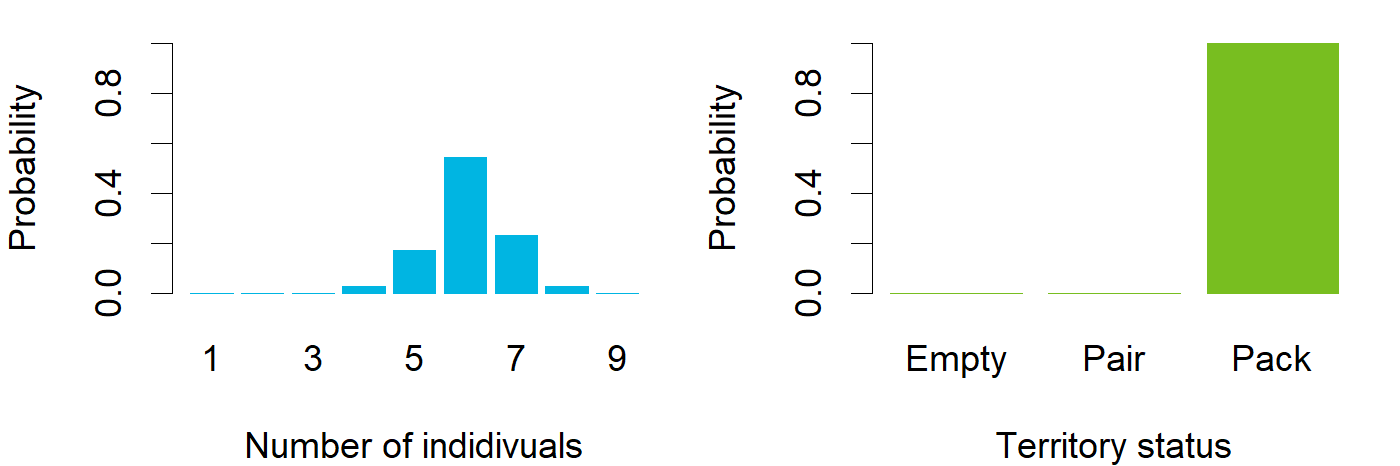

### Nov_ENG_Halivaara.png

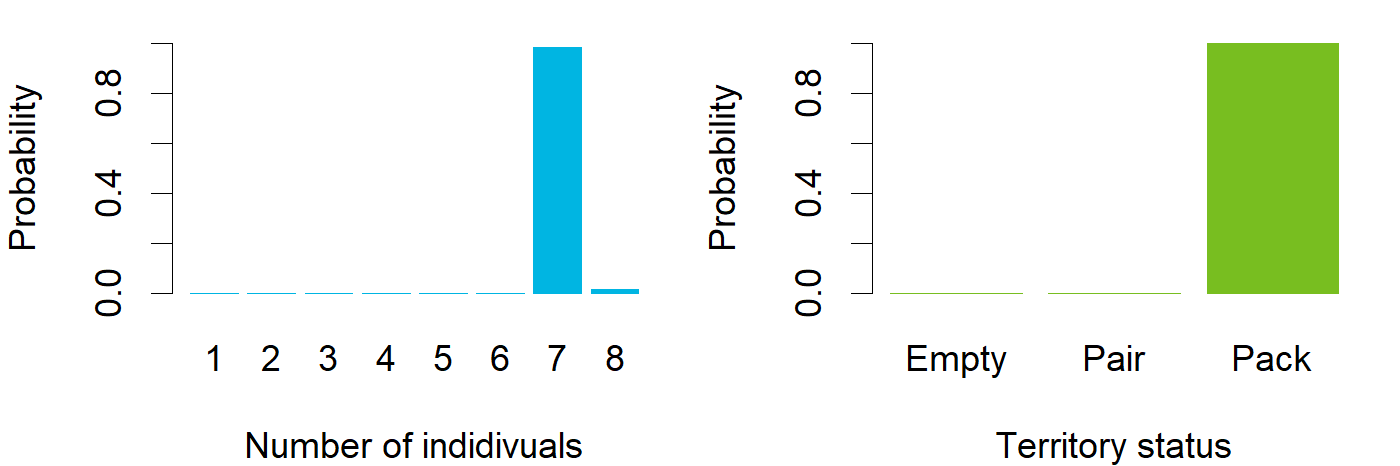

### Nov_ENG_Honkajoki.png

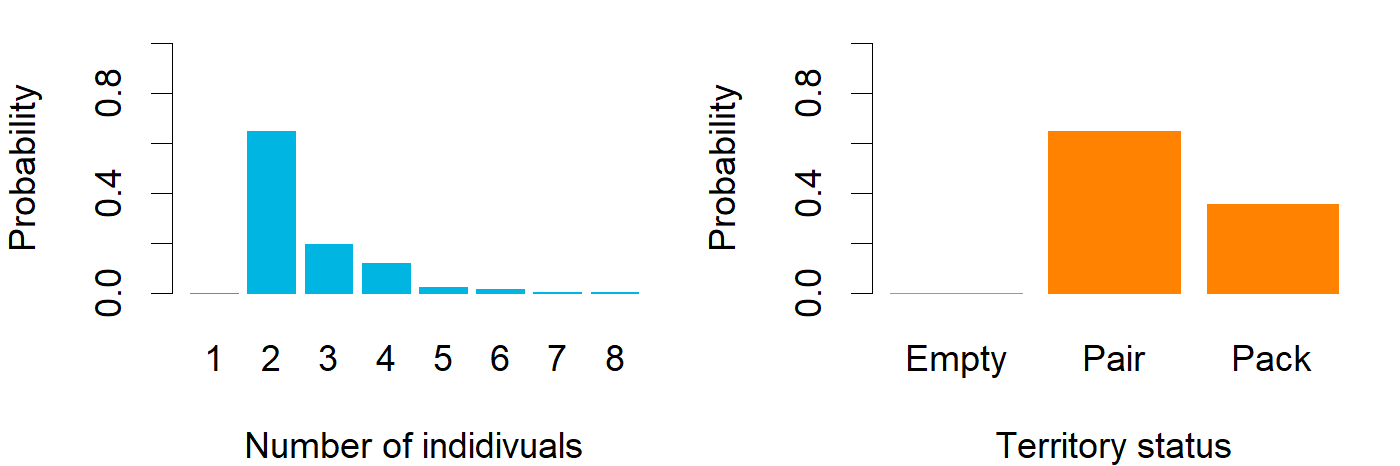

### Nov_ENG_Höljäkkä.png

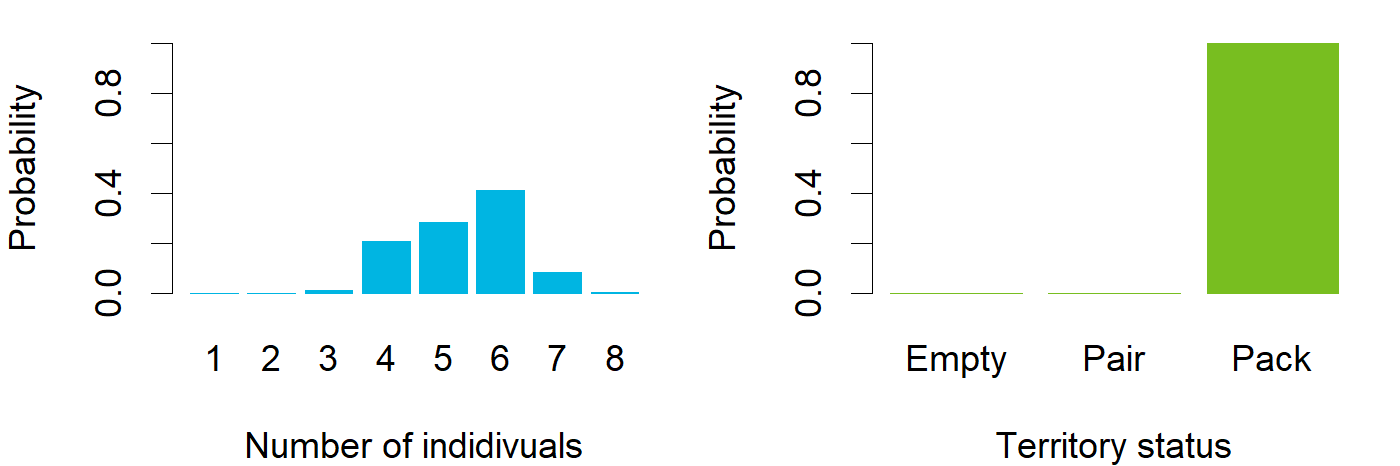

### Nov_ENG_Ihode.png

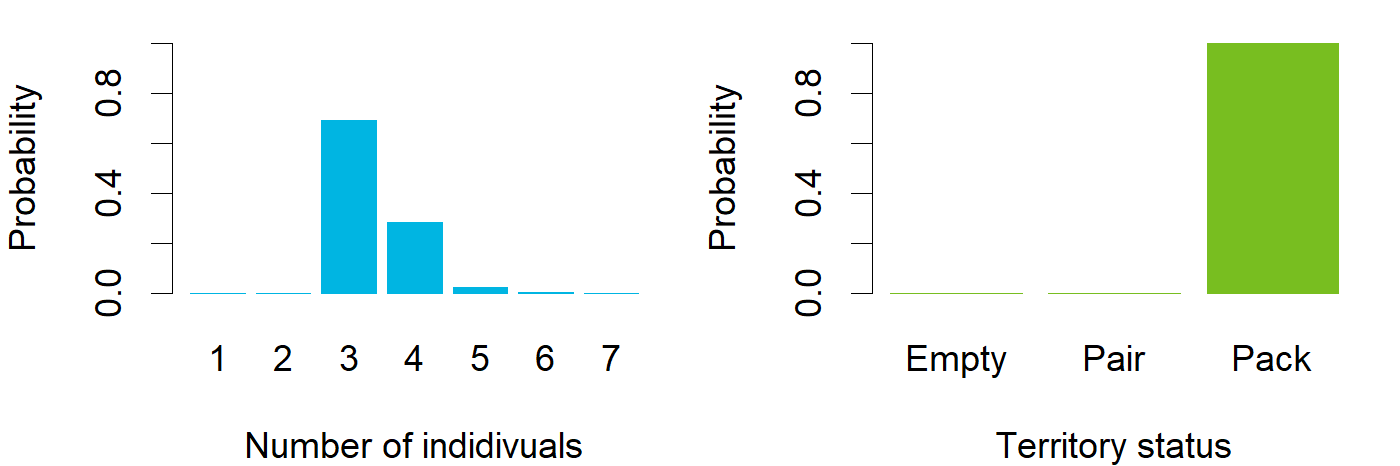

### Nov_ENG_Inari.png

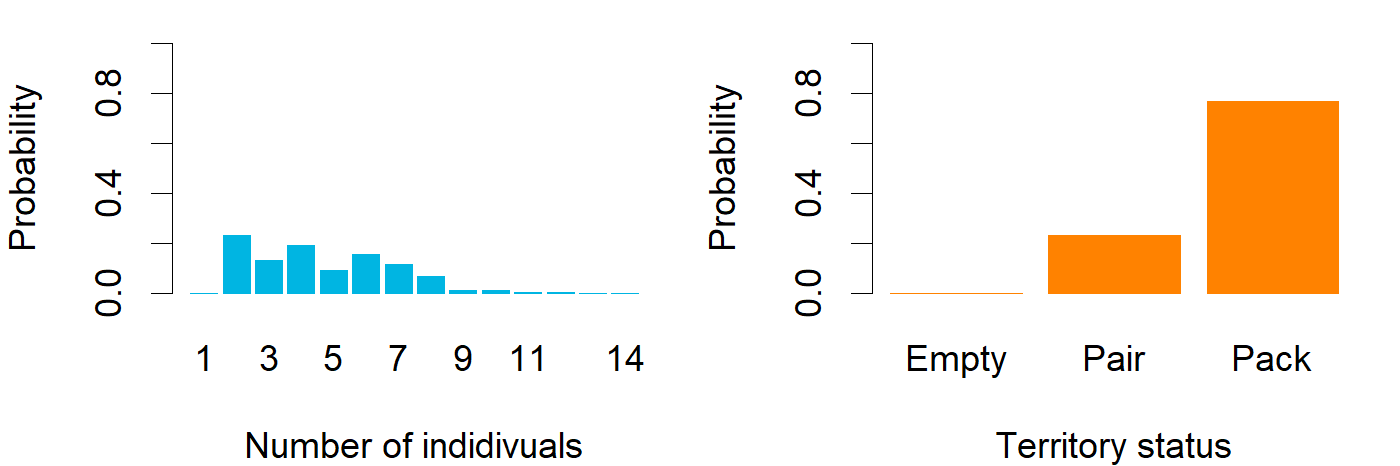

### Nov_ENG_Isojoki.png

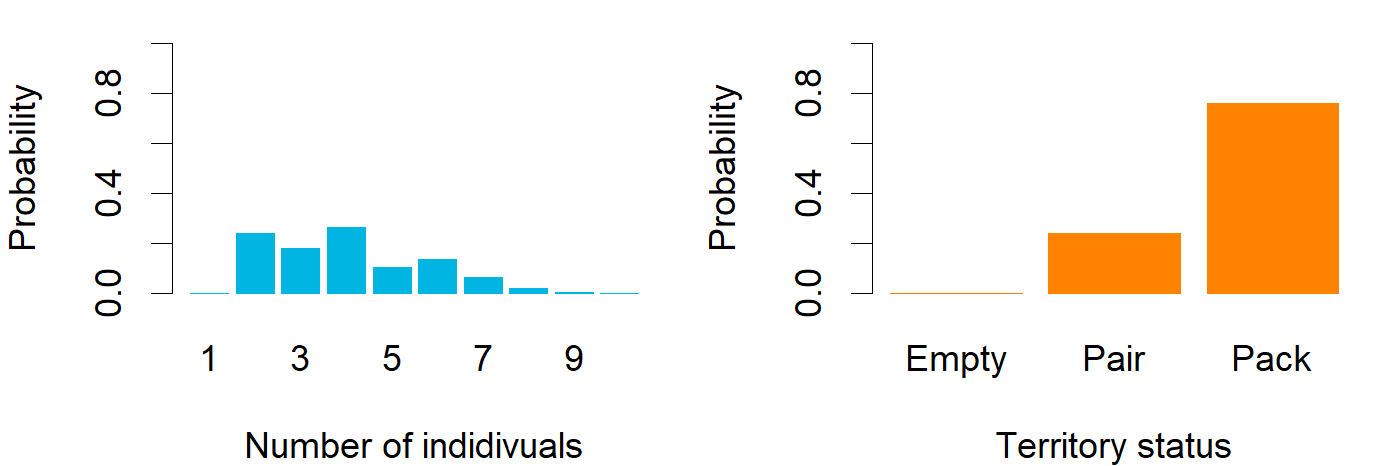

### Nov_ENG_Jeppo.png

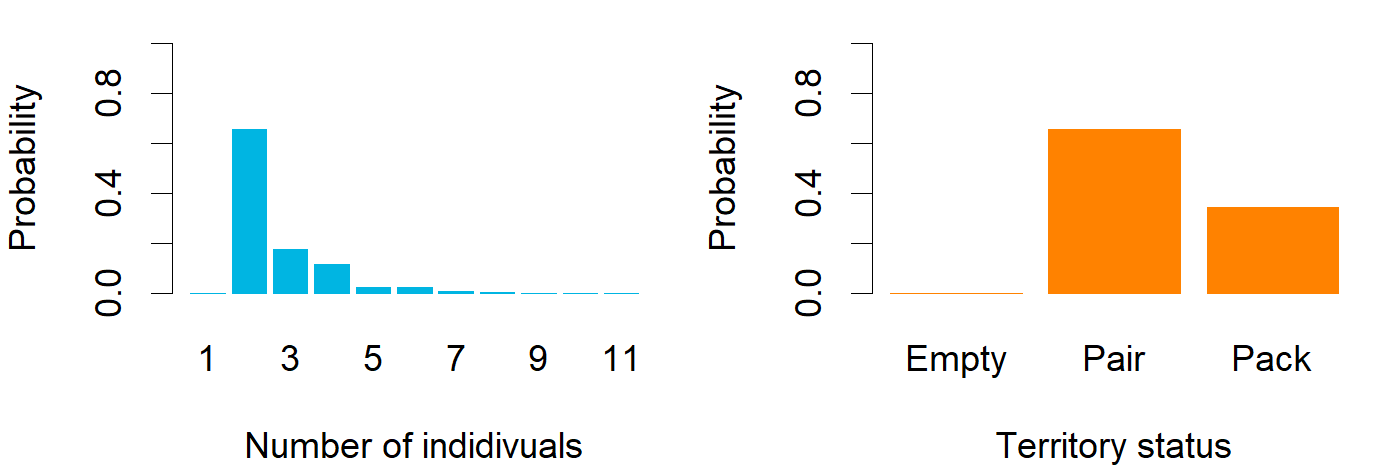

### Nov_ENG_Juntti.png

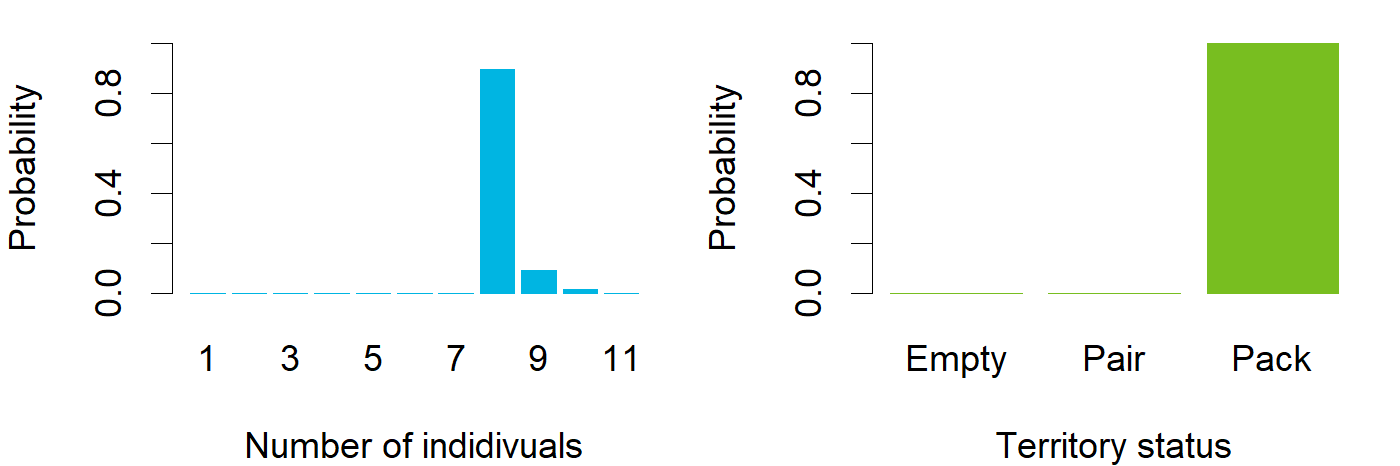

### Nov_ENG_Jurva.png

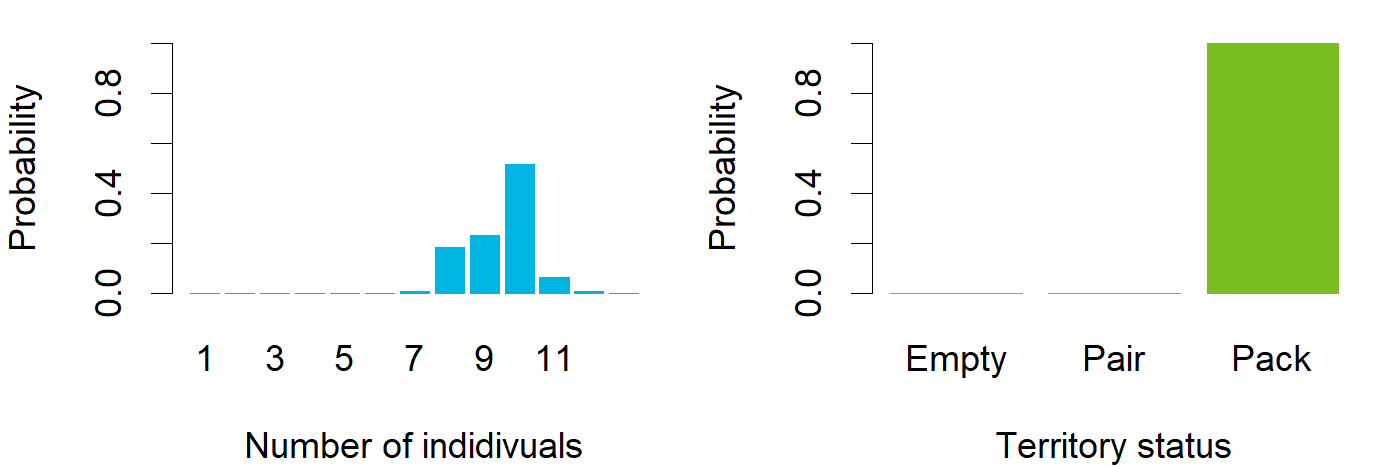

### Nov_ENG_Juupajoki.png

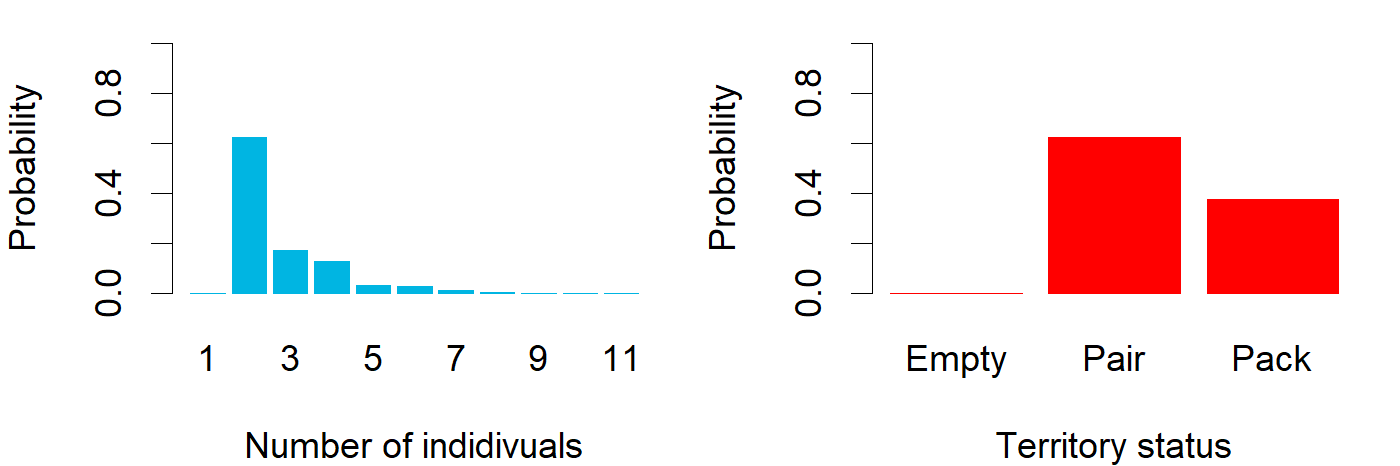

### Nov_ENG_Kaivola.png

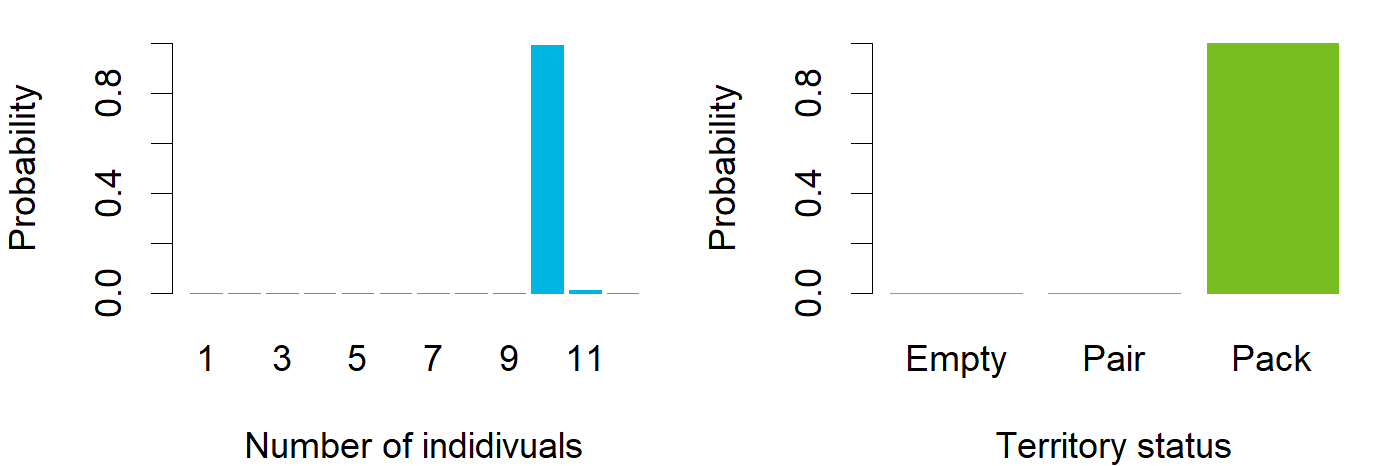

### Nov_ENG_Kallioluoma.png

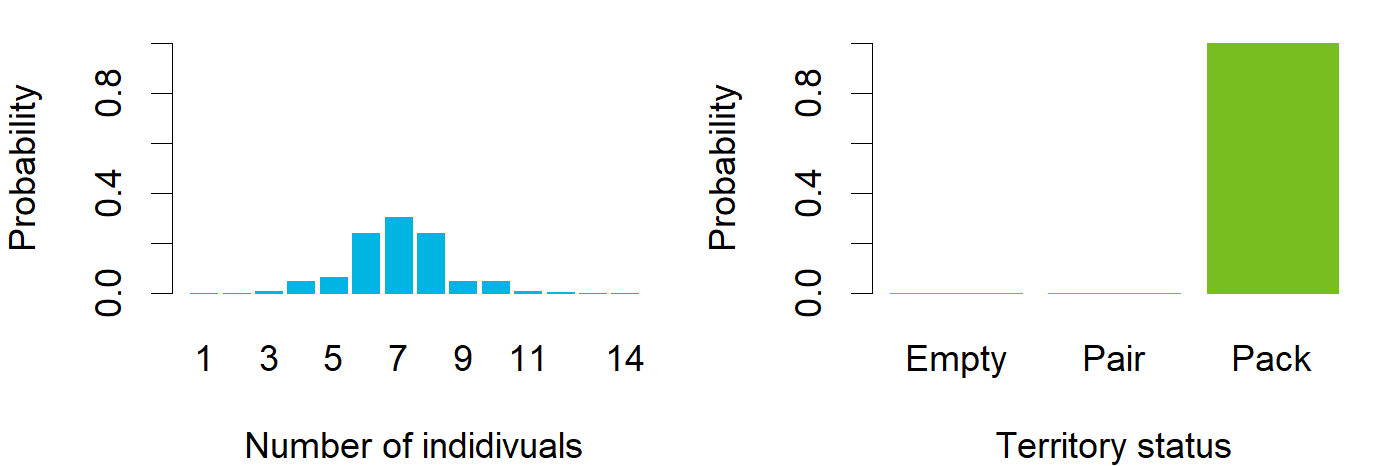

### Nov_ENG_Kankaanpää.png

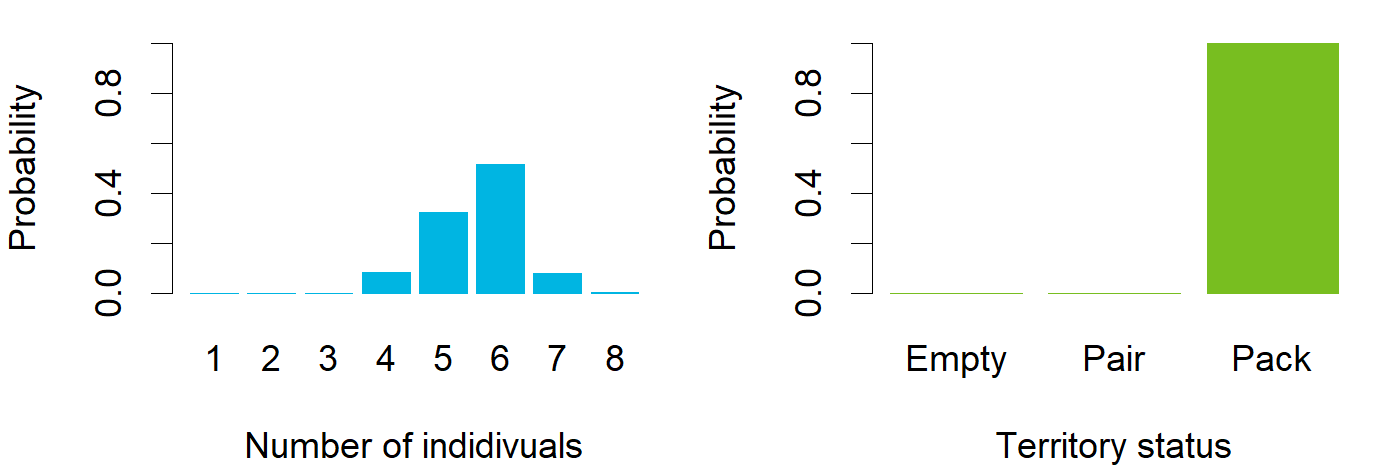

### Nov_ENG_Karvia.png

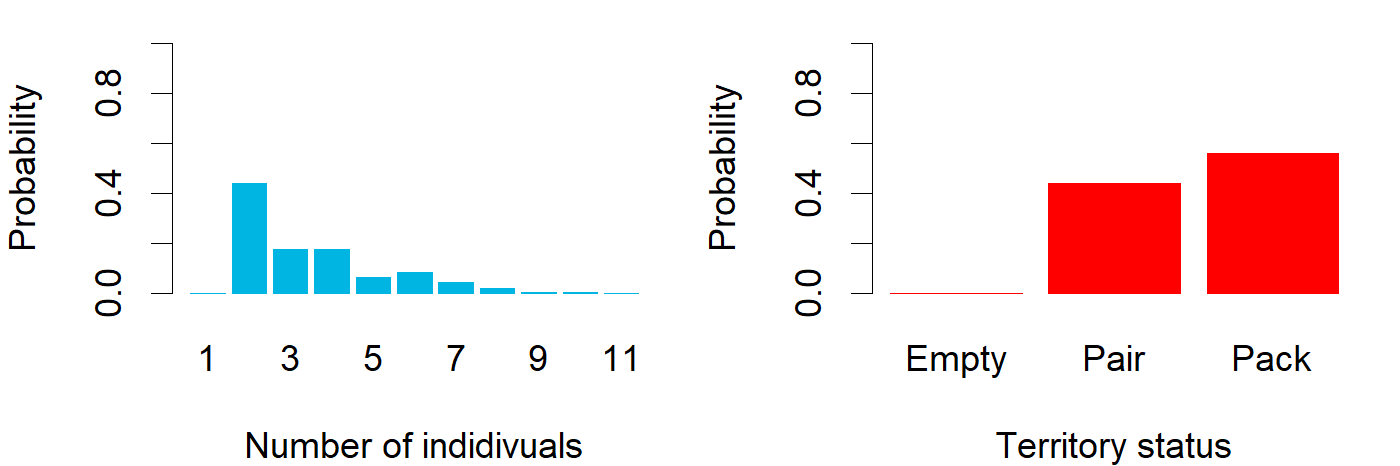

### Nov_ENG_Kelsimä.png

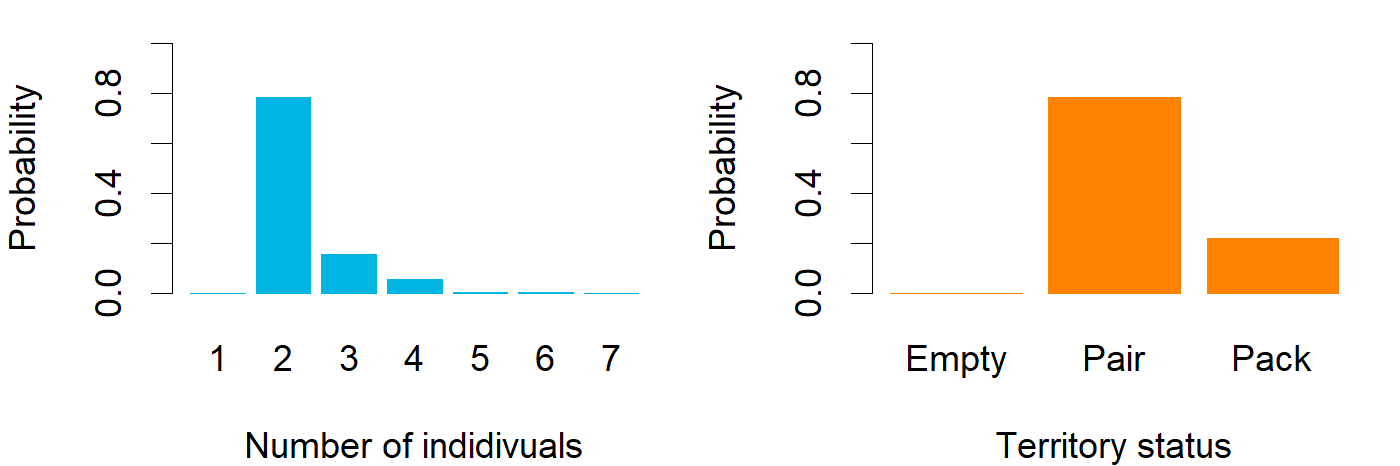

### Nov_ENG_Kemiö.png

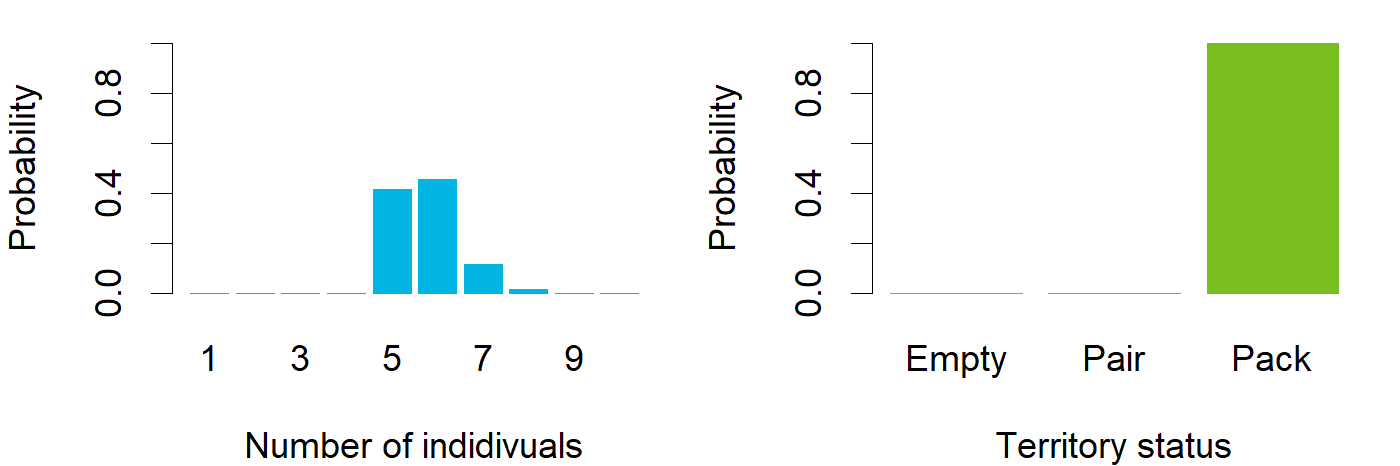

### Nov_ENG_Kiuruvesi.png

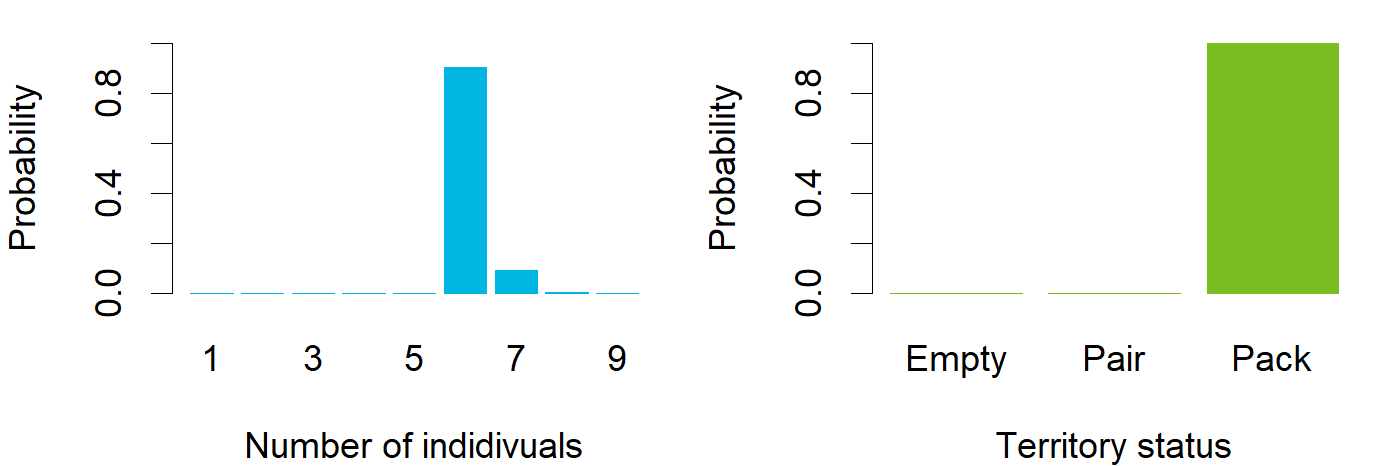

### Nov_ENG_Kivikiekki.png

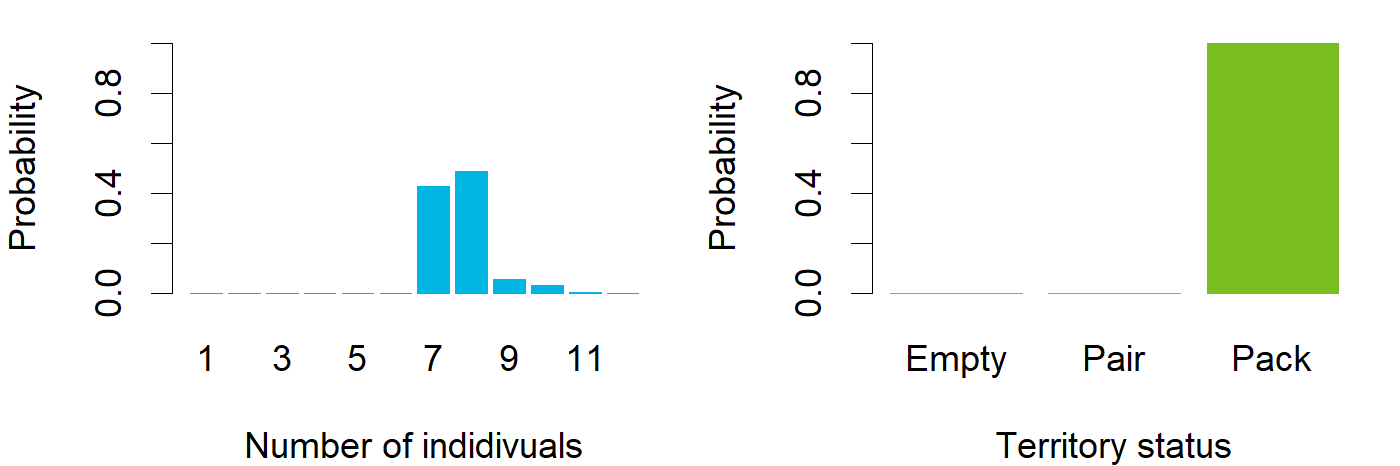

### Nov_ENG_Kivivaara.png

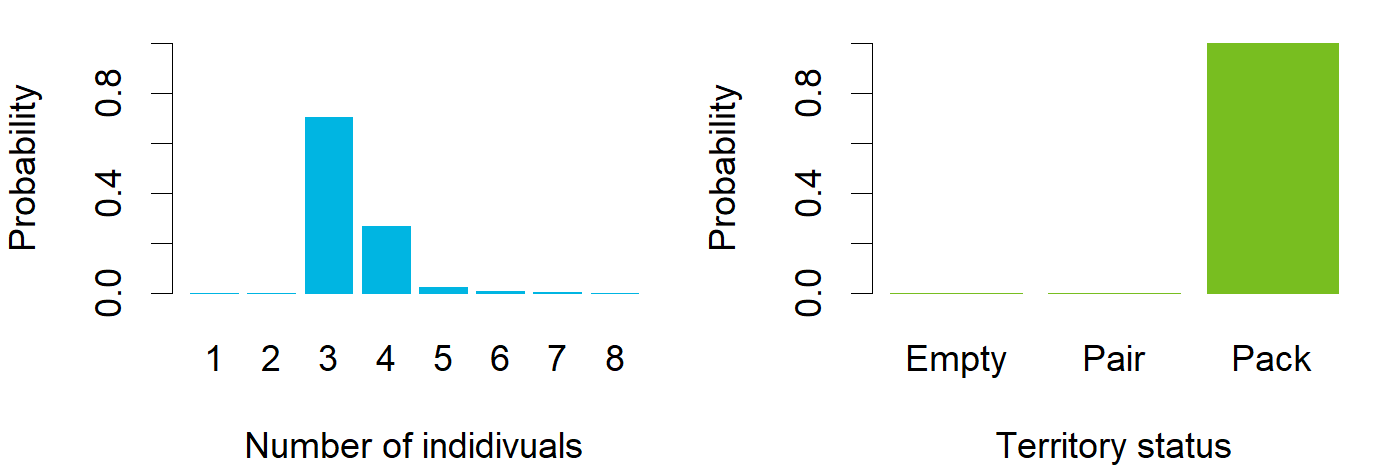

### Nov_ENG_Kovero.png

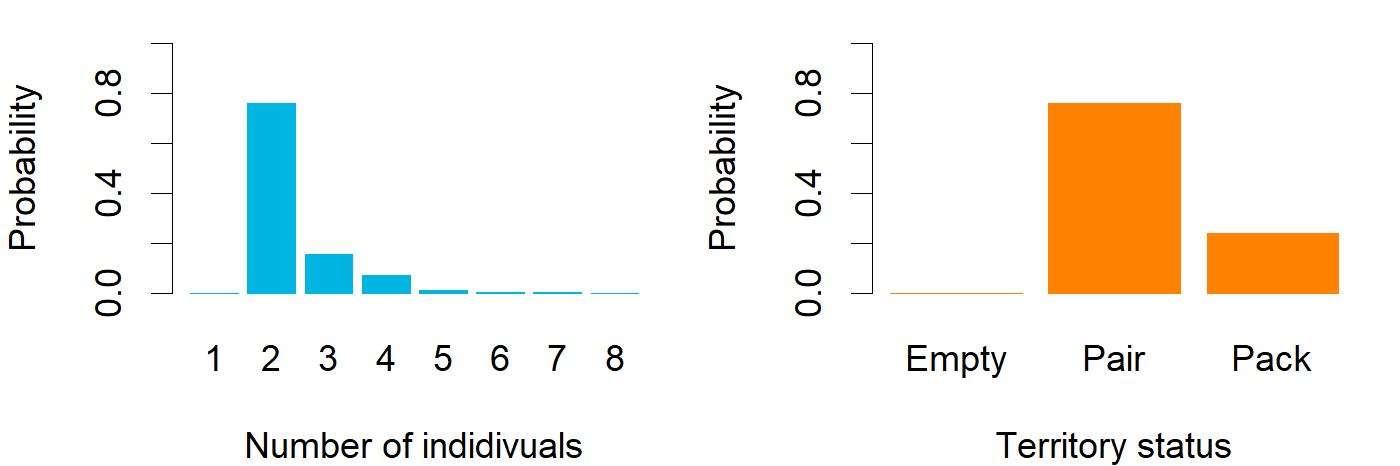

### Nov_ENG_Köyliö.png

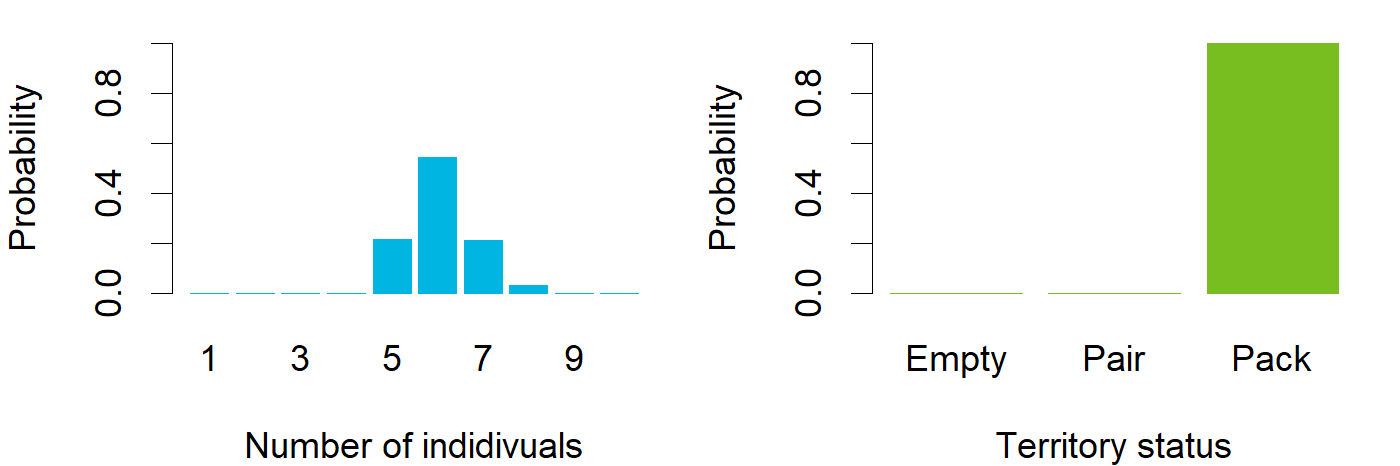

### Nov_ENG_Laakajärvi.png

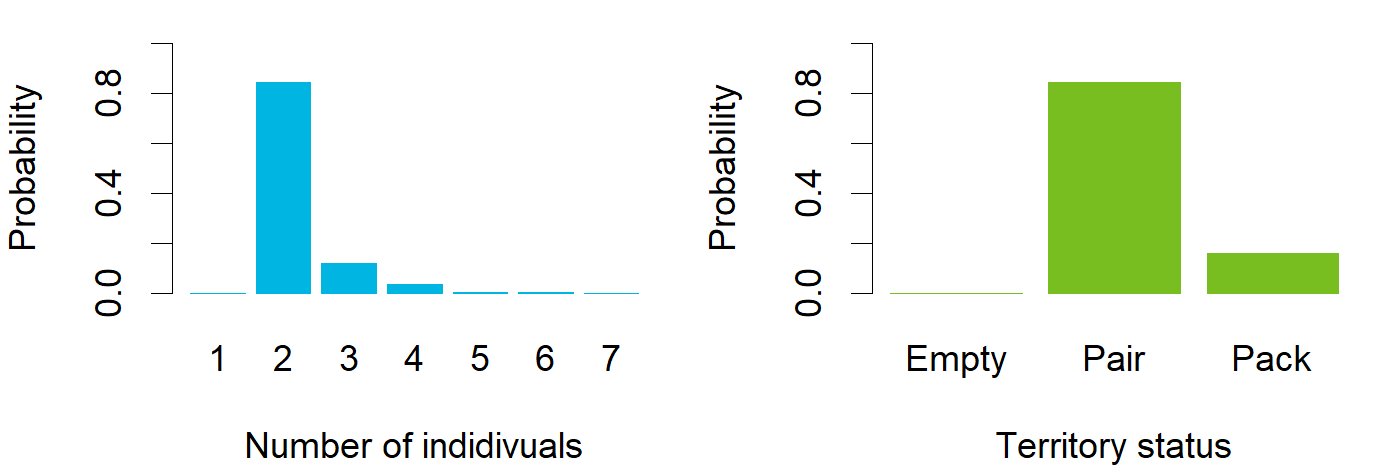

### Nov_ENG_Laihia.png

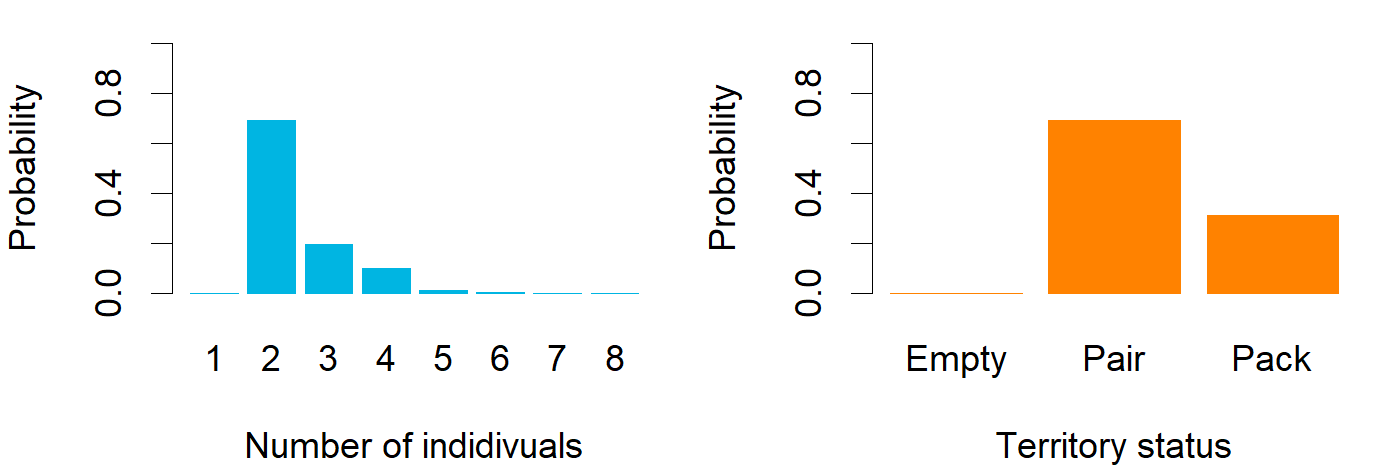

### Nov_ENG_Lauhavuori.png

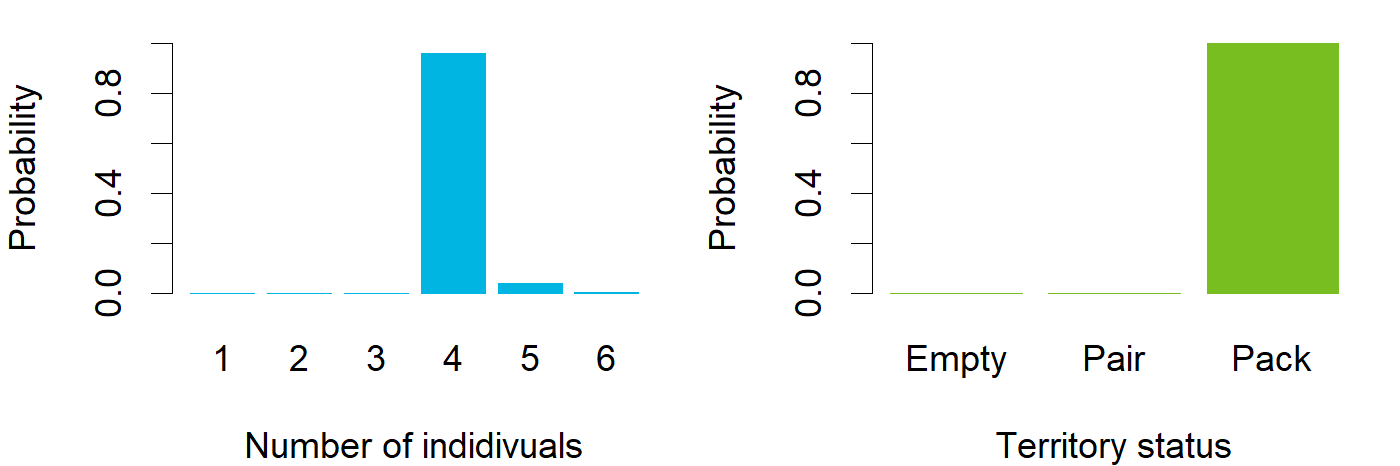

### Nov_ENG_Leppävirta.png

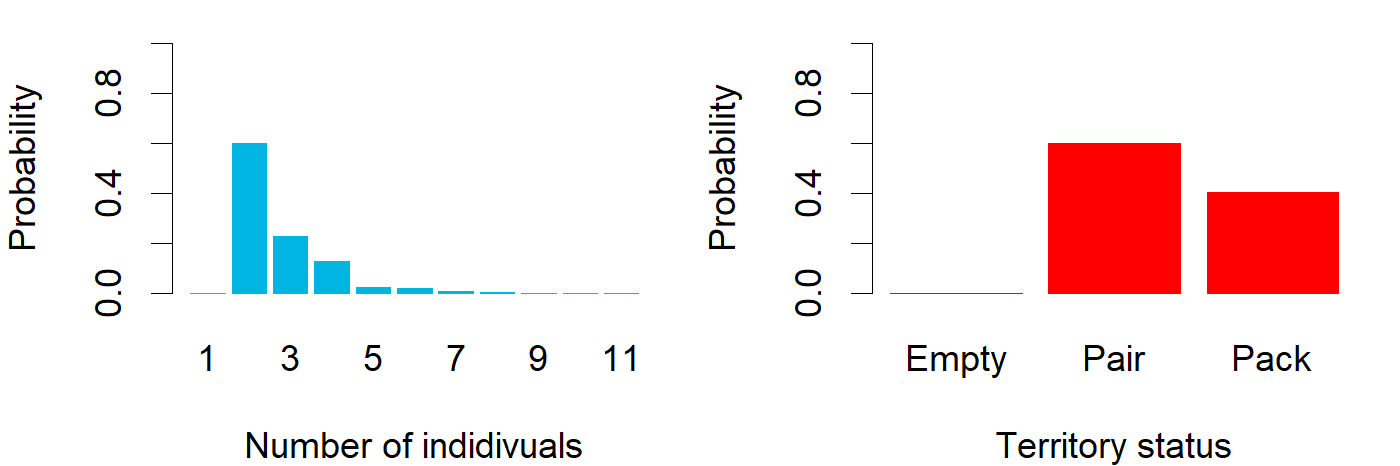

### Nov_ENG_Marttinen.png

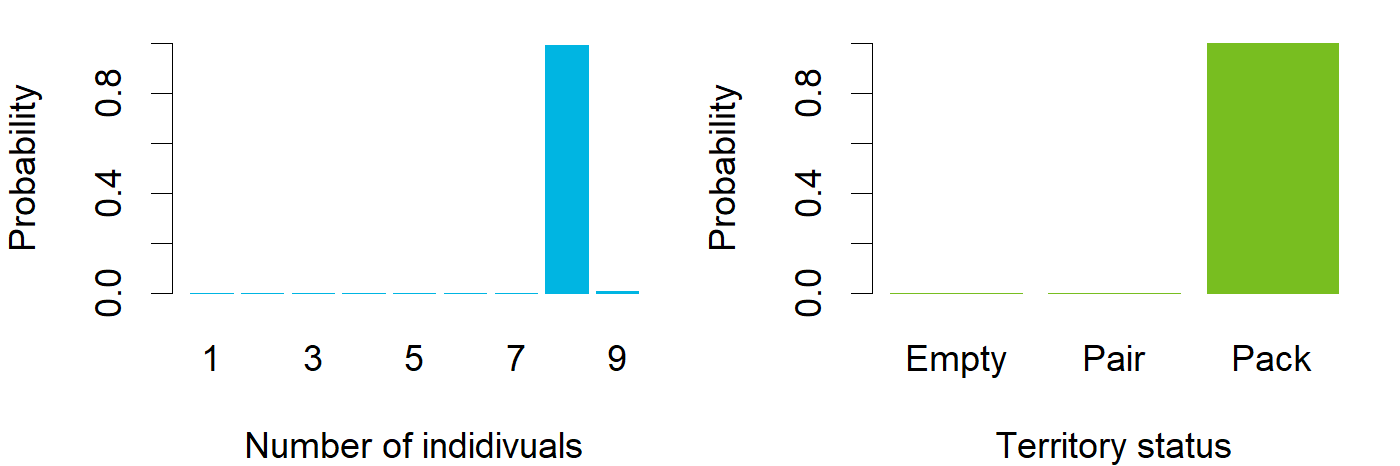

### Nov_ENG_Mynämäki.png

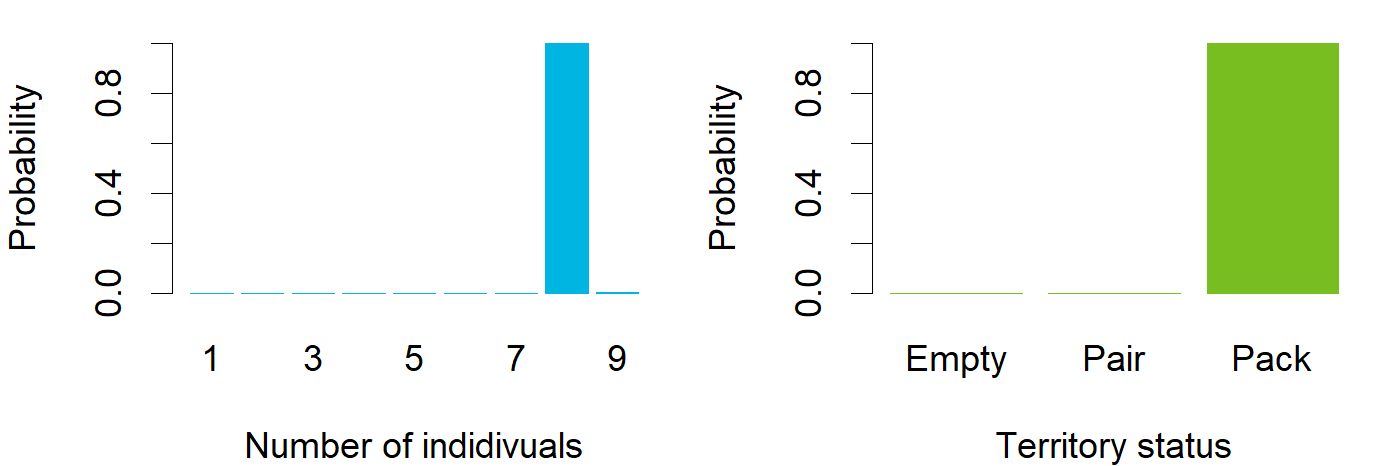

### Nov_ENG_Möhkö.png

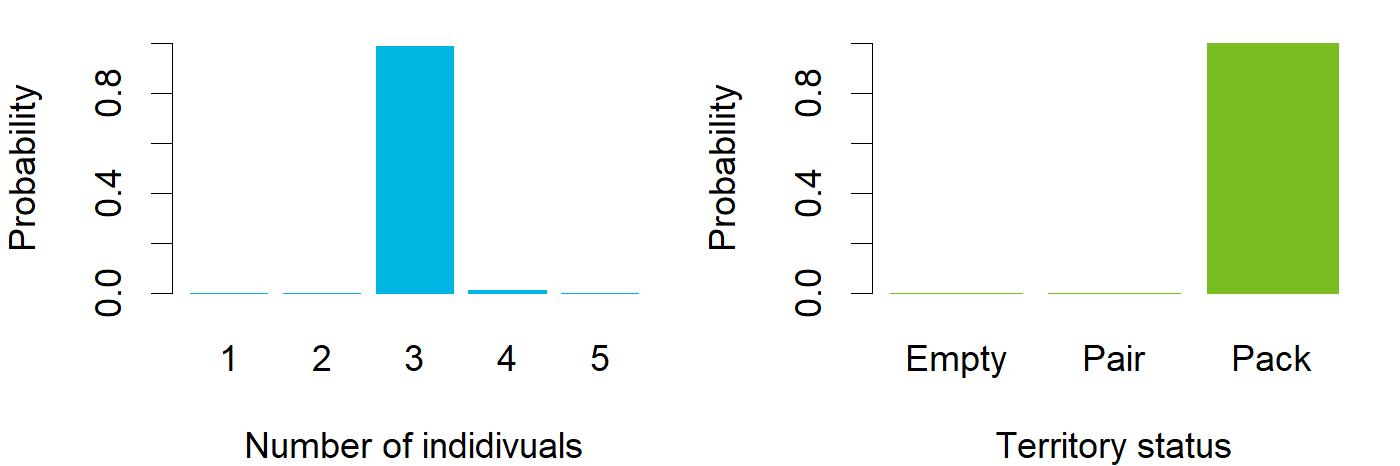

### Nov_ENG_Nivala.png

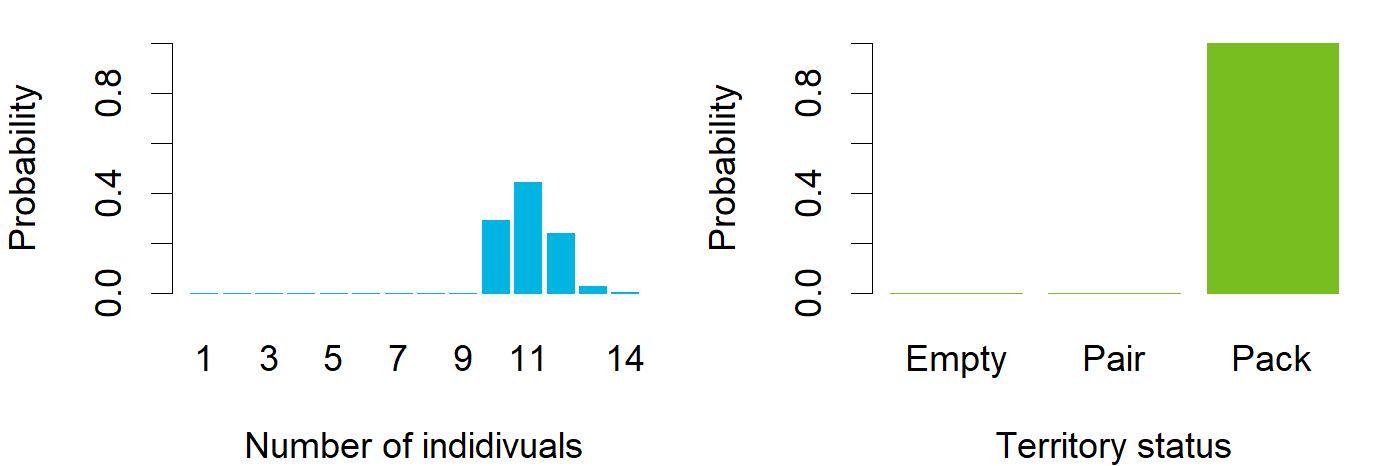

### Nov_ENG_Närvijoki.png

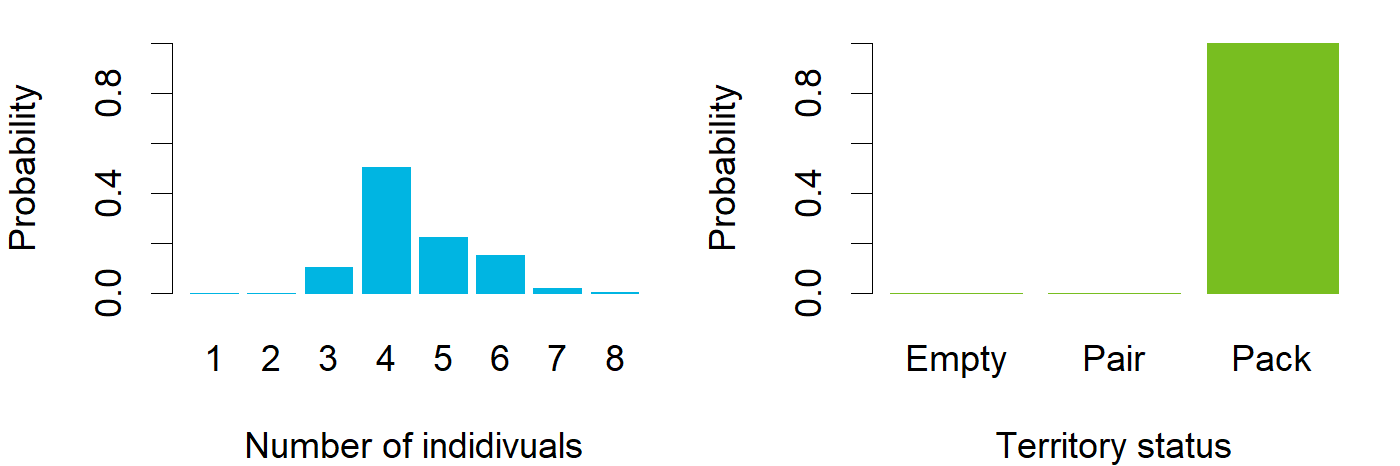

### Nov_ENG_Panja.png

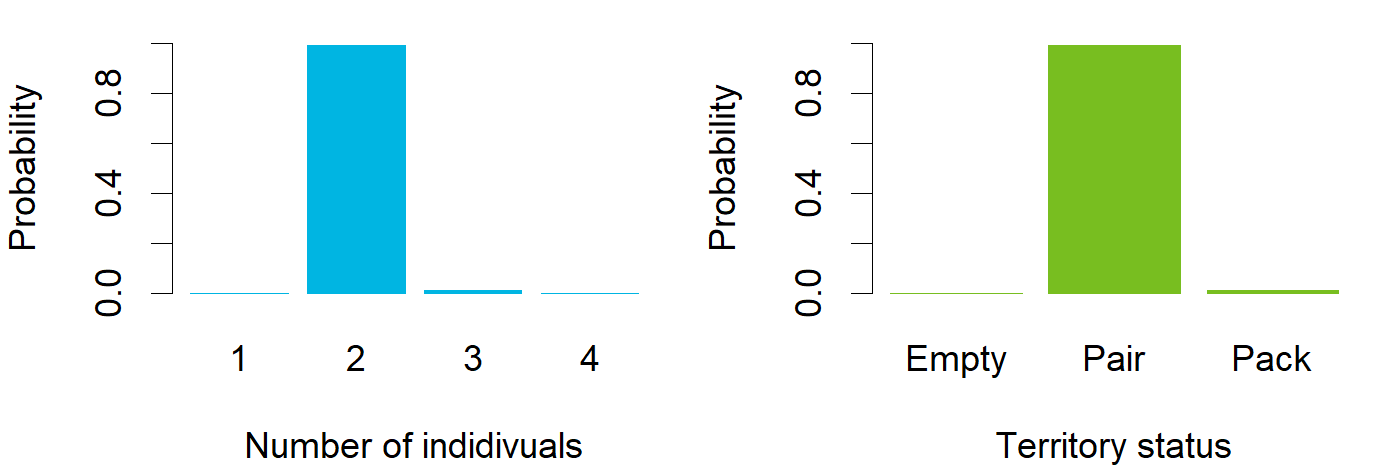

### Nov_ENG_Perho.png

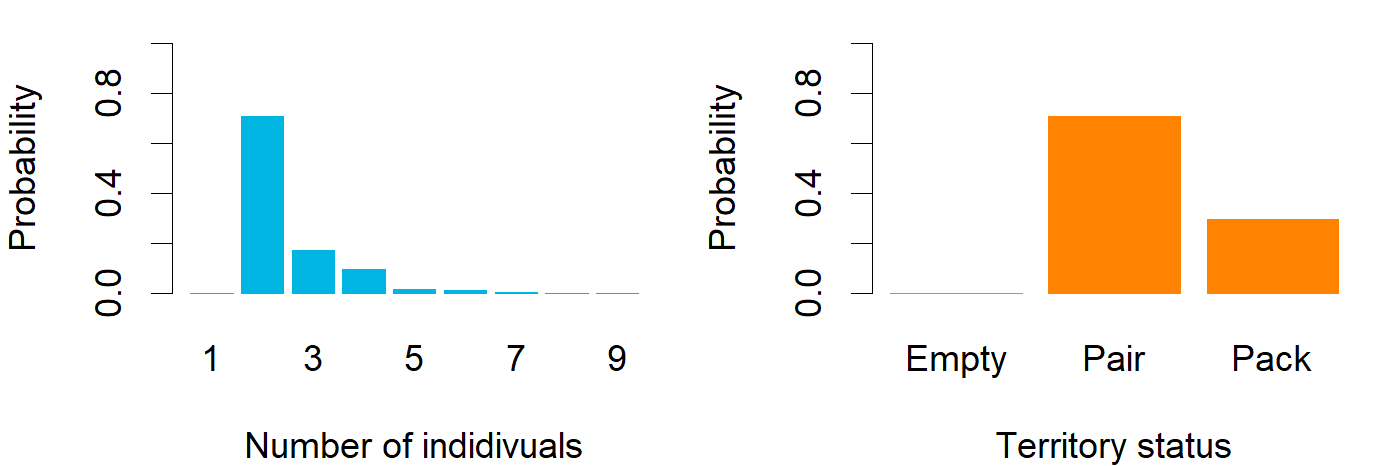

### Nov_ENG_Peurajärvi.png

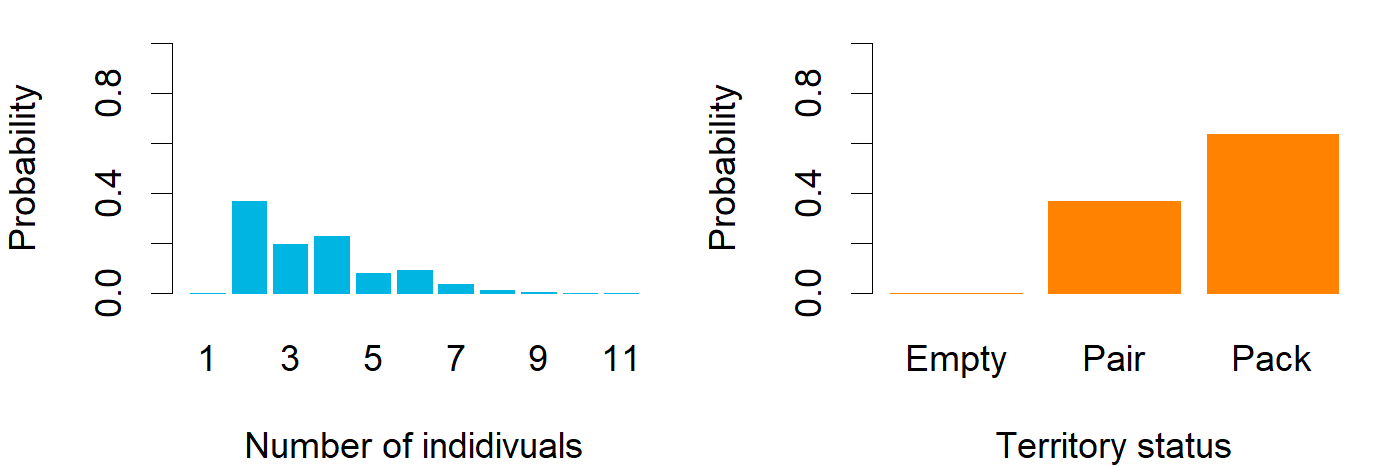

### Nov_ENG_Pirttivaara.png

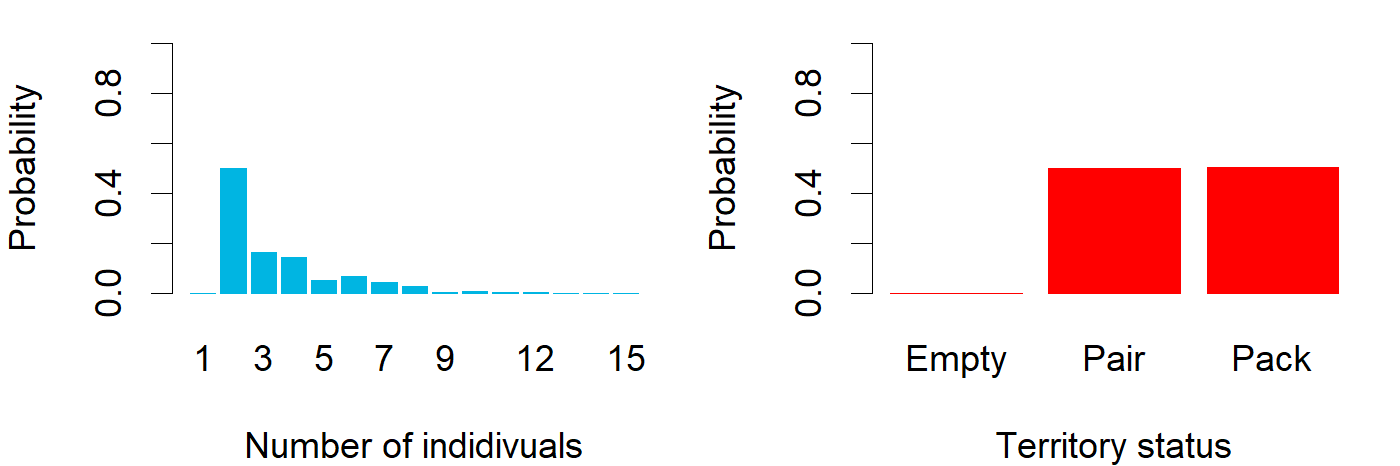

### Nov_ENG_Pulkkila.png

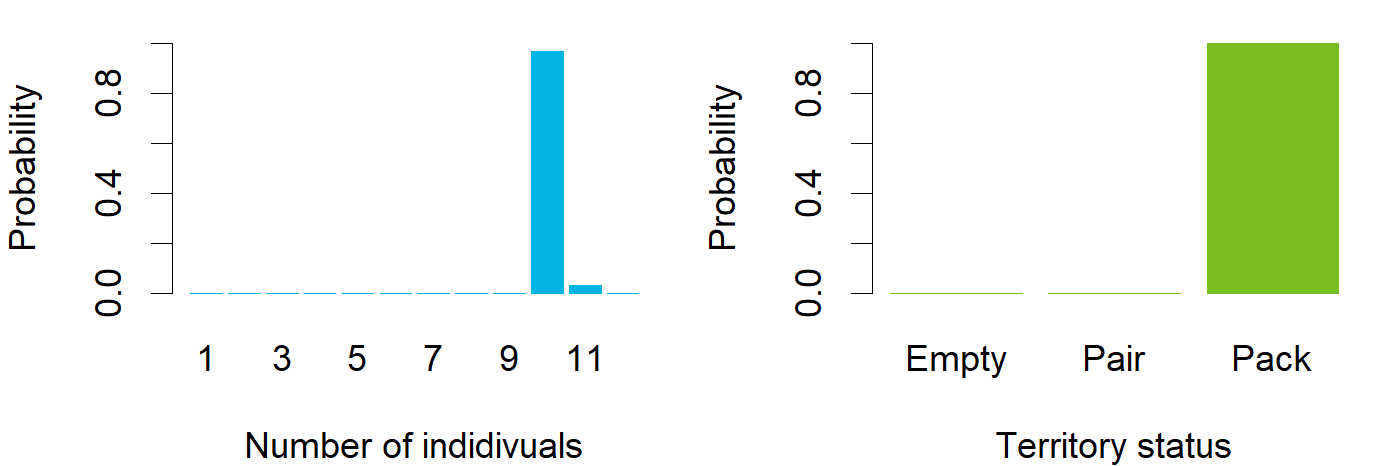

### Nov_ENG_Punkalaidun.png

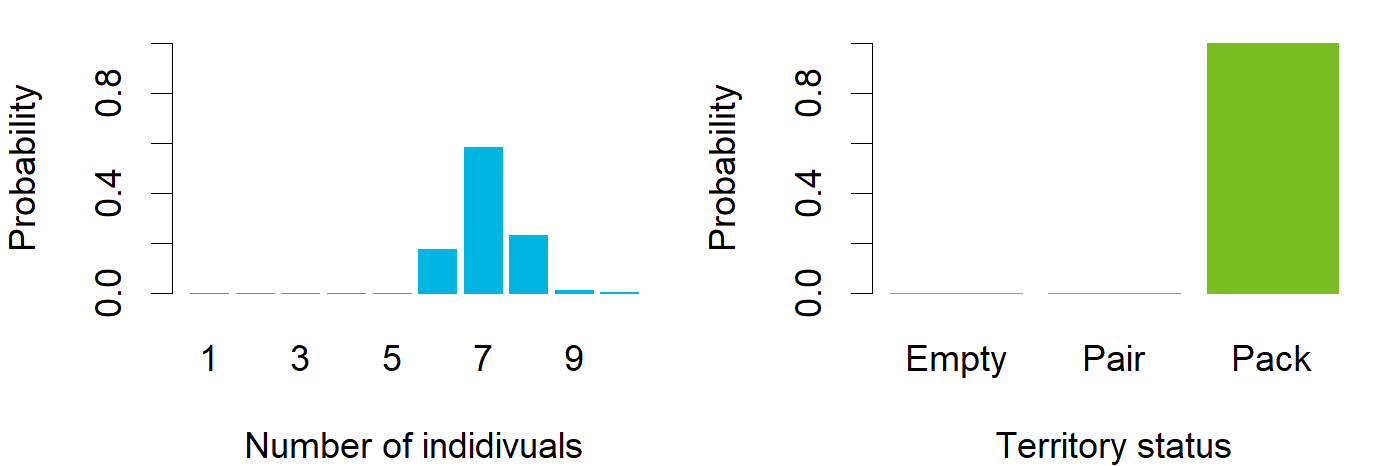

### Nov_ENG_Pyhäjoki.png

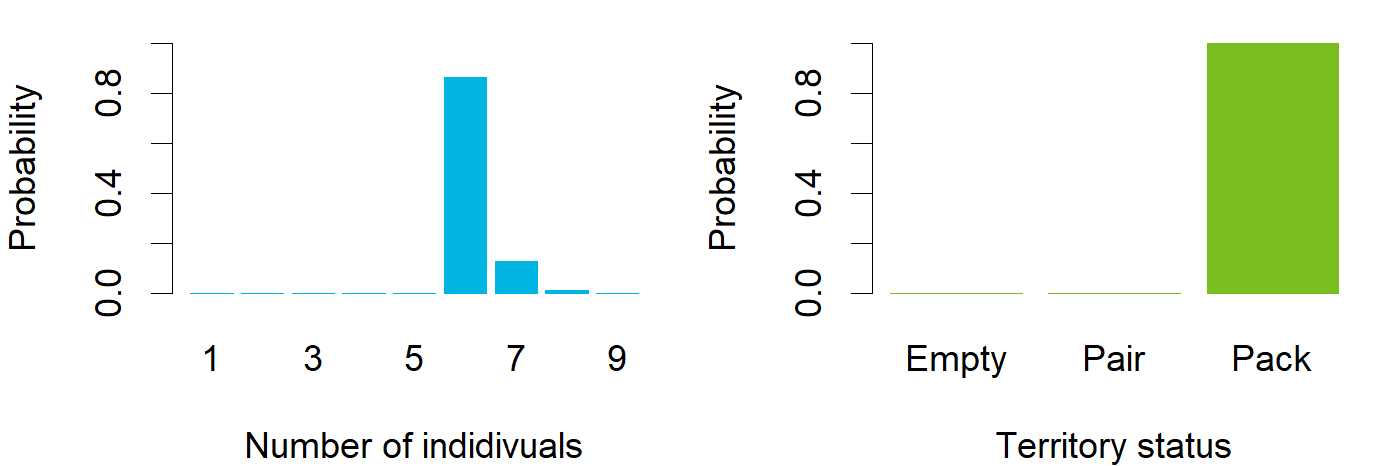

### Nov_ENG_Pöytyä.png

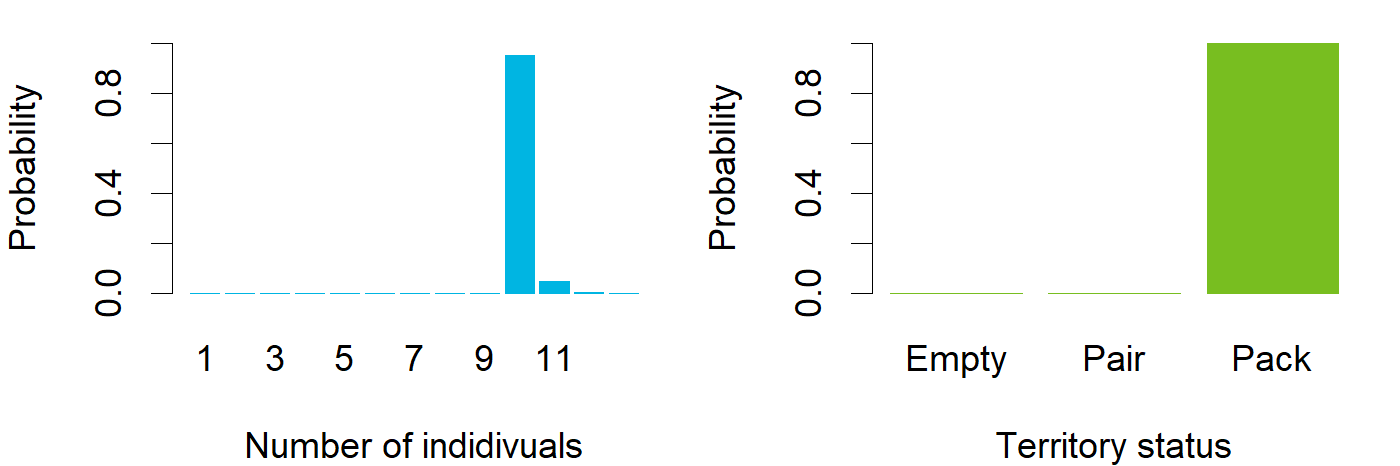

### Nov_ENG_Raasepori.png

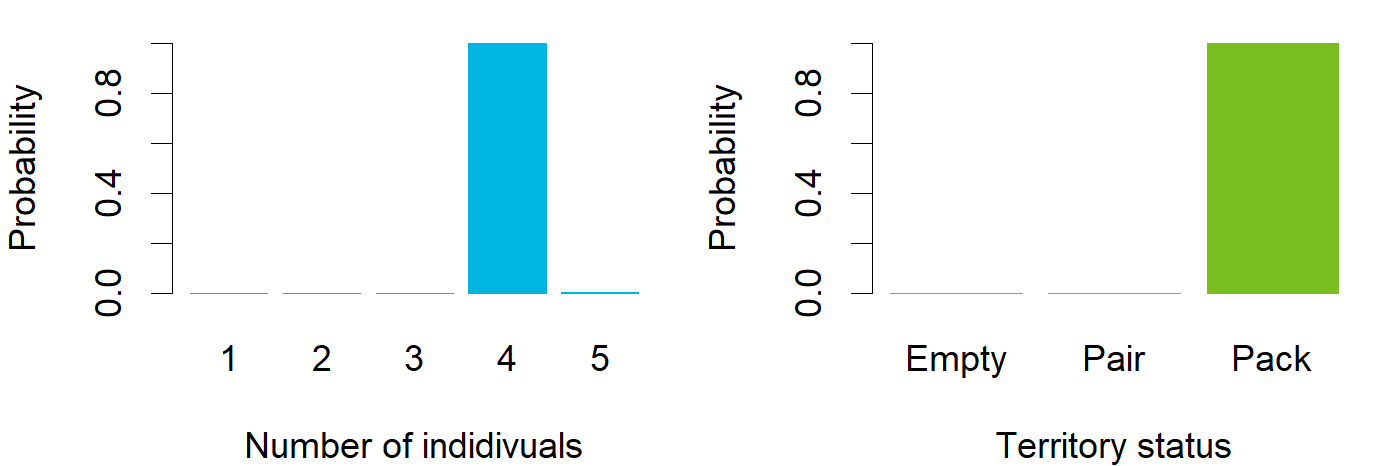

### Nov_ENG_Rantsila.png

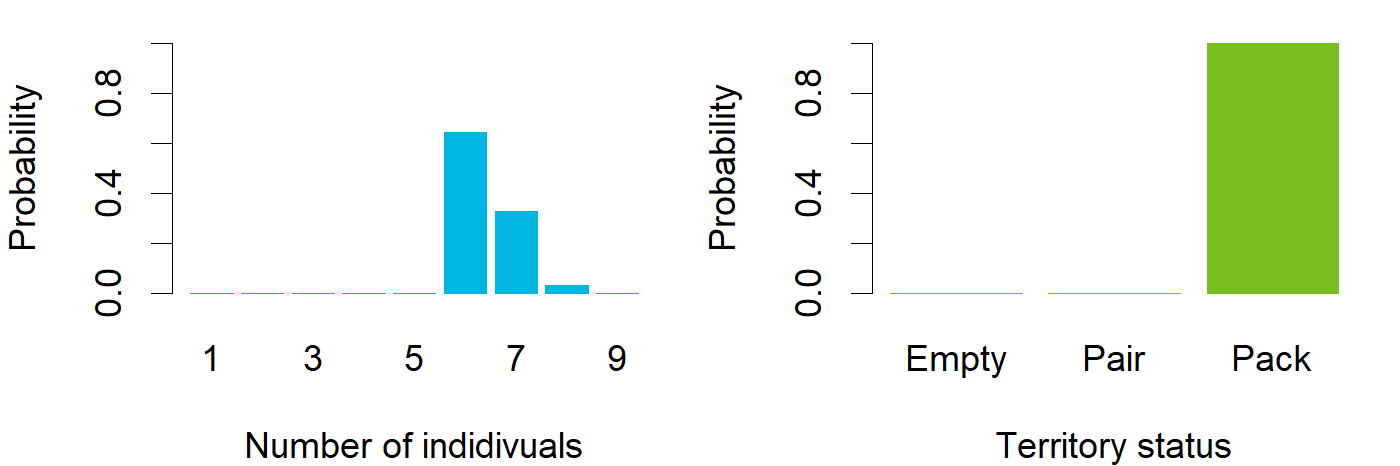
